# Isolation of an archaeon at the prokaryote-eukaryote interface

**DOI:** 10.1101/726976

**Authors:** Hiroyuki Imachi, Masaru K. Nobu, Nozomi Nakahara, Yuki Morono, Miyuki Ogawara, Yoshihiro Takaki, Yoshinori Takano, Katsuyuki Uematsu, Tetsuro Ikuta, Motoo Ito, Yohei Matsui, Masayuki Miyazaki, Kazuyoshi Murata, Yumi Saito, Sanae Sakai, Chihong Song, Eiji Tasumi, Yuko Yamanaka, Takashi Yamaguchi, Yoichi Kamagata, Hideyuki Tamaki, Ken Takai

## Abstract

The origin of eukaryotes remains enigmatic. Current data suggests that eukaryotes may have risen from an archaeal lineage known as “Asgard archaea”. Despite the eukaryote-like genomic features found in these archaea, the evolutionary transition from archaea to eukaryotes remains unclear due to the lack of cultured representatives and corresponding physiological insight. Here we report the decade-long isolation of a Lokiarchaeota-related Asgard archaeon from deep marine sediment. The archaeon, “*Candidatus* Prometheoarchaeum syntrophicum strain MK-D1”, is an anaerobic, extremely slow-growing, small cocci (∼550 nm), that degrades amino acids through syntrophy. Although eukaryote-like intracellular complexities have been proposed for Asgard archaea, the isolate has no visible organella-like structure. *Ca*. P. syntrophicum instead displays morphological complexity – unique long, and often, branching protrusions. Based on cultivation and genomics, we propose an “Entangle-Engulf-Enslave (E^3^) model” for eukaryogenesis through archaea-alphaproteobacteria symbiosis mediated by the physical complexities and metabolic dependency of the hosting archaeon.

---

How did the first eukaryotic cell emerge? So far, among various competing evolutionary models, the most widely accepted are the symbiogenetic models in which an archaeal host cell and an alphaproteobacterial endosymbiont merged to become the first eukaryotic cell^1–4^. Recent metagenomic discovery of Lokiarchaeota (and the Asgard archaea superphylum) led to the theory that eukaryotes originated from an archaeon closely related to Asgard archaea^5,6^. The Asgard archaea genomes encode a repertory of proteins hitherto only found in *Eukarya* (eukaryotic signature proteins – ESPs), including those involved in membrane trafficking, vesicle formation/transportation, ubiquitin and cytoskeleton formation^6^. Subsequent metagenomic studies have suggested that Asgard archaea have a wide variety of physiological properties, including hydrogen-dependent anaerobic autotrophy^7^, peptide or short-chain hydrocarbon-dependent organotrophy^8–11^and rhodopsin-based phototrophy^12,13^. A recent study suggests that an ancient Asgard archaea degraded organic substances and syntrophically handed off reducing equivalents (*e.g.*, hydrogen and electrons) to a bacterial partner, and further proposes a symbiogenetic model for the origin of eukaryotes based on this interaction^14^. However, at present, no single representative of the Asgard archaea has been cultivated and, thus, the physiology and cell biology of this clade remains unclear. In an effort to close this knowledge gap, we successfully isolated the first Asgard archaeon and here report the physiological characteristics, potentially key insights into the evolution of eukaryotes.

## Isolation of an Asgard archaeon

Setting out to isolate uncultivated deep marine sediment microorganisms, we engineered and operated a methane-fed continuous-flow bioreactor system for over 2000 days to enrich such organisms from anaerobic marine methane-seep sediments^15^(Supplementary Text 1). We successfully enriched many phylogenetically diverse yet-to-be cultured microorganisms, including Asgard archaea members (Loki-, Heimdall- and Odinarchaeota)^15^. For further enrichment and isolation, sample of the bioreactor community was inoculated in glass tubes with simple substrates and basal media. After approximately one year, we found faint cell turbidity in a Casamino acids-fed culture supplemented with four bacteria-suppressing antibiotics (*i.e.*, ampicillin, kanamycin, streptomycin, and vancomycin; Supplementary Text 2) and incubated at 20°C. Clone library-based small subunit (SSU) rRNA gene analysis revealed a simple community containing many *Halodesulfovibrio* and, excitingly, a small population of Lokiarchaeota (Extended Data Table 1). In pursuit of this archaeon, named strain MK-D1, we repeated the subcultures at the time when MK-D1 cell yield was maximized by means of quantitative PCR (qPCR) monitoring. Repeated subcultures gradually enriched the archaeon with extremely slow growth rate and low cell yield (Fig. 1a). The culture consistently had a 30–60 days of lag phase and required over 3 months to reach full growth with a yield of ∼10^5^ 16S rRNA gene copies/ml (Fig. 1a). The doubling time was estimated to be approximately 14–25 days. Variation of cultivation temperatures (Extended Data Fig. 1), and substrate combinations and concentrations did not significantly improve the lag phase, growth rate or cell yield (data not shown), while the static cultivation supplemented with 20 amino acids (AAs) and powdered milk resulted in the stable growth. For further characterization, we cultured the archaeon under the optimal conditions determined above.

**Fig. 1.**
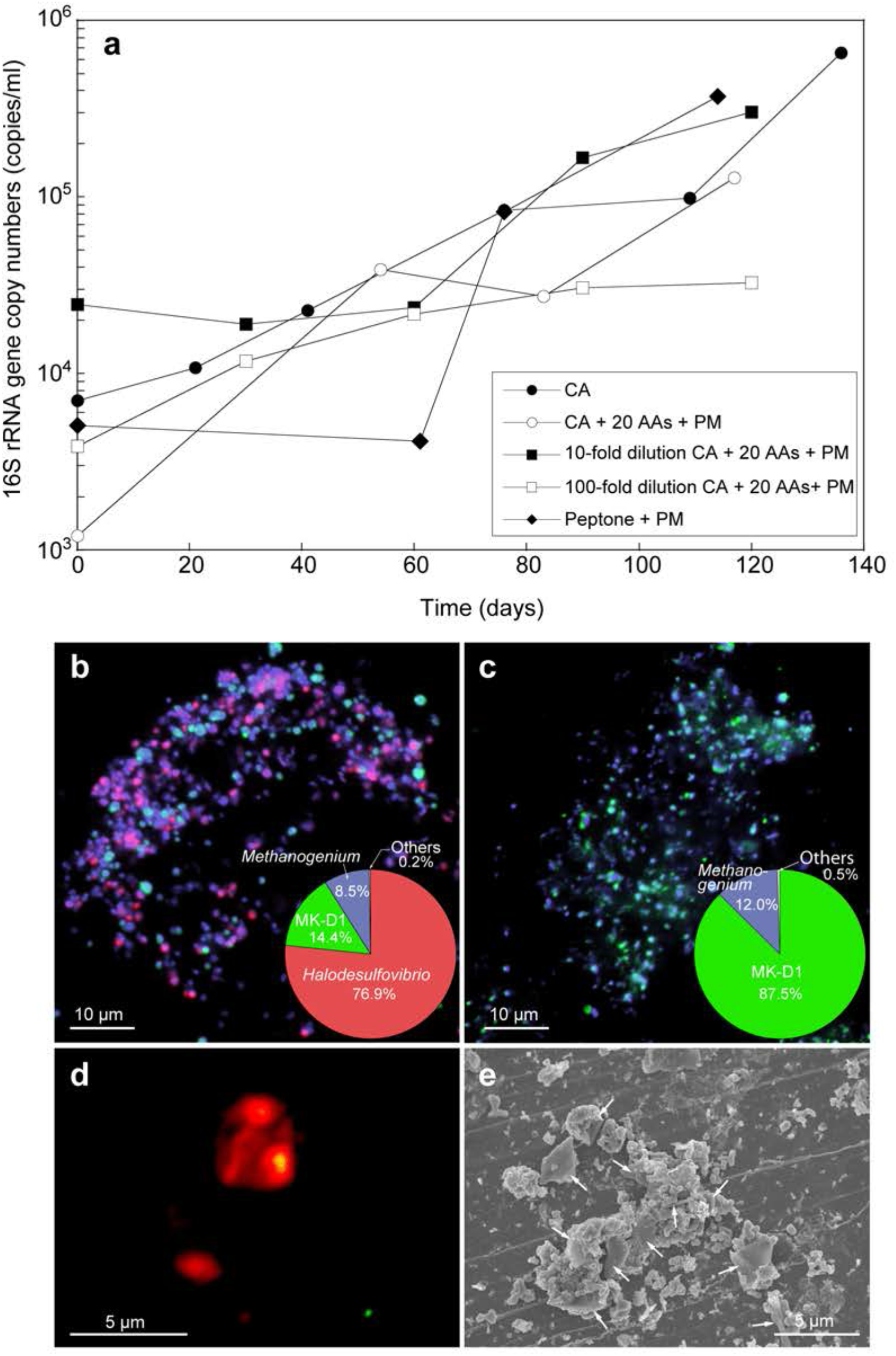
Growth curves and photomicrographs of the cultured Lokiarchaeota strain MK-Dl. **a**, Growth curves of MK-DI in anaerobic media supplemented with Casamino acids (CA; 0.05%, w/v) alone; CA with 20 amino acids (20 AAs; 0.1 mM of each) and powdered milk (PM; 0.1 %, w/v); or peptone (0.1 %, w/v) with PM. Results are also shown for cultures fed with 10- and 100-fold dilution of CA, 20 AAs, and PM. **b, c**, Fluorescence images of cells from enrichment cultures after eight (b) and eleven (c) transfers stained with DAPI (violet) and hybridized with nucleotide probes targeting MK-DI (green) and *Bacteria* (red). Pie charts show relative abundance of microbial populations based on SSU rRNA gene tag-sequencing (iTAG) analysis. **d**, A fluorescence image of cells from enrichment cultures after eleven transfers hybridized with nucleotide probes targeting MK-DI (green) and *Methanogenium* (red). **e**, SEM image of a highly purified co-culture of MK-DI and *Methanogenium.* White arrows indicate *Methanogenium* cells. The detailed iTAG-based community compositions of cultures corresponding to each of the images are shown in Supplementary Table S2.

After six transfers, MK-D1 reached 13% abundance in a tri-culture containing *Halodesulfovibrio* (85%) and *Methanogenium* (2%) (Extended Data Table 1). Fluorescence *in situ* hybridization (FISH) and scanning electron microscopic (SEM) observation revealed close physical association of the archaeon with the other microorganisms (Figs. 1b–e, Extended Data Fig. 2). Through metagenome-based exploration of this archaeon’s metabolic potential and stable isotope probing experiment, we discovered that MK-D1 can catabolize some AAs and peptides through syntrophic growth with *Halodesulfovibrio* and *Methanogenium* via interspecies hydrogen (and/or formate) transfer (Fig. 2 and Supplementary Table S1 and Fig. S1, details in latter section)^16^. Indeed, addition of hydrogen scavenger-inhibiting compounds (*i.e*., 10 mM molybdate and 10 mM 2-bromoethanesulfonate for sulfate-reducing *Halodesulfovibrio* and methanogenic *Methanogenium*, respectively) significantly impeded growth of MK-D1. Through subsequent transfers, we were able to eliminate the *Halodesulfovibrio* population, allowing us to obtain a pure co-culture of the archaeon and *Methanogenium* after a twelve-year journey – starting from deep-sea sediments to a bioreactor-based “pre-enrichment” and a final seven-year *in vitro* enrichment. We here propose the name “*Candidatus* Prometheoarchaeum syntrophicum strain MK-D1” for the isolated Lokiarchaeon.

**Fig. 2.**
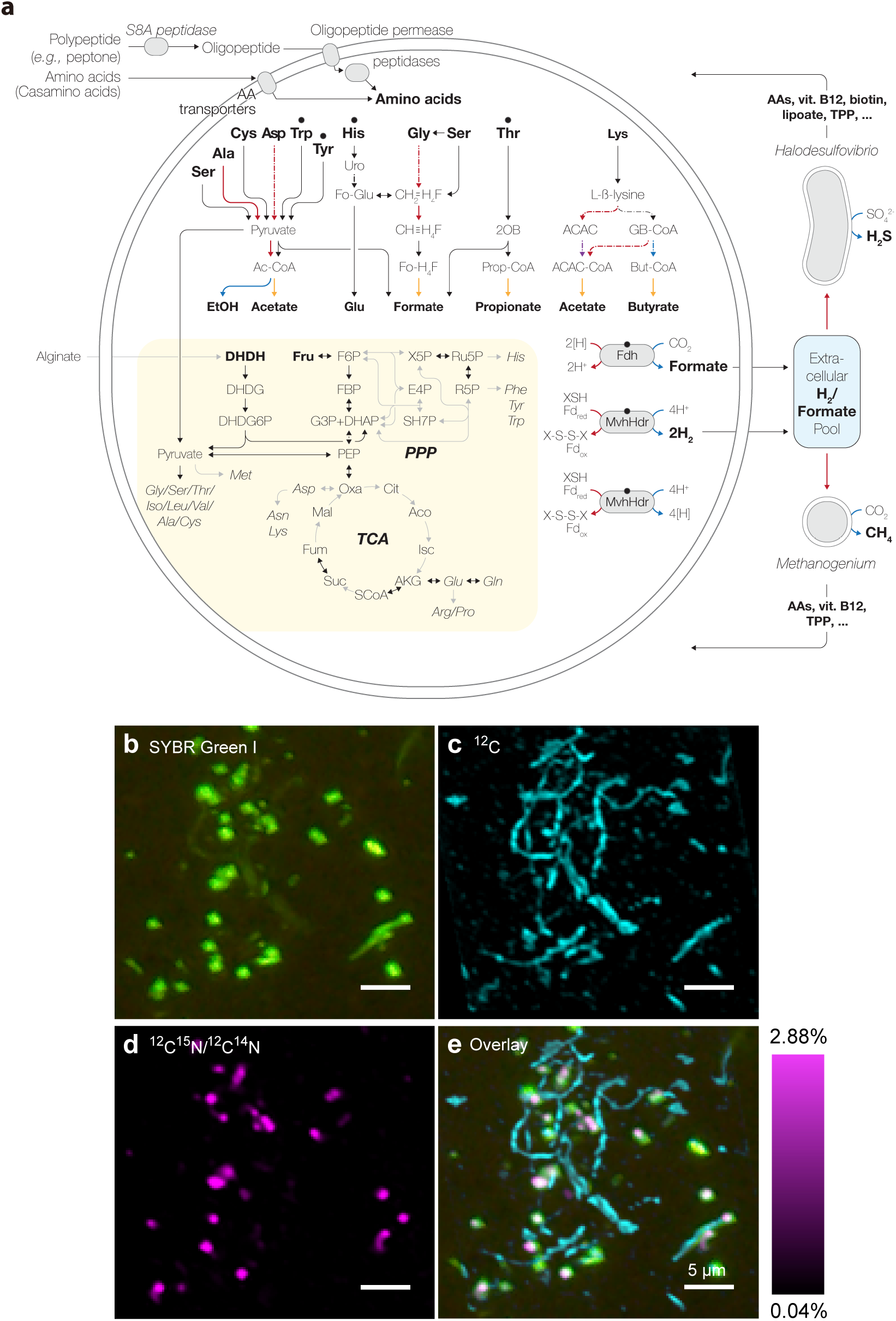
Syntrophic amino acid utilization of MK-D1. **a**, Genome-based metabolic reconstruction of MK-D1. Metabolic pathways identified (colored or black) and not identified (gray) are shown. For identified pathways, each step (solid line) or process (dotted) is marked by whether it is oxidative (red), reductive (blue), ATP-yielding (orange), or ATP-consuming (purple). Wavy arrows indicate exchange of compounds: formate, H_2_, AAs, vitamin B_12_, biotin, lipoate, and thiamine pyrophosphate (tpp) which are predicted to be metabolized or synthesized by the partnering *Halodesulfovibrio* and/or Methanogenium. Biosynthetic pathways are indicated with a yellow background. Metatranscriptomics-detected AA-catabolizing pathways are indicated (black dots above AAs). Abbreviations: 4,5-dihydroxy-2,6-dioxohexanoate (DHDH), 2-dehydro-3-deoxy -D-gluconate (DHDG), 3-dehydro-3-deoxy-D-gluconate 6-phosphate (DHDG6P), acetyl-CoA (Ac-CoA), urocanate (uro), formyl glutamate (Fo-Glu), methylene-tetrahydrofolate (CH_3_=H_4_F), methenyl-tetrahydrofolate (CH≡H_4_F), formyl-tetrahydrofolate (Fo-H_4_F), 2-oxobutyrate (2OB), propionyl-CoA (Prop-CoA), acetoacetate (ACAC), gamma-amino-butyryl-CoA (GB-CoA), butyryl-CoA (But-CoA), ferredoxin (Fd), thiol/disulfide pair (XSH/X-S-S-X), tricarboxylic acid (TCA) cycle, and pentose-phosphate pathway (PPP). **b-e**, NanoSIMS analysis of a highly purified MK-DI culture incubated with ^13^C- and ^15^N-labeled AA mixture. **b**, Green fluorescent micrograph of SYBR Green **I**-stained cells. Aggregates are MK-DI, and filamentous cells are *Methanobacterium* sp. strain MO-MB1 (fluorescence can be weak due to high rigidity and low permeability of cell membrane [Supplementary Fig. S1]). **c**, NanoSIMS ion image of ^12^C (cyan). **d**, NanoSIMS ion image of ^12^C ^l5^N/^12^C^l4^N (magenta). e, Overlay image of **b-d**. The right-hand scale bar indicates the relative abundance of ^15^N expressed as ^15^N/^14^N. The iTAG analysis of the imaged culture is shown in Supplementary Table S2.

## Cell biology and physiology of MK-D1

We further characterized strain MK-D1 using the highly purified cultures and pure co-cultures. Microscopic observations showed that the cells are small cocci, ca. 300-750 nm in diameter (average 550 nm, n=15), and generally form aggregates surrounded with extracellular polysaccharide (EPS)-like materials (Fig. 3a, b and Extended Data Fig. 2), consistent with previous observations using FISH^15,17^. Dividing cells had less EPS-like materials and a ring-like structure around the middle of cells (Fig. 3c and Extended Data Fig. 2). Cryo-electron and transmission electron microscopic observations revealed that the cells contain no visible organelle-like inclusions (Fig. 3d–f, Extended Data Fig. 2 and Supplementary Movies S1–S3). The cells produce membrane vesicles (MVs; 50–280 nm in diameter) (Fig. 3d–f and Extended Data Fig. 2) and chains of blebs (Fig. 3c and Extended Data Fig. 2e). The cells also form unique membrane-based protrusions with a diameter of about 80–100 nm and various lengths (Fig. 3g–i and Extended Data Fig. 2). Some protrusions remarkably display complex branching, unlike known archaeal protrusions^18^. These protrusions were especially abundant after late exponential growth phase. Lipid composition analysis of the MK-D1 and *Methanogenium* co-culture revealed typical archaeal signatures – a C_20_-phytane and C_40_-biphytanes (BPs) with 0–2 cyclopentane rings (Fig. 3j). Considering the lipid data obtained from a reference *Methanogenium* isolate (99.3% 16S rRNA gene identity; Supplementary Fig. S3), MK-D1 probably contains C_20_-phytane and C_40_-BPs with 0–2 rings.

**Fig. 3.**
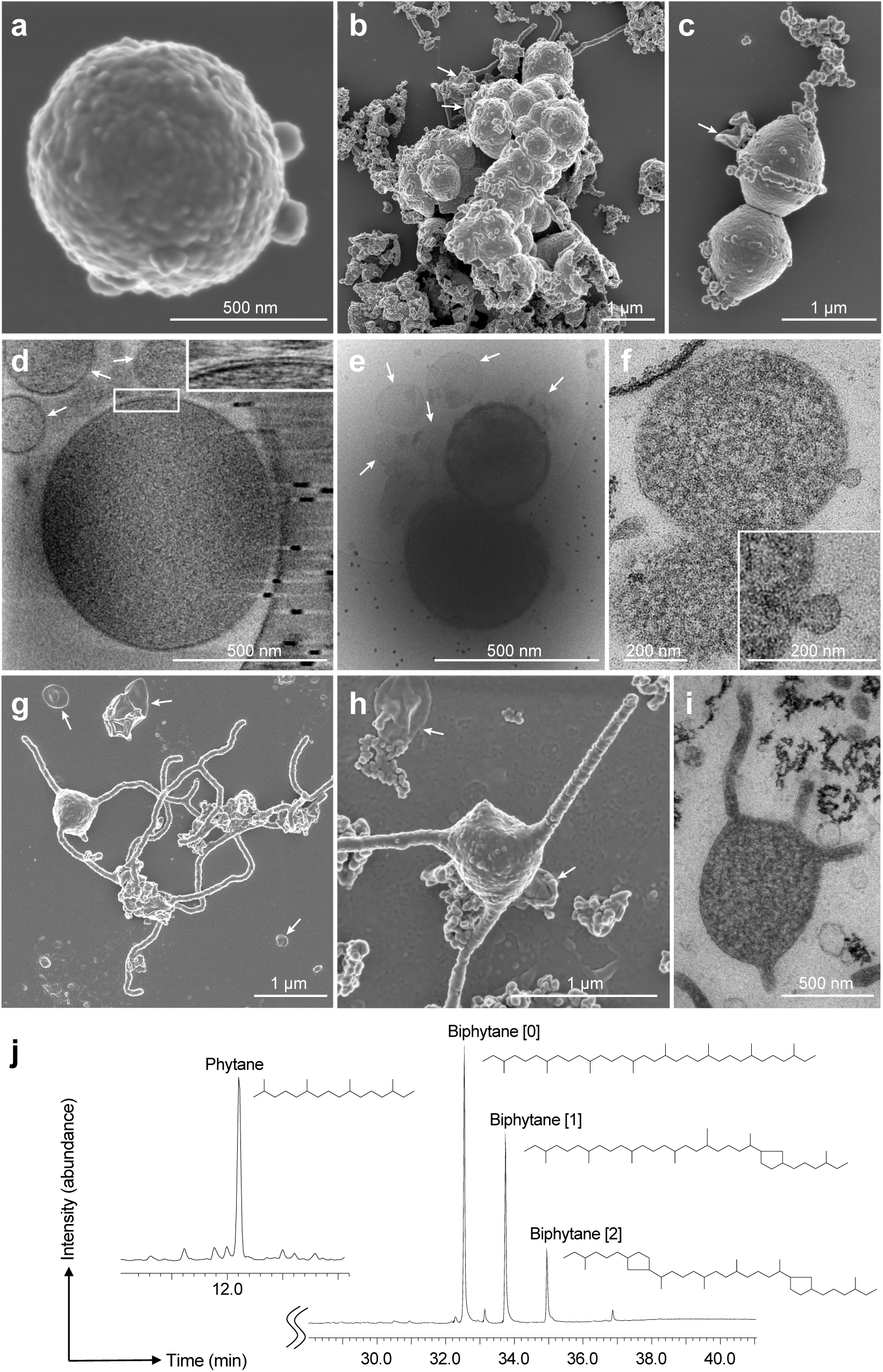
Microscopic characterization and lipid composition of MK-D1. **a-c**, SEM images of MK-Dl. Single cell (**a**), aggregated cells covered with EPS-like materials (**b**), and a dividing cell with polar chains of blebs (**c**). **d**, Cryo-electron tomography image of MK-Dl. The upper-right inset image shows a close-up of the boxed area for showing cell envelope structure. **e**, Cryo-electron microscopy (EM) image of large MVs attached and surrounding MK-D1 cells. **f**, Ultrathin section of an MK-D1 cell and an MV. The lower-right inset image shows a magnified view of the MV. **g**, **h**, SEM images of MK-D1 cells producing long branching (**g**) and straight **(h)** membrane protrusions. **i**, Ultrathin section of a MK-D1 cell with protrusions. **j**, A total ion chromatogram of mass spectrometry for lipids extracted from a highly purified MK-D1 culture. The chemical structures of isoprenoid lipids are also shown (see also Supplementary Fig. S2). The experiment were repeated twice and gave similar results. White arrows in the images indicate large MVs. Detailed iTAG-based community compositions of the cultures are shown in Supplementary Table S2.

MK-D1 can degrade AAs anaerobically, as confirmed by monitoring AAs depletion during the growth of pure co-cultures (Extended Data Fig. 3). We further verify AA utilization by quantifying the uptake of a mixture of ^13^C- and ^15^N-labeled AAs through nanometer-scale secondary ion mass spectrometry (NanoSIMS) (Fig. 2b–e). Cell aggregates of MK-D1 incorporated more nitrogen than carbon, suggesting that the possible mixotrophy. Interestingly, the ^13^C-labeling of methane and carbon dioxide varied depending on the methanogenic partner, indicating that MK-D1 produces both hydrogen and formate from AAs for interspecies electron transfer (Extended Data Table 2, see later section). Indeed, addition of high concentrations of hydrogen or formate completely suppressed growth of MK-D1 (Extended Data Table 3). The syntrophic partner was replaceable – MK-D1 could also grow syntrophically with *Methanobacterium* sp. strain MO-MB1^19^instead of *Methanogenium*, which was originally co-enriched with MK-D1 (Fig. 2b–e). Although 14 different culture conditions were applied, none of substances (*e.g.*, sugars, electron acceptors, and cell building blocks) enhanced the cell yield, implying specialization to degradation of AAs or peptides (Extended Data Table 3).

### Etymology

*Prometheoarchaeum, Prometheus* (Greek): a Greek god who shaped man out of mud and gave them the ability to create fire; *archaeum* from *archaea* (Greek): an ancient life. The genus name is an analogy between this organism’s evolutionary relationship with the origin of eukaryotes and the involvement of Prometheus in man’s origin from sediments and acquisition of an unprecedented oxygen-driven energy-harnessing ability. The species name, *syntrophicum, syn* (Greek): together with; *trephein* (Greek) nourish; *icus* (Latin) pertaining to. The species name referred to syntrophic substrate utilization property of this strain.

### Locality

Isolated from deep-sea methane seep sediment of the Nankai Trough at 2533 m water depth, off Kumano area, Japan.

### Diagnosis

Anaerobic, AA-oxidizing archaeon, small cocci, ca. 550 nm in diameter, syntrophically grows with hydrogen- and formate-utilizing microorganisms. It produces MVs, chains of blebs, and membrane-based protrusions.

## Reconstruction of extant and ancestral features

MK-D1 encodes genes for degradation of 10 AAs and reductive generation of H_2_ and formate for electron disposal. Most of the identified AA-catabolizing pathways only recover energy through degradation of a 2-oxoacid intermediate (*i.e.*, pyruvate or 2-oxobutyrate; Fig. 2a). MK-D1 can degrade 2-oxoacids hydrolytically (2-oxoacid--formate lyases) or oxidatively (2-oxoacid:ferredoxin oxidoreductases) to yield acyl-CoA intermediates that can be further hydrolyzed for ATP generation. The hydrolytic and oxidative paths release the AA carboxylate group as formate and CO_2_ respectively. For the former, formate can be directly handed off to a partnering methanogenic archaea or sulfate-reducing bacteria (SRB). For the latter, reduced ferredoxin generated from 2-oxoacid oxidation can drive reduction of H^+^ to H_2_ (electron-confurcating NiFe hydrogenase MvhADG-HdrABC) or CO_2_ to formate (formate dehydrogenase FdhA) for interspecies electron transfer. This suggests that MK-D1 has two approaches for syntrophic interaction. A ^13^C-AA-fed co-culture of MK-D1 with *Methanobacterium* generated ^13^C-enriched CH_4_ (Extended Data Table 2), indicating syntrophy mediated by the hydrolytic path (*i.e.*, AA-derived ^13^C-formate transferred to partner, oxidized to ^13^CO_2_, and further reduced to ^13^CH_4_). On the other hand, a ^13^C-AA-fed tri-culture of MK-D1 with *Halodesulfovibrio* and *Methanogenium* generated ^13^C-enriched CO_2_, suggesting the oxidative path (*i.e.*, AA-derived ^13^CO_2_ released and mixed with ^12^C-bicarbonate pool in mineral medium). Thus, we confirm that MK-D1 can switch between syntrophic interaction via 2-oxoacid hydrolysis and oxidation depending on the partner(s).

The evolutionary relationship between archaea and eukaryotes has been under debate, hinging on the incompleteness and contamination associated with metagenome-derived genomes and variation in results depending on tree construction protocols^20–23^. By isolating strain MK-D1, we were able to obtain a closed genome (Supplementary Table S1 and Fig. S1) and construct a ribosomal protein-based phylogenomic tree that shows clear phylogenetic sistering between MK-D1 and *Eukarya* (Fig. 4a and Supplementary Tables S4 and S5, and Fig. S4). Thus, strain MK-D1 represents the closest cultured archaeal relative of eukaryotes. We confirmed the presence of many ESPs identified in related Asgard archaea (Supplementary Fig. S5) and obtained the first RNA-based evidence for expression of such genes (Supplementary Table S6).

**Fig. 4.**
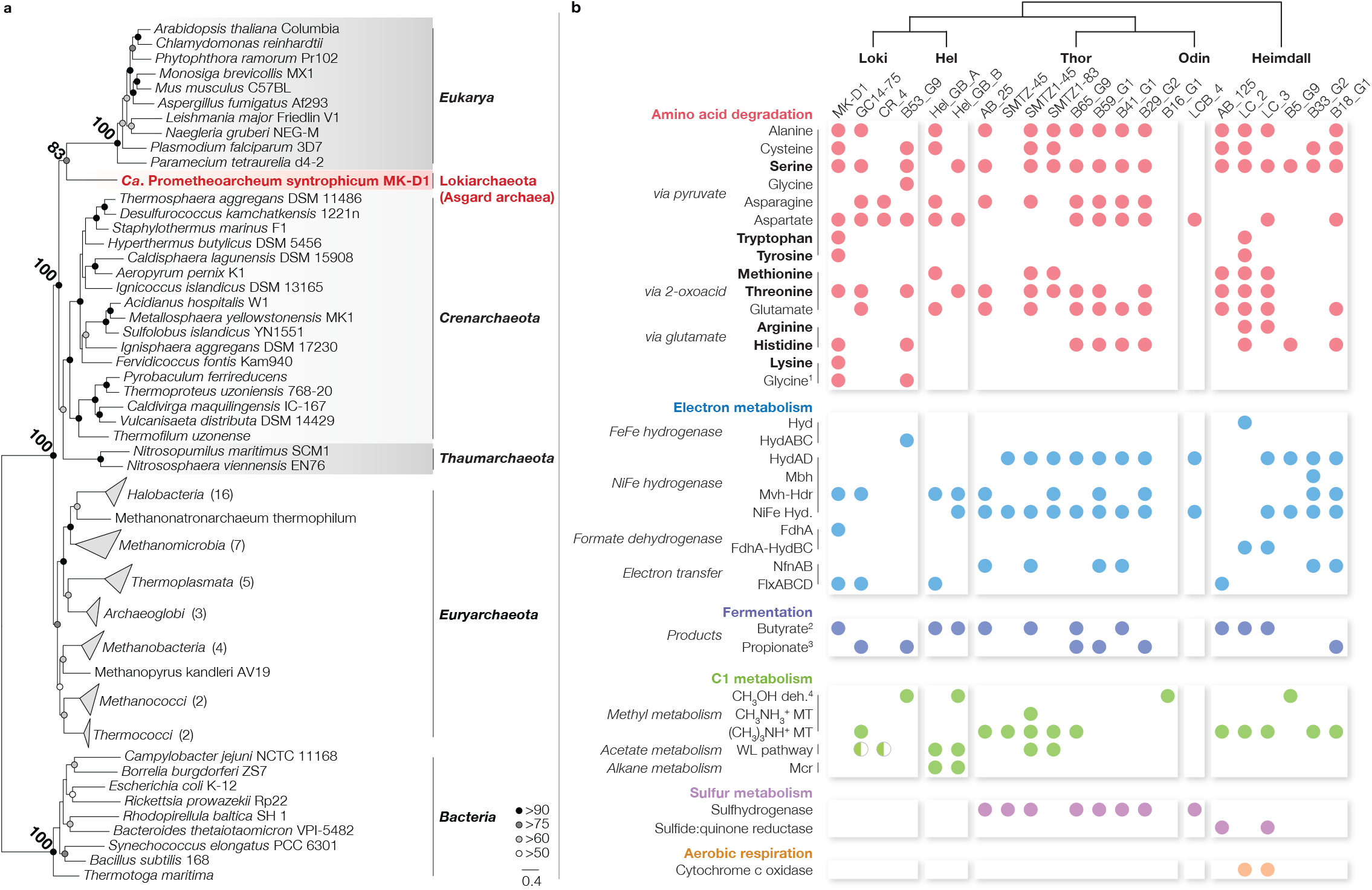
phylogeny of MK-D1 and catabolic features of Asgard archaea. **a**, Phylogenomic tree of MK-D1 and select cultured karyotes, and bacteria based on 31 ribosomal proteins conserved across the three domains (Supplementary Table S4). protein sequences were collected from MK-D1 and other representative cultured organisms (Supplementary Table S5) individually using MAFFT (--linsi). After removing all-gap positions and concatenation, the maximum likelihood tree ucted using RAxML-ng (fixed empirical substitution matrix [LG], 4 discrete GAMMA categories, empirical AA frequen-0 bootstrap replicates). Bootstrap values around critical branching points are also shown. **b**, The presence/absence of AA, electron metabolism, fermentation, C1 metabolism, sulfur metabolism, and aerobic respiration in individual genomes complete pathway – full circle; mostly complete pathway – half circle). For AA metabolism, pathways that are exclusive-catabolism/degradation are bolded. ^1^ Glycine metabolism through pyruvate (above) or formate (below). ^2^ Butyrate metab-versible (fermentation or beta oxidation), but the butyryl-CoA dehydrogenases tend to be associated with EtfAB in the uggesting formation of an electron-confurcating complex for butyrate fermentation. ^3^ Determined by presence of methyl-oA decarboxylase, biotin carboxyl carrier protein, and pyruvate carboxylase. Propionate metabolism is also reversible, but archaea member encodes the full set of genes necessary for syntrophic propionate degradation. ^4^ Alcohol dehydrogenases iverse substrate specificities. Abbreviations: monomeric FeFe hydrogenase (Hyd), trimeric electron-confurcating FeFe e (HydABC), reversible NADPH-dependent NiFe hydrogenase (HydAD), reversible heterodisulfide-dependent nfurcating hydrogenase (Mvh-Hdr), other NiFe hydrogenases (NiFe Hyd.), formate dehydrogenase (FdhA), putative nfurcating formate dehydrogenase (FdhA-HydBC), NADH-dependent NADPH:ferredoxin oxidoreductase (NfnAB), fide- and flavin-dependent oxidoreductase of unknown function (FlxABCD), tetrahydromethanopterin methyltransferase (MT) wood--Ljungdahl (WL), and methyl-CoM reductase (Mcr).

Given the phylogenetic relationship of MK-D1, other Asgard archaea, and eukaryotes, estimating the physiological traits of the last Asgard archaea common ancestor is of utmost importance. Comparative genomics of MK-D1 and published metagenome-assembled genomes of Asgard archaea revealed that most of the members encode AA-catabolizing pathways, reversible NiFe hydrogenases (MvhADG-HdrABC^24^ and/or HydAD^25^) (Fig. 4b), and restricted biosynthetic capacities (*i.e.*, AA and vitamin synthesis; Extended Data Fig. 4), indicating H_2_-evolving AA degradation and partner dependence may be a common feature across the superphylum. Like MK-D1, other Asgard archaea members of Lokiarchaeota, Helarchaeota, and Heimdallarchaeota may be capable of syntrophic AA degradation given that they encode an electron transfer complex FlxABCD-HdrABC associated with syntrophic bacteria^26^ or formate dehydrogenases. Many lineages also possess genes for alternative electron disposal through fermentation – *i.e*., reduction of pyruvate and acetyl-CoA to propionate and butyrate correspondingly (see Fig. 4b for details). Many lineages also encode the potential for other metabolisms – mono/tri-methylamine-driven homoacetogenesis and coupled H_2_/S^0^ metabolism in Thorarchaeota; H_2_S metabolism in Heimdallarchaeota; and, as pointed out by pioneering studies, Wood-Ljungdahl pathway in several genomes^7,8,10,14^; alkane metabolism in Helarchaeota^11^; and aerobic respiration in Heimdallarchaeota^6^. Although these metabolisms are highly unique and ecologically important, they are either only sporadically present or confined to specific phylum-level lineages. To identify potentially ancestral features, we searched for catabolic genes conserved across phylum-level lineages including Heimdallarchaeota (the most deep-branching Asgard archaea) that form monophyletic clusters in phylogenetic analyses. We found key catabolic genes for serine, threonine, and histidine degradation (serine/threonine dehydratase and urocanate hydratase; Supplementary Figs. S6 and S7), butyrate fermentation (3-ketoacyl-CoA thiolase and fatty-acid--CoA ligase; Supplementary Figs. S8 and S9), and propionate fermentation (succinate dehydrogenase flavoprotein subunit, methylmalonyl-CoA transcarboxylase-associated biotin ligase, and biotin carboxyl carrier protein; Supplementary Figs. S10–S12). Given the physiology of the isolated MK-D1, presence of AA catabolism, H_2_ metabolism, and lack of biosynthetic pathways in nearly all extant Asgard archaea lineages, and conservation of the above metabolisms, we propose that the last Asgard archaea common ancestor was an AA-degrading anaerobe producing H_2_ and fatty acids as byproducts that acquired ATP primarily from substrate-level phosphorylation from catabolizing 2-oxoacid intermediates and depended on metabolic partners, though we do not reject the possibility of other additional lifestyles.

## Proposal of new eukaryogenesis model

We demonstrate that Asgard archaea are capable of syntrophic AA degradation and identify related metabolic features conserved across the superphylum. This provides tangible evidence for the recent proposal that the ancestral Asgard archaeon was a syntrophic organotroph based on the prevalence of NiFe hydrogenases and hydrogenogenic organotrophy across the superphylum^14^. In Earth’s early ocean, partners were likely methanogenic archaea rather than SRB due to low ocean sulfate concentrations prior to the Great Oxidation Event (G.O.E.; 2.7 Ga∼)^27^ (Fig. 5a). As the ocean and atmosphere became oxygenated, marine sulfate concentrations rose^28^ and syntrophy likely shifted to interaction with SRB as observed in this study, which is more thermodynamically favorable^29^. During the G.O.E., cyanobacterial activity (and concomitant marine organic matter production) increased, leading to transfer of excess organic matter from the photic zone to marine sediments^30,31^. The ancient anaerobic Asgard archaea could have taken one of two paths for survival and adaptation: to remain confined in strictly anaerobic habitats or to advance towards the anoxic-oxic interface with greater substrate and “energy” availability. The archaeon at the last Archaea-Eukarya division (*i.e.*, Heimdallarchaeota-Eukarya ancestor) likely preferentially grew closer to the sulfate- and organics-rich anoxic-oxic interface environments (SRB could have continued syntrophic interaction at the anoxic-oxic interface as many extant SRB are aerotolerant and can perform sulfate reduction in the presence of O_2_^32^). However, to further adapt to higher O_2_ concentrations and also compete with facultatively aerobic organotrophs, acquisition of the capacity for O_2_ utilization would have been necessary.

**Fig. 5.**
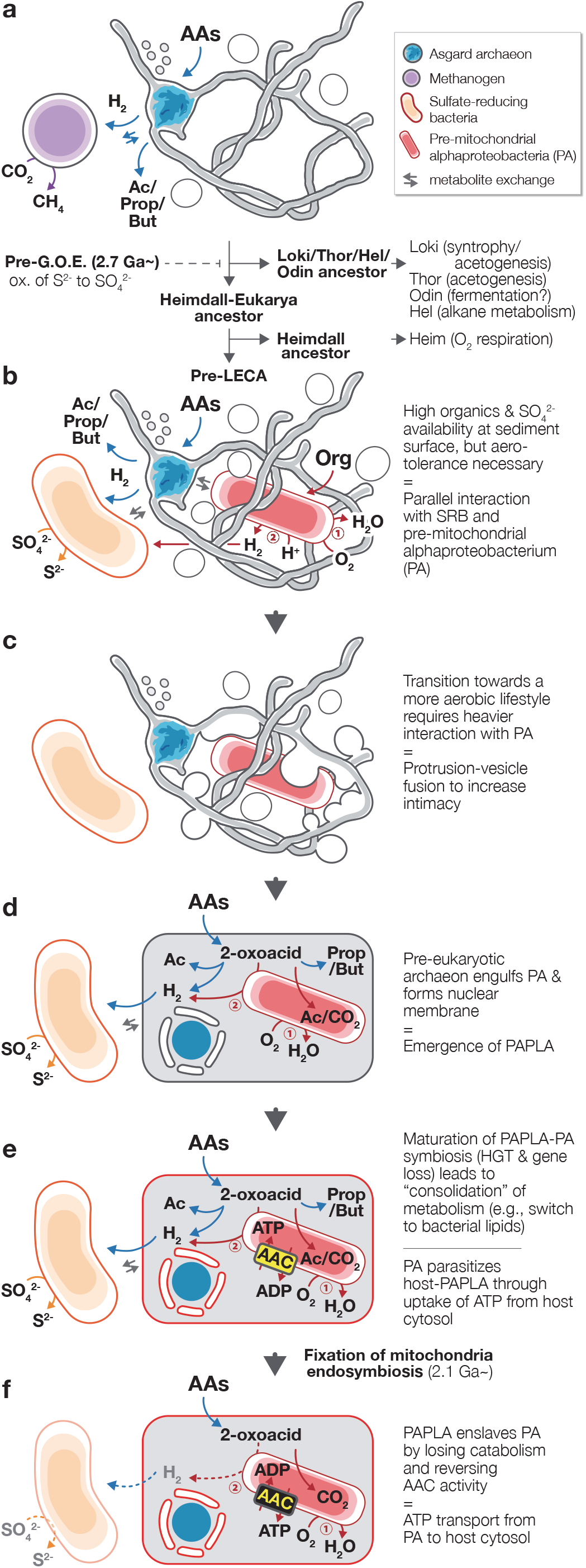
A new evolutionary model “Entange-Engulf-En-slave (E^3^)” for eukaryogenesis. **a**, Syntrophic/fermentative Asgard archaea ancestor likely degraded AAs to short-chain fatty acids and H2. With the rising of Earth’s O2 concentrations due to oxygenic photosynthesis (around the Great Oxidation Event [G.O.E.]), Asgard archaea diverged towards specialized anaerobic niches (Lokiarchaeota [Loki], Thorarchaeota [Thor], Odinarchaeota [Odin], and Helarchaeota [Hel]) and aerobiosis (Heimdallarchaeota [Heimdall] and Eukarya). **b**, To thrive at the oxic-anoxic interface with high organic and sulfate availability, pre-LECA (last eukaryote common ancestor) archaeon syntrophically interacted with H2-scavenging SRB (orange) and O2-scavenging organotrophic prmito-chondrial alphaproteobacterium (PA; red). PA could likely degrade organics (1) aerobically or (2) anaerobically in interaction with SRB. Given the restricted biosynthetic capacities of all extant Asgard archaea, the pre-LECA archaeon necessitated metabolite exchange with SRB and PA. **c**, Protrusions and MVs tangle with PA and enhance physical interaction; protrusion-MV fusion mediates further intimate interactions, and ultimately leads to PA engulfment. This mechanism for engulfment allows for formation of a nucleoid-bounding membrane topologically similar to the eukaryote nuclear membrane. **d**, After engulfment, the pre-LECA archaeon and PA can continue the interaction shown in (**b**), leading to emergence of a PA-containing pre-LECA archaeon (PAPLA) with primitive endosymbiosis. **e**, Maturation of PAPLA-PA symbiosis and development of PA parasitism. **f**, Enslavement of PA by PAPLA through delegation of ATP generating metabolism to PA.

Two routes may be possible: acquisition of aerobic respiration (*i.e.*, electron transport chain and terminal oxidases) or an O_2_-utilizing endosymbiont. We hypothesize that the ancestral Heimdallarchaeon (or a specific sub-lineage) adopted the former route (Fig. 4b) and the pre-last eukaryotic common ancestor (LECA) archaeon took the latter. Prior to endosymbiosis, the pre-LECA archaeon likely interacted with SRB and O_2_-utilizing organotrophs, who maintained the local habitats O_2_ concentrations low (Fig. 5b). The O_2_-utilizing partner was likely a facultative aerobe capable of aerobic and anaerobic H_2_-generating organotrophy. In this three-member interaction, the SRB could syntrophically scavenge H_2_ from both the pre-LECA archaeon and facultatively aerobic partner. The dynamic oxic-anoxic-adaptable symbiosis could have strengthened the three-member interaction and physical association. Moreover, the pre-LECA archaeon is predicted to lack many biosynthetic pathways (Extended Data Fig. 4) and, thus, would have still depended on metabolite exchange with partners for growth. One of the facultatively aerobic partners was likely the pre-mitochondrial alphaproteobacterium (PA; *i.e.*, future mitochondrion) as it has been proposed that PA would be capable of aerobic and anaerobic H_2_-generating organotrophy^4^. Evolution of the symbiosis likely led to PA endosymbiosis into the pre-LECA archaeon, resulting in a transitional PA-containing pre-LECA archaeon (PAPLA) using PA as an O_2_-scavenging and building-block-providing symbiont essential for growth under microaerobic conditions even without SRB.

Note that it is entirely possible for the H_2_-consuming partner to have become endosymbionts of the pre-LECA archaeon as proposed previously^14,33^. However, H_2_-consuming partners would have had less advantage as endosymbionts. H_2_ consumers would prefer higher H_2_ concentrations, but an enlarged host would require larger amounts of substrate to accumulate H_2_ compared to a small prokaryotic H_2_ producers with concentrated H_2_ generation. Moreover, H_2_ is membrane-permeable, so there is little benefit to being inside an H_2_ producer. Thus, endosymbiosis of an H_2_ utilizer is unfavorable without uncompartmentalized H_2_ generation and unlikely to have stabilized. In fact, extant methanogenic endosymbionts are observed only in hydrogenosome-possessing protozoa^34^.

How did the endosymbiosis physically manifest? Given the structure of extant eukaryotic cells, it is logical to presume that the pre-LECA archaeon engulfed their metabolic partner. Although a phagocytosis-like process has been previously proposed^6^, (i) the observed MK-D1 cells are much too small to engulf their metabolic partner in this way, (ii) Asgard archaea lack phagocytotic machinery^35^, and (iii) a pre-mitochondriate organism lacks sufficient energy to perform phagocytosis^36^. Based on the observation of unusual morphological structures of MK-D1 cells (Fig. 3 and Extended Data Fig. 2), the pre-LECA Asgard archaeon may have produced protrusions and/or MVs (Fig. 5b). For an archaeon syntrophically growing in a narrow space (*e.g.*, sediment pore), it may have been possible for the protrusions/MVs to fuse and inadvertently surround its partner, resulting in phagocytosis-independent engulfment (Fig. 5c). There are many possible triggers for membrane fusion, including mechanical stress, electric current, or even evolution of membrane-fusing proteins (*e.g.*, SNARE)^37^. Unlike phagocytosis, such a process would assimilate the partner and simultaneously form a chromosome-bounding membrane structure topologically similar to that of the eukaryotic nuclear membrane (Fig. 5d), a scheme similar to the “Inside-out model” presented by Baum and Baum (2014)^38^.

PAPLA likely shared 2-oxoacids with the endosymbiotic PA, given that AA-degrading pathways widely encoded by Asgard archaea primarily recover ATP from 2-oxoacid degradation (*i.e.*, 2-oxoacid oxidation and acyl-CoA hydrolysis; Figs. 4b and 5d). Under anaerobic conditions, PAPLA and PA may have shared AA-derived 2-oxoacids and both produced H_2_ for syntrophic interaction with SRB. Conversely, when exposed to microaerobic conditions, PAPLA likely catabolized AA in syntrophy with SRB but also provided 2-oxacids to PA and stimulate O_2_ consumption. PAPLA could have theoretically absorbed the endosymbiotic PA’s metabolic capacity to become a more metabolically versatile unicellular organism; however, maintaining a respiratory endosymbiont was likely beneficial because the high surface-area-to-volume ratio of endosymbiotic (smaller) cells can maximize electron-driven energy synthesis (*i.e.*, oxidative phosphorylation), which is kinetically limited by membrane surface area. Moreover, the O_2_-consuming symbiont could theoretically produce large amounts of energy for biosynthesis and allow the host PAPLA access to an intracellular pool of biological building blocks (*e.g.*, cofactors) without the need for active transport by PAPLA itself.

To mature the endosymbiosis, streamlining of metabolic processes is paramount. Two major redundancies are lipid biosynthesis and 2-oxoacid-driven ATP generation. As the hosting PAPLA had ether-type lipids (as evidenced by MK-D1; Fig. 3j) and PA likely had ester-type, two lipid types coexisted in the hybrid cell (Fig. 5d). As horizontal gene transfer between the host and symbiont (or potentially other bacterial source) proceeded, PAPLA likely lost synthesis of ether-type lipids and acquired that of ester-type to resolve redundancy (*i.e.*, streamline genome) in lipid biosynthesis, and passively exchanged the ether-type lipids with ester-type through dilution via cell division (lipid types can mix without compromising structure/fluidity^39^). Combining the engulfment described above and this lipid exchange, we reach a PAPLA with single-layered ester-type lipid plasma and nuclear membranes, which is consistent with extant eukaryotes (Fig. 5e).

Although PAPLA and PA initially shared 2-oxoacids, to streamline, this symbiosis must transition towards delegation of this catabolism to one side and sharing of the generated ATP. For maximizing aerobic oxidative phosphorylation, it is reasonable for PA to catabolize 2-oxoacids, generate ATP, and transport ATP to the host cytosol. This is analogous to the symbiosis between extant eukaryotes and their mitochondria (or hydrogenosomes) and, thus, evolution of the ATP transporter (*i.e.*, ADP/ATP carrier or AAC) was a major factor in fixing the symbiosis^40^. However, it is unrealistic to think that ATP-providing machinery evolved altruistically. We hypothesize that PA first developed AAC as a parasitic tool to absorb host-synthesized ATP, as in extant pathogenic *Rickettsia* and *Chlamydiae*^41^ (Fig. 5e). To combat this parasitism, PAPLA could have evolved to (i) lose pyruvate (and other 2-oxoacids) catabolism, then (ii) delegate 2-oxoacid degradation and ATP synthesis to PA, (iii) generate an ATP concentration gradient from PA to the cytosol, and ultimately (iv) reverse the direction of AAC activity^41^ (Fig. 5f). For PAPLA to evolve towards retaining both aerobic and anaerobic metabolism, the progression above is critical because PAPLA-PA symbiosis would have heavily leaned towards parasitism under anaerobic conditions (*i.e.*, taking both 2-oxoacids and ATP and ATP with little to no return). On the other hand, adaptation to aerobic conditions through loss of O_2_-sensitive ferredoxin-dependent 2-oxoacid metabolism (*e.g.*, pyruvate:ferredoxin oxidoreductase) and consequent delegation of 2-oxoacid-degrading ATP generation to PA (via 2-oxoacid dehydrogenases) would have allowed evolution towards aerobiosis. In this symbiosis, PAPLA and PA mutually benefit – PAPLA can allot energy metabolism to PA and indirectly obtain energy from organotrophy via AAC while PA is fed 2-oxoacids for energy production (Fig. 5f). Here, PAPLA enslaves PA and we arrive at LECA possessing symbiosis congruent with that of extant eukaryotes and their mitochondria.

In summary, we obtained the first isolate of Asgard archaea with unique metabolic and morphological features through a total of 12 years cultivation, and combining these observations with genomic analyses, propose the “Entangle-Engulf-Enslave (E^3^) model” for eukaryogenesis from archaea. Maturation of this model requires elucidation and incorporation of the timing/progression of lateral gene transfer between PAPLA and PA or other bacteria, PA simplification and organellogenesis towards the first mitochondrion, cell complexification (*e.g.*, non-mitochondrial organelle development), and elaboration of eukaryotic cell division and their ties with Earth’s history. Endeavors in cultivation of other Asgard archaea and more deep-branching eukaryotes are essential to further unveil the road from archaea to eukaryotes.

## Supplementary Information

is available in the online version of the paper.

## Acknowledgements

We thank Hiroyuki Ohno and Tsuyoshi Yamaguchi for assistance with HCR-FISH analysis, Takeshi Terada for help of NanoSIMS sample preparation, Masami Isozaki for assistance with cultivation experiments, Takaaki Kubota for assistance with chemical analysis, Kiyotaka Takishita, Akinori Yabuki, Takashi Shiratori, Akiyoshi Ohashi, Fumio Inagaki, Takuro Nunora, Shinsuke Kawagucci, Takazo Shibuya, Shun’ichi Ishii, Yusuke Tsukatani and Yutetsu Kuruma for useful advice and discussion and Ai Miyashita, Yuto Yashiro, Ken Aoi, Masayuki Ehara, Masataka Aoki and Yayoi Saito for assistance with the bioreactor operation. We also thank Juichiro Ashi and the R/V Yokosuka and “*Shinkai 6500*” operation team during cruise YK06-03 (JAMSTEC) and the shipboard scientists and crews of the *Chikyu* Shakedown Cruise CK06-06 for their assistance in collecting samples. This study was partially supported by grants from the Japan Society for the Promotion of Science (JSPS) (KAKENHI Grants 18687006, 21687006, 24687011, 15H02419 and 19H01005 to H.I., 18H03367 to M.K.N., 26710012, 18H02426, 18H05295 to H.T., and Grant-in-Aid for JSPS Fellow 16J10845 to N.N.). This work was also supported by JSPS KAKENHI Grant Number JP16H06280, Grant-in-Aid for Scientific Research on Innovative Areas – Platforms for Advanced Technologies and Research Resources “Advanced Bioimaging Support” and the Cooperative Study Program (19-504) of National Institute for Physiological Sciences.

## Author contributions

H.I. conceived the study and deep marine sediment sampling. H.I., N.N., M.O., M.M. and S.S. conducted cultivation and culture-based experiments. M.K.N. and Y.Takaki performed genome analysis. H.I., N.N., Y.Morono, M.O., T.I., M.I, K.M., C.S. and K.U. undertook the microscopy and NanoSIMS work. M.O., Y.S. and Y.Y. performed qPCR, SSU rRNA gene analysis and DNA/RNA sequencing. Y.Takano, Y. Matsui and E.T. performed chemical analysis. H.I., M.K.N., N.N., Y.Morono, Y. Takaki, Y.Takano, K.M., C.S., T.Y. Y.K. H.T. and K.T. conducted data interpretation. H.I., M.K.N., Y.Takano, H.T., Y.K. and K.T. wrote the manuscript with input from all co-authors. All authors have read and approved the manuscript submission.

## Author Information

The authors declare no competing financial interests. Correspondence and requests for materials should be addressed to H.I or M.K.N.

## Methods

No statistical methods were used to predetermine sample size.

### Sampling site and sample description

A 25-cm long sediment core (949C3) was collected from a methane-seep site at the Omine Ridge, Nankai Trough, off the Kumano area, Japan (33°7.2253’N, 136°28.6672’E), 2,533 m below the sea level, via the manned submersible “*Shinkai 6500*” (cruise YK06-03, dive no. 6K949, May 6th, 2006). The detailed sediment core sample and site information has been described previously^15,42,43^. Our previous geochemical and 16S rRNA gene analysis indicated that the occurrence of anaerobic oxidation of methane (AOM) reactions mediated by archaeal anaerobic methanotrophs (ANMEs) in the sediment^15,42^. The SSU rRNA gene analysis also showed that the sediment contained abundant and diverse microorganisms, most of which were affiliated with uncultured microbial groups, including Asgard archaea^15,42^.

### Culturing

The deep-sea methane-seep sediment sample was first enriched using a continuous-flow bioreactor system supplemented with methane as the major energy source. The bioreactor, called a down-flow hanging sponge (DHS) bioreactor, has operated in our laboratory, JAMSTEC, Yokosuka Headquarters, since December 28, 2006. The detailed operation conditions for the DHS bioreactor have been described previously^15^. To isolate anaerobic microorganisms, including Asgard archaea, from the DHS reactor, 2 ml samples of the bioreactor enrichment sediment slurry were inoculated in 15 ml glass tubes with a simple substrate and a basal medium. The composition of the basal medium was almost similar to that used for cultivation in the DHS bioreactor^15^, but it did not contain sulfate (i.e., Na_2_SO_4_). The basal medium composition was as follows (per liter): 9.47 g MgCl_2_·6H_2_O, 1.36 g CaCl_2_·2H_2_O, 20.7 g NaCl, 0.54 g NH_4_Cl, 0.14 g KH_2_PO_4_, 2.7 g NaHCO_3_, 0.3 g Na_2_S·9H_2_O, 0.3 g cysteine·HCl, 1 ml trace element solution^15^, 1 ml Se/W solution, 2 ml vitamin solution^15^ and resazurin solution (1 mg/ml). The medium was purged with N_2_/CO_2_ gas (80:20, v/v), and the pH was adjusted to 7.5 at 25°C. The culture tubes were sealed with butyl rubber stoppers and screw caps. Autoclaved or filter-sterilized organic substances (such as protein-derived materials, sugars, and fatty acids) were added to the tubes with stock solutions prior to inoculation with the bioreactor enriched community. After establishing a stable *Ca*. P. syntrophicum culture, cultivations were performed at 20°C in 50-ml serum vials containing 20 ml basal medium supplemented with CA (0.05%, w/v), 20 AAs (0.1 mM each), and PM (0.1%, w/v, Hohoemi, Meiji Co., Ltd.) under an atmosphere of N_2_/CO_2_ (80:20, v/v) in the dark without shaking, unless mentioned otherwise. Information regarding the purity check of MK-D1 cultures, as well as additional information about cultivation, is described in the Supplementary Methods.

### SSU rRNA gene-based analysis

DNA extraction and PCR mixture preparation were performed on a clean bench to reduce contamination. DNA extraction from culture samples was performed as described previously^44^. The concentration of extracted DNA was measured using a Quant-iT dsDNA High-Sensitivity Assay Kit (Life Technologies). PCR amplification was performed using the TaKaRa Ex *Taq* (for conventional clone analysis) or TaKaRa LA *Taq* (for Illumina-based amplicon sequencing [iTAG] for targeted sequencing for the SSU rRNA gene analysis) (TaKaRa Bio Inc.), and the reaction mixtures for PCR were prepared according to the manufacturer’s instructions. For the conventional clone analysis, a universal primer pair 530F/907R^42^ and an archaeal primer pair 340F/932R^15,45^ were used for PCR amplification. For iTAG analysis, the universal primer pair 530F/907R, which contained overhang adapters at 5’ ends, was used. The procedures used for library construction, sequencing, and data analysis were described previously^19,46^.

### Growth monitoring using qPCR

For the quantitative analysis, a StepOnePlus Real-Time PCR System (Thermo Fisher Scientific) with a SYBR Premix Ex Taq II kit (TaKaRa Bio Inc.) was used. The candidate phylum Lokiarchaeota-specific primer pair MBGB525F/Ar912r was used for amplification of 16S rRNA genes. Primer MBGB525F is the complementary sequence of the MGBG525 probe^17^, while Ar912r is an archaeal universal primer that is a slightly modified version of the original designed primer^47^. The detailed procedure for qPCR is described in the Supplementary Methods. The doubling times of MK-D1 were calculated based on the semilogarithmic plot of the qPCR data.

### Growth test with multiple substrates

To examine effect of the presence of other substances on growth of MK-D1, CA–20 AAs–PM medium supplemented with an individual substrate (Extended Data Table 3) was prepared, followed by qPCR and iTAG analyses. Each cultivation condition was set in duplicate; however, the H_2_-fed culture was prepared in triplicate because Sousa *et al*. (2016)^7^ reported that a Lokiarchaeum has potential to grow with hydrogen based on a comparative genome analysis. Detailed culture liquid sampling and the subsequent qPCR and iTAG analyses are described in the Supplementary Materials.

### Evaluation of growth temperature

The test was performed using a basal medium containing CA and PM, with a pure co-culture of MK-D1 and *Methanogenium* as the inoculum (20%, v/v). The cultures were incubated at 4, 10, 15, 20, 25, 30, 37, and 40°C. All incubations for the test were performed in triplicate. After 100 days of incubation, 16S rRNA gene copy numbers of MK-D1 were evaluated through qPCR technique.

### FISH

Fixation of microbial cells, storage of the fixed cells, and standard FISH were performed in accordance with a previously described protocol^19^. The 16S rRNA-targeted oligonucleotide probes used in this study are listed in Supplementary Table S3. The designing of MK-D1-specific probes is described in the Supplementary Methods. As clear fluorescent signals were not obtained using the standard FISH technique, we employed an *in situ* DNA-hybridization chain reaction (HCR) technique^48^. The FISH samples were observed using epifluorescence microscopes (BX51 or BX53, Olympus) and a confocal laser scanning microscope (Nikon A1RMP, Nikon Instech).

### SEM

Microbial cells were fixed overnight in 2.5% (w/v) glutaraldehyde in the CA–20 AAs medium at 20°C. The sample preparation procedure has been described previously^49^. The cell samples were observed under a field emission (FE)-SEM (JSM-6700F, JEOL) or an extreme high-resolution FIB-SEM (Helios G4 UX, ThermoFisher Scientific).

### Ultrathin sectioning and TEM

Cells were prefixed with 2.5% (w/v) glutaraldehyde for 2 h. The specimens were frozen in a high-pressure freezing apparatus (EM-PACT2, Leica)^50^. The frozen samples were substituted with 2% OsO_4_ in acetone for 3–4 days at - 80°C, and the samples were warmed gradually to room temperature, rinsed with acetone embedded in epoxy resin (TAAB) Thin sections (70 nm) were cut with a ultramicrotome (EM-UC7, Leica). Ultra-thin sections of the cells were stained with 2% uranyl acetate and lead-stained solution (0.3% lead nitrate and 0.3% lead acetate, Sigma-Aldrich), and were observed by a transmission electron microscopy (Tecnai 20, FEI) at an acceleration voltage of 120 kV.

### Cryo-EM

Due to the low cell yield-culture, 400 ml of the culture of MK-D1 was prepared and concentrated to about 5 ml using a 0.22 µm-pore-size polyethersulfone (PES) filter unit (Corning) in an anaerobic chamber (95:5 [v/v] N_2_:H_2_ atmosphere; COY Laboratory Products). The concentrated culture liquid was placed in a glass vial in the anaerobic chamber. After that, the head space of the glass vial was replaced by N_2_/CO_2_ gas (80:20, v/v). Immediately before the electron microscopic observation, the glass vial was opened, and the liquid culture was concentrated to about 200 µl by centrifugation at 20,400 g for 10 min at 20°C. Subsequently, 3 µl of the concentrated liquid culture was applied onto a Quantifoil Mo grid R1.2/1.3 (Quantifoil MicroTools) pretreated with glow-discharge, and was plunged-frozen in liquid ethane using a Vitrobot Mark IV (FEI Company) at 4°C and 95% humidity.

The frozen grid was mounted onto a 914 liquid-nitrogen cryo-specimen holder (Gatan Inc.) and loaded into a JEM2200FS electron microscope (JEOL) equipped with a field emission electron source operating at 200 kV and an omega-type in-column energy filter (slit width: 20 eV). The images were recorded on a DE-20 direct detector camera (Direct Electron LP.) at a nominal magnification of 15,000 x, which resulted in an imaging resolution of 3.66 Å per pixel, with the total dose under 20 electrons per Å^2^ using a low dose system. For electron tomography, tilt series images were collected manually in a range of ∼± 62° at 2° increments. The total electron dose on the specimen per tilt series was kept under 100 electrons per Å^2^ to minimize radiation damage. The tilt series were aligned using gold fiducials, and tomograms were reconstructed using filtered back projection or SIRT in the IMOD software^51^ with an image binning of 5.

### Lipid analysis

About 120 ml of a highly purified culture sample were concentrated using the same method described above, except that the filtration concentration procedure was performed on a clean bench instead of the anaerobic chamber. Following cell collection, the cells were washed with the anaerobic basal medium to eliminate the interfering matrix. Subsequently, lipid analysis was conducted for the collected cells after the improved method^52^. For precise qualitative liquid analysis, gas chromatography (GC) combined with mass spectrometry (MS) on the 7890 system (Agilent Technologies Inc.) was conducted to compare the retention time and mass fragmentation signatures.

### Stable isotope probing and NanoSIMS analysis

To confirm utilization of amino acids by MK-D1, a stable isotope probing experiment was performed using a ^13^C- and ^15^N-labeled amino acids mixture (Cambridge Isotope Laboratories). Briefly, 120 ml serum vials containing 40 ml basal medium were prepared and supplemented with the stable isotope labeled 20 AAs (roughly 0.1 mM of each), CA (0.05%, w/v) and non-labeled 20 AAs mixture (0.1 mM of each). Two types of highly purified cultures of MK-D1 were used as inocula: a co-culture with *Methanobacterium* sp. strain MO-MB1 and a tri-culture with *Halodesulfovibrio* and *Methanogenium*. The vials were incubated at 20°C in the dark without shaking for 120 days. A reference cultivation was also performed under the same cultivation condition without the addition of the stable isotope labeled 20 AAs mixture (Extended Data Table 2). The detailed sample preparation and analysis method using NanoSIMS is described in the Supplementary Methods.

### Chemical analysis

The stable carbon isotope compositions of methane and carbon dioxide in the sampled gas phase were analyzed as described previously (Okumura et al. 2016). Methane concentrations were measured by gas chromatography (GC-4000, GL Science Inc., Tokyo, Japan) using a Shincarbon ST 50/80 column (1.0 m x 3.0 mm ID, Shinwa Chem. Ind.) and a flame ionization detector with nitrogen as a carrier gas.

Amino acid concentrations in pure co-cultures of MK-D1 and *Methanogenium* were quantified through a previously described method^53,54^. In brief, we processed the acid hydrolysis with 6 M HCl (110°C, 12 h) for the culture liquid samples after filtration using a 0.2 µm pore-size polytetrafluoroethylene filter unit (Millipore). The amino acid fraction was derivatized to N-pivaloyl iso-propyl esters prior to GC using a 6890N GC instrument connected to the nitrogen phosphorus and flame ionization detectors (Agilent Technologies Inc.). For cross-validation of qualitative identification of amino acids, GC-MS on the 7890 system (Agilent Technologies Inc.) was used^52^.

### Genome sequencing and assembly

DNA extraction was performed as described previously^44^. Mate-paired library with an average insert size of 3000 bp was constructed according to the manufacturer’s instructions with Nextera Mate Pair Library Preparation kit (Illumina). Library sequencing was performed using Illumina MiSeq platform (2 x 300 bp), which resulted in 3,822,290 paired reads. The mate pair reads were processed as follows: adapters and low-quality sequences were removed using Trimmomatic ver. 0.33^55^, and the linker sequences were removed using NextClip ver. 1.3.1^56^. *De novo* assembly was performed using SPAdes ver 3.1.1^57^ with multiple k-mer sizes (21, 33, 55, 77, and 99), which resulted in 3,487 contigs with lengths >500 bp, totaling upto 14.68 Mbp. The software MyCC^58^ was used with default parameters for binning based on genomic signatures, marker genes, and contig coverages. As heterogeneity in the sequence can cause highly fragmented or redundant contigs, the ambiguous contigs (sequence coverage < 5 or a length < 1kb) and redundant contigs were discarded from binning. This resulted in the recovery of genomes related to Lokiarchaeum (i.e., *Ca*. P. syntrophicum MK-D1, 4.46 Mbp), *Halodesulfovibrio* (4.13 Mbp) and *Methanogenium* (2.33 Mbp). Scaffold for each bin were constructed using SSPACE ver. 3.0^59^ with mate-paired information of Illumina reads. To obtain the complete genome sequence of *Ca.* P. syntrophicum, the gaps were filled using Sanger sequencing method. Genomes were annotated using Prokka v1.12^60^ and manually curated. The curation involved functional domain analysis through CD-Search with its corresponding conserved domain database^61,62^; signal peptide and transmembrane domain prediction through SignalP v4.1^63^; carbohydrate-active enzyme, peptidase, and lipase prediction through dbCAN 5.0^64^, MEROPS^65^, and lipase engineering database^66^; and hydrogenase annotation with assistance from HydDB^67^. In addition, to further verify the function, we compared the sequence similarity of each gene to UNIPROTKB/SWISSPROT containing enzymes with experimentally verified catalytic activity and genes with extensive genetic, phylogenetic, and/or genomic characterizations^68,69^ with a 40% amino acid similarity cutoff. For enzymes that have divergent functions even with a 40% similarity cutoff (e.g., [FeFe] and [NiFe] hydrogenases, 3-oxoacid oxidoreductases, glutamate dehydrogenases, and sugar kinases), phylogenetic trees were constructed with reference sequences to identify association of the query sequences to phylogenetic clusters containing enzymes with characterized catalytic activity.

### Phylogenetic analysis

Phylogenomic tree of MK-D1 and select cultured archaea, eukaryotes, and bacteria. 31 ribosomal proteins conserved across the three domains (Supplementary Table S4) were collected from MK-D1, the organisms shown in the tree, and metagenome-assembled genomes (MAGs) of uncultured archaeal lineages (Supplementary Table S5). Two alignments were performed in parallel: (i) only including sequences from cultured organisms and (ii) also including MAG-derived sequences. MAFFT v7 (--linsi) was used for alignment in both cases^70^. For the latter, MAG-derived sequences were included to generate an alignment that maximizes the archaeal diversity taken into account, but removed for subsequent tree construction to avoid any influence of contamination (*i.e.*, concatenation of sequences that do not belong to the same organism). *Ca.* Korarchaeum sequences were kept in the tree based on the cultured+uncultured alignment due to its critical phylogenetic position in TACK phylogeny. After removing all-gap positions and concatenation, the maximum likelihood trees were constructed using RAxML-NG (fixed empirical substitution matrix [LG], 4 discrete GAMMA categories, empirical AA frequencies, and 100 bootstrap replicates)^71^. Bootstrap values around critical branching points are also shown. For 16S ribosomal RNA phylogeny, sequences were aligned using SINA^72^ against the Silva v132 alignment^73^. The maximum likelihood tree was calculated using RAxML^74^ using fixed empirical substitution matrix (LG), 4 discrete GAMMA categories, empirical amino acid frequencies from the alignment, and 100 bootstrap replicates. For analysis of urocanate hydratase, serine/threonine dehydratase, succinate dehydrogenase flavoprotein, fatty-acid--CoA ligase, and 3-ketoacyl-CoA thiolase homologs were collected through BLASTp analysis of the Asgard archaea sequences against the UniProt database (release 2019_06). Of homologs with sequence similarity ≥40% and overlap ≥70%, representative sequences were selected using CD-HIT with a clustering cutoff of 70% similarity (default settings otherwise). Additional homologs with verified biochemical activity, sequence similarity ≥30%, and overlap ≥70% were collected through BLASTp analysis of the Asgard archaea sequences against the UniProt/SwissProt database. Sequences were aligned using MAFFT v7^70^ with default settings and trimmed using trimAl^75^ with default settings. The phylogenetic tree was constructed using RAxML-NG^71^ using fixed empirical substitution matrix (LG), 4 discrete GAMMA categories, empirical amino acid frequencies from the alignment, and 100 bootstrap replicates.

For analysis of biotin ligase and biotin carboxyl carrier protein, homologs were collected through BLASTp analysis of the Asgard archaea sequences against the UniProt database (release 2019_06). Of homologs with sequence similarity ≥40% and overlap ≥70%, representative sequences were selected using CD-HIT with a clustering cutoff of 70% similarity (default settings otherwise). Additional homologs with verified biochemical activity, sequence similarity ≥30%, and overlap ≥70% were collected through BLASTp analysis of the Asgard archaea sequences against the UniProt/SwissProt database. Sequences were aligned using MAFFT v7^70^ with default settings and trimmed using trimAl^75^ with default settings. The phylogenetic tree was constructed using FastTree^76^ using fixed empirical substitution matrix (LG) and 1000 bootstrap replicates.

### RNA based sequencing analysis

To perform RNA based sequencing analysis, 100 ml of culture liquid were prepared from five highly purified cultures that were incubated with CA, 20 AAs, and PM for about 100 days at 20°C. Before RNA extraction, the growth of MK-D1 was confirmed using the qPCR technique, and the cells density levels were ∼10^5^ copies/ml in each culture.

To harvest microbial cells, the culture liquid was filtered through a 0.22-µm pore-size mixed cellulose ester membrane filter (GSWP01300, Merck MilliPore) on a clean bench. After filtration, the membrane was cut in half with a sterilized scissors and then directly inserted into the PowerBiofilm bead tubes of a PowerBiofilm RNA Isolation kit (MO BIO Laboratories). The following RNA extraction procedures were performed according to the manufacturer’s instructions. The extracted RNA was applied to an RNA clean & concentrator kit-5 (Zymo Research) for concentration. The obtained RNA was quantified using an Agilent 2100 Bioanalyzer system with an RNA Pico kit (Agilent Technologies) and then applied to an Ovation Universal RNA-Seq System (NuGEN Technologies) for the construction of an RNA sequence library. At the step for Insert Dependent Adaptor Cleavage technology mediated adaptor cleavage during the library construction, specific primers for 16S rRNA and 23S rRNA genes of MK-D1 were used to reduce rRNA gene sequences from the cDNA pool. The constructed cDNA library was sequenced using the MiSeq platform (Illumina).

The raw RNA sequencing data were trimmed by removal of the adapters and low-quality sequences using Trimmomatic ver. 0.33^55^. The expression abundance of all coding transcripts was estimated in RPKM values using EDGE-pro ver. 1.3.1^77^.

### Data availability

Genomes for *Ca*. Prometheoarchaeum syntrophicum MK-D1, *Halodesulfovibrio* sp. MK-HDV, and *Methanogenium* sp. MK-MG are available under Genbank BioProjects PRJNA557562, PRJNA557563, and PRJNA557565 respectively. The iTAG sequence data was deposited in Bioproject PRJDB8518 with the accession numbers DRR184081–DRR184101. The 16S rRNA gene sequences of MK-D1, *Halodesulfovibrio* sp. MK-HDV, *Methanogenium* sp. MK-MG and clones obtained from primary enrichment culture were deposited in the DDBJ/EMBL/GenBank database under accession numbers LC490619–LC490624.

