## Extended Data for "Isolation of an archaeon at the prokaryote-eukaryote interface"

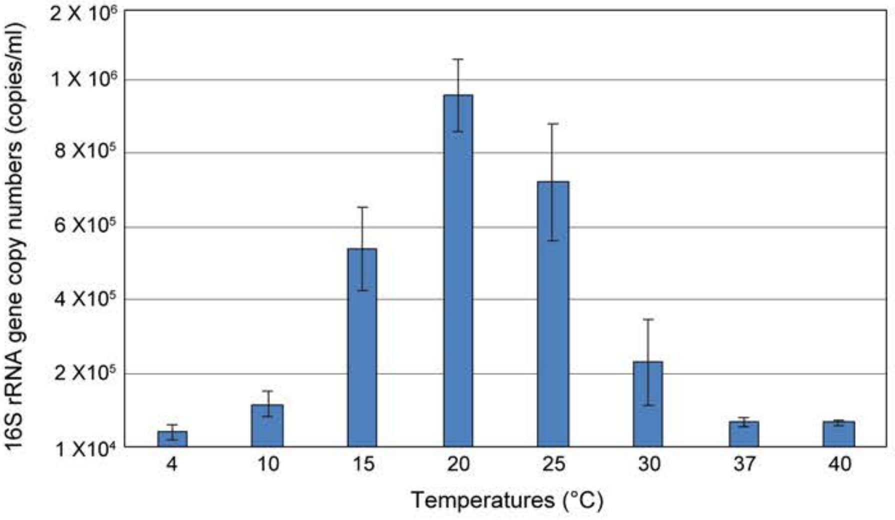


**Extended Data Fig. 1 | Effect of temperature on growth of MK-D1.** Error bars indicate standard deviations of triplicate determinations. The temperature range test was performed twice, and both results displayed similar results.

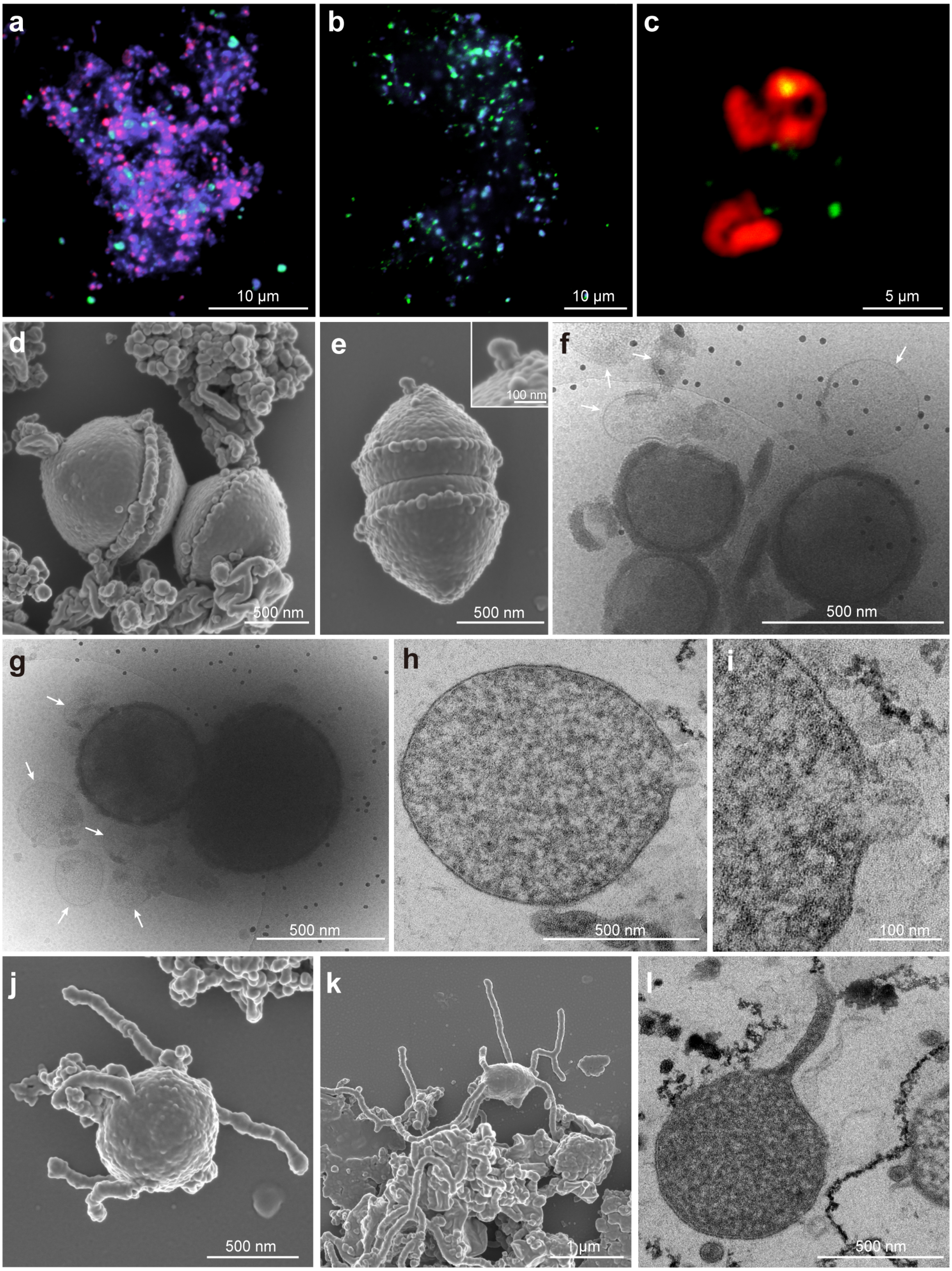


**Extended Data Fig. 2 | Other representative photomicrographs of MK-D1. a, b,** Fluorescence images of cells from enrichment cultures after eight (**a**) and eleven (**b**) transfers stained with DAPI (violet) and hybridized with nucleotide probes targeting MK-D1 (green) and *Bacteria* (red). The images are other fields of view, which were taken at the same time for the Figs. 1b and c images. **c**, A fluorescence image of cells in the enrichments after eleven transfers hybridized with nucleotide probes targeting MK-D1 (green) and *Archaea* (but with one mismatch against MK-D1; red). (Large and irregular coccoid-shaped cells stained by ARC915 only are likely *Methanogenium*.) **d**, **e**, Dividing cells of MK-D1 with a bleb. The upper-right inset image in **e** shows close-up of the bleb. f, g, Cryo-EM images of MK-D1 cells and large MVs (white arrows). **h**, **i**, Ultrathin sections of MK-D1 cells with an MV. The image **i** shows a close-up image of **h**. **j**, **k**, SEM images of MK-D1 cells with protrusions. **l**, Ultrathin section of a MK-D1 cell with a protrusion. Detailed iTAG analyses of cultures are shown in Supplementary Table S2.

**a**

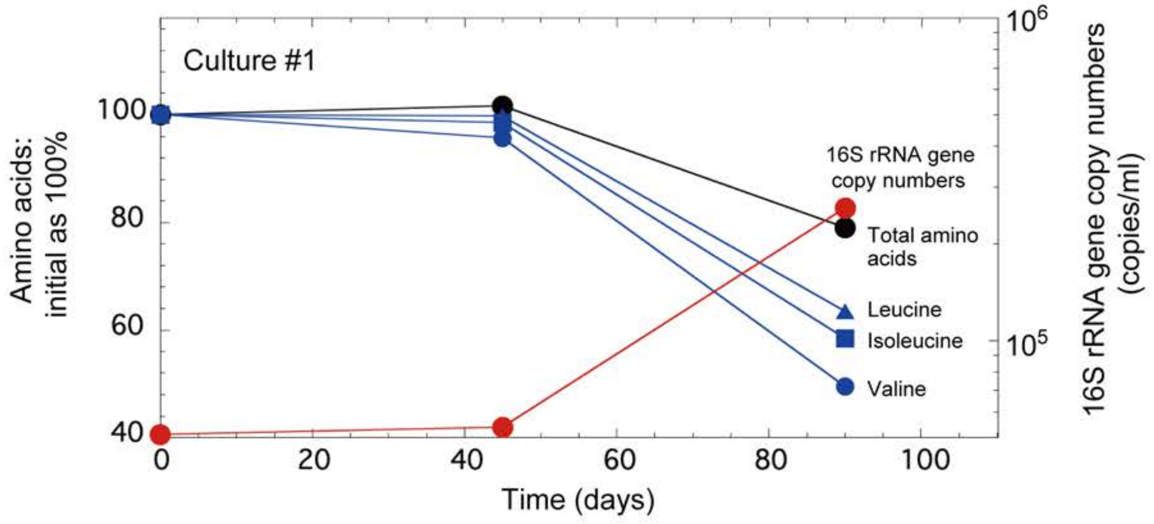


**b**

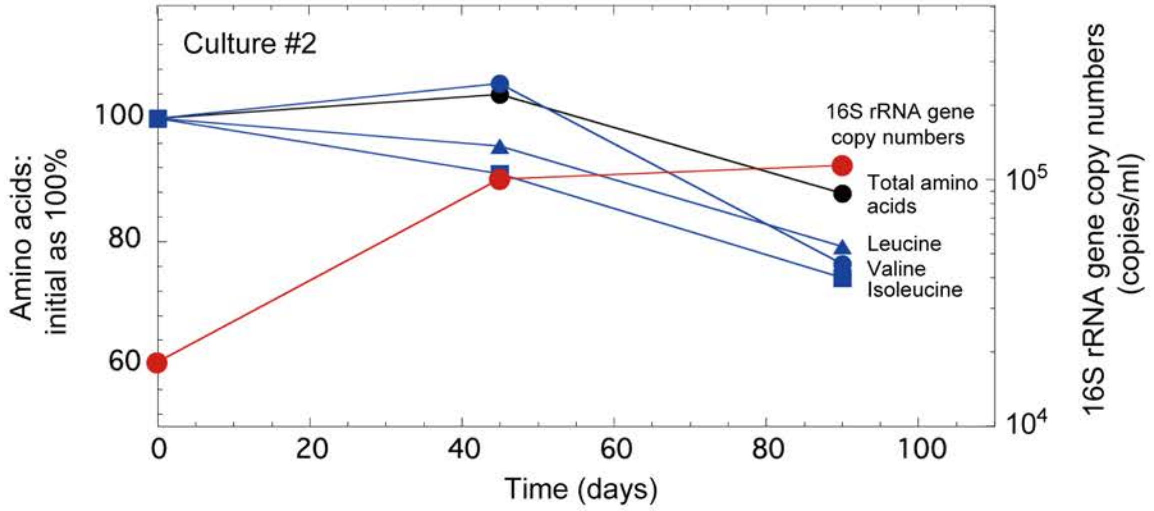


**Extended Data Fig. 3 | The amino acid concentrations and growth curves of MK-D1 in pure co-cultures. a, b,** Results from cultures #1 (**a**) and #2 (**b**) are shown. Please note that the initial concentrations of amino acids were normalized as 100%. Total amino acids and several representative amino acids (valine, leucine and isoleucine) are independently shown for the duplicate culture samples. Detailed iTAG-based community compositions of the cultures are shown in Supplementary Table S2.

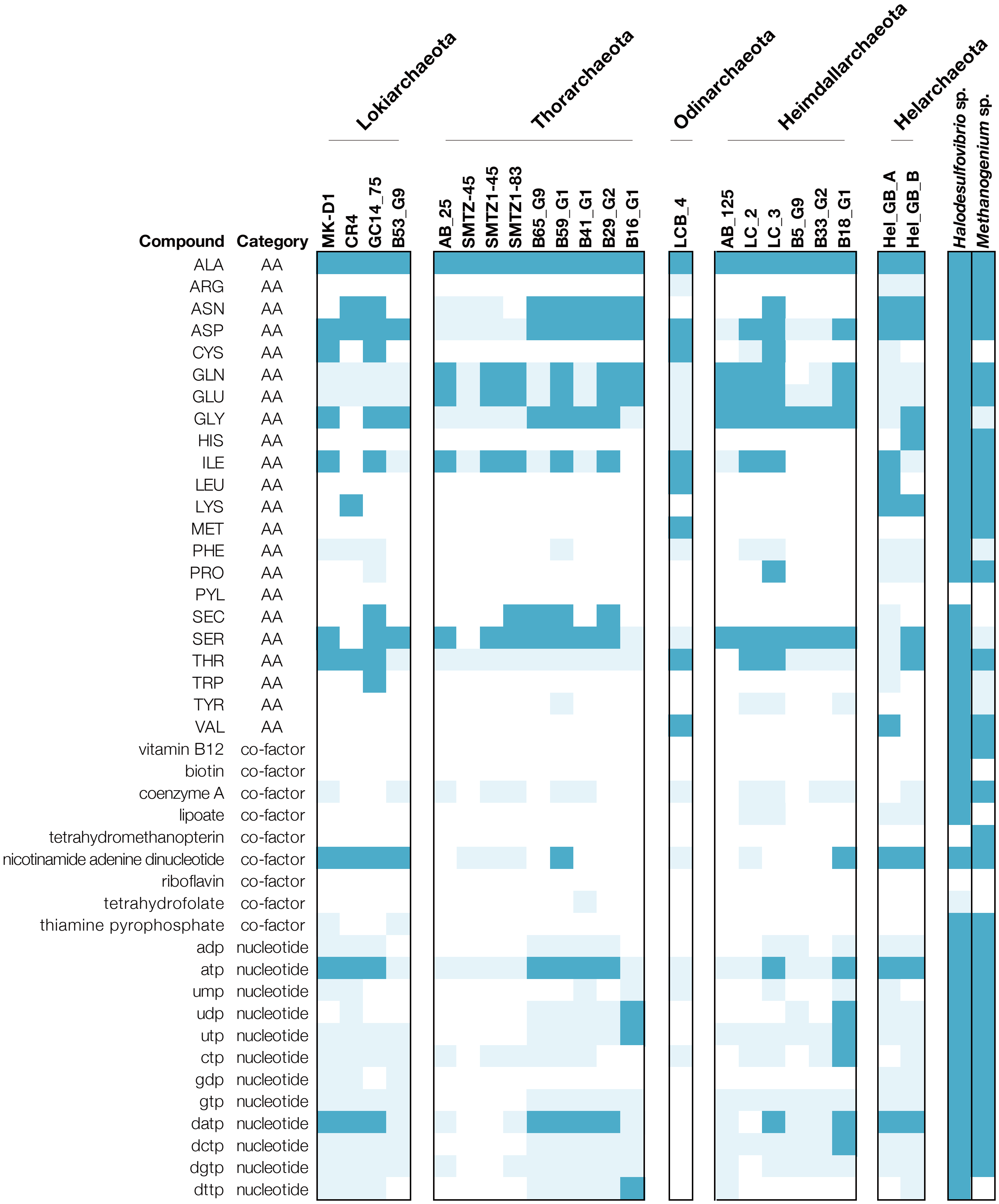


**Extended Data Fig. 4 | AA, co-factor, and nucleotide biosynthesis capacities of MK-D1 and other Asgard archaea.** Genomes encoding genes for synthesis from pyruvate or acetyl-CoA (dark blue) and synthesis from other intermediates (light blue) are indicated. Those without complete pathways for either are indicated white. *Halodesulfovibrio* sp. strain MK-HDV and *Methanogenium* sp. strain MK-MG isolated in this study are also shown.

**Extended Data Table 1 | SSU rRNA gene clones obtained from the primary and six successive transferred enrichment culture**

**<Primary enrichment culture>**

Clone library using universal primers (530F/907R)

| Phylotype name | No. of clones | Accession no. | Sequence length (bp) | Closest cultured species or clone (accession number) | Sequence identity (%) | Phylogenetic affiliation | Identical or almost identical clones detected from the AOM bioreactor enrichment (accession number, sequence identity %)[a] |
|---|---|---|---|---|---|---|---|
| 111_U1 | 40 | LC490621 | 374 | *Halodesulfovibrio aestuarii* strain Sylt 3 (NR_116770) | 99 | genus *Halodesulfovibrio* | — |
| 111_U2 | 3 | — | 377 | *Methylobacter marinus* strain A45 (NR_025132) | 100 | genus *Methylobacter* | MK903D_B19 (AB831411, 100%) |
| 111_U3 | 2 | — | 374 | *Photobacterium indicum* strain NBRC 14233 (NR_113657 ) | 100 | genus *Photobacterium* | MK903D_B9 (AB831402, 100%) |
| 111_U4 | 1 | LC490622 | 377 | subseafloor sediment clone ODP1251B13.14 (AB177314) | 99 | subgroup 21 within the phylum *Acidobacteria* | — |
| 111_U5[b] | 1 | LC490623 | 377 | hydrothermal seep sediment BAC_OTU_13 (KP091106 ) | 100 | GIF9 group within the class *Dehalococcoidia* | MK0D_B60 (AB831337, 99.5%) |
| 111_U6 | 1 | LC490624 | 374 | *Roseovarius gaetbuli* strain YM-20 (NR_134163) | 99 | genus *Roseovarius* | — |

Clone library using archaeal primers (340F/932R)

| Phylotype name | No. of clones | Accession no. | Sequence length (bp) | Closest cultured species or clone (accession number) | Sequence identity (%) | Phylogenetic affiliation | Identical or almost identical clones detected from the AOM bioreactor enrichment (accession bumber, sequence identity %)[a] |
|---|---|---|---|---|---|---|---|
| 111_A1 | 6 | — | 535 | *Methanococcoides burtonii* strain DSM 6242 (NR_074242 ) | 99 | genus *Methanococcoides* | MK903D_A2 (AB831282, 100%) |
| 111_A2 | 5 | LC490620 | 513 | *Methanogenium cariaci* strain JR1 (NR_104730) | 99 | genus *Methanogenium* | — |
| 111_A3 | 2 | — | 534 | methane seep clone AN_5119N_arc_E4_T3 (KM356859) | 99 | ANME-2a | MK0D_A9 (AB831268, 100%) |
| 111_A4 | 2 | LC490619 | 516 | methane seep clone AC_5120_arc_D2_T3 (KM356804) | 99 | Lokiarchaeota (*Ca*. P. syntrophicum strain MK-D1) | MK903R_A35 (AB831305, 99.0%) |

**<Six successive transferred enrichment culture>**

Clone library using universal primers (530F/907R)

| Phylotype name | No. of clones | Accession no. | Sequence length (bp) | Closest cultured species or clone (accession number) | Sequence identity (%) | Phylogenetic affiliation | Identical or almost identical clones detected from the AOM bioreactor enrichment (accession bumber, sequence identity %)[a] |
|---|---|---|---|---|---|---|---|
| 111-5_U1 | 40 | — | 374 | *Halodesulfovibrio oceani* strain I.8.1 (NR_116768) | 100 | genus *Halodesulfovibrio* | — |
| 111-5_U2 | 6 | — | 380 | methane seep clone AC_5120_arc_D2_T3 (KM356804) | 100 | Lokiarchaeota (*Ca*. P. syntrophicum strain MK-D1) | — |
| 111-5_U3 | 1 | — | 380 | *Methanogenium boonei* strain AK-7 (NR_115706) | 99 | genus *Methanogenium* | — |

[a]The clone sequences have been reported in our previous study[15].

[b]An anaerobic bacterium strain MK-GIF9, which has the identical 16S rRNA gene sequence of the OTU, has successfully been isolated from the enrichment culture (Nakahara *et al*. Cultivation of previously uncultured *Chloroflexi*, a bacterial dark matter in subseafloor biosphere. 16th International Symposiumon Microbial Ecology [ISME16], Montreal, Canada [2016]). Detailed infomation about the cultivation, and physiological and genomic properties of the bacterium will be reported in the near future.

**Extended Data Table 2 | Carbon isotope fractionation values in MK-D1 cultures after 120 days incubation with and without stable isotope labeled amino acids**

| Culture ID | $\delta^{13}C\text{-}CO_2$ (‰ VPDB)[a] | $\delta^{13}C\text{-}CH_4$ (‰ VPDB)[a] |
|---|---|---|
| ***Co-cultures with Methanobacterium*** | | |
| No.1 with stable istope labeled AAs | -12.3 | 4094.8 |
| No.2 with stable istope labeled AAs | -9.3 | 6990.7 |
| No.3 w/o stable istope labeled AAs | -23.1 | -36.7 |
| No.4 w/o stable istope labeled AAs | -23.1 | -33.1 |
| ***Tri-cultures with Halodesulfovibrio and Methanogenium*** | | |
| No.5 with stable istope labeled AAs | 318.5 | 86.0 |
| No.6 with stable istope labeled AAs | 309.3 | 87.8 |
| No.7 w/o stable istope labeled AAs | -22.6 | -95.5 |
| No.8 w/o stable istope labeled AAs | -22.8 | -97.8 |

[a]‰ versus the Vienna Pee Dee Belemnite.

**Extended Data Table 3 | Growth of *Ca.* P. syntrophicum strain MK-D1 for 120 days incubation with a range of substrates**

| Culture name | Substrate | Initial MK-D1 16S rRNA gene copies per ml of culture | Final MK-D1 16S rRNA gene copies per ml of culture | No. of MK-D1 16S rRNA gene copies relative to initial culture | Community compositions evaluated by iTAG analysis (%)[a] | | | |
|---|---|---|---|---|---|---|---|---|
| | | | | | MK-D1 | *Methanogenium* sp. | *Methenobacterium* sp. strain MO-MB1 | Others |
| Inoculum | Casamino acids (CA)[b] + 20 amino acids mixture (AAs)[c] + powdered milk (PM)[d] | — | 5.91E+05 | — | 39.8 | 36.8 | 23.3 | 0.01 |
| Control-1 | CA + 20 AAs + PM | 1.42E+04 | 1.62E+05 | 11.36 | 76.7 | 21.8 | 1.4 | 0.03 |
| Control-2 | CA + 20 AAs + PM | 4.67E+03 | 6.55E+04 | 14.03 | 60.3 | 38.0 | 1.6 | 0.04 |
| H2-1 | CA + 20 AAs + PM + 1.5 kPa $H_2$[e] + 10 mM 2-bromoethane sulfonate (2-BES)[f] | 9.46E+03 | 4.35E+03 | 0.46 | — | — | — | — |
| H2-2 | CA + 20 AAs + PM + 1.5 kPa $H_2$ + 10 mM 2-BES | 1.37E+04 | 3.28E+03 | 0.24 | — | — | — | — |
| H2-3 | CA + 20 AAs + PM + 1.5 kPa $H_2$ + 10 mM 2-BES | 3.10E+04 | 8.27E+03 | 0.27 | — | — | — | — |
| Formate-1 | CA + 20 AAs + PM + 1 mM Formate + 10 mM 2-BES | 2.76E+04 | 2.00E+03 | 0.07 | — | — | — | — |
| Formate-2 | CA + 20 AAs + PM + 1 mM Formate + 10 mM 2-BES | 1.46E+04 | 9.49E+03 | 0.65 | — | — | — | — |
| Nitrate-1 | CA + 20 AAs + PM + 500 µM Nitrate[g] | 2.13E+04 | 8.43E+03 | 0.40 | — | — | — | — |
| Nitrate-2 | CA + 20 AAs + PM + 500 µM Nitrate | 1.47E+04 | 5.19E+03 | 0.35 | — | — | — | — |
| Sulfate-1 | CA + 20 AAs + PM + 500 µM Sulfate | 5.28E+03 | 9.21E+04 | 17.42 | 79.5 | 19.5 | 1.0 | 0.03 |
| Sulfate-2 | CA + 20 AAs + PM + 500 µM Sulfate | 3.39E+04 | 5.28E+04 | 1.56 | — | — | — | — |
| Thiosulfate-1 | CA + 20 AAs + PM + 500 µM Thiosulfate | 1.23E+04 | 5.00E+04 | 4.05 | — | — | — | — |
| Thiosulfate-2 | CA + 20 AAs + PM + 500 µM Thiosulfate | 2.29E+04 | 6.09E+04 | 2.66 | — | — | — | — |
| Lactate-1 | CA + 20 AAs + PM + 1 mM Lactate | 5.31E+03 | 1.31E+04 | 2.46 | — | — | — | — |
| Lactate-2 | CA + 20 AAs + PM + 1 mM Lactate | 1.53E+04 | 1.91E+04 | 1.25 | — | — | — | — |
| Acetate-1 | CA + 20 AAs + PM + 1 mM Acetate | 2.63E+04 | 9.17E+04 | 3.48 | — | — | — | — |
| Acetate-2 | CA + 20 AAs + PM + 1 mM Acetate | 1.56E+04 | 2.13E+04 | 1.36 | — | — | — | — |
| Glucose-1 | CA + 20 AAs + PM + 1 mM Glucose | 1.12E+04 | 1.16E+05 | 10.33 | 73.8 | 24.3 | 1.9 | 0.03 |
| Glucose-2 | CA + 20 AAs + PM + 1 mM Glucose | 1.06E+04 | 1.06E+05 | 10.01 | 70.3 | 28.0 | 1.7 | Not detected |
| Fructose-1 | CA + 20 AAs + PM + 1 mM Fructose | 3.18E+04 | 3.31E+04 | 1.04 | — | — | — | — |
| Fructose-2 | CA + 20 AAs + PM + 1 mM Fructose | 1.79E+04 | 1.44E+05 | 8.08 | — | — | — | — |
| Xylose-1 | CA + 20 AAs + PM + 1 mM Xylose | 2.82E+04 | 6.79E+03 | 0.24 | — | — | — | — |
| Xylose-2 | CA + 20 AAs + PM + 1 mM Xylose | 9.25E+03 | 1.18E+05 | 12.73 | 61.4 | 36.5 | 2.1 | 0.01 |
| Ribose-1 | CA + 20 AAs + PM + 1 mM Ribose | 1.42E+04 | 2.88E+04 | 2.02 | — | — | — | — |
| Ribose-2 | CA + 20 AAs + PM + 1 mM Ribose | 7.34E+03 | 2.29E+04 | 3.13 | — | — | — | — |
| Maltose-1 | CA + 20 AAs + PM + 1 mM Maltose | 2.84E+04 | 1.21E+05 | 4.25 | — | — | — | — |
| Maltose-2 | CA + 20 AAs + PM + 1 mM Maltose | 2.17E+04 | 4.55E+04 | 2.09 | — | — | — | — |
| Citrate-1 | CA + 20 AAs + PM + 1 mM Citrate | 3.36E+04 | 1.20E+05 | 3.56 | — | — | — | — |
| Citrate-2 | CA + 20 AAs + PM + 1 mM Citrate | 1.82E+04 | 5.73E+04 | 3.15 | — | — | — | — |
| Pyruvate-1 | CA + 20 AAs + PM + 1 mM Pyruvate | 1.73E+04 | 9.37E+04 | 5.42 | — | — | — | — |
| Pyruvate-2 | CA + 20 AAs + PM+ 1 mM Pyruvate | 2.22E+04 | 4.86E+03 | 0.22 | — | — | — | — |
| Fumarate-1 | CA + 20 AAs + PM + 1 mM Fumarate | 3.16E+04 | 7.20E+04 | 2.28 | — | — | — | — |
| Fumarate-2 | CA + 20 AAs + PM + 1 mM Fumarate | 1.94E+04 | 2.35E+04 | 1.21 | — | — | — | — |
| Archaeal cell-1 | CA + 20 AAs + PM + archaeal cell membrane components[h] | 1.53E+04 | 1.42E+05 | 9.27 | 81.5 | 17.5 | 0.8 | 0.3 |
| Archaeal cell-2 | CA + 20 AAs + PM + archaeal cell membrane components | 4.17E+04 | 1.05E+05 | 2.52 | — | — | — | — |

A dash indicates that data were not taken for that sample.

[a]The iTAG analysis was performed on the samples in which an increase of 16S rRNA gene copy numbers of MK-D1 about 10 times or more after incubation was observed by the qPCR assay. The detailed results are shown in Supplementary Table S2.

[b]Final concentration of Casamino acids was 0.05% (w/v).

[c]Final concentration of each amino acid was 0.1 mM.

[d]A powdered milk for baby (Hohoemi, Meiji Co., Ltd., Tokyo, Japan) was used at a final concentation of 0.1 % (w/v).

[e]The concentration of hydrogen gas was in the head space of the culture bottle.

[f]2-BES was added to inhibit methanogens.

[g]Addition of nitrate completely supressed growth of the *Ca*. P. syntrophicum. This is probably bacause nitrate inhibited fomate dehydrogenase activity of *Ca*. P. syntrophicum[78].

[h]Archaeal cell membrane components were mixture of phytol, intact polar lipid (IPL)-glycerol-dialkyl-glycerol tetraesthers (GDGTs), and core lipid (CL)-GDGTs (each at a final concentration 50 ng/ml). The reason for using the archaeal membrane components is that these have positive effect on the growth for some archaeal species: (i) archaeal cell extract including membrane lipids stimulates growth of an extremely thermophilic archaeon *Thermocaldium modestius*[79], and (ii) a hyperthermophilic archaeon *Thermofilum pendes* requires the polar lipids for the growth, which was obtained from an archaeal species of *Thermoproteus tenax*[80].
