## Supplementary Information for "Isolation of an archaeon at the prokaryote-eukaryote interface"

**Table of Contents**

**Supplementary Texts………02**
**Supplementary Methods…..05**
**Supplementary Figures……09**
**Supplementary Tables……..22**
**Supplementary References...80**

### Supplementary Texts

#### Supplementary Text 1

**Reason for using a continuous-flow bioreactor system to enrich deep marine sedimentary microorganisms and deep-sea methane seep sediment as an inoculum source.** Culture-independent molecular studies showed that deep marine sediment harbors phylogenetically diverse microorganisms, most of which belong to uncultured taxa and are distinct from those living on the Earth's surface[1-4]. Hence, their physiology and metabolic functions still remain largely unknown[5,6]. To gain insight into deep marine sedimentary microbes, they need to be cultivated, and this has been a significant challenge. However, only a small fraction of indigenous deep subseafloor microbes has been successfully isolated and characterized[7,8]. It is unclear why the cultivation of deep marine sedimentary microbes is difficult, but the batch-type cultivation techniques commonly used in previous studies may have been inadequate for this purpose. Therefore, the development of a new cultivation technique is needed. We have, therefore, employed a continuous-flow bioreactor technique for the cultivation of deep marine sedimentary microbes since 2006. The bioreactor is called a down-flow hanging sponge (DHS) reactor, which was originally developed to treat municipal sewage at a low cost in developing countries[9-11]. Specifically, a polyurethane sponge packed in the DHS reactor column provides a large surface area for microbial colonization and a longer cell residence time. As such, this type of continuous-flow reactor cultivation can provide substrates at low concentrations, similarly to those found in the natural environment. In addition, continuous-flow bioreactors allow the outflow of metabolic products that may inhibit microbial growth if accumulated. These continuous-flow reactors thereby might increase the culturability of deep marine sedimentary microorganisms in a controlled manner and serve as better sources (incubators) for microbial isolation than the original samples[12]. In fact, using DHS reactors, we have successfully enriched phylogenetically diverse microorganisms from deep marine sediments[12-15] and isolated and characterized various microorganisms using enriched microbial community from the bioreactors[16-20].

In this study, we used deep-sea methane-seep sediments collected off Kumano area, Japan. In deep-sea methane seep sediments, anaerobic oxidation of methane (AOM) reaction is the major microbial process and is mediated by a syntrophic association of euryarchaeal anaerobic methanotrophs (ANMEs) and deltaproteobacterial sulfatededucing bacteria (SRB)[21]. In addition to ANMEs and SRB, abundant and diverse microorganisms, most of which are affiliated with uncultured microbial groups of high taxonomic ranks, such as phylum, class, and order, live in methane-seep sediments[22-24]. Therefore, deep-sea methane-seep sediment can be regard as a hot spot for uncultured microorganisms. As such, cultivation and characterization of these uncultured microorganisms can greatly expand our knowledge regarding microbial physiology, genetics, and ecology. This was our rationale for using deep-sea methane seep sediment as an inoculum source for uncultured microorganism cultivation. However, in 2006, when we started the DHS bioreactor cultivation, there was extremely limited information about the metabolism of uncultured microorganisms because the metagenomic approach was not a common technique. We, therefore, could not predict the appropriate carbon and energy sources to culture uncultured microorganisms. However, we were aware that methane-seep microbial communities were sustained by methane released below the sea floor. Thus, we expected that if we provided methane as a major energy source in the DHS reactor system, the uncultured microorganisms could be cultivated from the methane-seep sediment under laboratory conditions, along with ANMEs and SRB. Indeed, using a combination of the DHS bioreactor "pre-enrichment" and subsequent *in vitro* cultivation, we have successfully cultured and isolated microorganisms representing predominant uncultured taxa. The "*Candidatus* Prometheoarchaeum syntrophicum" strain MK-D1 reported in this study is an example of cultured microorganism using our deep-sea methane seep-derived bioreactor enrichment.

**Supplementary Text 2**

**Reason for use of the four antibiotics to isolate the Lokiarchaota.** In parallel to the attempted the cultivation of microorganisms from methane-seep sediment, we tried to isolate anaerobic microorganisms from the enriched methanogenic microbial community in another DHS reactor, which was established from deep marine sediments collected off the Shimokita Peninsula, Japan[15]. During the isolation attempt, we detected few Lokiarchaeota sequences in a propionate-fed culture (Supplementary Table S8 in Imachi et al. [2011][15]). However, the Lokiarchaeota sequences became undetectable after five successive transfers. After this, we could not revive the culture any longer. However, we detected some Lokiarchaeota sequences in several anaerobic enrichment cultures supplemented with four antibiotics (i.e., ampicillin, vancomycin, kanamycin, and streptomycin, each at a final concentration of 50 µg/ml), via archaeal 16S rRNA gene-based clone analysis (data not shown). This finding suggested that Lokiarchaeota members can tolerate these antibiotics. Therefore, we added these four antibiotics into the media to serve as selective agents for the isolation of Lokiarchaeota members from the enriched AOM microbial community in the DHS reactor.

### Supplementary Methods

**Culturing.** The purity of *Ca*. P. syntrophicum strain MK-D1 was routinely examined by microscopy and iTAG analysis. In addition, the purity was verified by the whole genome shotgun sequencing, which only detected the sequences of MK-D1 and *Methanogenium* genomes. We also confirmed the culture purity based on failure to amplify the bacterial 16S rRNA gene through PCR suing the bacterial primer pairs 27F/907R[24] and EUB338F*/1492R[25-27]. Moreover, we evaluated the culture purity based on the failure of microbial growth in the following media at 10°C, 20°C, 30°C, 37°C, and 55°C: (i) thioglycolate medium (Difco); (ii) basal medium supplemented with 1 mM sucrose, 1 mM glucose, 1 mM fructose, 1 mM xylose, and 0.01% (w/v) yeast extract; and (iii) basal medium containing 5 mM lactate, 10 mM sulfate, 0.05% (w/v) CA, and 0.01% (w/v) yeast extract.

To confirm *Halodesulfovibrio* has ability to use $H_2$ and formate, we isolated the bacterium from the Lokiarchaeum enrichment culture using a roll-tube technique, with lactate (10 mM) and sulfate (10 mM), acting as an electron donor and acceptor, respectively. After isolation, we confirmed that the *Halodesulfovibrio*, designated strain MK-HDV, could grow on a hydrogen- or formate-fed medium supplemented with sulfate at 20°C.

*Methanobacterium* sp. strain MO-MB1 was previously isolated from subseafloor sediment in our laboratory as a hydrogen- and formate-utilizing methanogenic archaea[15]. *Methanogenium cariaci* strain JR1 was obtained from the Japan Collection of Microorganisms (Tsukuba, Japan) and cultured the basal medium supplemented with $H_2$ (ca. 150 kPa in head space of culture bottle) and acetate (1 mM) and yeast extract (0.01%) at 20°C

Cultures of *Halodesulfovibrio* sp. strain MK-HDV and *Methanobacterium* sp. strain MO-MB1 have been deposited in Japan Collection for Microorganisms (JCM 32479 and JCM 18473, respectively).

**Growth monitoring using qPCR.** The reaction mixture for qPCR was prepared according to the manufacturer's protocol. To construct a template standard for the primer set, we used a dilution series of the 16S rRNA gene amplicon of MK-D1, which was obtained via clone analysis using an archaeal primer pair Arch21F/1492R[28,29]. The dilution series of the PCR product was used in each qPCR analysis to calculate the 16S rRNA gene copy number. Template DNA was quantified using a Quant-iT dsDNA High-Sensitivity Assay Kit (Life Technologies). The PCR condition was as follows: initial denaturation at 95°C for 30 s, followed by 40 cycles of denaturation at 95°C for 10 s, annealing at 58°C for 30 s, and extension at 72°C for 31 s. The annealing temperature was optimized empirically through the amplification of the 16S rRNA gene of MK-D1. To verify the specificity of the qPCR assay, we performed two types of experiments. First, a melting-curve analysis was performed for every qPCR assay. Second, the PCR product size was confirmed by gel electrophoresis and subsequent clone library analysis. The clone library experiment was performed only for three DNA samples, which were obtained from two- and three-successive transferred Lokiarchaeota enrichment cultures and the DHS bioreactor enrichment. We confirmed that all the PCR products showed the expected PCR amplicon size (i.e., about 390 bp) and all retrieved clones were identical to MK-D1 or were affiliated with the candidate phylum Lokiarchaeota (16 clones were randomly collected from each library).

**FISH.** MK-D1-specific probes were designed using the probe design tool of the ARB program[30]. The specificity of the probes was confirmed using the BLAST and the ARB-SILVA databases[31]. The $\Delta G^0_{overall}$ values of the probes and target MK-D1 16S rRNA sequence were calculated using the mathFISH web server[32]. Both probes exhibited high hybridization efficiencies (-15.5 kcal/mol for DSAG-Gr2-1142, -14.4 kcal/mol for DSAG-Gr2-1432).

**Growth test using multiple substrates.** A highly purified culture of MK-D1 was inoculated in the medium (15% inoculum, v/v). Then, 1 ml of the culture liquid was immediately taken from each culture to examine the initial 16S rRNA gene copy numbers of MK-D1 using the qPCR technique. The liquid culture samples for qPCR analysis were stored at -80°C until further processing. After the sampling, all the cultures were incubated at 20°C for 120 days in the dark without shaking. After 120 days of incubation, 1 ml of liquid culture was sampled from each culture vial to quantify the final 16S rRNA gene copy numbers of MK-D1. DNA extraction and qPCR analysis were performed, as the methods mentioned in the Methods section. In the qPCR assay, samples taken from the same culture vial (e.g., 0-day and 120-day samples of H2-1 culture) were applied to the same PCR plate for accurate quantification. To confirm microbial community structure after incubation, iTAG analysis was performed on the samples, which showed an increase of about 10-fold or more in the 16S rRNA gene copy numbers of MK-D1 after incubation, as observed by the qPCR assay.

**Stable isotope probing incubation and NanoSIMS analysis.** During incubation, 5 ml of culture liquid was taken from the vials once every 30 days, 1 ml of which was used for qPCR and iTAG analyses, and the remaining 4 ml was processed for NanoSIMS analysis. The samples for NanoSIMS analysis were fixed in 2% PFA under anaerobic condition for approximately 2 h. The fixed cells were washed twice in PBS and stored in a 1:1 mixture of PBS and ethanol at -20°C until further processing.

Due to the low-biomass sample, the fixed cells were concentrated in a small analysis area (0.5–1 $mm^2$) of indium tin oxide-coated polycarbonate membranes using a fluorescence-activated cell sorting[33]. Microbial cells on membranes were stained with SYBR Green I and observed with an epifluorescence microscope (BX-51, Olympus) prior to NanoSIMS analysis.

Samples were analyzed by raster ion imaging with a CAMECA NanoSIMS 50L at the Kochi Institute for Core Sample Research, JAMSTEC. A focused primary $Cs^+$ beam of ~1 pA for carbon and nitrogen isotopic analysis was rastered over 30 × 30 $\mu m^2$ areas on the samples. Negative secondary ions of $^{12}C$ (EM#1), $^{13}C$ (EM#2), $^{16}O$ (EM#4), $^{12}C^{14}N$ (EM#5), $^{12}C^{15}N$ (EM#6), and $^{32}S$ (EM#7) were measured using six electron multipliers (EMs) in a multi-detection mode at a high mass resolving power of ~7,000 (CAMECA NanoSIMS definition), which is sufficient to separate all relevant isobaric interferences (i.e., $^{13}C$ on $^{12}C^{1}H$, $^{12}C^{14}N$ on $^{13}C^{++}$, and $^{12}C^{14}N^{1}H$). Each run was initiated after the stabilization of the secondary ion beam intensity following a pre-sputtering of approximately 2 min with a strong primary ion beam current. The same area was repeatedly scanned (20–30 times) in each run, with individual images consisting of 256 × 256 pixels and with a dwell time of 2,000 µs. The total acquisition time was approximately 1 h.

The recorded images and data were analyzed using IDL based NASA JSC imaging software for NanoSIMS[34], OpenMIMS (https://github.com/BWHCNI/OpenMIMS) and Look@NanoSIMS[35]. The images were corrected for quasi-simultaneous arrival effect and detector dead time. Different scans of each image were aligned to correct image drift during acquisition. The final images were generated by adding the secondary ion counts of each recorded secondary ion from each pixel for all scans.

### Supplementary Figures

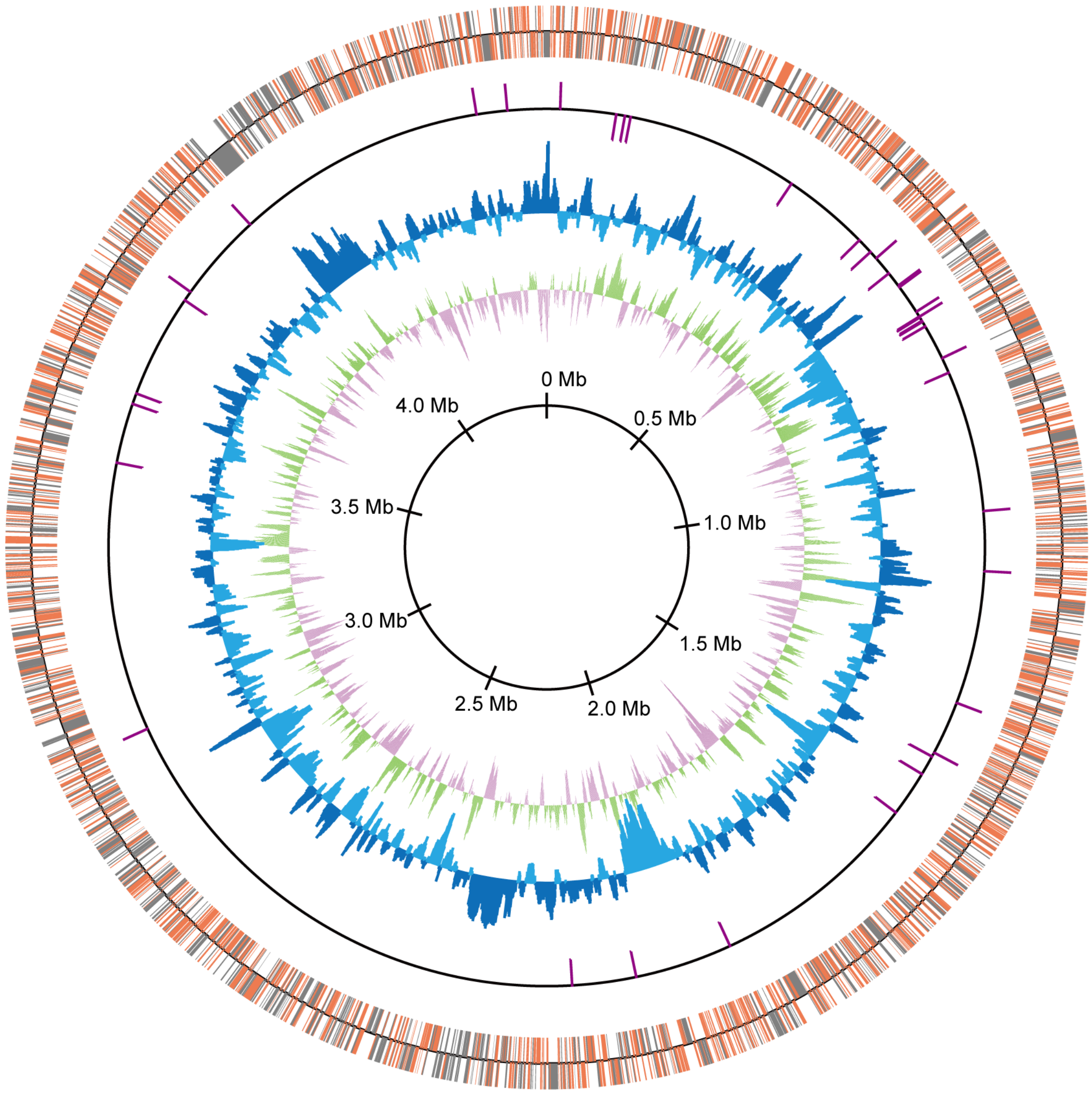


**Supplementary Figure S1 | Circular representation of the *Ca.* Prometheoarchaeum syntrophicum strain MK-D1 genome.** From the outside to the center: the distribution of the CDSs based on the conserved (orange) or non-conserved (gray) genes in the first circle, non-coding RNAs in the second circle, GC content showing deviation from average (40.7%) in the third circle, and GC skew in the fourth circle. The GC content and GC skew were calculated using a sliding window of 2 kb in step of 10 kb. The CDS and RNA genes illustrate the findings for plus and minus strands.

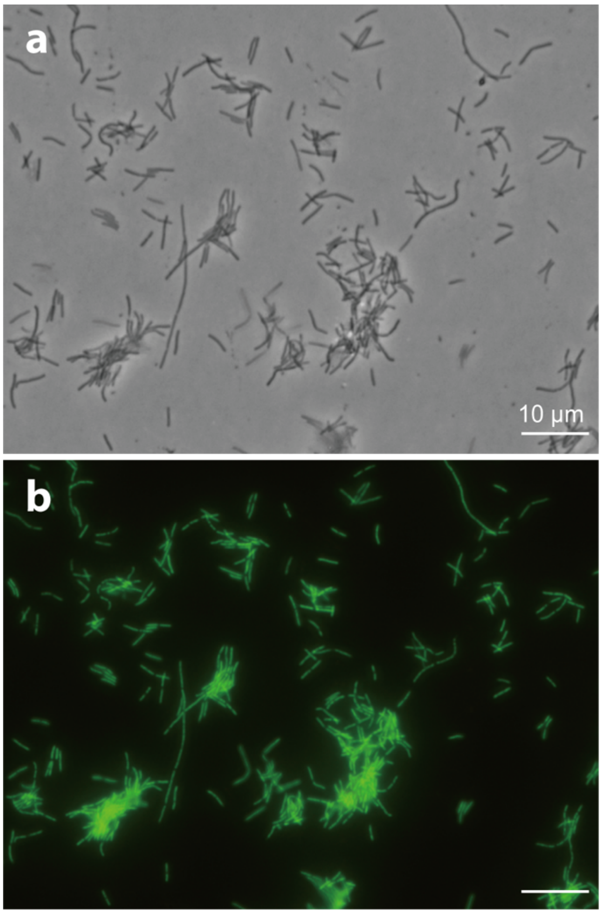


**Supplementary Figure S2 | Photomicrographs of *Methanobacterium* sp. strain MO-MB1 cells stained with SYBR Green I.** Phase-contrast (**a**) and fluorescence (**b**) micrographs of the same field are shown. Cells of strain MO-MB1 were fixed with 2% PFA after culturing with the basal medium under the optimal cultivation condition (i.e., supplementation with $H_2$ [ca. 150 kPa in head space of culture bottle] and acetate [1 mM] and yeast extract [0.01%], and incubation temperature 30°C). All the *Methanobacterium* cells were stained well with SYBR Green I. The weak fluorescence intensities of some *Methanobacterium* cells in Fig. 2b were likely due to increased rigidity and reduced permeability of their membranes incurred from growth under low hydrogen concentrations (*c.f.*, Nakamura et al. [2006][36]). Bars, 10 µm.

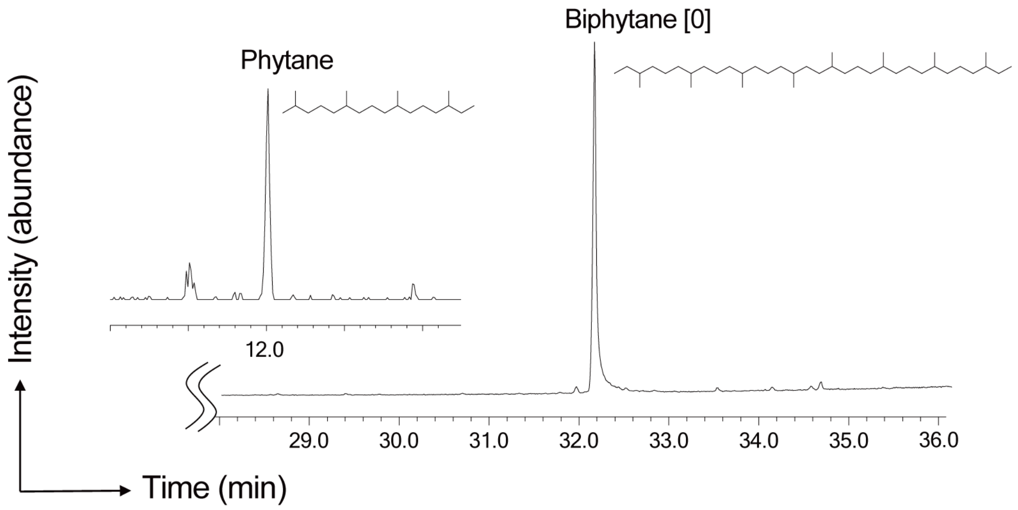


**Supplementary Figure S3 |** A total ion chromatogram of gas chromatography/mass spectrometry (GC/MS) for lipid obtained from *Methanogenium cariac*i strain JR1 (JCM 10550). The chemical structures of isoprenoid lipids are also shown.

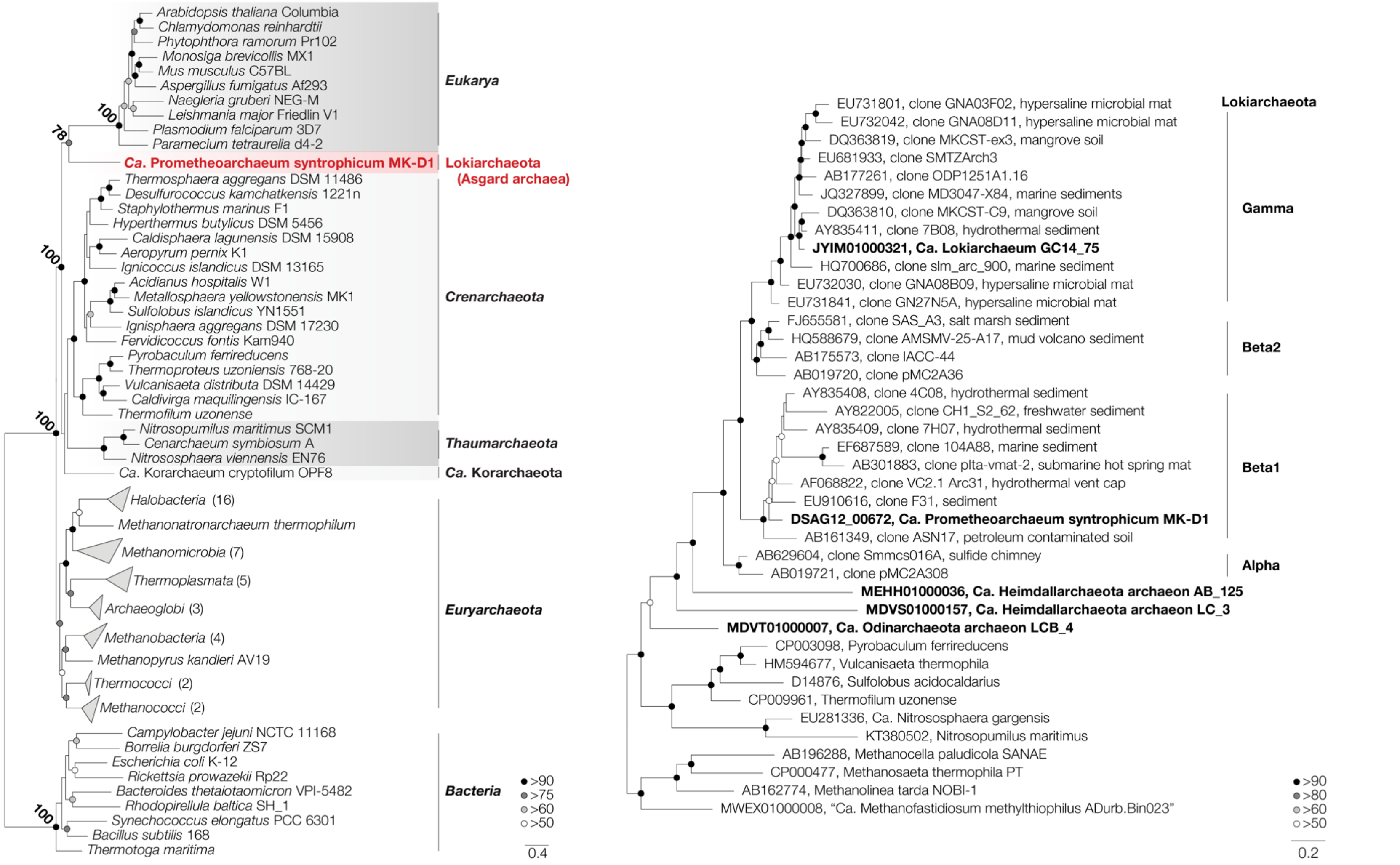


**Supplementary Figure S4 | Ribosomal protein-(left) and 16S rRNA gene-(right) based phylogeny of MK-D1.** (see next page)

**Supplementary Figure S4 (cont.) | Ribosomal protein- (left) and 16S ribosomal RNA gene- (right) based phylogeny of MK-D1.** Phylogenomic tree of MK-D1 and select cultured archaea, eukaryotes, and bacteria based on 31 ribosomal proteins conserved across the three domains (Supplementary Table S4). Ribosomal protein sequences were collected from MK-D1, the organisms shown in the tree, and metagenome-assembled genomes (MAGs) of uncultured archaeal lineages (Supplementary Table S5) and aligned individually using MAFFT (--linsi). MAG-derived sequences were included to generate an alignment that maximizes the archaeal diversity taken into account, but removed for subsequent tree construction to avoid any influence of contamination (*i.e.*, concatenation of sequences that do not belong to the same organism). *Ca.* Korarchaeum sequences were kept due to its critical phylogenetic position in TACK phylogeny. After removing all-gap positions and concatenation, the maximum likelihood tree was constructed using RAxML-NG (fixed empirical substitution matrix [LG], 4 discrete GAMMA categories, empirical AA frequencies, and 100 bootstrap replicates). Bootstrap values around critical branching points are also shown. Phylogenetic tree of MK-D1 and related archaea based on 16S rRNA genes. 16S rRNA gene sequences were aligned using SINA[37] against the Silva v132 alignment[31]. The maximum likelihood tree was calculated using RAxML[38] using the generalized time reversible (GTR) model, 4 discrete GAMMA categories, generalized time reversible model, and 100 bootstrap replicates.

283

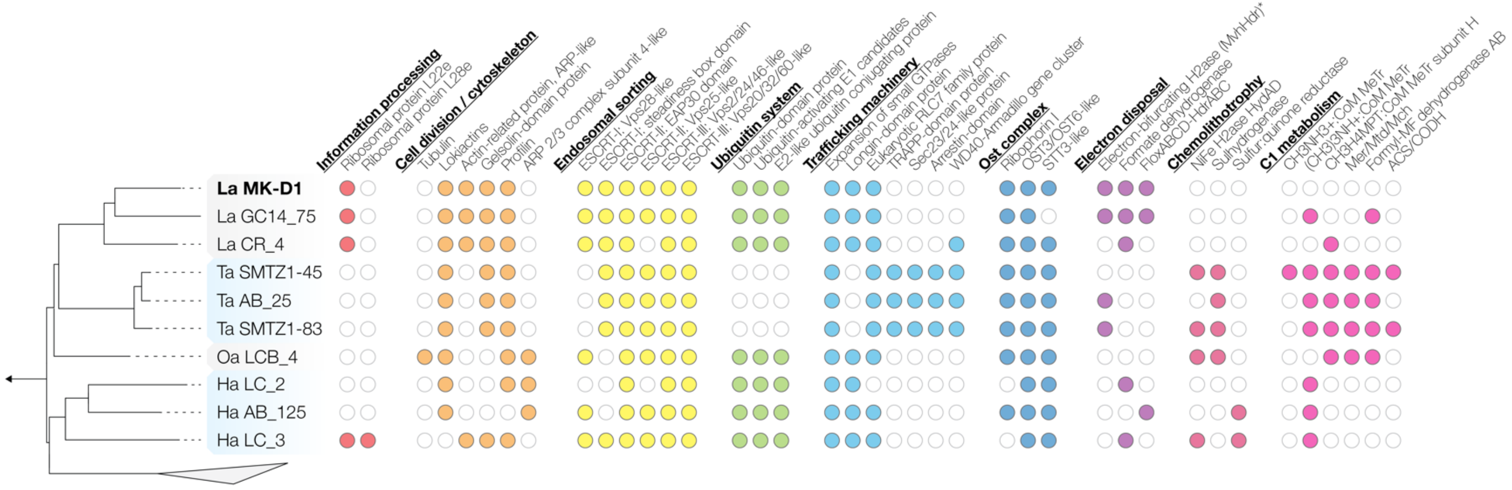


**Supplementary Figure S5 | Eukaryotic signature proteins and anaerobic metabolism of select Asgard archaea.** Asgard archaea candidate phyla are abbreviated as follows: Lokiarchaeota (La), Thorarchaeota (Ta), Odinarchaeota (Oa), and Heimdallarchaeota (Ha).

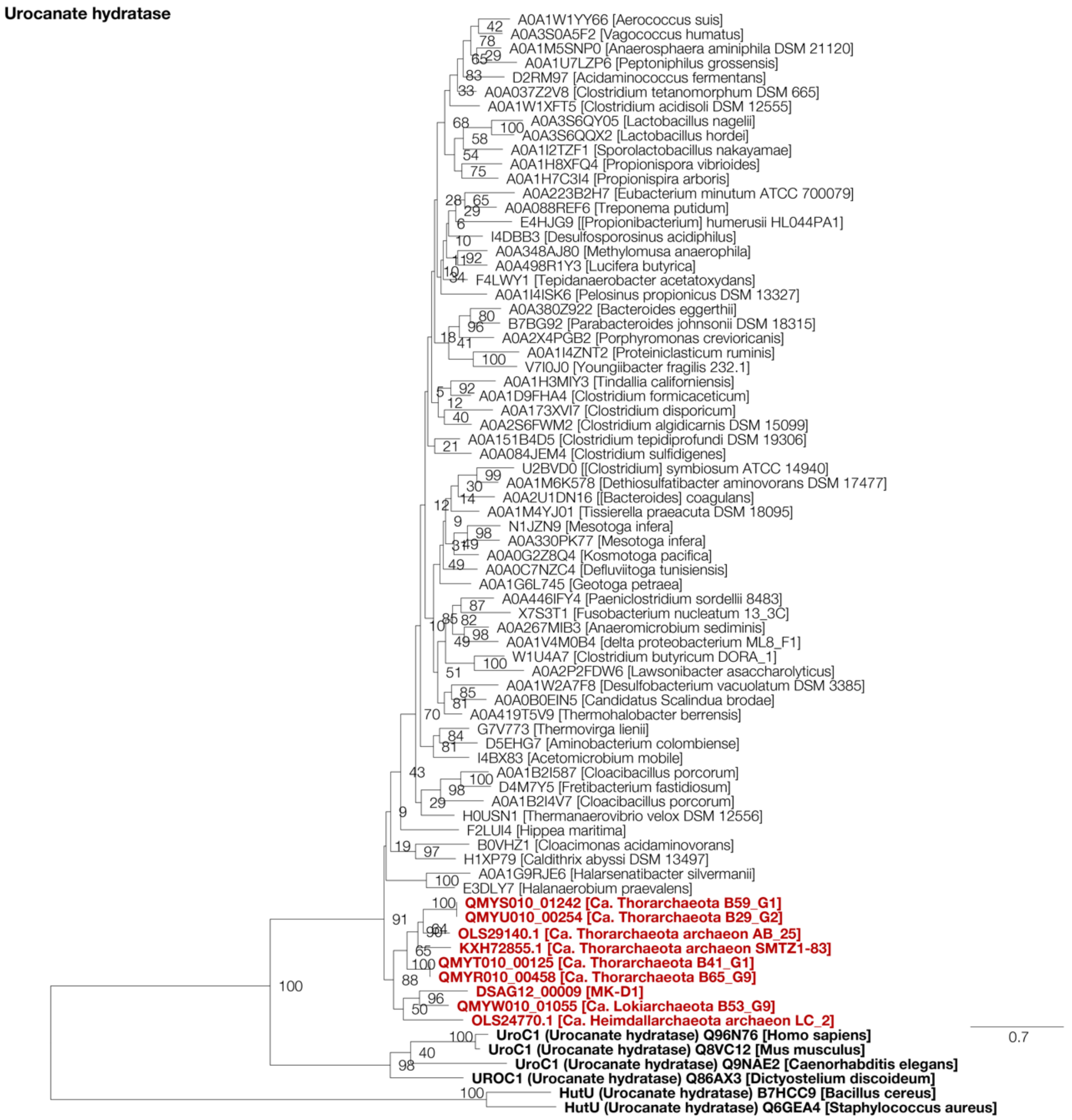


**Supplementary Figure S6 | Maximum likelihood tree of Asgard archaea urocanate hydratase (HutU).** HutU homologs were collected through BLASTp analysis of the Asgard archaea sequences against the UniProt database (release 2019_06). Of homologs with sequence similarity ≥40% and overlap ≥70%, representative sequences were selected using CD-HIT with a clustering cutoff of 70% similarity (default settings otherwise). Additional homologs with verified biochemical activity, sequence similarity ≥30%, and overlap ≥70% were collected through BLASTp analysis of the Asgard archaea sequences against the UniProt/SwissProt database. Sequences were aligned using MAFFT v7[39] with default settings and trimmed using trimAl[40] with default settings. The phylogenetic tree was constructed using RAxML-NG[38] using fixed empirical substitution matrix (LG), 4 discrete GAMMA categories, empirical amino acid frequencies from the alignment, and 100 bootstrap replicates.

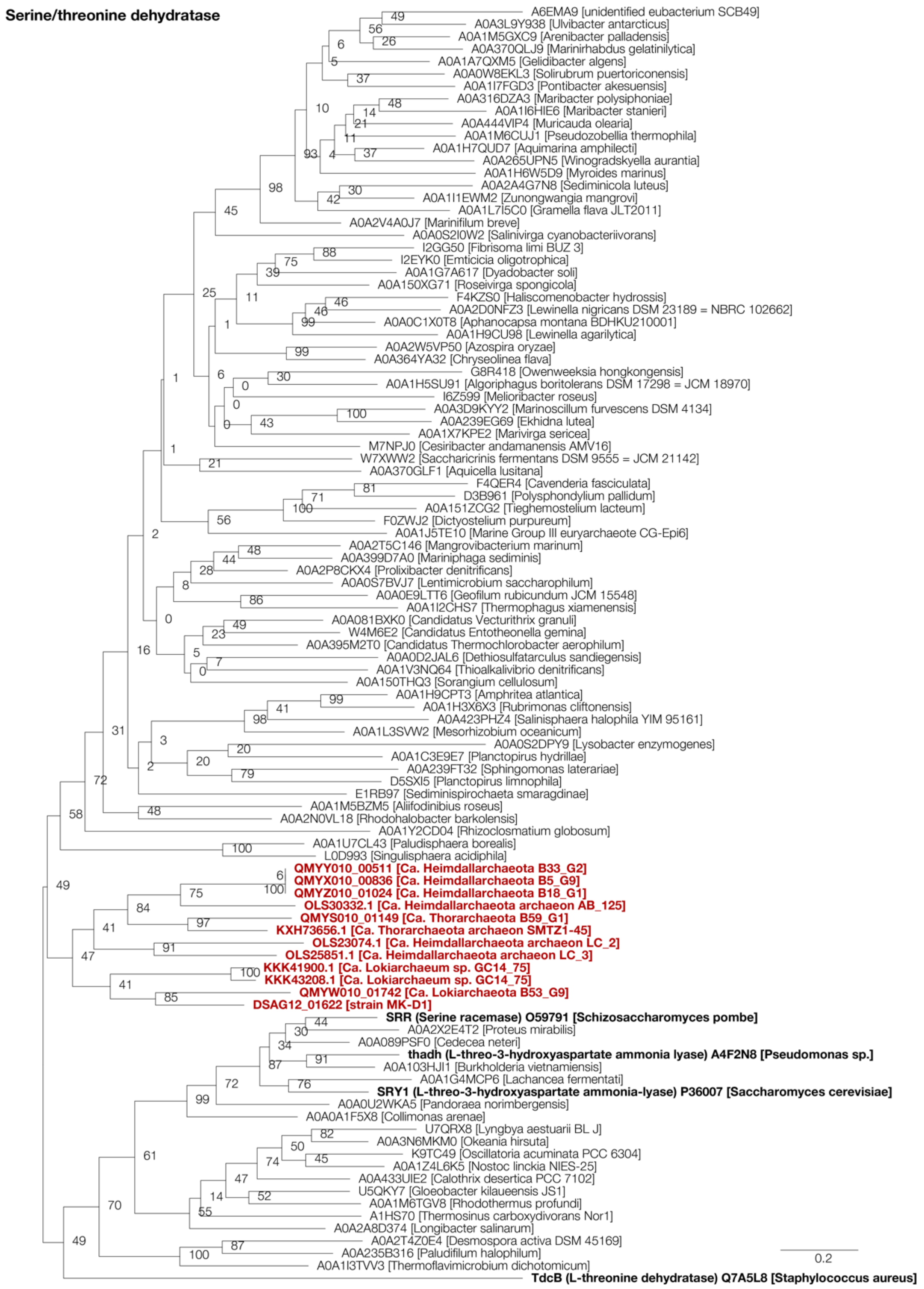


**Supplementary Figure S7 | Maximum likelihood tree of Asgard archaea L-threonine/L-serine dehydratase (TdcB).** See Fig S4 caption for details.

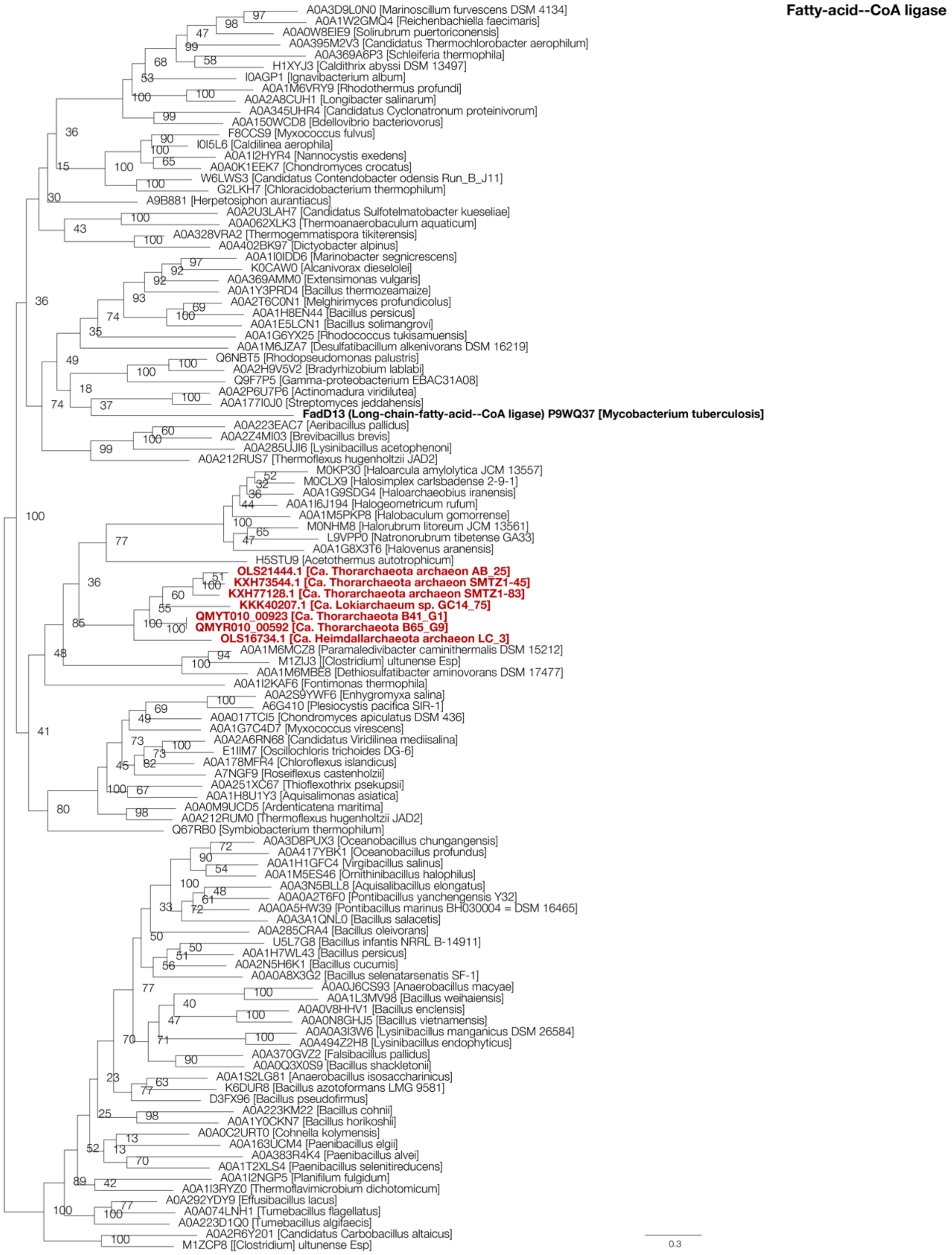


**Supplementary Figure S8 | Maximum likelihood tree of Asgard archaea fatty-acid--CoA ligase.** Although the closest characterized relative of the Asgard archaea genes is a long-chain-fatty-acid--CoA ligase, the preferred substrate remains unclear as fatty-acid--CoA ligases have broad/diverse substrate specificities. See Fig S4 caption for details.

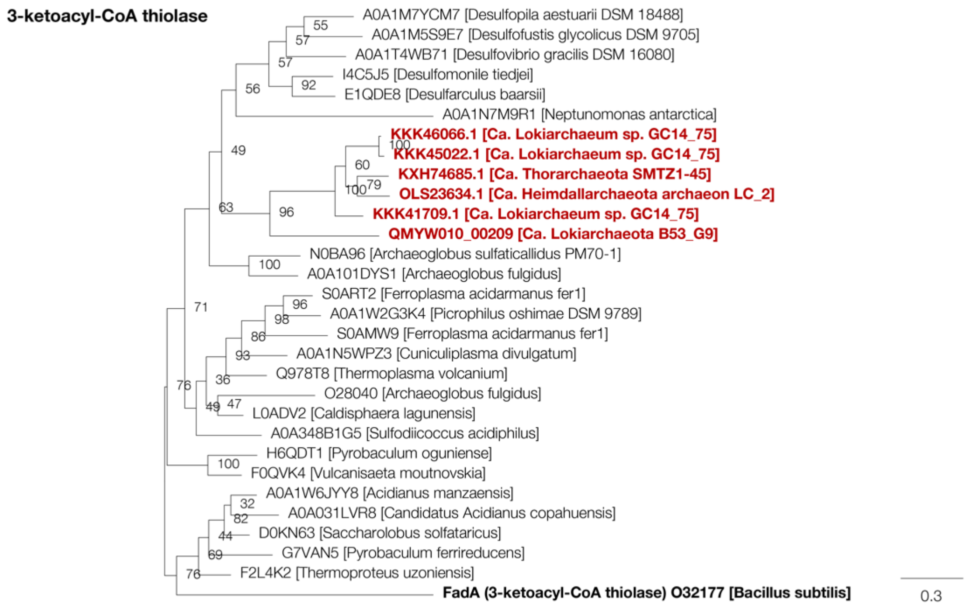


**Supplementary Figure S9 | Maximum likelihood tree of Asgard archaea 3-ketoacyl-CoA thiolase (FadA).** See Fig S4 caption for details.

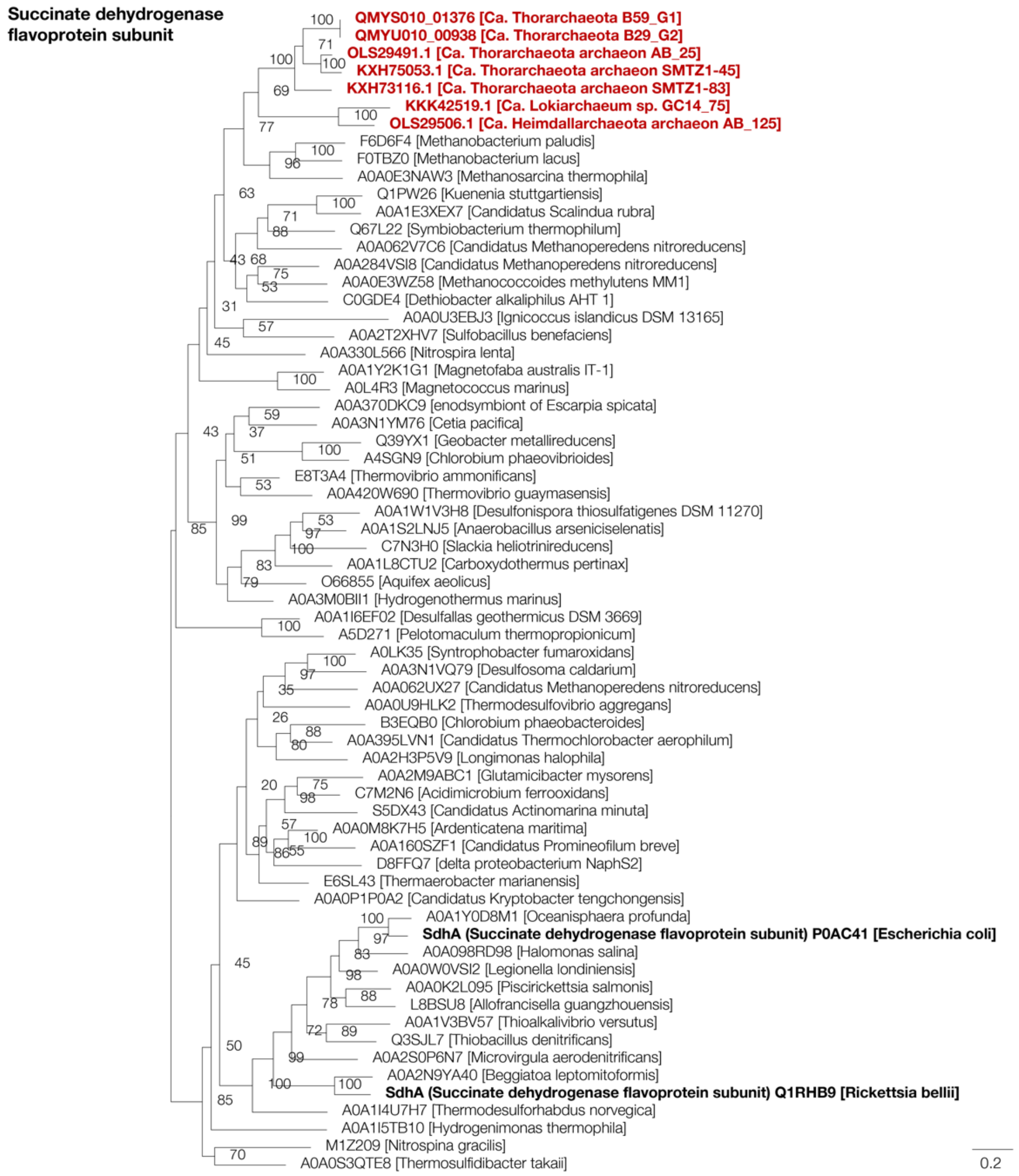


**Supplementary Figure S10 | Maximum likelihood tree of Asgard archaea succinate dehydrogenase flavoprotein subunit (SdhA).** See Fig S4 caption for details.

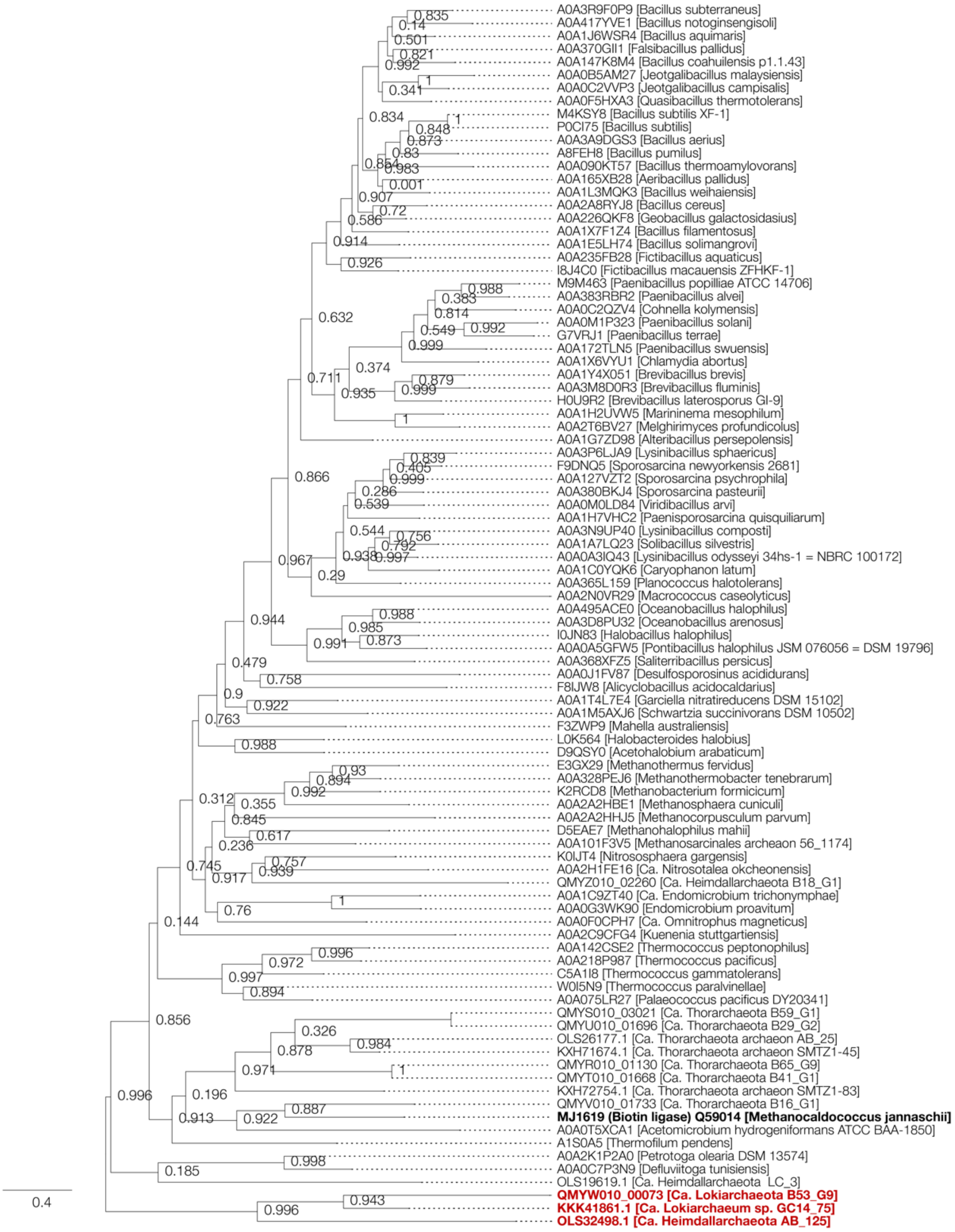


**Supplementary Figure S11 | Maximum likelihood tree of Asgard archaea biotin ligase (BirA).** BirA homologs were collected through BLASTp analysis of the Asgard archaea sequences against the UniProt database (release 2019_06). Of homologs with sequence similarity ≥40% and overlap ≥70%, representative sequences were selected using CD-HIT with a clustering cutoff of 70% similarity (default settings otherwise). Additional homologs with verified biochemical activity, sequence similarity ≥30%, and overlap ≥70% were collected through BLASTp analysis of the Asgard archaea sequences against the UniProt/SwissProt database. Sequences were aligned using MAFFT v7[39]with default settings and trimmed using trimAl[40] with default settings. The phylogenetic tree was constructed using FastTree[41] using fixed empirical substitution matrix (LG) and 1000 bootstrap replicates.

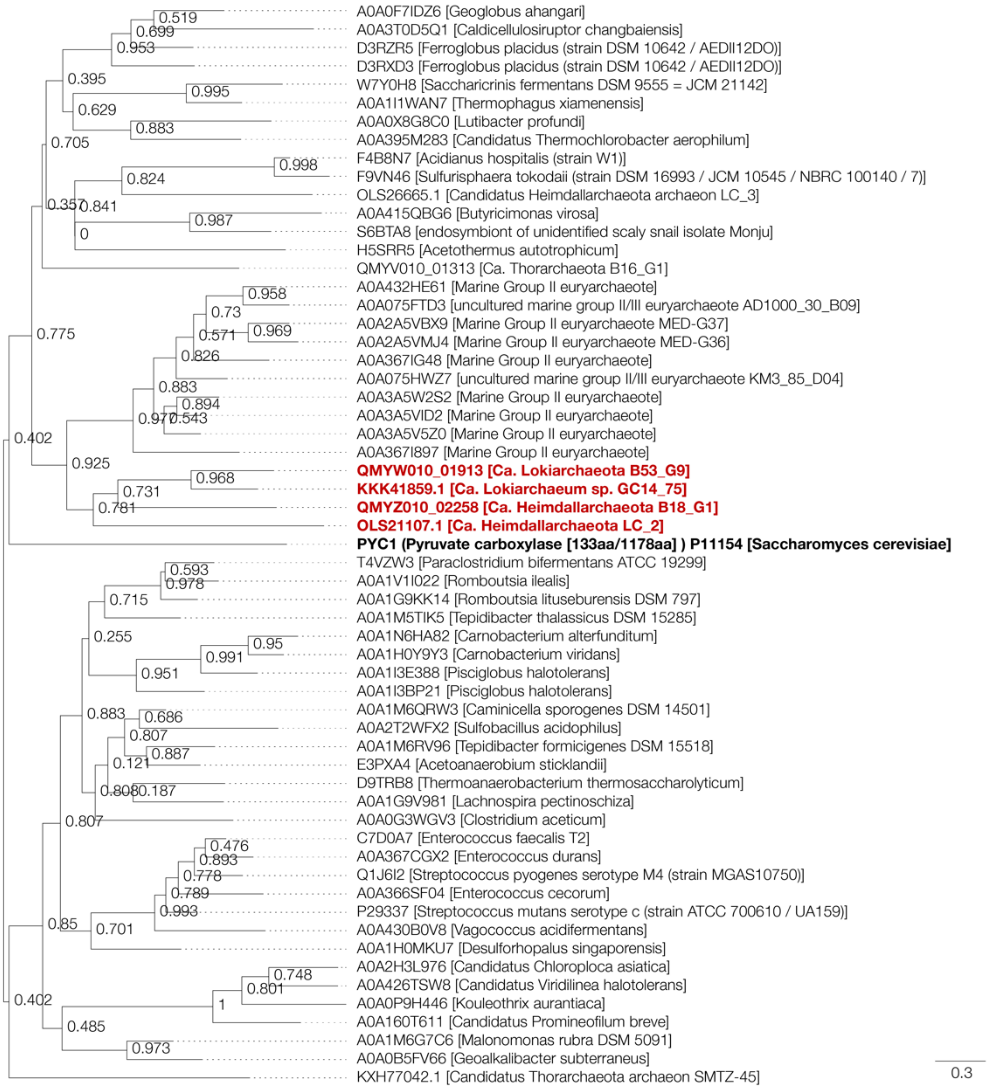


**Supplementary Figure S12 | Maximum likelihood tree of Asgard archaea transcarboxylase biotin carboxyl carrier protein.** See Fig S9 caption for details.

**Supplementary Table S1 | Genome information of *Ca*. P. syntrophicum, *Halodesulfovibrio* and *Methanogenium* cultured in this study.**

| | *Ca*. P. syntrophicum MK-D1 | *Halodesulfovibrio* MK-HDV | *Methanogenium* MK-MG |
|---|---|---|---|
| Size (bp) | 4,427,796 | 4,171,812 | 2,331,727 |
| Contigs | 1 | 3 | 13 |
| Coverage (times) | 52 | 189 | 66 |
| G+C content (%) | 31.1 | 44.28 | 49.59 |
| Number of ORF | 3946 | 3594 | 2294 |
| rRNAs | 2 | 7 | 16 |
| tRNAs | 24 | 100 | 49 |
| Genome completeness (%)[a] | 100 | 99.38 | 98.37 |
| Contamination (%)[a] | 0 | 0.59 | 1.96 |

[a]Genome completeness and contamination were estimated using CheckM (Parks, D. H., Imelfort, M., Skennerton, C. T., Hugenholtz, P. & Tyson, G. W. CheckM: assessing the quality of microbial genomes recovered from isolates, single cells, and metagenomes. *Genome Res*. **25,** 1043–1055 [2015]).

**Supplementary Table 2 | iTAG analysis of cultures used for characterization of MK-D1.** Sequencing was performed on a MiSeq platform using universal primer pair 530F and 907R. OTUs that have only two or fewer tag-sequences were not included in the microbial community analysis of the MK-D1 cultures.

**<Fig. 1b, and Extended Data Fig. 2a (FISH image for a tri-culture)>**
Accession number: DRR184081

| Phylogenetic group | Sequence read number | Relative abundance (%) |
|---|---|---|
| Bacteria, Halodesulfovibrio | 66871 | 76.9 |
| Archaea, Lokiarchaeota (*Ca*. P. syntrophicum) | 12502 | 14.4 |
| Archaea, Methanogenium | 7398 | 8.5 |
| Archaea, Methanococcoides | 179 | 0.2 |
| Bacteria, Dehalococcoidia GIF9 group | 12 | 0.01 |
| Total | 86962 | |

**<Fig. 1c, d, and Extended Data Fig. 2b, c (FISH image for a co-culture)>**
Accession number: DRR184082

| Phylogenetic group | Sequence read number | Relative abundance (%) |
|---|---|---|
| Archaea, Lokiarchaeota (*Ca*. P. syntrophicum) | 112611 | 87.5 |
| Archaea, Methanogenium | 15458 | 12.0 |
| Archaea, Methanobacterium | 612 | 0.5 |
| Total | 128681 | |

**<Fig. 1e>**
Accession number: DRR184083

| Phylogenetic group | Sequence read number | Relative abundance (%) |
|---|---|---|
| Archaea, Lokiarchaeota (*Ca*. P. syntrophicum) | 72338 | 75.8 |
| Archaea, Methanogenium | 23120 | 24.2 |
| Bacteria, Sphingomonas | 16 | 0.02 |
| Bacteria, Atribacteria | 15 | 0.02 |
| Bacteria, Lactobacillus | 5 | 0.01 |
| Total | 95494 | |

**<Fig. 2b–e, and Extended Data Table 2 (NanoSIMS analysis, co-culture with *Methanobacterium*)>**
Accession number: DRR184084

| Phylogenetic group | Sequence read number | Relative abundance (%) |
|---|---|---|
| Archaea, Lokiarchaeota (*Ca*. P. syntrophicum) | 87664 | 98.9 |
| Archaea, Methanobacterium | 954 | 1.1 |
| Archaea, Methanococcoides | 14 | 0.02 |
| Bacteria, Halodesulfovibrio | 10 | 0.01 |
| Bacteria, Dehalococcoidia GIF9 group | 9 | 0.01 |
| Total | 88651 | |

**<Extended Data Table 2 (tri-culture with *Halodesulfovibrio* and *Methanogenium*)>**
Accession number: DRR184085

| Phylogenetic group | Sequence read number | Relative abundance (%) |
|---|---|---|
| Bacteria, Halodesulfovibrio | 67531 | 77.7 |
| Archaea, Methanogenium | 20972 | 24.1 |
| Archaea, Lokiarchaeota (*Ca*. P. syntrophicum) | 15774 | 18.1 |
| Archaea, Methanococcoides | 705 | 0.8 |
| Archaea, Methanobacterium | 13 | 0.01 |
| Total | 104995 | |

**<Fig. 3a, b, and Extended Data Fig. 2d (SEM images)>**
Accession number: DRR184086

| Phylogenetic group | Sequence read number | Relative abundance (%) |
|---|---|---|
| Archaea, Lokiarchaeota (*Ca*. P. syntrophicum) | 115119 | 89.26 |
| Archaea, Methanogenium | 13846 | 10.74 |
| Total | 128965 | |

**<Fig. 3c, f, i and Extended Data Fig. 2e, h, i. l (SEM and TEM images)>**
Accession number: DRR184087

| Phylogenetic group | Sequence read number | Relative abundance (%) |
|---|---|---|
| Archaea, Lokiarchaeota (*Ca*. P. syntrophicum) | 130923 | 87.0 |
| Archaea, Methanogenium | 19496 | 13.0 |
| Bacteria, Phenylobacterium | 14 | 0.01 |
| Total | 150433 | |

**<Fig. 3d, e, and Extended Data Fig. 2f, g (Cryo-EM images)>**
Accession number: DRR184088

| Phylogenetic group | Sequence read number | Relative abundance (%) |
|---|---|---|
| Archaea, Lokiarchaeota (*Ca*. P. syntrophicum) | 86292 | 84.6 |
| Archaea, Methanogenium | 15647 | 15.3 |
| Archaea, Methanobacterium | 82 | 0.1 |
| Bacteria, Dehalococcoidia GIF9 group | 19 | 0.02 |
| Bacteria, Halodesulfovibrio | 8 | 0.01 |
| Bacteria, Lysinibacillus | 7 | 0.01 |
| Total | 102055 | |

**<Fig. 3g, h and Extended Data Fig. 2j, k (SEM images)>**
Accession number: DRR184089

| Phylogenetic group | Sequence read number | Relative abundance (%) |
|---|---|---|
| Archaea, Lokiarchaeota (*Ca*. P. syntrophicum) | 104002 | 90.0 |
| Archaea, Methanogenium | 11610 | 10.0 |
| Total | 115612 | |

**<Fig. 3j (Lipid analysis)>**
Accession number: DRR184090

| Phylogenetic group | Sequence read number | Relative abundance (%) |
|---|---|---|
| Archaea, Lokiarchaeota (*Ca*. P. syntrophicum) | 163886 | 91.5 |
| Archaea, Methanogenium | 15257 | 8.5 |
| Bacteria, Sphingomonas | 22 | 0.01 |
| Total | 179165 | |

**<Extended Data Fig. 3a (Amino acids concentrations at 90 days)>**
Accession number: DRR184091

| Phylogenetic group | Sequence read number | Relative abundance (%) |
|---|---|---|
| Archaea, Lokiarchaeota (*Ca*. P. syntrophicum) | 109046 | 86.8 |
| Archaea, Methanogenium | 16580 | 13.2 |
| Total | 125626 | |

**<Extended Data Fig. 3b (Amino acids concentrations at 90 days)>**
Accession number: DRR184092

| Phylogenetic group | Sequence read number | Relative abundance (%) |
|---|---|---|
| Archaea, Lokiarchaeota (*Ca*. P. syntrophicum) | 126553 | 91.0 |
| Archaea, Methanogenium | 12539 | 9.0 |
| Total | 139092 | |

**<Extended Data Table 3 (Inoculum)>**
Accession number: DRR184093

| Phylogenetic group | Sequence read number | Relative abundance (%) |
|---|---|---|
| Archaea, Lokiarchaeota (*Ca*. P. syntrophicum) | 78508 | 39.8 |
| Archaea, Methanogenium | 72659 | 36.8 |
| Archaea, Methanobacterium | 46017 | 23.3 |
| Bacteria, Massilia | 10 | 0.005 |
| Bacteria, Bacillus | 4 | 0.002 |
| Bacteria, Paenibacillus | 3 | 0.002 |
| Bacteria, Pseudomonas | 3 | 0.002 |
| Total | 197204 | |

**<Extended Data Table 3 (Control-1)>**
Accession number: DRR184094

| Phylogenetic group | Sequence read number | Relative abundance (%) |
|---|---|---|
| Archaea, Lokiarchaeota (*Ca*. P. syntrophicum) | 140431 | 76.7 |
| Archaea, Methanogenium | 39929 | 21.8 |
| Archaea, Methanobacterium | 2655 | 1.4 |
| Bacteria, Veillonellaceae | 60 | 0.03 |
| Total | 183075 | |

**<Extended Data Table 3 (Control-2)>**
Accession number: DRR184095

| Phylogenetic group | Sequence read number | Relative abundance (%) |
|---|---|---|
| Archaea, Lokiarchaeota (*Ca*. P. syntrophicum) | 70273 | 60.3 |
| Archaea, Methanogenium | 44282 | 38.0 |
| Archaea, Methanobacterium | 1906 | 1.6 |
| Bacteria, Hydrogenophilus | 25 | 0.02 |
| Bacteria, Deinococcus | 19 | 0.02 |
| Bacteria, Propionibacterium | 4 | 0.003 |
| Archaea, Thaumarchaeota | 3 | 0.003 |
| Total | 116512 | |

**<Extended Data Table 3 (Sulfate-1)>**
Accession number: DRR184096

| Phylogenetic group | Sequence read number | Relative abundance (%) |
|---|---|---|
| Archaea, Lokiarchaeota (*Ca*. P. syntrophicum) | 84925 | 79.5 |
| Archaea, Methanogenium | 20854 | 19.5 |
| Archaea, Methanobacterium | 1017 | 1.0 |
| Archaea, Thaumarchaeota | 14 | 0.01 |
| Bacteria, Oceanospirillales | 7 | 0.01 |
| Archaea, Marine Group II | 5 | 0.005 |
| Bacteria, Veillonellaceae | 5 | 0.005 |
| Total | 106827 | |

**<Extended Data Table 3 (Glucose-1)>**
Accession number: DRR184097

| Phylogenetic group | Sequence read number | Relative abundance (%) |
|---|---|---|
| Archaea, Lokiarchaeota (*Ca*. P. syntrophicum) | 118208 | 73.8 |
| Archaea, Methanogenium | 38864 | 24.3 |
| Archaea, Methanobacterium | 2998 | 1.9 |
| Bacteria, Sedimentibacter | 30 | 0.02 |
| Archaea, Thaumarchaeota | 9 | 0.01 |
| Total | 160109 | |

**<Extended Data Table 3 (Glucose-2)>**
Accession number: DRR184098

| Phylogenetic group | Sequence read number | Relative abundance (%) |
|---|---|---|
| Archaea, Lokiarchaeota (*Ca*. P. syntrophicum) | 85521 | 70.3 |
| Archaea, Methanogenium | 34100 | 28.0 |
| Archaea, Methanobacterium | 2010 | 1.7 |
| Total | 121631 | |

**<Extended Data Table 3 (Xylose-2)>**
Accession number: DRR184099

| Phylogenetic group | Sequence read number | Relative abundance (%) |
|---|---|---|
| Archaea, Lokiarchaeota (*Ca*. P. syntrophicum) | 92052 | 61.4 |
| Archaea, Methanogenium | 54674 | 36.5 |
| Archaea, Methanobacterium | 3128 | 2.1 |
| Bacteria, Sphingomonas | 15 | 0.01 |
| Total | 149869 | |

**<Extended Data Table 3 (Archaeal cell-1)>**
Accession number: DRR184100

| Phylogenetic group | Sequence read number | Relative abundance (%) |
|---|---|---|
| Archaea, Lokiarchaeota (*Ca*. P. syntrophicum) | 115040 | 81.5 |
| Archaea, Methanogenium | 24713 | 17.5 |
| Archaea, Methanobacterium | 1064 | 0.8 |
| Bacteria, Afipia | 391 | 0.3 |
| Total | 141208 | |

**<Pure co-culture>**
Accession number: DRR184101

| Phylogenetic group | Sequence read number | Relative abundance (%) |
|---|---|---|
| Archaea, Lokiarchaeota (*Ca*. P. syntrophicum) | 95965 | 91.4 |
| Archaea, Methanogenium | 9049 | 8.6 |
| Total | 105014 | |

No bacterial PCR amplicons and no bacterial growth were observed from the culture. The purity check of culture is described in the Supplementary Methods.

**Supplementary Table S3 | DNA probes used in this study.**

| Probe | Target group | Probe sequence (5' to 3')[a] | Labeling | %FA[b] | Reference |
|---|---|---|---|---|---|
| **Standard-FISH** | | | | | |
| DSAG-Gr2-1142[c] | *Ca*. P. syntrophicum strain MK-D1 and its relatives | CAGTCCGCTTAGCGTTCC | Alexa Fluor 488 | 35 | This study |
| MBGB-525[c,d] | Almost all members of Lokiarchaeota | AGAGCTGGTTTTACCGCG | Alexa Fluor 488 | 10 | Knittel *et al*., 2005 |
| EUB338 | Most *Bacteria* | GCTGCCTCCCGTAGGAGT | Alexa Fluor 555 | 20 | Amann *et al*., 1990 |
| ARC915 | Most *Archaea* | GTGCTCCCCCGCCAATTCCT | Alexa Fluor 555 | 35 | Stahl *et al*., 1991 |
| ***in situ* DNA-HCR-FISH** | | | | | |
| DSAG-Gr2-1142-initiatorH | *Ca*. P. syntrophicum strain MK-D1 and its relatives | CCGAATACAAAGCATCAACGACTAGAAAAAACAGTCCGCTTAGCGTTCC | — | 35 | This study |
| DSAG-Gr2-1432-initiatorH[e] | *Ca*. P. syntrophicum strain MK-D1 | CCGAATACAAAGCATCAACGACTAGAAAAAACGACCCTTAGGACCGTTTTTC | — | 30 | This study |
| EUB338-initiatorC | Most *Bacteria* | CCAGTTATCAGTAGTCCGTCCTTCATTTTTTGCTGCCTCCCGTAGGAGT | — | 20 | Yamaguchi *et al*., 2015 |
| MG1200-initiatorC | most *Methanomicrobiales* (including *Methanogenium*) | CCAGTTATCAGTAGTCCGTCCTTCATTTTTTCGGATAATTCGGGGCATGCTG | — | 10[f] | Raskin *et al*., 1994. |
| ARC915-initiatorC | Most *Archaea* | CCAGTTATCAGTAGTCCGTCCTTCATTTTTTGTGCTCCCCCGCCAATTCCT | — | 35 | Yamaguchi *et al*., 2015 |
| **Amplifier probes** | | | | | |
| H1 | — | TCTAGTCGTTgatgctttgtattcggCGACAGATAAccgaatacaaagcatc | Alexa Fluor 488 | — | Choi *et al*., 2010[g] |
| H2 | — | ccgaatacaaagcatcAACGACTAGAgatgctttgtattcggTTATCTGTCG | Alexa Fluor 488 | — | Choi *et al*., 2010[g] |
| C1 | — | ATGAAGGACGgactactgataactggGACTTCCATAccagttatcagtagtc | Alexa Fluor 555 | — | Choi *et al*., 2010[g] |
| C2 | — | ccagttatcagtagtcCGTCCTTCATgactactgataactggTATGGAAGTC | Alexa Fluor 555 | — | Choi *et al*., 2010[g] |

[a]Double underlined sequences are the initiator sequence. Lowercase letters represent stem structures of amplifier probe. Underlined sequence of amplifier probes are complementary to the initiator sequnces.

[b]%FA represent formamide concentration (v/v). For *Ca*. P. syntrophicum-specific probes, FA concentrations were determined by Clone-FISH method (Schramm *et al*., 2002). The Clone-FISH sample insereted a neally full 16S rRNA gene sequence of MK-D1 was prepared as described in Kubota *et al*. (2006).

[c]For the probes, both of 5'-singly fluoresence labaled and 5'-,3'-doubly fluoresence labaled probes (Stoecker *et al*., 2010) were applied to detect *Ca*. P. syntrophicum strain MK-D1 cells.

[d]The probe also targets sequences of the some Thorarchaeota, some Bathyarchaeota and many Marine Hydrothermal Vent Group.

[e]When HCR-FISH detection using DSAG-Gr2-1142 probe, we obtained clear fluorescence signals of MK-D1 cells from the tri-cuture, but not from the co-cultures. Therefore, we designed a new probe DSAG-Gr2-1432 and mixed with DSAG-Gr2-1142 probe for clear detection of the cells in the co-cultures. FA concentration was 30% when we used two probes simultaneously.

[f]FA concentration for the MG1200 probe was re-evaluated using a pure culture of *Methanogenium cariaci* strain JR1 (JCM 10550).

[g]Amplifier probe sequences were changed to DNA probe from RNA probe.

**Supplementary Table S4 | Ribosomal proteins used for phylogenomic tree construction.**

| COG | Ribosomal protein |
|---|---|
| COG00081 | Ribosomal protein L1 |
| COG00090 | Ribosomal protein L2 |
| COG00087 | Ribosomal protein L3 |
| COG00088 | Ribosomal protein L4 |
| COG00094 | Ribosomal protein L5 |
| COG00097 | Ribosomal protein L6P/L9E |
| COG00080 | Ribosomal protein L11 |
| COG00102 | Ribosomal protein L13 |
| COG00093 | Ribosomal protein L14 |
| COG00200 | Ribosomal protein L15 |
| COG00256 | Ribosomal protein L18 |
| COG00091 | Ribosomal protein L22 |
| COG00089 | Ribosomal protein L23 |
| COG00198 | Ribosomal protein L24 |
| COG00255 | Ribosomal protein L29 |
| COG01841 | Ribosomal protein L30/L7E |
| COG00052 | Ribosomal protein S2 |
| COG00092 | Ribosomal protein S3 |
| COG00522 | Ribosomal protein S4 and related proteins |
| COG00098 | Ribosomal protein S5 |
| COG00049 | Ribosomal protein S7 |
| COG00096 | Ribosomal protein S8 |
| COG00103 | Ribosomal protein S9 |
| COG00051 | Ribosomal protein S10 |
| COG00100 | Ribosomal protein S11 |
| COG00048 | Ribosomal protein S12 |
| COG00099 | Ribosomal protein S13 |
| COG00184 | Ribosomal protein S15P/S13E |
| COG00186 | Ribosomal protein S17 |
| COG00185 | Ribosomal protein S19 |

**Supplementary Table S5 | Genomes included in phylogenomic analysis.** For genomes included in tree construction (Fig. 3a or Fig. S4), the number of ribosomal proteins (those listed in Table S4) identified in each genome is indicated.

| Domain | Genome | Alignment (Fig. S4) | Tree (Fig. S4) | Alignment (Fig. 3a) | Tree (Fig. 3a) | # RPs |
|---|---|---|---|---|---|---|
| | Acidianus hospitalis W1 | X | X | X | X | 31 |
| | Acidiplasma cupricumulans | X | X | X | X | 31 |
| | Aeropyrum pernix K1 | X | X | X | X | 30 |
| | Archaeoglobus fulgidus | X | X | X | X | 31 |
| | Caldisphaera lagunensis DSM 15908 | X | X | X | X | 31 |
| | Caldivirga maquilingensis IC-167 | X | X | X | X | 31 |
| | Cuniculiplasma divulgatum | X | X | X | X | 31 |
| | **Ca. Prometheoarchaeum syntrophicum MK-D1** | X | X | X | X | 31 |
| | Desulfurococcus kamchatkensis 1221n | X | X | X | X | 31 |
| | Ferroglobus placidus DSM 10642 | X | X | X | X | 31 |
| | Ferroplasma acidiphilum | X | X | X | X | 29 |
| | Fervidicoccus fontis Kam940 | X | X | X | X | 31 |
| | Geoglobus acetivorans | X | X | X | X | 31 |
| | Haladaptatus litoreus | X | X | X | X | 31 |
| | Halalkalicoccus jeotgali B3 | X | X | X | X | 31 |
| | Halanaeroarchaeum sulfurireducens | X | X | X | X | 31 |
| | Haloarchaeobius iranensis | X | X | X | X | 31 |
| | Haloarcula japonica DSM 6131 | X | X | X | X | 31 |
| | Halobacterium salinarum NRC-1 | X | X | X | X | 31 |
| | Halobiforma nitratireducens JCM 10879 | X | X | X | X | 31 |
| | Halococcus saccharolyticus DSM 5350 | X | X | X | X | 31 |
| | Haloferax denitrificans ATCC 35960 | X | X | X | X | 31 |
| | Halomicrobium mukohataei DSM 12286 | X | X | X | X | 31 |
| | Halopiger xanaduensis SH-6 | X | X | X | X | 31 |
| | Haloquadratum walsbyi DSM 16790 | X | X | X | X | 31 |
| | Halorubrum californiensis DSM 19288 | X | X | X | X | 31 |
| | Haloterrigena limicola JCM 13563 | X | X | X | X | 31 |
| | Hyperthermus butylicus DSM 5456 | X | X | X | X | 30 |
| | Ignicoccus islandicus DSM 13165 | X | X | X | X | 31 |
| Archaea | Ignisphaera aggregans DSM 17230 | X | X | X | X | 31 |
| | Metallosphaera yellowstonensis MK1 | X | X | X | X | 31 |
| | Methanobacterium formicicum DSM 3637 | X | X | X | X | 31 |
| | Methanobrevibacter smithii ATCC 35061 | X | X | X | X | 31 |
| | Methanocaldococcus jannaschii DSM 2661 | X | X | X | X | 31 |
| | Methanococcus aeolicus Nankai-3 | X | X | X | X | 31 |
| | Methanocorpusculum labreanum Z | X | X | X | X | 31 |
| | Methanoculleus bourgensis MS2 | X | X | X | X | 31 |
| | Methanomethylovorans hollandica DSM 15978 | X | X | X | X | 31 |
| | Methanonatronarchaeum thermophilum | X | X | X | X | 31 |
| | Methanopyrus kandleri AV19 | X | X | X | X | 31 |
| | Methanoregula boonei 6A8 | X | X | X | X | 31 |
| | Methanosaeta concilii GP6 | X | X | X | X | 31 |
| | Methanosarcina barkeri MS | X | X | X | X | 30 |
| | Methanosphaera stadtmanae DSM 3091 | X | X | X | X | 31 |
| | Methanothermobacter thermautotrophicus str Delta H | X | X | X | X | 31 |
| | Natrinema altunense JCM 12890 | X | X | X | X | 31 |
| | Natronorubrum sulfidifaciens JCM 14089 | X | X | X | X | 31 |
| | Nitrosopumilus maritimus SCM1 | X | X | X | X | 31 |
| | Nitrososphaera viennensis EN76 | X | X | X | X | 31 |
| | Pyrobaculum ferrireducens | X | X | X | X | 31 |
| | Pyrococcus furiosus DSM 3638 | X | X | X | X | 31 |
| | Staphylothermus marinus F1 | X | X | X | X | 31 |
| | Sulfolobus islandicus YN1551 | X | X | X | X | 31 |
| | Thermococcus kodakarensis KOD1 | X | X | X | X | 31 |
| | Thermofilum uzonense | X | X | X | X | 31 |
| | Thermoplasma volcanium GSS1 | X | X | X | X | 30 |
| | Thermoproteus uzoniensis 768-20 | X | X | X | X | 31 |
| | Thermosphaera aggregans DSM 11486 | X | X | X | X | 31 |
| | Vulcanisaeta distributa DSM 14429 | X | X | X | X | 31 |
| | Arabidopsis thaliana Columbia | X | X | X | X | 31 |
| | Aspergillus fumigatus Af293 | X | X | X | X | 30 |
| | Chlamydomonas reinhardtii | X | X | X | X | 31 |
| | Leishmania major Friedlin V1 | X | X | X | X | 31 |
| Eukarya | Monosiga brevicollis MX1 | X | X | X | X | 29 |
| | Mus musculus C57BL | X | X | X | X | 30 |
| | Naegleria gruberi NEG-M | X | X | X | X | 30 |
| | Paramecium tetraurelia d4-2 | X | X | X | X | 31 |
| | Phytophthora ramorum Pr102 | X | X | X | X | 29 |
| | Plasmodium falciparum 3D7 | X | X | X | X | 29 |
| | Bacillus subtilis subsp subtilis str 168 | X | X | X | X | 31 |
| | Bacteroides thetaiotaomicron VPI-5482 | X | X | X | X | 31 |
| | Borrelia burgdorferi ZS7 | X | X | X | X | 31 |
| | Campylobacter jejuni NCTC 11168 | X | X | X | X | 31 |
| Bacteria | Escherichia coli str K-12 | X | X | X | X | 31 |
| | Rhodopirellula baltica SH 1 | X | X | X | X | 31 |
| | Rickettsia prowazekii str Rp22 | X | X | X | X | 31 |
| | Synechococcus elongatus PCC 6301 | X | X | X | X | 31 |
| | Thermotoga maritima | X | X | X | X | 31 |
| | Candidatus Korarchaeum cryptofilum OPF8 | X | X | | | 31 |
| | Cenarchaeum symbiosum A | X | X | | | 31 |
| | Arc I group archaeon ADurb1013 Bin02101 | X | | | | |
| | Bathyarchaeota BA1 | X | | | | |
| | Bathyarchaeota BA2 | X | | | | |
| | Candidatus Aenigmarchaeota GW2011_AR5 | X | | | | |
| | Candidatus Aenigmarchaeota archaeon CG1_02_38_14 | X | | | | |
| | Candidatus Caldiarchaeum subterraneum | X | | | | |
| | Candidatus Haloredivivus sp G17 | X | | | | |
| | Candidatus Heidallarchaeota AB_125 | X | | | | |
| | Candidatus Heimdallarchaeota LC_2 | X | | | | |
| | Candidatus Heimdallarchaeota LC_3 | X | | | | |
| | Candidatus Lokiarchaeota archaeon CR 4 | X | | | | |
| | Candidatus Methanohalarchaeum thermophilum | X | | | | |
| | Candidatus Methanomassiliicoccus intestinalis Issoire-Mx1 | X | | | | |
| | Candidatus Methanomethylicus mesodigestum V2 | X | | | | |
| | Candidatus Methanomethylicus oleusabulum V3 | X | | | | |
| | Candidatus Methanomethylophilus alvus Mx1201 | X | | | | |
| | Candidatus Methanoperedens nitroreducens | X | | | | |
| | Candidatus Methanoplasma termitum | X | | | | |
| Archaea (uncultured) | Candidatus Micrarchaeum acidiphilum ARMAN-1 | X | | | | |
| | Candidatus Nanosalina sp J07AB43 | X | | | | |
| | Candidatus Nanosalinarum sp J07AB56 | X | | | | |
| | Candidatus Nitrocosmicus oleophilus | X | | | | |
| | Candidatus Nitrosoarchaeum koreensis MY1 | X | | | | |
| | Candidatus Nitrosomarinus catalina | X | | | | |
| | Candidatus Nitrosopelagicus brevis | X | | | | |
| | Candidatus Nitrososphaera gargensis Ga92 | X | | | | |
| | Candidatus Nitrosotalea devanaterra | X | | | | |
| | Candidatus Nitrosotenuis cloacae | X | | | | |
| | Candidatus Odinarchaeota archaeon LCB_4 | X | | | | |
| | Candidatus Syntrophoarchaeum butanivorans | X | | | | |
| | Candidatus Thorarchaeota archaeon AB_25 | X | | | | |
| | Candidatus Thorarchaeota archaeon SMTZ-45 | X | | | | |
| | Candidatus Thorarchaeota archaeon SMTZ1-45 | X | | | | |
| | Candidatus Thorarchaeota archaeon SMTZ1-83 | X | | | | |
| | Candidatus Woesearchaeota AR18 | X | | | | |
| | Candidatus Woesearchaeota AR19 | X | | | | |
| | Candidatus Woesearchaeota archaeon CG1_02_57_44 | X | | | | |
| | GWA2_AR13_28_113 | X | | | | |
| | Lokiarchaeum sp GC14_75 | X | | | | |
| | Parvarchaeum acidiphilum | X | | | | |
| | Aciduliprofundum boonei T469 | X | | | | |
| Archaea (low completeness) | Methanogenium cariaci JCM 10550 | X | | | | |
| | Thermocladium sp ECH_B | X | | | | |
| Eukarya (low completeness) | Giardia lamblia ATCC 50803 | X | | | | |
| | Ustilago maydis 521 | X | | | | |

**Supplementary Table S6 | MK-D1 gene expression data.** For each gene, the locus tag, predicted product, KEGG annotation (if available), and gene expression level (reads per kilobase of transcript per million mapped reads; RPKM).

| Locus Tag | Product | KEGG | RPKM | ESPs |
|---|---|---|---|---|
| DSAG12_00001 | hypothetical protein | -- | 204 | |
| DSAG12_00002 | hypothetical protein | -- | 213 | |
| DSAG12_00003 | | K00837 | 204 | |
| DSAG12_00004 | hypothetical protein | -- | 1148 | |
| DSAG12_00005 | hypothetical protein | -- | 529 | |
| DSAG12_00006 | HSP20 family protein | K13993 | 283 | |
| DSAG12_00007 | hypothetical protein | -- | 170 | |
| DSAG12_00008 | hypothetical protein | -- | 162 | |
| DSAG12_00009 | hypothetical protein | -- | 314 | |
| DSAG12_00010 | urocanate hydratase | K01712 | 238 | |
| DSAG12_00011 | histidine ammonia-lyase | K01745 | 259 | |
| DSAG12_00012 | imidazolonepropionase | K01468 | 236 | |
| DSAG12_00013 | formate--tetrahydrofolate ligase | K01938 | 161 | |
| DSAG12_00014 | hypothetical protein | -- | 276 | |
| DSAG12_00015 | ferredoxin hydrogenase large subunit | K00533 | 132 | |
| DSAG12_00016 | NADH-quinone oxidoreductase subunit E | K00334 | 189 | |
| DSAG12_00017 | putative transposase | K07496 | 0 | |
| DSAG12_00018 | tRNA-Trp | | 187 | |
| DSAG12_00019 | O-methyltransferase | K15471 | 155 | |
| DSAG12_00020 | hypothetical protein | -- | 224 | |
| DSAG12_00021 | polar amino acid transport system ATP-binding protein | K02028 | 223 | |
| DSAG12_00022 | arginine/lysine/histidine/glutamine transport system substrate-binding and permease protein | K17062 | 309 | |
| DSAG12_00023 | polar amino acid transport system substrate-binding protein | K02030 | 182 | |
| DSAG12_00024 | uncharacterized protein | K06864 | 0 | |
| DSAG12_00025 | hypothetical protein | -- | 157 | |
| DSAG12_00026 | Lrp/AsnC family transcriptional regulator, leucine-responsive regulatory protein | K03719 | 174 | |
| DSAG12_00027 | Ras-related protein Rab-43 | K07930 | 130 | small GTP-binding domain protein |
| DSAG12_00028 | hypothetical protein | -- | 175 | |
| DSAG12_00029 | hypothetical protein | -- | 105 | |
| DSAG12_00030 | hypothetical protein | -- | 172 | |
| DSAG12_00031 | proteasome beta subunit | K03433 | 239 | |
| DSAG12_00032 | hypothetical protein | -- | 158 | |
| DSAG12_00033 | glutamyl-tRNA synthetase | K01885 | 250 | |
| DSAG12_00034 | hypothetical protein | -- | 112 | |
| DSAG12_00035 | hypothetical protein | -- | 164 | |
| DSAG12_00036 | hypothetical protein | -- | 83 | |
| DSAG12_00037 | tRNA pseudouridine38-40 synthase | K06173 | 158 | |
| DSAG12_00038 | hypothetical protein | -- | 223 | |
| DSAG12_00039 | replication factor C small subunit | K04801 | 211 | |
| DSAG12_00040 | hypothetical protein | -- | 164 | |
| DSAG12_00041 | hypothetical protein | -- | 167 | |
| DSAG12_00042 | hypothetical protein | -- | 181 | |
| DSAG12_00043 | UDP-2,3-diacylglucosamine hydrolase | K03269 | 197 | |
| DSAG12_00044 | hypothetical protein | -- | 263 | |
| DSAG12_00045 | hypothetical protein | -- | 166 | |
| DSAG12_00046 | hypothetical protein | -- | 221 | |
| DSAG12_00047 | hypothetical protein | -- | 96 | |
| DSAG12_00048 | hypothetical protein | -- | 32 | |
| DSAG12_00049 | | K06889 | 0 | |
| DSAG12_00050 | dihydrodiol dehydrogenase / D-xylose 1-dehydrogenase (NADP) | K00078 | 169 | |
| DSAG12_00051 | MFS transporter, SP family, sugar:H+ symporter | K08139 | 228 | |
| DSAG12_00052 | peptide methionine sulfoxide reductase msrA/msrB | K12267 | 234 | |
| DSAG12_00053 | phosphohistidine phosphatase | K08296 | 188 | |
| DSAG12_00054 | hypothetical protein | -- | 171 | |
| DSAG12_00055 | hypothetical protein | K09164 | 194 | |
| DSAG12_00056 | hypothetical protein | -- | 166 | |
| DSAG12_00057 | hypothetical protein | -- | 171 | |
| DSAG12_00058 | ADP-ribosylation factor-like protein 5B | K07950 | 172 | |
| DSAG12_00059 | hypothetical protein | -- | 302 | |
| DSAG12_00060 | hypothetical protein | -- | 420 | |
| DSAG12_00061 | internalin A | K13730 | 115 | |
| DSAG12_00062 | hypothetical protein | -- | 224 | |
| DSAG12_00063 | peptide-methionine (R)-S-oxide reductase | K07305 | 155 | |
| DSAG12_00064 | aminopeptidase N | K01256 | 178 | |
| DSAG12_00065 | hypothetical protein | K09120 | 228 | |
| DSAG12_00066 | hypothetical protein | -- | 128 | |
| DSAG12_00067 | hypothetical protein | -- | 152 | |
| DSAG12_00068 | hypothetical protein | -- | 147 | |
| DSAG12_00069 | exosome complex component CSL4 | K07573 | 59 | |
| DSAG12_00070 | putative methylase | K07579 | 160 | |
| DSAG12_00071 | hypothetical protein | -- | 169 | |
| DSAG12_00072 | Trp repressor binding protein | K03809 | 19 | |
| DSAG12_00073 | hypothetical protein | -- | 213 | |
| DSAG12_00074 | enolase | K01689 | 333 | |
| DSAG12_00075 | serine/threonine-protein phosphatase 4 catalytic subunit | K15423 | 236 | |

| | | | |
|---|---|---|---|
| DSAG12_00076 | hypothetical protein | -- | 189 |
| DSAG12_00077 | alanine dehydrogenase | K00259 | 147 |
| DSAG12_00078 | acetyl-CoA C-acetyltransferase | K00626 | 274 |
| DSAG12_00079 | hypothetical protein | -- | 304 |
| DSAG12_00080 | butyryl-CoA dehydrogenase | K00248 | 205 |
| DSAG12_00081 | 3-hydroxybutyryl-CoA dehydratase | K17865 | 284 |
| DSAG12_00082 | hypothetical protein | -- | 175 |
| DSAG12_00083 | 3-oxoacid CoA-transferase subunit B | K01029 | 223 |
| DSAG12_00084 | 3-oxoacid CoA-transferase subunit A | K01028 | 272 |
| DSAG12_00085 | putative membrane protein | K08981 | 273 |
| DSAG12_00086 | hypothetical protein | -- | 90 |
| DSAG12_00087 | hypothetical protein | -- | 174 |
| DSAG12_00088 | | K00936 | 164 |
| DSAG12_00089 | hypothetical protein | -- | 238 |
| DSAG12_00090 | hypothetical protein | -- | 201 |
| DSAG12_00091 | elongation factor 1-alpha | K03231 | 146 |
| DSAG12_00092 | magnesium-protoporphyrin IX monomethyl ester (oxidative) cyclase | K04035 | 289 |
| DSAG12_00093 | hypothetical protein | -- | 200 |
| DSAG12_00094 | archaea-specific DNA-binding protein | K03622 | 42 |
| DSAG12_00095 | | K06883 | 138 |
| DSAG12_00096 | 2-isopropylmalate synthase | K01649 | 218 |
| DSAG12_00097 | hypothetical protein | -- | 231 |
| DSAG12_00098 | small nuclear ribonucleoprotein D2 | K11096 | 216 |
| DSAG12_00099 | hypothetical protein | -- | 51 |
| DSAG12_00100 | hypothetical protein | -- | 163 |
| DSAG12_00101 | hypothetical protein | -- | 114 |
| DSAG12_00102 | hypothetical protein | -- | 150 |
| DSAG12_00103 | hypothetical protein | -- | 145 |
| DSAG12_00104 | hypothetical protein | -- | 65 |
| DSAG12_00105 | | K07051 | 225 |
| DSAG12_00106 | hypothetical protein | K09125 | 245 |
| DSAG12_00107 | tocopherol O-methyltransferase | K05928 | 209 |
| DSAG12_00108 | tRNA-Ser | | 0 |
| DSAG12_00109 | large subunit ribosomal protein L14e | K02875 | 98 |
| DSAG12_00110 | cytidylate kinase | K00945 | 207 |
| DSAG12_00111 | large subunit ribosomal protein L34e | K02915 | 105 |
| DSAG12_00112 | hypothetical protein | -- | 0 |
| DSAG12_00113 | hypothetical protein | -- | 229 |
| DSAG12_00114 | hypothetical protein | -- | 151 |
| DSAG12_00115 | hypothetical protein | -- | 221 |
| DSAG12_00116 | hypothetical protein | -- | 156 |
| DSAG12_00117 | RNA 3'-terminal phosphate cyclase (ATP) | K01974 | 173 |
| DSAG12_00118 | nicotinamide-nucleotide amidase | K03742 | 348 |
| DSAG12_00119 | hypothetical protein | -- | 129 |
| DSAG12_00120 | hypothetical protein | K09134 | 156 |
| DSAG12_00121 | hypothetical protein | -- | 195 |
| DSAG12_00122 | hypothetical protein | -- | 109 |
| DSAG12_00123 | hypothetical protein | -- | 162 |
| DSAG12_00124 | hypothetical protein | -- | 162 |
| DSAG12_00125 | hypothetical protein | -- | 180 |
| DSAG12_00126 | tRNA-Ile | Intron(12645 | 169 |
| DSAG12_00127 | hypothetical protein | -- | 254 |
| DSAG12_00128 | valyl-tRNA synthetase | K01873 | 131 |
| DSAG12_00129 | hypothetical protein | -- | 205 |
| DSAG12_00130 | hypothetical protein | -- | 129 |
| DSAG12_00131 | hypothetical protein | -- | 0 |
| DSAG12_00132 | tRNA-Ala | | 207 |
| DSAG12_00133 | hypothetical protein | -- | 212 |
| DSAG12_00134 | 4-hydroxybutyryl-CoA synthetase (ADP-forming) | K18593 | 0 |
| DSAG12_00135 | glucosamine--fructose-6-phosphate aminotransferase (isomerizing) | K00820 | 272 |
| DSAG12_00136 | hypothetical protein | -- | 120 |
| DSAG12_00137 | pyruvate, water dikinase | K01007 | 216 |
| DSAG12_00138 | MFS transporter, DHA1 family, tetracycline resistance protein | K08151 | 179 |
| DSAG12_00139 | hypothetical protein | -- | 213 |
| DSAG12_00140 | hypothetical protein | -- | 0 |
| DSAG12_00141 | hypothetical protein | -- | 206 |
| DSAG12_00142 | leucyl-tRNA synthetase | K01869 | 27 |
| DSAG12_00143 | hypothetical protein | -- | 250 |
| DSAG12_00144 | hypothetical protein | -- | 173 |
| DSAG12_00145 | meiotic recombination protein SPO11 | K10878 | 162 |
| DSAG12_00146 | hypothetical protein | -- | 214 |
| DSAG12_00147 | phosphopantothenoylcysteine decarboxylase / phosphopantothenate--cysteine ligase | K13038 | 224 |
| DSAG12_00148 | hypothetical protein | -- | 207 |
| DSAG12_00149 | archaea-specific RecJ-like exonuclease | K07463 | 127 |
| DSAG12_00150 | hypothetical protein | -- | 198 |
| DSAG12_00151 | O-phospho-L-seryl-tRNASec:L-selenocysteinyl-tRNA synthase | K03341 | 196 |
| DSAG12_00152 | hypothetical protein | -- | 220 |
| DSAG12_00153 | nucleolar GTP-binding protein | K06943 | 196 |

| | | | |
|---|---|---|---|
| DSAG12_00154 | hypothetical protein | -- | 156 |
| DSAG12_00155 | hypothetical protein | -- | 211 |
| DSAG12_00156 | hypothetical protein | -- | 250 |
| DSAG12_00157 | | K00837 | 168 |
| DSAG12_00158 | D-3-phosphoglycerate dehydrogenase | K00058 | 271 |
| DSAG12_00159 | hypothetical protein | -- | 260 |
| DSAG12_00160 | retinol dehydrogenase 13 | K11161 | 236 |
| DSAG12_00161 | TldD protein | K03568 | 218 |
| DSAG12_00162 | PmbA protein | K03592 | 272 |
| DSAG12_00163 | acetyltransferase | K09181 | 238 |
| DSAG12_00164 | | K06873 | 339 |
| DSAG12_00165 | hypothetical protein | -- | 266 |
| DSAG12_00166 | hypothetical protein | -- | 131 |
| DSAG12_00167 | putative protease | K08303 | 197 |
| DSAG12_00168 | hypothetical protein | -- | 235 |
| DSAG12_00169 | hypothetical protein | -- | 191 |
| DSAG12_00170 | presenilin 1 | K04505 | 116 |
| DSAG12_00171 | hypothetical protein | -- | 232 |
| DSAG12_00172 | starch synthase (maltosyl-transferring) | K16147 | 276 |
| DSAG12_00173 | nitrogen fixation protein NifU and related proteins | K04488 | 220 |
| DSAG12_00174 | putative ABC transport system permease protein | K02004 | 175 |
| DSAG12_00175 | putative ABC transport system permease protein | K02004 | 171 |
| DSAG12_00176 | putative ABC transport system ATP-binding protein | K02003 | 138 |
| DSAG12_00177 | hypothetical protein | -- | 188 |
| DSAG12_00178 | hypothetical protein | -- | 270 |
| DSAG12_00179 | hypothetical protein | K09732 | 128 |
| DSAG12_00180 | hypothetical protein | -- | 65 |
| DSAG12_00181 | hypothetical protein | -- | 51 |
| DSAG12_00182 | beta-phosphoglucomutase | K01838 | 283 |
| DSAG12_00183 | hypothetical protein | -- | 165 |
| DSAG12_00184 | glycerophosphoryl diester phosphodiesterase | K01126 | 161 |
| DSAG12_00185 | hypothetical protein | -- | 186 |
| DSAG12_00186 | lipoprotein-releasing system ATP-binding protein | K09810 | 219 |
| DSAG12_00187 | hypothetical protein | -- | 202 |
| DSAG12_00188 | hypothetical protein | -- | 190 |
| DSAG12_00189 | centriolar protein POC1 | K16482 | 116 |
| DSAG12_00190 | hypothetical protein | -- | 165 |
| DSAG12_00191 | Xaa-Pro aminopeptidase | K01262 | 133 |
| DSAG12_00192 | hypothetical protein | -- | 219 |
| DSAG12_00193 | hypothetical protein | -- | 187 |
| DSAG12_00194 | | K06933 | 192 |
| DSAG12_00195 | hypothetical protein | -- | 150 |
| DSAG12_00196 | Lrp/AsnC family transcriptional regulator, leucine-responsive regulatory protein | K03719 | 255 |
| DSAG12_00197 | hypothetical protein | -- | 180 |
| DSAG12_00198 | MFS transporter, DHA1 family, tetracycline resistance protein | K08151 | 233 |
| DSAG12_00199 | DNA mismatch repair protein MutL | K03572 | 171 |
| DSAG12_00200 | elongation factor 1-beta | K03232 | 192 |
| DSAG12_00201 | hypothetical protein | -- | 100 |
| DSAG12_00202 | hypothetical protein | -- | 137 |
| DSAG12_00203 | long-chain acyl-CoA synthetase | K01897 | 153 |
| DSAG12_00204 | hypothetical protein | -- | 270 |
| DSAG12_00205 | carboxypeptidase T | K05996 | 169 |
| DSAG12_00206 | DNA adenine methylase | K06223 | 187 |
| DSAG12_00207 | phosphate transport system protein | K02039 | 113 |
| DSAG12_00208 | hypothetical protein | -- | 179 |
| DSAG12_00209 | hypothetical protein | -- | 173 |
| DSAG12_00210 | 2-succinyl-6-hydroxy-2,4-cyclohexadiene-1-carboxylate synthase | K08680 | 181 |
| DSAG12_00211 | hypothetical protein | -- | 173 |
| DSAG12_00212 | ArsR family transcriptional regulator | K03892 | 52 |
| DSAG12_00213 | hypothetical protein | -- | 132 |
| DSAG12_00214 | | K07089 | 113 |
| DSAG12_00215 | hypothetical protein | -- | 210 |
| DSAG12_00216 | hypothetical protein | -- | 175 |
| DSAG12_00217 | hypothetical protein | -- | 197 |
| DSAG12_00218 | hypothetical protein | -- | 140 |
| DSAG12_00219 | hypothetical protein | -- | 142 |
| DSAG12_00220 | hypothetical protein | -- | 118 |
| DSAG12_00221 | hypothetical protein | -- | 264 |
| DSAG12_00222 | ethanolamine utilization protein EutA | K04019 | 241 |
| DSAG12_00223 | hypothetical protein | -- | 258 |
| DSAG12_00224 | hypothetical protein | -- | 205 |
| DSAG12_00225 | HSP20 family protein | K13993 | 0 |
| DSAG12_00226 | hypothetical protein | -- | 309 |
| DSAG12_00227 | HSP20 family protein | K13993 | 269 |
| DSAG12_00228 | hypothetical protein | -- | 361 |
| DSAG12_00229 | DNA mismatch repair protein MutS | K03555 | 216 |
| DSAG12_00230 | hypothetical protein | -- | 218 |
| DSAG12_00231 | hypothetical protein | -- | 226 |

| | | | |
|---|---|---|---|
| DSAG12_00232 | hypothetical protein | -- | 182 |
| DSAG12_00233 | hypothetical protein | -- | 206 |
| DSAG12_00234 | hypothetical protein | -- | 164 |
| DSAG12_00235 | hypothetical protein | -- | 60 |
| DSAG12_00236 | hypothetical protein | -- | 477 |
| DSAG12_00237 | | K07052 | 203 |
| DSAG12_00238 | hypothetical protein | -- | 238 |
| DSAG12_00239 | | K07138 | 185 |
| DSAG12_00240 | hypothetical protein | -- | 246 |
| DSAG12_00241 | hypothetical protein | -- | 233 |
| DSAG12_00242 | archaea-specific RecJ-like exonuclease | K07463 | 175 |
| DSAG12_00243 | hypothetical protein | -- | 226 |
| DSAG12_00244 | ABC-2 type transport system ATP-binding protein | K01990 | 256 |
| DSAG12_00245 | ABC-2 type transport system permease protein | K01992 | 258 |
| DSAG12_00246 | hypothetical protein | -- | 195 |
| DSAG12_00247 | hypothetical protein | -- | 167 |
| DSAG12_00248 | hypothetical protein | -- | 113 |
| DSAG12_00249 | proteasome regulatory subunit | K03420 | 175 |
| DSAG12_00250 | hypothetical protein | -- | 329 |
| DSAG12_00251 | 2-oxoisovalerate ferredoxin oxidoreductase, beta subunit | K00187 | 57 |
| DSAG12_00252 | 2-oxoisovalerate ferredoxin oxidoreductase, alpha subunit | K00186 | 343 |
| DSAG12_00253 | pyruvate ferredoxin oxidoreductase, delta subunit | K00171 | 278 |
| DSAG12_00254 | 2-oxoisovalerate ferredoxin oxidoreductase, gamma subunit | K00189 | 171 |
| DSAG12_00255 | transcription initiation factor TFIIB | K03124 | 209 |
| DSAG12_00256 | hypothetical protein | -- | 307 |
| DSAG12_00257 | hypothetical protein | -- | 68 |
| DSAG12_00258 | | K01567 | 277 |
| DSAG12_00259 | transcription initiation factor TFIIB | K03124 | 227 |
| DSAG12_00260 | hypothetical protein | -- | 246 |
| DSAG12_00261 | hypothetical protein | -- | 175 |
| DSAG12_00262 | | K07073 | 186 |
| DSAG12_00263 | | K07129 | 242 |
| DSAG12_00264 | hypothetical protein | -- | 223 |
| DSAG12_00265 | hypothetical protein | -- | 212 |
| DSAG12_00266 | GMP synthase (glutamine-hydrolysing) | K01951 | 231 |
| DSAG12_00267 | | K06935 | 218 |
| DSAG12_00268 | protein pelota | K06965 | 163 |
| DSAG12_00269 | hypothetical protein | -- | 194 |
| DSAG12_00270 | xanthine phosphoribosyltransferase | K00769 | 155 |
| DSAG12_00271 | energy-coupling factor transport system permease protein | K16785 | 136 |
| DSAG12_00272 | energy-coupling factor transport system ATP-binding protein / energy-coupling factor transport system ATP-binding protein | K16786 K16 | 180 |
| DSAG12_00273 | phosphoesterase RecJ domain-containing protein | K06881 | 207 |
| DSAG12_00274 | prefoldin beta subunit | K04798 | 178 |
| DSAG12_00275 | hypothetical protein | -- | 142 |
| DSAG12_00276 | putative transcriptional regulator | K07728 | 153 |
| DSAG12_00277 | ribonuclease HII | K03470 | 179 |
| DSAG12_00278 | gluconate 5-dehydrogenase | K00046 | 222 |
| DSAG12_00279 | acyl-CoA thioesterase | K02614 | 202 |
| DSAG12_00280 | hypothetical protein | -- | 198 |
| DSAG12_00281 | 3-phenylpropionate/trans-cinnamate dioxygenase ferredoxin subunit | K05710 | 203 |
| DSAG12_00282 | fibrillarin-like pre-rRNA processing protein | K04795 | 132 |
| DSAG12_00283 | nucleolar protein 56 | K14564 | 186 |
| DSAG12_00284 | dihydroorotate dehydrogenase (NAD+) catalytic subunit | K17828 | 207 |
| DSAG12_00285 | dihydroorotate dehydrogenase electron transfer subunit | K02823 | 218 |
| DSAG12_00286 | elongator complex protein 3 | K07739 | 225 |
| DSAG12_00287 | hypothetical protein | -- | 298 |
| DSAG12_00288 | hypothetical protein | -- | 218 |
| DSAG12_00289 | hypothetical protein | -- | 193 |
| DSAG12_00290 | hypothetical protein | -- | 59 |
| DSAG12_00291 | hypothetical protein | -- | 274 |
| DSAG12_00292 | hypothetical protein | -- | 126 |
| DSAG12_00293 | hypothetical protein | -- | 253 |
| DSAG12_00294 | | K01529 | 163 |
| DSAG12_00295 | hypothetical protein | -- | 256 |
| DSAG12_00296 | hypothetical protein | -- | 144 |
| DSAG12_00297 | hypothetical protein | -- | 128 |
| DSAG12_00298 | hypothetical protein | -- | 191 |
| DSAG12_00299 | hypothetical protein | -- | 149 |
| DSAG12_00300 | hypothetical protein | -- | 179 |
| DSAG12_00301 | serine/threonine protein kinase, bacterial | K08884 | 225 |
| DSAG12_00302 | hypothetical protein | -- | 160 |
| DSAG12_00303 | putative ATP-dependent endonuclease of the OLD family | K07459 | 162 |
| DSAG12_00304 | putative adenine-specific DNA-methyltransferase | K07319 | 177 |
| DSAG12_00305 | hypothetical protein | -- | 107 |
| DSAG12_00306 | radical S-adenosyl methionine domain-containing protein 2 | K15045 | 138 |
| DSAG12_00307 | hypothetical protein | -- | 118 |
| DSAG12_00308 | hypothetical protein | -- | 0 |

| | | | | |
|---|---|---|---|---|
| DSAG12_00309 | hypothetical protein | -- | 187 | |
| DSAG12_00310 | hypothetical protein | -- | 187 | |
| DSAG12_00311 | hypothetical protein | -- | 242 | |
| DSAG12_00312 | hypothetical protein | -- | 223 | |
| DSAG12_00313 | hypothetical protein | -- | 204 | |
| DSAG12_00314 | hypothetical protein | -- | 189 | |
| DSAG12_00315 | hypothetical protein | -- | 48 | |
| DSAG12_00316 | hypothetical protein | -- | 20 | |
| DSAG12_00317 | elongator complex protein 3 | K07739 | 141 | |
| DSAG12_00318 | GMP synthase (glutamine-hydrolysing) | K01951 | 133 | |
| DSAG12_00319 | hypothetical protein | -- | 0 | |
| DSAG12_00320 | putative DNA methylase | K07445 | 207 | |
| DSAG12_00321 | hypothetical protein | -- | 192 | |
| DSAG12_00322 | serine/threonine protein kinase, bacterial | K08884 | 101 | |
| DSAG12_00323 | hypothetical protein | -- | 207 | |
| DSAG12_00324 | hypothetical protein | -- | 181 | |
| DSAG12_00325 | hypothetical protein | -- | 203 | |
| DSAG12_00326 | hypothetical protein | -- | 226 | |
| DSAG12_00327 | TetR/AcrR family transcriptional regulator, transcriptional repressor for nem operon | K16137 | 136 | |
| DSAG12_00328 | glutamyl-tRNA(Gln) amidotransferase subunit D | K09482 | 139 | |
| DSAG12_00329 | glutamyl-tRNA(Gln) amidotransferase subunit E | K03330 | 265 | |
| DSAG12_00330 | hypothetical protein | -- | 223 | |
| DSAG12_00331 | hypothetical protein | -- | 114 | |
| DSAG12_00332 | hypothetical protein | -- | 157 | |
| DSAG12_00333 | hypothetical protein | -- | 208 | |
| DSAG12_00334 | molybdopterin synthase sulfur carrier subunit | K03636 | 145 | |
| DSAG12_00335 | hypothetical protein | -- | 84 | |
| DSAG12_00336 | aldehyde:ferredoxin oxidoreductase | K03738 | 179 | |
| DSAG12_00337 | glutamyl-tRNA(Gln) amidotransferase subunit E | K03330 | 266 | |
| DSAG12_00338 | glutamyl-tRNA(Gln) amidotransferase subunit D | K09482 | 206 | |
| DSAG12_00339 | hypothetical protein | -- | 256 | |
| DSAG12_00340 | hypothetical protein | -- | 62 | |
| DSAG12_00341 | hypothetical protein | -- | 265 | |
| DSAG12_00342 | hypothetical protein | -- | 201 | |
| DSAG12_00343 | hypothetical protein | -- | 411 | |
| DSAG12_00344 | hypothetical protein | -- | 337 | |
| DSAG12_00345 | hypothetical protein | -- | 188 | |
| DSAG12_00346 | hypothetical protein | -- | 219 | |
| DSAG12_00347 | hypothetical protein | -- | 191 | |
| DSAG12_00348 | ribosome biogenesis ATPase | K14571 | 193 | |
| DSAG12_00349 | hypothetical protein | -- | 195 | |
| DSAG12_00350 | hypothetical protein | -- | 129 | |
| DSAG12_00351 | hypothetical protein | -- | 131 | |
| DSAG12_00352 | galactokinase | K00849 | 168 | |
| DSAG12_00353 | hypothetical protein | -- | 260 | |
| DSAG12_00354 | hypothetical protein | -- | 245 | |
| DSAG12_00355 | | K07059 | 145 | |
| DSAG12_00356 | hypothetical protein | -- | 165 | |
| DSAG12_00357 | peptide/nickel transport system substrate-binding protein | K02035 | 215 | |
| DSAG12_00358 | alanyl-tRNA synthetase | K01872 | 215 | |
| DSAG12_00359 | hypothetical protein | -- | 118 | |
| DSAG12_00360 | | K07050 | 235 | |
| DSAG12_00361 | ATP-binding cassette, subfamily B, bacterial | K06147 | 199 | |
| DSAG12_00362 | ATP-binding cassette, subfamily B, bacterial MsbA | K11085 | 231 | |
| DSAG12_00363 | hypothetical protein | -- | 262 | |
| DSAG12_00364 | hypothetical protein | -- | 0 | |
| DSAG12_00365 | hypothetical protein | -- | 263 | |
| DSAG12_00366 | tRNA 2-thiouridine synthesizing protein D | K07235 | 50 | |
| DSAG12_00367 | tRNA 2-thiouridine synthesizing protein A | K04085 | 213 | |
| DSAG12_00368 | tRNA-Lys | Intron(41766 | 134 | |
| DSAG12_00369 | hypothetical protein | -- | 48 | |
| DSAG12_00370 | hypothetical protein | -- | 154 | |
| DSAG12_00371 | hypothetical protein | -- | 132 | |
| DSAG12_00372 | hypothetical protein | -- | 150 | |
| DSAG12_00373 | hypothetical protein | -- | 141 | |
| DSAG12_00374 | hypothetical protein | -- | 146 | |
| DSAG12_00375 | hypothetical protein | -- | 143 | |
| DSAG12_00376 | H/ACA ribonucleoprotein complex subunit 4 | K11131 | 238 | small GTP-binding domain protein |
| DSAG12_00377 | 16S rRNA (guanine1207-N2)-methyltransferase | K00564 | 246 | |
| DSAG12_00378 | MFS transporter, DHA1 family, multidrug resistance protein | K08153 | 189 | |
| DSAG12_00379 | DNA-directed RNA polymerase subunit H | K03053 | 190 | |
| DSAG12_00380 | DNA-directed RNA polymerase subunit B | K13798 | 109 | |
| DSAG12_00381 | hypothetical protein | -- | 311 | |
| DSAG12_00382 | hypothetical protein | -- | 176 | |
| DSAG12_00383 | aminoglycoside N6'-acetyltransferase | K00663 | 44 | |
| DSAG12_00384 | tRNA/rRNA methyltransferase | K02533 | 172 | |
| DSAG12_00385 | hypothetical protein | -- | 209 | |
| DSAG12_00386 | hypothetical protein | -- | 186 | |

| | | | | |
|---|---|---|---|---|
| DSAG12_00387 | hypothetical protein | -- | 64 | |
| DSAG12_00388 | hypothetical protein | -- | 196 | |
| DSAG12_00389 | transcription initiation factor TFIIE subunit alpha | K03136 | 182 | |
| DSAG12_00390 | hypothetical protein | -- | 180 | |
| DSAG12_00391 | peptidyl-prolyl cis-trans isomerase A (cyclophilin A) | K03767 | 129 | |
| DSAG12_00392 | RIO kinase 2 | K07179 | 156 | |
| DSAG12_00393 | demethylmenaquinone methyltransferase / 2-methoxy-6-polyprenyl-1,4-benzoquinol methylase | K03183 | 221 | |
| DSAG12_00394 | small nuclear ribonucleoprotein | K04796 | 140 | |
| DSAG12_00395 | hypothetical protein | -- | 69 | |
| DSAG12_00396 | ArsR family transcriptional regulator | K07721 | 103 | |
| DSAG12_00397 | hypothetical protein | -- | 127 | |
| DSAG12_00398 | choloylglycine hydrolase | K01442 | 157 | |
| DSAG12_00399 | hypothetical protein | -- | 202 | |
| DSAG12_00400 | hypothetical protein | -- | 213 | |
| DSAG12_00401 | hypothetical protein | -- | 247 | |
| DSAG12_00402 | ABC-2 type transport system ATP-binding protein | K01990 | 228 | |
| DSAG12_00403 | ribosome-dependent ATPase | K13926 | 259 | |
| DSAG12_00404 | NADPH2:quinone reductase | K00344 | 238 | |
| DSAG12_00405 | hypothetical protein | -- | 242 | |
| DSAG12_00406 | hypothetical protein | -- | 111 | |
| DSAG12_00407 | opine dehydrogenase | K04940 | 91 | |
| DSAG12_00408 | hypothetical protein | -- | 204 | |
| DSAG12_00409 | hypothetical protein | -- | 197 | |
| DSAG12_00410 | hypothetical protein | -- | 101 | |
| DSAG12_00411 | adenine-specific DNA-methyltransferase | K07316 | 178 | |
| DSAG12_00412 | protein phosphatase | K01090 | 200 | |
| DSAG12_00413 | | K06933 | 226 | |
| DSAG12_00414 | ArsR family transcriptional regulator | K03892 | 158 | |
| DSAG12_00415 | hypothetical protein | -- | 178 | |
| DSAG12_00416 | hypothetical protein | -- | 269 | |
| DSAG12_00417 | hypothetical protein | -- | 267 | |
| DSAG12_00418 | hypothetical protein | -- | 245 | |
| DSAG12_00419 | hypothetical protein | -- | 327 | |
| DSAG12_00420 | hypothetical protein | -- | 567 | |
| DSAG12_00421 | UDP-N-acetylglucosamine 3-dehydrogenase | K18855 | 216 | |
| DSAG12_00422 | hypothetical protein | -- | 218 | |
| DSAG12_00423 | hypothetical protein | -- | 144 | |
| DSAG12_00424 | RIO kinase 1 | K07178 | 133 | hypothetical protein with ubiquitin-like domain |
| DSAG12_00425 | hypothetical protein | -- | 215 | |
| DSAG12_00426 | hypothetical protein | -- | 54 | |
| DSAG12_00427 | hypothetical protein | -- | 200 | |
| DSAG12_00428 | hypothetical protein | -- | 199 | |
| DSAG12_00429 | hypothetical protein | -- | 118 | |
| DSAG12_00430 | hypothetical protein | -- | 164 | |
| DSAG12_00431 | hypothetical protein | -- | 165 | |
| DSAG12_00432 | putative pyruvate formate lyase activating enzyme | K04070 | 107 | |
| DSAG12_00433 | hypothetical protein | -- | 204 | |
| DSAG12_00434 | | K07068 | 9541 | |
| DSAG12_00435 | acetyl-CoA C-acetyltransferase | K00626 | 199 | |
| DSAG12_00436 | hypothetical protein | -- | 297 | |
| DSAG12_00437 | hypothetical protein | -- | 186 | |
| DSAG12_00438 | UDPglucose--hexose-1-phosphate uridylyltransferase | K00965 | 283 | |
| DSAG12_00439 | hypothetical protein | -- | 221 | |
| DSAG12_00440 | long-chain acyl-CoA synthetase | K01897 | 222 | |
| DSAG12_00441 | aldehyde:ferredoxin oxidoreductase | K03738 | 248 | |
| DSAG12_00442 | hypothetical protein | -- | 293 | |
| DSAG12_00443 | acetaldehyde dehydrogenase / alcohol dehydrogenase | K04072 | 143 | |
| DSAG12_00444 | hypothetical protein | -- | 249 | |
| DSAG12_00445 | hypothetical protein | -- | 210 | |
| DSAG12_00446 | hypothetical protein | -- | 232 | |
| DSAG12_00447 | hypothetical protein | -- | 254 | |
| DSAG12_00448 | MoxR-like ATPase | K03924 | 251 | |
| DSAG12_00449 | hypothetical protein | -- | 225 | |
| DSAG12_00450 | large conductance mechanosensitive channel | K03282 | 219 | |
| DSAG12_00451 | hypothetical protein | -- | 205 | |
| DSAG12_00452 | hypothetical protein | -- | 249 | |
| DSAG12_00453 | zinc finger protein | K06874 | 50 | |
| DSAG12_00454 | 2-oxoisovalerate ferredoxin oxidoreductase, gamma subunit | K00189 | 218 | |
| DSAG12_00455 | 2-oxoisovalerate ferredoxin oxidoreductase, delta subunit | K00188 | 233 | |
| DSAG12_00456 | pyruvate ferredoxin oxidoreductase, alpha subunit | K00169 | 169 | |
| DSAG12_00457 | pyruvate ferredoxin oxidoreductase, beta subunit | K00170 | 334 | |
| DSAG12_00458 | tRNA acetyltransferase TAN1 | K06963 | 322 | |
| DSAG12_00459 | 6-phosphofructokinase 1 | K00850 | 174 | |
| DSAG12_00460 | hypothetical protein | -- | 263 | |
| DSAG12_00461 | hypothetical protein | -- | 10 | |
| DSAG12_00462 | transposase | K07486 | 46 | |
| DSAG12_00463 | hypothetical protein | -- | 184 | |

| | | | |
|---|---|---|---|
| DSAG12_00464 | hypothetical protein | -- | 157 |
| DSAG12_00465 | hypothetical protein | -- | 237 |
| DSAG12_00466 | 5-methyltetrahydrofolate--homocysteine methyltransferase | K00548 | 233 |
| DSAG12_00467 | 5-methyltetrahydrofolate--homocysteine methyltransferase | K00548 | 173 |
| DSAG12_00468 | methylenetetrahydrofolate reductase (NADPH) | K00297 | 219 |
| DSAG12_00469 | | K01076 | 172 |
| DSAG12_00470 | propionyl-CoA synthetase | K01908 | 236 |
| DSAG12_00471 | hypothetical protein | -- | 221 |
| DSAG12_00472 | hypothetical protein | -- | 223 |
| DSAG12_00473 | hypothetical protein | -- | 173 |
| DSAG12_00474 | aminomethyltransferase | K00605 | 226 |
| DSAG12_00475 | glycine cleavage system H protein | K02437 | 167 |
| DSAG12_00476 | glycine dehydrogenase subunit 1 | K00282 | 221 |
| DSAG12_00477 | glycine dehydrogenase subunit 2 | K00283 | 285 |
| DSAG12_00478 | hypothetical protein | -- | 188 |
| DSAG12_00479 | hypothetical protein | -- | 135 |
| DSAG12_00480 | hypothetical protein | -- | 171 |
| DSAG12_00481 | hypothetical protein | -- | 201 |
| DSAG12_00482 | hypothetical protein | -- | 78 |
| DSAG12_00483 | hypothetical protein | K09707 | 118 |
| DSAG12_00484 | hypothetical protein | -- | 46 |
| DSAG12_00485 | aminotransferase | K10907 | 197 |
| DSAG12_00486 | hypothetical protein | -- | 63 |
| DSAG12_00487 | Ca2+-transporting ATPase | K01537 | 244 |
| DSAG12_00488 | hypothetical protein | -- | 57 |
| DSAG12_00489 | proteasome beta subunit | K03433 | 0 |
| DSAG12_00490 | hypothetical protein | -- | 0 |
| DSAG12_00491 | hypothetical protein | -- | 0 |
| DSAG12_00492 | 4-alpha-glucanotransferase | K00705 | 186 |
| DSAG12_00493 | hypothetical protein | -- | 194 |
| DSAG12_00494 | structural maintenance of chromosome 4 | K06675 | 215 |
| DSAG12_00495 | hypothetical protein | -- | 229 |
| DSAG12_00496 | hypothetical protein | -- | 108 |
| DSAG12_00497 | hypothetical protein | -- | 156 |
| DSAG12_00498 | hypothetical protein | -- | 140 |
| DSAG12_00499 | cold shock protein (beta-ribbon, CspA family) | K03704 | 134 |
| DSAG12_00500 | hypothetical protein | -- | 89 |
| DSAG12_00501 | hypothetical protein | -- | 123 |
| DSAG12_00502 | hypothetical protein | -- | 155 |
| DSAG12_00503 | transcription initiation factor TFIIB | K03124 | 211 |
| DSAG12_00504 | hypothetical protein | -- | 158 |
| DSAG12_00505 | tRNA-Thr | | 102 |
| DSAG12_00506 | aminopeptidase | K01269 | 314 |
| DSAG12_00507 | tRNA-splicing ligase RtcB | K14415 | 0 |
| DSAG12_00508 | hypothetical protein | -- | 265 |
| DSAG12_00509 | 3,4-dihydroxy 2-butanone 4-phosphate synthase / GTP cyclohydrolase II | K14652 | 270 |
| DSAG12_00510 | basic amino acid/polyamine antiporter, APA family | K03294 | 133 |
| DSAG12_00511 | hypothetical protein | -- | 258 |
| DSAG12_00512 | hypothetical protein | -- | 212 |
| DSAG12_00513 | hypothetical protein | -- | 136 |
| DSAG12_00514 | hypothetical protein | -- | 206 |
| DSAG12_00515 | glucosamine--fructose-6-phosphate aminotransferase (isomerizing) | K00820 | 191 |
| DSAG12_00516 | glucosamine--fructose-6-phosphate aminotransferase (isomerizing) | K00820 | 256 |
| DSAG12_00517 | hypothetical protein | -- | 171 |
| DSAG12_00518 | archaeal cell division control protein 6 | K10725 | 186 |
| DSAG12_00519 | hypothetical protein | -- | 150 |
| DSAG12_00520 | DNA polymerase II small subunit | K02323 | 217 |
| DSAG12_00521 | DNA polymerase II large subunit | K02322 | 142 |
| DSAG12_00522 | serine/threonine protein kinase, bacterial | K08884 | 218 |
| DSAG12_00523 | hypothetical protein | -- | 235 |
| DSAG12_00524 | putative hydrolase of the HAD superfamily | K07025 | 0 |
| DSAG12_00525 | threonyl-tRNA synthetase | K01868 | 237 |
| DSAG12_00526 | tRNA (guanine10-N2)-dimethyltransferase | K07446 | 176 |
| DSAG12_00527 | tRNA-Lys | | 128 |
| DSAG12_00528 | large subunit ribosomal protein L2 | K02886 | 103 |
| DSAG12_00529 | S-adenosylmethionine synthetase | K00789 | 229 |
| DSAG12_00530 | hypothetical protein | -- | 0 |
| DSAG12_00531 | hypothetical protein | -- | 273 |
| DSAG12_00532 | hypothetical protein | -- | 236 |
| DSAG12_00533 | hypothetical protein | -- | 85 |
| DSAG12_00534 | hypothetical protein | K09741 | 161 |
| DSAG12_00535 | large subunit ribosomal protein L37Ae | K02921 | 255 |
| DSAG12_00536 | exosome complex component RRP42 | K12589 | 205 |
| DSAG12_00537 | exosome complex component RRP41 | K11600 | 95 |
| DSAG12_00538 | exosome complex component RRP4 | K03679 | 195 |
| DSAG12_00539 | ribosome maturation protein SDO1 | K14574 | 280 |
| DSAG12_00540 | proteasome alpha subunit | K03432 | 282 |
| DSAG12_00541 | hypothetical protein | -- | 192 |

| | | | | |
|---|---|---|---|---|
| DSAG12_00542 | hypothetical protein | -- | 194 | |
| DSAG12_00543 | ribosomal-protein-alanine N-acetyltransferase | K03789 | 236 | |
| DSAG12_00544 | hypothetical protein | -- | 204 | |
| DSAG12_00545 | hypothetical protein | -- | 134 | |
| DSAG12_00546 | tRNA-Val | | 216 | |
| DSAG12_00547 | tricorn protease interacting factor F2/3 | K13722 | 181 | profilin-like domain protein |
| DSAG12_00548 | hypothetical protein | -- | 227 | |
| DSAG12_00549 | Ca2+-transporting ATPase | K01537 | 226 | |
| DSAG12_00550 | beta-lysine 5,6-aminomutase beta subunit | K18011 | 50 | |
| DSAG12_00551 | beta-lysine 5,6-aminomutase alpha subunit | K01844 | 237 | |
| DSAG12_00552 | hypothetical protein | -- | 204 | |
| DSAG12_00553 | hypothetical protein | -- | 267 | |
| DSAG12_00554 | lysine 2,3-aminomutase | K01843 | 173 | |
| DSAG12_00555 | L-erythro-3,5-diaminohexanoate dehydrogenase | K18012 | 180 | |
| DSAG12_00556 | hypothetical protein | -- | 251 | |
| DSAG12_00557 | hypothetical protein | -- | 297 | |
| DSAG12_00558 | alpha-aminoadipic semialdehyde synthase | K14157 | 0 | |
| DSAG12_00559 | saccharopine dehydrogenase (NADP+, L-glutamate forming) | K00293 | 309 | |
| DSAG12_00560 | hypothetical protein | -- | 197 | |
| DSAG12_00561 | hypothetical protein | -- | 249 | |
| DSAG12_00562 | 23S rRNA (adenine2503-C2)-methyltransferase | K06941 | 136 | |
| DSAG12_00563 | L-serine dehydratase | K01752 | 561 | |
| DSAG12_00564 | L-serine dehydratase | K01752 | 0 | |
| DSAG12_00565 | 3-keto-5-aminohexanoate cleavage enzyme | K18013 | 247 | |
| DSAG12_00566 | 3-aminobutyryl-CoA ammonia-lyase | K18014 | 205 | |
| DSAG12_00567 | hypothetical protein | -- | 264 | |
| DSAG12_00568 | diamine N-acetyltransferase | K00657 | 267 | |
| DSAG12_00569 | tRNA-Ser | | 184 | |
| DSAG12_00570 | GTP-binding protein HflX | K03665 | 0 | |
| DSAG12_00571 | putative transcription factor | K03627 | 149 | |
| DSAG12_00572 | hypothetical protein | -- | 151 | |
| DSAG12_00573 | hypothetical protein | -- | 0 | |
| DSAG12_00574 | tyrosine decarboxylase / aspartate 1-decarboxylase | K18933 | 230 | |
| DSAG12_00575 | 3-methyl-2-oxobutanoate hydroxymethyltransferase | K00606 | 194 | |
| DSAG12_00576 | hypothetical protein | -- | 21 | |
| DSAG12_00577 | 2-dehydropantoate 2-reductase | K00077 | 160 | |
| DSAG12_00578 | hypothetical protein | -- | 275 | |
| DSAG12_00579 | hypothetical protein | -- | 726 | |
| DSAG12_00580 | hypothetical protein | -- | 229 | |
| DSAG12_00581 | hypothetical protein | -- | 165 | |
| DSAG12_00582 | transcription initiation factor TFIIB | K03124 | 205 | |
| DSAG12_00583 | hypothetical protein | -- | 151 | |
| DSAG12_00584 | translation initiation factor 1 | K03113 | 274 | |
| DSAG12_00585 | | K06885 | 131 | |
| DSAG12_00586 | phosphoadenosine phosphosulfate reductase | K00390 | 221 | |
| DSAG12_00587 | Rab family, other | K07976 | 0 | |
| DSAG12_00588 | hypothetical protein | -- | 127 | |
| DSAG12_00589 | hypothetical protein | -- | 217 | |
| DSAG12_00590 | hypothetical protein | -- | 225 | |
| DSAG12_00591 | uridylate kinase | K09903 | 236 | |
| DSAG12_00592 | pantoate kinase | K06982 | 239 | small GTP-binding domain protein |
| DSAG12_00593 | peptide chain release factor subunit 1 | K03265 | 116 | |
| DSAG12_00594 | 4-phosphopantoate---beta-alanine ligase | K09722 | 72 | |
| DSAG12_00595 | peptide alpha-N-acetyltransferase | K00670 | 159 | |
| DSAG12_00596 | Ras-related GTP-binding protein A/B | K16185 | 221 | |
| DSAG12_00597 | 23S ribosomal RNA | -- | 186 | |
| DSAG12_00598 | hypothetical protein | -- | 202 | |
| DSAG12_00599 | internalin A | K13730 | 197 | |
| DSAG12_00600 | internalin A | K13730 | 196 | |
| DSAG12_00601 | hypothetical protein | -- | 190 | small GTP-binding domain protein |
| DSAG12_00602 | hypothetical protein | -- | 0 | |
| DSAG12_00603 | hypothetical protein | -- | 0 | |
| DSAG12_00604 | hypothetical protein | -- | 166 | |
| DSAG12_00605 | hypothetical protein | -- | 244 | |
| DSAG12_00606 | hypothetical protein | -- | 0 | |
| DSAG12_00607 | hypothetical protein | -- | 184 | |
| DSAG12_00608 | | K06885 | 99 | |
| DSAG12_00609 | hypothetical protein | -- | 199 | |
| DSAG12_00610 | hypothetical protein | -- | 217 | |
| DSAG12_00611 | hypothetical protein | -- | 119 | |
| DSAG12_00612 | hypothetical protein | -- | 182 | |
| DSAG12_00613 | hypothetical protein | -- | 151 | |
| DSAG12_00614 | aspartate racemase | K01779 | 160 | |
| DSAG12_00615 | hypothetical protein | -- | 183 | |
| DSAG12_00616 | hypothetical protein | -- | 289 | |
| DSAG12_00617 | hypothetical protein | -- | 0 | |
| DSAG12_00618 | aminoglycoside N6'-acetyltransferase | K00663 | 164 | |
| DSAG12_00619 | hypothetical protein | -- | 0 | |

| | | | | |
|---|---|---|---|---|
| DSAG12_00620 | hypothetical protein | -- | 128 | |
| DSAG12_00621 | hypothetical protein | -- | 130 | |
| DSAG12_00622 | hypothetical protein | -- | 190 | |
| DSAG12_00623 | hypothetical protein | -- | 225 | |
| DSAG12_00624 | hypothetical protein | -- | 186 | |
| DSAG12_00625 | hypothetical protein | -- | 132 | |
| DSAG12_00626 | hypothetical protein | -- | 187 | |
| DSAG12_00627 | proteasome regulatory subunit | K03420 | 166 | |
| DSAG12_00628 | conserved protein with predicted RNA binding PUA domain | K07398 | 152 | |
| DSAG12_00629 | nascent polypeptide-associated complex subunit alpha | K03626 | 162 | |
| DSAG12_00630 | hypothetical protein | -- | 78 | |
| DSAG12_00631 | hypothetical protein | -- | 145 | |
| DSAG12_00632 | hypothetical protein | -- | 106 | hypothetical protein with vacuolar fusion domain MON1 |
| DSAG12_00633 | hypothetical protein | -- | 25 | |
| DSAG12_00634 | hypothetical protein | -- | 161 | |
| DSAG12_00635 | hypothetical protein | -- | 298 | |
| DSAG12_00636 | hypothetical protein | -- | 140 | |
| DSAG12_00637 | hypothetical protein | -- | 190 | |
| DSAG12_00638 | diaminopimelate decarboxylase | K01586 | 116 | |
| DSAG12_00639 | cell division protein FtsZ | K03531 | 86 | |
| DSAG12_00640 | ribosomal RNA assembly protein | K06961 | 155 | |
| DSAG12_00641 | RIO kinase 1 | K07178 | 238 | |
| DSAG12_00642 | nonsense-mediated mRNA decay protein 3 | K07562 | 85 | |
| DSAG12_00643 | hypothetical protein | K09148 | 0 | |
| DSAG12_00644 | translation initiation factor 2 subunit 2 | K03238 | 112 | |
| DSAG12_00645 | replication factor A2 | K10739 | 87 | |
| DSAG12_00646 | tRNA-Met | Intron(70709 | 145 | |
| DSAG12_00647 | hypothetical protein | -- | 148 | |
| DSAG12_00648 | hypothetical protein | -- | 255 | |
| DSAG12_00649 | hypothetical protein | -- | 147 | |
| DSAG12_00650 | replication factor A1 | K07466 | 146 | |
| DSAG12_00651 | fructokinase | K00847 | 132 | |
| DSAG12_00652 | protein-L-isoaspartate(D-aspartate) O-methyltransferase | K00573 | 83 | |
| DSAG12_00653 | thiamine biosynthesis protein ThiI | K03151 | 137 | |
| DSAG12_00654 | | K07159 | 173 | |
| DSAG12_00655 | tRNA-Asn | | 161 | |
| DSAG12_00656 | hypothetical protein | -- | 163 | |
| DSAG12_00657 | tRNA-Glu | Intron(71654 | 144 | domain |
| DSAG12_00658 | hypothetical protein | -- | 161 | |
| DSAG12_00659 | tRNA-Leu | | 196 | |
| DSAG12_00660 | pre-rRNA-processing protein TSR3 | K09140 | 213 | |
| DSAG12_00661 | hypothetical protein | -- | 129 | |
| DSAG12_00662 | hypothetical protein | -- | 189 | |
| DSAG12_00663 | | K06942 | 0 | |
| DSAG12_00664 | tRNA-Ala | | 26 | |
| DSAG12_00665 | 16S ribosomal RNA | | 212 | |
| DSAG12_00666 | hypothetical protein | -- | 153 | |
| DSAG12_00667 | hypothetical protein | -- | 207 | |
| DSAG12_00668 | | K06883 | 125 | |
| DSAG12_00669 | hypothetical protein | -- | 169 | |
| DSAG12_00670 | hypothetical protein | -- | 0 | |
| DSAG12_00671 | hypothetical protein | -- | 0 | |
| DSAG12_00672 | hypothetical protein | -- | 126 | |
| DSAG12_00673 | transcription initiation factor TFIIB | K03124 | 118 | |
| DSAG12_00674 | threonylcarbamoyladenosine tRNA methylthiotransferase CDKAL1 | K15865 | 126 | small GTP-binding domain protein |
| DSAG12_00675 | tRNA-Glu | | 76 | |
| DSAG12_00676 | UPF0148 protein | K07143 | 97 | |
| DSAG12_00677 | small subunit ribosomal protein S17e | K02962 | 84 | |
| DSAG12_00678 | hypothetical protein | -- | 64 | |
| DSAG12_00679 | hypothetical protein | -- | 119 | |
| DSAG12_00680 | archaeal cell division control protein 6 | K10725 | 124 | |
| DSAG12_00681 | hypothetical protein | -- | 0 | |
| DSAG12_00682 | hypothetical protein | -- | 106 | |
| DSAG12_00683 | archaeal flagellar protein FlaI | K07332 | 62 | |
| DSAG12_00684 | hypothetical protein | -- | 120 | |
| DSAG12_00685 | hypothetical protein | -- | 16 | |
| DSAG12_00686 | large subunit ribosomal protein L23 | K02892 | 100 | |
| DSAG12_00687 | large subunit ribosomal protein L4e | K02930 | 32 | |
| DSAG12_00688 | large subunit ribosomal protein L3 | K02906 | 48 | |
| DSAG12_00689 | ATP-binding cassette, sub-family E, member 1 | K06174 | 107 | |
| DSAG12_00690 | lysyl-tRNA synthetase, class I | K04566 | 61 | |
| DSAG12_00691 | hypothetical protein | -- | 98 | |
| DSAG12_00692 | hypothetical protein | -- | 87 | |
| DSAG12_00693 | | K06940 | 349 | |
| DSAG12_00694 | hypothetical protein | -- | 263 | |
| DSAG12_00695 | aspartate carbamoyltransferase regulatory subunit | K00610 | 204 | |
| DSAG12_00696 | hypothetical protein | -- | 189 | |

| | | | |
|---|---|---|---|
| DSAG12_00697 | hypothetical protein | -- | 95 |
| DSAG12_00698 | hypothetical protein | K07746 | 179 |
| DSAG12_00699 | hypothetical protein | -- | 0 |
| DSAG12_00700 | hypothetical protein | -- | 71 |
| DSAG12_00701 | hypothetical protein | K07746 | 115 |
| DSAG12_00702 | hypothetical protein | -- | 119 |
| DSAG12_00703 | hypothetical protein | -- | 187 |
| DSAG12_00704 | thymidylate synthase (FAD) | K03465 | 161 |
| DSAG12_00705 | hypothetical protein | -- | 18 |
| DSAG12_00706 | hypothetical protein | -- | 126 |
| DSAG12_00707 | hypothetical protein | -- | 179 |
| DSAG12_00708 | phosphomethylpyrimidine synthase | K03147 | 22 |
| DSAG12_00709 | histidine triad (HIT) family protein | K02503 | 133 |
| DSAG12_00710 | hypothetical protein | -- | 128 |
| DSAG12_00711 | DNA polymerase II | K02336 | 186 |
| DSAG12_00712 | DNA repair protein RadA | K04483 | 179 |
| DSAG12_00713 | 3-dehydroquinate synthase | K01735 | 141 |
| DSAG12_00714 | hypothetical protein | -- | 188 |
| DSAG12_00715 | hypothetical protein | -- | 224 |
| DSAG12_00716 | hypothetical protein | -- | 181 |
| DSAG12_00717 | | K07131 | 63 |
| DSAG12_00718 | elongation factor 1-alpha | K03231 | 259 |
| DSAG12_00719 | hypothetical protein | -- | 217 |
| DSAG12_00720 | hypothetical protein | -- | 121 |
| DSAG12_00721 | peroxisomal coenzyme A diphosphatase NUDT7 | K17879 | 160 |
| DSAG12_00722 | putative mRNA 3-end processing factor | K07577 | 181 |
| DSAG12_00723 | 5'-methylthioadenosine phosphorylase | K00772 | 122 |
| DSAG12_00724 | hypoxanthine phosphoribosyltransferase | K00760 | 139 |
| DSAG12_00725 | carbonic anhydrase | K01673 | 178 |
| DSAG12_00726 | adenine deaminase | K01486 | 158 |
| DSAG12_00727 | Ca-activated chloride channel homolog | K07114 | 0 |
| DSAG12_00728 | thiamine transport system permease protein | K02063 | 98 |
| DSAG12_00729 | spermidine/putrescine transport system ATP-binding protein | K11072 | 150 |
| DSAG12_00730 | inositol-phosphate transport system ATP-binding protein | K17240 | 226 |
| DSAG12_00731 | hypothetical protein | K07220 | 245 |
| DSAG12_00732 | tRNA-Ile | | 162 |
| DSAG12_00733 | sulfoxide reductase catalytic subunit YedY | K07147 | 144 |
| DSAG12_00734 | hypothetical protein | -- | 202 |
| DSAG12_00735 | hypothetical protein | -- | 192 |
| DSAG12_00736 | hypothetical protein | -- | 231 |
| DSAG12_00737 | secernin | K14358 | 207 |
| DSAG12_00738 | hypothetical protein | -- | 234 |
| DSAG12_00739 | hypothetical protein | -- | 235 |
| DSAG12_00740 | adenylate kinase | K18532 | 0 |
| DSAG12_00741 | phenylacetic acid degradation protein | K02617 | 198 |
| DSAG12_00742 | Ca-activated chloride channel homolog | K07114 | 280 |
| DSAG12_00743 | hypothetical protein | -- | 157 |
| DSAG12_00744 | hypothetical protein | -- | 165 |
| DSAG12_00745 | hypothetical protein | -- | 243 |
| DSAG12_00746 | hypothetical protein | -- | 196 |
| DSAG12_00747 | serine/threonine protein kinase, bacterial | K08884 | 161 |
| DSAG12_00748 | protein phosphatase | K01090 | 134 |
| DSAG12_00749 | serine/threonine-protein phosphatase 5 | K04460 | 175 |
| DSAG12_00750 | serine/threonine-protein phosphatase PP1 catalytic subunit | K06269 | 220 |
| DSAG12_00751 | 7-cyano-7-deazaguanine tRNA-ribosyltransferase | K18779 | 243 |
| DSAG12_00752 | hypothetical protein | K09738 | 135 |
| DSAG12_00753 | tRNA (guanine37-N1)-methyltransferase | K15429 | 181 |
| DSAG12_00754 | hypothetical protein | -- | 154 |
| DSAG12_00755 | hypothetical protein | -- | 244 |
| DSAG12_00756 | hypothetical protein | -- | 216 |
| DSAG12_00757 | tRNA-Arg | | 165 |
| DSAG12_00758 | hypothetical protein | -- | 135 |
| DSAG12_00759 | phenylacetic acid degradation protein | K02617 | 193 |
| DSAG12_00760 | transcription initiation factor TFIIB | K03124 | 111 |
| DSAG12_00761 | Ras-related GTP-binding protein A/B | K16185 | 148 |
| DSAG12_00762 | foldase protein PrsA | K07533 | 221 |
| DSAG12_00763 | putative hydrolase of the HAD superfamily | K07025 | 201 |
| DSAG12_00764 | hypothetical protein | -- | 183 |
| DSAG12_00765 | myo-inositol-1-phosphate synthase | K01858 | 0 |
| DSAG12_00766 | hypothetical protein | -- | 132 |
| DSAG12_00767 | hypothetical protein | -- | 249 |
| DSAG12_00768 | hypothetical protein | -- | 297 |
| DSAG12_00769 | ribonuclease P/MRP protein subunit POP5 | K03537 | 128 |
| DSAG12_00770 | ribonuclease P/MRP protein subunit RPP1 | K03539 | 132 |
| DSAG12_00771 | hypothetical protein | K07581 | 121 |
| DSAG12_00772 | large subunit ribosomal protein L15e | K02877 | 186 |
| DSAG12_00773 | | K06883 | 209 |
| DSAG12_00774 | hypothetical protein | -- | 146 |

| | | | | |
|---|---|---|---|---|
| DSAG12_00775 | enoyl-CoA hydratase | K01715 | 0 | |
| DSAG12_00776 | seryl-tRNA synthetase | K01875 | 114 | |
| DSAG12_00777 | hypothetical protein | -- | 98 | |
| DSAG12_00778 | phytol kinase | K18678 | 142 | |
| DSAG12_00779 | geranylgeranylglycerol-phosphate geranylgeranyltransferase | K17105 | 186 | |
| DSAG12_00780 | hypothetical protein | -- | 135 | |
| DSAG12_00781 | L-threonylcarbamoyladenylate synthase | K07566 | 184 | |
| DSAG12_00782 | hypothetical protein | -- | 232 | small GTP-binding domain protein |
| DSAG12_00783 | hypothetical protein | -- | 122 | |
| DSAG12_00784 | hypothetical protein | -- | 161 | |
| DSAG12_00785 | hypothetical protein | -- | 217 | |
| DSAG12_00786 | hypothetical protein | -- | 202 | |
| DSAG12_00787 | hypothetical protein | -- | 180 | |
| DSAG12_00788 | MFS transporter, DHA1 family, multidrug resistance protein | K08153 | 184 | |
| DSAG12_00789 | hypothetical protein | -- | 254 | |
| DSAG12_00790 | hypothetical protein | -- | 217 | |
| DSAG12_00791 | hypothetical protein | -- | 49 | |
| DSAG12_00792 | hypothetical protein | -- | 0 | |
| DSAG12_00793 | hypothetical protein | -- | 191 | |
| DSAG12_00794 | hypothetical protein | -- | 309 | |
| DSAG12_00795 | hypothetical protein | -- | 151 | |
| DSAG12_00796 | pyruvate ferredoxin oxidoreductase, beta subunit | K00170 | 124 | |
| DSAG12_00797 | pyruvate ferredoxin oxidoreductase, alpha subunit | K00169 | 0 | |
| DSAG12_00798 | pyruvate ferredoxin oxidoreductase, gamma subunit | K00172 | 12 | |
| DSAG12_00799 | hypothetical protein | -- | 174 | |
| DSAG12_00800 | hypothetical protein | -- | 141 | |
| DSAG12_00801 | hypothetical protein | -- | 185 | |
| DSAG12_00802 | Ras-related protein Rab-22 | K07891 | 77 | |
| DSAG12_00803 | glutamate formiminotransferase / formiminotetrahydrofolate cyclodeaminase | K13990 | 145 | |
| DSAG12_00804 | hypothetical protein | -- | 144 | |
| DSAG12_00805 | UDP-N-acetylglucosamine 3-dehydrogenase | K18855 | 140 | |
| DSAG12_00806 | segregation and condensation protein B | K06024 | 93 | |
| DSAG12_00807 | hypothetical protein | -- | 242 | |
| DSAG12_00808 | chromosome segregation protein | K03529 | 326 | |
| DSAG12_00809 | hypothetical protein | -- | 266 | |
| DSAG12_00810 | hypothetical protein | -- | 121 | |
| DSAG12_00811 | hypothetical protein | -- | 90 | |
| DSAG12_00812 | hypothetical protein | -- | 175 | |
| DSAG12_00813 | hypothetical protein | -- | 143 | |
| DSAG12_00814 | hypothetical protein | -- | 281 | |
| DSAG12_00815 | hypothetical protein | -- | 202 | |
| DSAG12_00816 | hypothetical protein | -- | 249 | |
| DSAG12_00817 | erbb2-interacting protein | K12796 | 255 | |
| DSAG12_00818 | internalin A | K13730 | 282 | |
| DSAG12_00819 | hypothetical protein | -- | 317 | |
| DSAG12_00820 | citronellol/citronellal dehydrogenase | K13775 | 201 | |
| DSAG12_00821 | hypothetical protein | -- | 71 | |
| DSAG12_00822 | hypothetical protein | -- | 216 | |
| DSAG12_00823 | oligogalacturonide transporter | K16210 | 226 | |
| DSAG12_00824 | hypothetical protein | -- | 245 | |
| DSAG12_00825 | | K06889 | 0 | |
| DSAG12_00826 | hypothetical protein | -- | 253 | |
| DSAG12_00827 | hypothetical protein | -- | 175 | |
| DSAG12_00828 | hypothetical protein | -- | 138 | |
| DSAG12_00829 | hypothetical protein | -- | 64 | |
| DSAG12_00830 | hypothetical protein | -- | 131 | |
| DSAG12_00831 | inorganic pyrophosphatase | K01507 | 170 | |
| DSAG12_00832 | hypothetical protein | -- | 291 | |
| DSAG12_00833 | DNA replication factor GINS | K09723 | 244 | |
| DSAG12_00834 | hypothetical protein | -- | 220 | |
| DSAG12_00835 | hypothetical protein | -- | 0 | |
| DSAG12_00836 | hypothetical protein | -- | 302 | |
| DSAG12_00837 | alpha-mannosidase | K01191 | 166 | |
| DSAG12_00838 | pyruvate formate lyase activating enzyme | K04069 | 235 | |
| DSAG12_00839 | hypothetical protein | -- | 63 | |
| DSAG12_00840 | hypothetical protein | -- | 66 | |
| DSAG12_00841 | 2-haloacid dehalogenase | K01560 | 144 | |
| DSAG12_00842 | hypothetical protein | -- | 234 | |
| DSAG12_00843 | Ras-related GTP-binding protein A/B | K16185 | 140 | |
| DSAG12_00844 | hypothetical protein | -- | 165 | |
| DSAG12_00845 | hypothetical protein | -- | 193 | |
| DSAG12_00846 | hypothetical protein | -- | 186 | |
| DSAG12_00847 | hypothetical protein | -- | 179 | |
| DSAG12_00848 | 1-acyl-sn-glycerol-3-phosphate acyltransferase | K00655 | 157 | |
| DSAG12_00849 | hypothetical protein | -- | 142 | |
| DSAG12_00850 | hypothetical protein | -- | 150 | |
| DSAG12_00851 | hypothetical protein | -- | 224 | |
| DSAG12_00852 | 2-hydroxy-3-oxopropionate reductase | K00042 | 196 | |

| | | | | |
|---|---|---|---|---|
| DSAG12_00853 | hypothetical protein | -- | 204 | |
| DSAG12_00854 | glycoside/pentoside/hexuronide:cation symporter, GPH family | K03292 | 160 | |
| DSAG12_00855 | hypothetical protein | -- | 175 | |
| DSAG12_00856 | | K00540 | 200 | |
| DSAG12_00857 | beta-galactosidase | K01190 | 22 | |
| DSAG12_00858 | hypothetical protein | -- | 227 | |
| DSAG12_00859 | zinc transport system ATP-binding protein | K09817 | 228 | |
| DSAG12_00860 | manganese/iron transport system permease protein | K09819 | 191 | |
| DSAG12_00861 | zinc/manganese transport system substrate-binding protein | K02077 | 217 | |
| DSAG12_00862 | DtxR family transcriptional regulator, Mn-dependent transcriptional regulator | K03709 | 175 | |
| DSAG12_00863 | molybdopterin synthase sulfur carrier subunit | K03636 | 169 | |
| DSAG12_00864 | hypothetical protein | -- | 42 | |
| DSAG12_00865 | hypothetical protein | -- | 239 | |
| DSAG12_00866 | hypothetical protein | -- | 250 | |
| DSAG12_00867 | hypothetical protein | -- | 181 | |
| DSAG12_00868 | hypothetical protein | -- | 224 | |
| DSAG12_00869 | methionyl-tRNA synthetase | K01874 | 209 | |
| DSAG12_00870 | ribonucleoside-diphosphate reductase alpha chain | K00525 | 214 | |
| DSAG12_00871 | Fe-S cluster assembly ATP-binding protein | K09013 | 135 | |
| DSAG12_00872 | | K07033 | 190 | |
| DSAG12_00873 | hypothetical protein | -- | 221 | |
| DSAG12_00874 | release factor glutamine methyltransferase | K02493 | 210 | |
| DSAG12_00875 | 16S rRNA (adenine1518-N6/adenine1519-N6)-dimethyltransferase | K02528 | 131 | |
| DSAG12_00876 | putative nucleotide binding protein | K07572 | 106 | |
| DSAG12_00877 | DNA-directed RNA polymerase subunit F | K03051 | 122 | |
| DSAG12_00878 | large subunit ribosomal protein L21e | K02889 | 233 | |
| DSAG12_00879 | endoglucanase | K01179 | 142 | |
| DSAG12_00880 | hypothetical protein | -- | 156 | |
| DSAG12_00881 | hypothetical protein | -- | 162 | |
| DSAG12_00882 | hypothetical protein | -- | 127 | |
| DSAG12_00883 | flotillin | K07192 | 260 | |
| DSAG12_00884 | hypothetical protein | -- | 193 | |
| DSAG12_00885 | | K07047 | 255 | |
| DSAG12_00886 | hypothetical protein | -- | 142 | |
| DSAG12_00887 | hypothetical protein | -- | 173 | |
| DSAG12_00888 | hypothetical protein | -- | 172 | |
| DSAG12_00889 | tRNA pseudouridine synthase 10 | K07583 | 202 | |
| DSAG12_00890 | adenylyltransferase and sulfurtransferase | K11996 | 149 | |
| DSAG12_00891 | hypothetical protein | -- | 199 | |
| DSAG12_00892 | hypothetical protein | -- | 269 | |
| DSAG12_00893 | excinuclease ABC subunit C | K03703 | 171 | |
| DSAG12_00894 | hypothetical protein | -- | 196 | |
| DSAG12_00895 | hypothetical protein | -- | 274 | |
| DSAG12_00896 | Xaa-Pro aminopeptidase | K01262 | 258 | |
| DSAG12_00897 | hypothetical protein | -- | 145 | |
| DSAG12_00898 | hypothetical protein | -- | 233 | |
| DSAG12_00899 | hypothetical protein | -- | 156 | |
| DSAG12_00900 | hypothetical protein | -- | 212 | |
| DSAG12_00901 | hypothetical protein | -- | 34 | |
| DSAG12_00902 | hypothetical protein | -- | 210 | |
| DSAG12_00903 | tryptophanyl-tRNA synthetase | K01867 | 193 | |
| DSAG12_00904 | protein transport protein SEC61 subunit alpha | K10956 | 44 | |
| DSAG12_00905 | hypothetical protein | -- | 233 | |
| DSAG12_00906 | hypothetical protein | -- | 222 | |
| DSAG12_00907 | hypothetical protein | -- | 191 | |
| DSAG12_00908 | | K01554 | 245 | |
| DSAG12_00909 | hypothetical protein | -- | 258 | |
| DSAG12_00910 | hypothetical protein | -- | 84 | |
| DSAG12_00911 | hypothetical protein | -- | 176 | |
| DSAG12_00912 | | K06937 | 119 | |
| DSAG12_00913 | hypothetical protein | -- | 104 | |
| DSAG12_00914 | hypothetical protein | -- | 156 | |
| DSAG12_00915 | proteasome alpha subunit | K03432 | 163 | |
| DSAG12_00916 | hypothetical protein | -- | 239 | |
| DSAG12_00917 | hypothetical protein | -- | 304 | |
| DSAG12_00918 | hypothetical protein | -- | 197 | |
| DSAG12_00919 | hypothetical protein | -- | 124 | |
| DSAG12_00920 | sphingosine-1-phosphate phosphatase 1 | K04716 | 194 | |
| DSAG12_00921 | hypothetical protein | -- | 213 | |
| DSAG12_00922 | hypothetical protein | -- | 112 | |
| DSAG12_00923 | hypothetical protein | -- | 165 | |
| DSAG12_00924 | hypothetical protein | -- | 275 | |
| DSAG12_00925 | hypothetical protein | -- | 322 | |
| DSAG12_00926 | hypothetical protein | -- | 213 | |
| DSAG12_00927 | hypothetical protein | -- | 230 | |
| DSAG12_00928 | hypothetical protein | -- | 266 | |
| DSAG12_00929 | | K06915 | 212 | small GTP-binding domain protein |
| DSAG12_00930 | hypothetical protein | -- | 182 | |

| | | | | |
|---|---|---|---|---|
| DSAG12_00931 | hypothetical protein | -- | 116 | |
| DSAG12_00932 | ATP-binding protein involved in chromosome partitioning | K03593 | 185 | |
| DSAG12_00933 | | K07096 | 253 | |
| DSAG12_00934 | hypothetical protein | -- | 197 | |
| DSAG12_00935 | | K00924 | 175 | |
| DSAG12_00936 | hypothetical protein | -- | 56 | |
| DSAG12_00937 | hypothetical protein | -- | 206 | |
| DSAG12_00938 | glycerophosphoryl diester phosphodiesterase | K01126 | 182 | |
| DSAG12_00939 | hypothetical protein | -- | 177 | |
| DSAG12_00940 | hypothetical protein | -- | 185 | |
| DSAG12_00941 | hypothetical protein | -- | 227 | |
| DSAG12_00942 | adenylyltransferase and sulfurtransferase | K11996 | 163 | |
| DSAG12_00943 | hypothetical protein | -- | 299 | |
| DSAG12_00944 | actin beta/gamma 1 | K05692 | 213 | |
| DSAG12_00945 | hypothetical protein | -- | 285 | |
| DSAG12_00946 | | K01175 | 244 | |
| DSAG12_00947 | hypothetical protein | -- | 133 | |
| DSAG12_00948 | hypothetical protein | -- | 65 | |
| DSAG12_00949 | hypothetical protein | -- | 160 | |
| DSAG12_00950 | NAD+ synthase | K01916 | 211 | |
| DSAG12_00951 | hypothetical protein | -- | 173 | |
| DSAG12_00952 | hypothetical protein | -- | 180 | |
| DSAG12_00953 | hypothetical protein | -- | 207 | |
| DSAG12_00954 | hypothetical protein | -- | 145 | |
| DSAG12_00955 | hypothetical protein | -- | 436 | Lokiactin |
| DSAG12_00956 | 3-hydroxypropionyl-coenzyme A dehydratase | K15019 | 89 | |
| DSAG12_00957 | septum formation protein | K06287 | 192 | |
| DSAG12_00958 | hypothetical protein | -- | 174 | |
| DSAG12_00959 | hypothetical protein | -- | 199 | |
| DSAG12_00960 | tRNA-Pro | | 191 | |
| DSAG12_00961 | hypothetical protein | -- | 197 | |
| DSAG12_00962 | ADP-ribose pyrophosphatase | K01515 | 152 | |
| DSAG12_00963 | membrane dipeptidase | K01273 | 175 | |
| DSAG12_00964 | Ras-related protein Rab-5C | K07889 | 142 | |
| DSAG12_00965 | hypothetical protein | -- | 175 | |
| DSAG12_00966 | hypothetical protein | -- | 125 | |
| DSAG12_00967 | adenine-specific DNA-methyltransferase | K07316 | 201 | |
| DSAG12_00968 | magnesium transporter | K03284 | 216 | |
| DSAG12_00969 | elongation factor 1-alpha | K03231 | 157 | |
| DSAG12_00970 | hypothetical protein | -- | 241 | |
| DSAG12_00971 | hypothetical protein | -- | 0 | |
| DSAG12_00972 | hypothetical protein | -- | 163 | |
| DSAG12_00973 | 2-iminobutanoate/2-iminopropanoate deaminase | K09022 | 214 | |
| DSAG12_00974 | hypothetical protein | -- | 247 | |
| DSAG12_00975 | phosphoglycolate phosphatase | K01091 | 209 | small GTP-binding domain protein |
| DSAG12_00976 | ATP-dependent RNA helicase DeaD | K05592 | 212 | |
| DSAG12_00977 | hypothetical protein | -- | 184 | |
| DSAG12_00978 | hypothetical protein | K09131 | 165 | |
| DSAG12_00979 | hypothetical protein | K09717 | 239 | |
| DSAG12_00980 | archaea-specific RecJ-like exonuclease | K07463 | 201 | |
| DSAG12_00981 | hypothetical protein | -- | 269 | |
| DSAG12_00982 | hypothetical protein | -- | 180 | |
| DSAG12_00983 | hypothetical protein | -- | 114 | |
| DSAG12_00984 | 1,3-propanediol dehydrogenase | K00086 | 198 | |
| DSAG12_00985 | ubiquitin-conjugating enzyme E2 H | K10576 | 163 | |
| DSAG12_00986 | hypothetical protein | -- | 267 | |
| DSAG12_00987 | chlorophyll(ide) b reductase | K13606 | 233 | |
| DSAG12_00988 | hypothetical protein | -- | 90 | |
| DSAG12_00989 | hypothetical protein | -- | 168 | |
| DSAG12_00990 | hypothetical protein | -- | 177 | |
| DSAG12_00991 | hypothetical protein | -- | 191 | |
| DSAG12_00992 | hypothetical protein | -- | 82 | |
| DSAG12_00993 | dihydroflavonol-4-reductase | K00091 | 118 | |
| DSAG12_00994 | 3-oxo-5-alpha-steroid 4-dehydrogenase 1 | K12343 | 237 | |
| DSAG12_00995 | thiamine biosynthesis protein ThiI | K03151 | 232 | hypothetical protein with ubiquitin-conjugating domain |
| DSAG12_00996 | hypothetical protein | -- | 171 | |
| DSAG12_00997 | hypothetical protein | -- | 208 | |
| DSAG12_00998 | hypothetical protein | -- | 200 | |
| DSAG12_00999 | hypothetical protein | -- | 133 | |
| DSAG12_01000 | hypothetical protein | -- | 197 | |
| DSAG12_01001 | hypothetical protein | -- | 118 | |
| DSAG12_01002 | nicotinamidase/pyrazinamidase | K08281 | 62 | |
| DSAG12_01003 | cyclic pyranopterin phosphate synthase | K03639 | 222 | |
| DSAG12_01004 | hypothetical protein | -- | 220 | |
| DSAG12_01005 | sulfate permease, SulP family | K03321 | 163 | |
| DSAG12_01006 | putative membrane protein | K08981 | 242 | |
| DSAG12_01007 | hypothetical protein | -- | 192 | |

| | | | |
|---|---|---|---|
| DSAG12_01008 | hypothetical protein | -- | 206 |
| DSAG12_01009 | hypothetical protein | -- | 159 |
| DSAG12_01010 | acyl-CoA thioester hydrolase | K07107 | 0 |
| DSAG12_01011 | | K01362 | 208 |
| DSAG12_01012 | hypothetical protein | -- | 188 |
| DSAG12_01013 | hypothetical protein | -- | 231 |
| DSAG12_01014 | demethylmenaquinone methyltransferase / 2-methoxy-6-polyprenyl-1,4-benzoquinol methylase | K03183 | 217 |
| DSAG12_01015 | | K07124 | 0 |
| DSAG12_01016 | hypothetical protein | -- | 68 |
| DSAG12_01017 | hypothetical protein | -- | 188 |
| DSAG12_01018 | hypothetical protein | -- | 270 |
| DSAG12_01019 | hypothetical protein | -- | 133 |
| DSAG12_01020 | hypothetical protein | K09116 | 152 |
| DSAG12_01021 | oligo-1,6-glucosidase | K01182 | 215 |
| DSAG12_01022 | hypothetical protein | -- | 183 |
| DSAG12_01023 | hypothetical protein | -- | 329 |
| DSAG12_01024 | hypothetical protein | -- | 244 |
| DSAG12_01025 | hypothetical protein | -- | 99 |
| DSAG12_01026 | hypothetical protein | -- | 136 |
| DSAG12_01027 | hypothetical protein | -- | 220 |
| DSAG12_01028 | hypothetical protein | -- | 315 |
| DSAG12_01029 | hypothetical protein | -- | 230 |
| DSAG12_01030 | long-chain acyl-CoA synthetase | K01897 | 238 |
| DSAG12_01031 | hypothetical protein | K09157 | 190 |
| DSAG12_01032 | hypothetical protein | -- | 230 |
| DSAG12_01033 | Lrp/AsnC family transcriptional regulator | K05800 | 269 |
| DSAG12_01034 | asparaginyl-tRNA synthetase | K01893 | 190 |
| DSAG12_01035 | threonine aldolase | K01620 | 234 |
| DSAG12_01036 | hypothetical protein | -- | 336 |
| DSAG12_01037 | beta-galactosidase | K12308 | 266 |
| DSAG12_01038 | adenosine kinase | K00856 | 151 |
| DSAG12_01039 | glycoside/pentoside/hexuronide:cation symporter, GPH family | K03292 | 67 |
| DSAG12_01040 | MFS transporter, DHA3 family, macrolide efflux protein | K08217 | 271 |
| DSAG12_01041 | molybdopterin synthase sulfur carrier subunit | K03636 | 340 |
| DSAG12_01042 | hypothetical protein | -- | 228 |
| DSAG12_01043 | hypothetical protein | -- | 177 |
| DSAG12_01044 | hypothetical protein | -- | 128 |
| DSAG12_01045 | NTE family protein | K07001 | 383 |
| DSAG12_01046 | hypothetical protein | -- | 237 |
| DSAG12_01047 | hypothetical protein | -- | 113 |
| DSAG12_01048 | hypothetical protein | -- | 166 |
| DSAG12_01049 | hypothetical protein | -- | 210 |
| DSAG12_01050 | tRNA-Cys | | 224 |
| DSAG12_01051 | hypothetical protein | -- | 207 |
| DSAG12_01052 | hypothetical protein | -- | 109 |
| DSAG12_01053 | hypothetical protein | -- | 502 |
| DSAG12_01054 | cysteinyl-tRNA synthetase | K01883 | 283 |
| DSAG12_01055 | voltage-gated potassium channel | K10716 | 639 |
| DSAG12_01056 | hypothetical protein | -- | 244 |
| DSAG12_01057 | hypothetical protein | -- | 283 |
| DSAG12_01058 | transposase | K07486 | 831 |
| DSAG12_01059 | hypothetical protein | -- | 379 |
| DSAG12_01060 | hypothetical protein | -- | 209 |
| DSAG12_01061 | MFS transporter, UMF1 family | K06902 | 0 |
| DSAG12_01062 | hypothetical protein | -- | 202 |
| DSAG12_01063 | hypothetical protein | -- | 239 |
| DSAG12_01064 | hypothetical protein | -- | 120 |
| DSAG12_01065 | glycerol-3-phosphate dehydrogenase (NAD(P)+) | K00057 | 192 |
| DSAG12_01066 | Cu+-exporting ATPase | K17686 | 191 |
| DSAG12_01067 | hypothetical protein | K09155 | 132 |
| DSAG12_01068 | hypothetical protein | -- | 49 |
| DSAG12_01069 | hypothetical protein | -- | 473 |
| DSAG12_01070 | ABC-2 type transport system ATP-binding protein | K01990 | 22 |
| DSAG12_01071 | hypothetical protein | -- | 271 |
| DSAG12_01072 | dihydrolipoamide dehydrogenase | K00382 | 160 |
| DSAG12_01073 | MarR family transcriptional regulator, transcriptional regulator for hemolysin | K06075 | 167 |
| DSAG12_01074 | putative ABC transport system ATP-binding protein | K02003 | 142 |
| DSAG12_01075 | putative ABC transport system permease protein | K02004 | 223 |
| DSAG12_01076 | acetolactate synthase I/II/III large subunit | K01652 | 244 |
| DSAG12_01077 | hypothetical protein | -- | 376 |
| DSAG12_01078 | hypothetical protein | -- | 200 |
| DSAG12_01079 | hypothetical protein | -- | 180 |
| DSAG12_01080 | ferritin | K02217 | 209 |
| DSAG12_01081 | tRNA 2-thiouridine synthesizing protein A | K04085 | 138 |
| DSAG12_01082 | | K07092 | 244 |
| DSAG12_01083 | cysteine desulfurase | K04487 | 170 |
| DSAG12_01084 | hypothetical protein | -- | 225 |
| DSAG12_01085 | hypothetical protein | -- | 258 |

| | | | |
|---|---|---|---|
| DSAG12_01086 | cold shock protein (beta-ribbon, CspA family) | K03704 | 321 |
| DSAG12_01087 | hypothetical protein | -- | 271 |
| DSAG12_01088 | hypothetical protein | -- | 267 |
| DSAG12_01089 | hypothetical protein | -- | 527 |
| DSAG12_01090 | hypothetical protein | -- | 470 |
| DSAG12_01091 | hypothetical protein | -- | 108 |
| DSAG12_01092 | hypothetical protein | -- | 191 |
| DSAG12_01093 | hypothetical protein | -- | 323 |
| DSAG12_01094 | | K00680 | 230 |
| DSAG12_01095 | hypothetical protein | -- | 322 |
| DSAG12_01096 | | K00519 | 154 |
| DSAG12_01097 | hypothetical protein | -- | 150 |
| DSAG12_01098 | hypothetical protein | -- | 154 |
| DSAG12_01099 | | K06883 | 155 |
| DSAG12_01100 | 2-oxoglutarate ferredoxin oxidoreductase subunit beta | K00175 | 107 |
| DSAG12_01101 | 2-oxoglutarate ferredoxin oxidoreductase subunit alpha | K00174 | 145 |
| DSAG12_01102 | rubredoxin-NAD+ reductase | K05297 | 96 |
| DSAG12_01103 | hypothetical protein | -- | 0 |
| DSAG12_01104 | hypothetical protein | -- | 117 |
| DSAG12_01105 | hypothetical protein | -- | 166 |
| DSAG12_01106 | hypothetical protein | -- | 222 |
| DSAG12_01107 | | K06888 | 325 |
| DSAG12_01108 | hypothetical protein | -- | 220 |
| DSAG12_01109 | hypothetical protein | -- | 133 |
| DSAG12_01110 | | K07066 | 179 |
| DSAG12_01111 | acyl-CoA dehydrogenase | K00249 | 263 |
| DSAG12_01112 | heterodisulfide reductase subunit D | K08264 | 275 |
| DSAG12_01113 | glycolate oxidase | K00104 | 24 |
| DSAG12_01114 | hypothetical protein | -- | 201 |
| DSAG12_01115 | hypothetical protein | -- | 127 |
| DSAG12_01116 | hypothetical protein | -- | 268 |
| DSAG12_01117 | hypothetical protein | -- | 208 |
| DSAG12_01118 | hypothetical protein | -- | 176 |
| DSAG12_01119 | glutaredoxin-like protein NrdH | K06191 | 132 |
| DSAG12_01120 | nitrite reductase (NADH) large subunit | K00362 | 111 |
| DSAG12_01121 | hypothetical protein | -- | 307 |
| DSAG12_01122 | hypothetical protein | -- | 235 |
| DSAG12_01123 | 4Fe-4S ferredoxin | K00205 | 252 |
| DSAG12_01124 | F420-non-reducing hydrogenase iron-sulfur subunit | K14127 | 183 |
| DSAG12_01125 | heterodisulfide reductase subunit A | K03388 | 263 |
| DSAG12_01126 | heterodisulfide reductase subunit B | K03389 | 180 |
| DSAG12_01127 | heterodisulfide reductase subunit C | K03390 | 282 |
| DSAG12_01128 | tRNA 2-thiouridine synthesizing protein A | K04085 | 141 |
| DSAG12_01129 | tRNA 2-thiouridine synthesizing protein D | K07235 | 90 |
| DSAG12_01130 | tRNA 2-thiouridine synthesizing protein C | K07236 | 198 |
| DSAG12_01131 | hypothetical protein | -- | 157 |
| DSAG12_01132 | hypothetical protein | -- | 252 |
| DSAG12_01133 | Lrp/AsnC family transcriptional regulator, regulator for asnA, asnC and gidA | K03718 | 189 |
| DSAG12_01134 | heterodisulfide reductase subunit C | K03390 | 190 |
| DSAG12_01135 | heterodisulfide reductase subunit B | K03389 | 267 |
| DSAG12_01136 | heterodisulfide reductase subunit A | K03388 | 210 |
| DSAG12_01137 | heterodisulfide reductase subunit A | K03388 | 245 |
| DSAG12_01138 | hypothetical protein | -- | 97 |
| DSAG12_01139 | hypothetical protein | -- | 155 |
| DSAG12_01140 | tRNA 2-thiouridine synthesizing protein A | K04085 | 150 |
| DSAG12_01141 | hypothetical protein | -- | 105 |
| DSAG12_01142 | glucose-1-phosphate thymidylyltransferase | K00973 | 264 |
| DSAG12_01143 | hypothetical protein | -- | 132 |
| DSAG12_01144 | hypothetical protein | -- | 218 |
| DSAG12_01145 | hypothetical protein | -- | 248 |
| DSAG12_01146 | 3-isopropylmalate dehydrogenase | K00052 | 281 |
| DSAG12_01147 | hypothetical protein | -- | 172 |
| DSAG12_01148 | hypothetical protein | -- | 222 |
| DSAG12_01149 | heterodisulfide reductase subunit B | K03389 | 275 |
| DSAG12_01150 | heterodisulfide reductase subunit C | K03390 | 65 |
| DSAG12_01151 | Ras-related protein Rab-8A | K07901 | 196 |
| DSAG12_01152 | hypothetical protein | -- | 171 |
| DSAG12_01153 | hypothetical protein | -- | 168 |
| DSAG12_01154 | hypothetical protein | -- | 230 |
| DSAG12_01155 | hypothetical protein | -- | 70 |
| DSAG12_01156 | superkiller protein 3 | K12600 | 223 |
| DSAG12_01157 | large subunit ribosomal protein L15 | K02876 | 972 |
| DSAG12_01158 | large subunit ribosomal protein L30 | K02907 | 212 |
| DSAG12_01159 | small subunit ribosomal protein S5 | K02988 | 0 |
| DSAG12_01160 | large subunit ribosomal protein L18 | K02881 | 239 |
| DSAG12_01161 | large subunit ribosomal protein L19e | K02885 | 241 |
| DSAG12_01162 | large subunit ribosomal protein L32e | K02912 | 152 |
| DSAG12_01163 | large subunit ribosomal protein L6 | K02933 | 224 |

| | | | | |
|---|---|---|---|---|
| DSAG12_01164 | small subunit ribosomal protein S8 | K02994 | 203 | |
| DSAG12_01165 | large subunit ribosomal protein L5 | K02931 | 198 | |
| DSAG12_01166 | small subunit ribosomal protein S4e | K02987 | 294 | Putative oligosaccharyl transferase STT3 subunit |
| DSAG12_01167 | large subunit ribosomal protein L24 | K02895 | 220 | |
| DSAG12_01168 | large subunit ribosomal protein L14 | K02874 | 163 | |
| DSAG12_01169 | small subunit ribosomal protein S17 | K02961 | 189 | |
| DSAG12_01170 | hypothetical protein | -- | 192 | |
| DSAG12_01171 | large subunit ribosomal protein L29 | K02904 | 203 | |
| DSAG12_01172 | small subunit ribosomal protein S3 | K02982 | 165 | |
| DSAG12_01173 | large subunit ribosomal protein L22 | K02890 | 282 | |
| DSAG12_01174 | hypothetical protein | -- | 202 | |
| DSAG12_01175 | small subunit ribosomal protein S19 | K02965 | 194 | |
| DSAG12_01176 | hypothetical protein | -- | 0 | |
| DSAG12_01177 | hypothetical protein | -- | 159 | |
| DSAG12_01178 | acyl-CoA dehydrogenase | K00249 | 218 | |
| DSAG12_01179 | hypothetical protein | -- | 142 | |
| DSAG12_01180 | hypothetical protein | -- | 149 | |
| DSAG12_01181 | | K07142 | 193 | |
| DSAG12_01182 | hypothetical protein | -- | 231 | |
| DSAG12_01183 | hypothetical protein | -- | 41 | |
| DSAG12_01184 | DNA repair protein RadA | K04483 | 250 | |
| DSAG12_01185 | hypothetical protein | -- | 204 | |
| DSAG12_01186 | 23S rRNA (uridine2552-2'-O)-methyltransferase | K02427 | 21 | |
| DSAG12_01187 | tRNA (guanine26-N2/guanine27-N2)-dimethyltransferase | K00555 | 195 | |
| DSAG12_01188 | DNA repair protein RadB | K04484 | 156 | |
| DSAG12_01189 | hypothetical protein | -- | 182 | |
| DSAG12_01190 | hypothetical protein | -- | 280 | |
| DSAG12_01191 | hypothetical protein | -- | 201 | |
| DSAG12_01192 | hypothetical protein | -- | 125 | |
| DSAG12_01193 | tritrans,polycis-undecaprenyl-diphosphate synthase [geranylgeranyl-diphosphate specific] | K15888 | 170 | |
| DSAG12_01194 | geranylgeranyl reductase | K10960 | 219 | |
| DSAG12_01195 | geranylgeranyl diphosphate synthase, type I | K13787 | 181 | Putative oligosaccharyltransferase complex subunit |
| DSAG12_01196 | isopentenyl-diphosphate delta-isomerase | K01823 | 168 | |
| DSAG12_01197 | isopentenyl phosphate kinase | K06981 | 132 | |
| DSAG12_01198 | mevalonate kinase | K00869 | 186 | |
| DSAG12_01199 | hydroxymethylglutaryl-CoA reductase | K00054 | 188 | |
| DSAG12_01200 | diphosphomevalonate decarboxylase | K01597 | 143 | |
| DSAG12_01201 | hydroxymethylglutaryl-CoA synthase | K01641 | 133 | |
| DSAG12_01202 | perosamine synthetase | K13010 | 214 | |
| DSAG12_01203 | hypothetical protein | -- | 153 | |
| DSAG12_01204 | hypothetical protein | -- | 207 | |
| DSAG12_01205 | 3-hydroxybutyryl-CoA dehydratase | K17865 | 145 | |
| DSAG12_01206 | | K00680 | 190 | |
| DSAG12_01207 | hypothetical protein | K09726 | 173 | |
| DSAG12_01208 | 4'-phosphopantetheinyl transferase | K06133 | 195 | |
| DSAG12_01209 | proline iminopeptidase | K01259 | 103 | |
| DSAG12_01210 | hypothetical protein | -- | 161 | |
| DSAG12_01211 | hypothetical protein | -- | 198 | |
| DSAG12_01212 | acetaldehyde dehydrogenase / alcohol dehydrogenase | K04072 | 191 | |
| DSAG12_01213 | long-chain acyl-CoA synthetase | K01897 | 232 | |
| DSAG12_01214 | acyl carrier protein | K02078 | 228 | |
| DSAG12_01215 | UDP-N-acetylglucosamine--dolichyl-phosphate N-acetylglucosaminephosphotransferase | K01001 | 104 | |
| DSAG12_01216 | hypothetical protein | -- | 0 | |
| DSAG12_01217 | 3-oxoadipate enol-lactonase | K01055 | 0 | |
| DSAG12_01218 | endonuclease III related protein | K07457 | 0 | |
| DSAG12_01219 | hypothetical protein | -- | 139 | |
| DSAG12_01220 | Ras-related protein Rab-11A | K07904 | 141 | |
| DSAG12_01221 | | K07158 | 226 | |
| DSAG12_01222 | hypothetical protein | -- | 220 | |
| DSAG12_01223 | hypothetical protein | -- | 93 | |
| DSAG12_01224 | hypothetical protein | -- | 138 | |
| DSAG12_01225 | hypothetical protein | -- | 173 | |
| DSAG12_01226 | hypothetical protein | -- | 221 | |
| DSAG12_01227 | hypothetical protein | -- | 185 | |
| DSAG12_01228 | hypothetical protein | -- | 193 | |
| DSAG12_01229 | hypothetical protein | -- | 57 | |
| DSAG12_01230 | protein phosphatase 1 regulatory subunit 7 | K17550 | 219 | |
| DSAG12_01231 | translation initiation factor 1A | K03236 | 212 | |
| DSAG12_01232 | hypothetical protein | -- | 197 | |
| DSAG12_01233 | hypothetical protein | -- | 171 | |
| DSAG12_01234 | triphosphoribosyl-dephospho-CoA synthase | K05966 | 330 | |
| DSAG12_01235 | hypothetical protein | -- | 127 | |
| DSAG12_01236 | ribulose-bisphosphate carboxylase large chain | K01601 | 305 | |
| DSAG12_01237 | ribose 1,5-bisphosphate isomerase | K18237 | 61 | |
| DSAG12_01238 | 5-(carboxyamino)imidazole ribonucleotide mutase | K01588 | 150 | |
| DSAG12_01239 | hypothetical protein | -- | 195 | |

| | | | | |
|---|---|---|---|---|
| DSAG12_01240 | adenylyltransferase and sulfurtransferase | K11996 | 311 | |
| DSAG12_01241 | hypothetical protein | -- | 215 | |
| DSAG12_01242 | hypothetical protein | -- | 141 | |
| DSAG12_01243 | beta-glucuronidase | K01195 | 258 | |
| DSAG12_01244 | hypothetical protein | -- | 237 | |
| DSAG12_01245 | hypothetical protein | -- | 185 | |
| DSAG12_01246 | hypothetical protein | -- | 187 | |
| DSAG12_01247 | hypothetical protein | -- | 104 | |
| DSAG12_01248 | 3-dehydroquinate dehydratase I | K03785 | 174 | |
| DSAG12_01249 | shikimate dehydrogenase | K00014 | 57 | putative homolog of eukaryotic ribosomal protein L22e |
| DSAG12_01250 | | K07041 | 258 | |
| DSAG12_01251 | ferredoxin-type protein NapH | K02574 | 241 | |
| DSAG12_01252 | hypothetical protein | -- | 179 | |
| DSAG12_01253 | hypothetical protein | -- | 210 | |
| DSAG12_01254 | hypothetical protein | K09121 | 210 | |
| DSAG12_01255 | hypothetical protein | -- | 356 | |
| DSAG12_01256 | ribosomal RNA methyltransferase Nop2 | K14835 | 84 | |
| DSAG12_01257 | 60S ribosome subunit biogenesis protein NIP7 | K07565 | 151 | |
| DSAG12_01258 | hypothetical protein | -- | 220 | |
| DSAG12_01259 | hypothetical protein | -- | 235 | |
| DSAG12_01260 | ABC-2 type transport system permease protein | K01992 | 195 | |
| DSAG12_01261 | ABC-2 type transport system ATP-binding protein | K01990 | 177 | |
| DSAG12_01262 | hypothetical protein | -- | 172 | |
| DSAG12_01263 | tRNA-Gln | | 186 | |
| DSAG12_01264 | adenylate cyclase, class 2 | K05873 | 272 | |
| DSAG12_01265 | uncharacterized protein | K06950 | 208 | |
| DSAG12_01266 | | K00680 | 203 | |
| DSAG12_01267 | hypothetical protein | -- | 165 | |
| DSAG12_01268 | hypothetical protein | -- | 175 | |
| DSAG12_01269 | hypothetical protein | -- | 99 | |
| DSAG12_01270 | hypothetical protein | -- | 202 | |
| DSAG12_01271 | hypothetical protein | -- | 188 | |
| DSAG12_01272 | hypothetical protein | -- | 114 | |
| DSAG12_01273 | FKBP-type peptidyl-prolyl cis-trans isomerase SlyD | K03775 | 184 | |
| DSAG12_01274 | | K07024 | 175 | |
| DSAG12_01275 | hypothetical protein | -- | 200 | |
| DSAG12_01276 | tRNA wybutosine-synthesizing protein 2 | K07055 | 198 | |
| DSAG12_01277 | hypothetical protein | -- | 0 | |
| DSAG12_01278 | hypothetical protein | -- | 140 | |
| DSAG12_01279 | L-aspartate oxidase | K00278 | 183 | |
| DSAG12_01280 | hypothetical protein | -- | 225 | |
| DSAG12_01281 | 3-hydroxypropionyl-coenzyme A dehydratase | K15019 | 177 | |
| DSAG12_01282 | butyryl-CoA dehydrogenase | K00248 | 188 | |
| DSAG12_01283 | glycoside/pentoside/hexuronide:cation symporter, GPH family | K03292 | 195 | |
| DSAG12_01284 | hypothetical protein | -- | 103 | |
| DSAG12_01285 | protein transport protein SEC24 | K14007 | 227 | |
| DSAG12_01286 | thioredoxin reductase (NADPH) | K00384 | 134 | |
| DSAG12_01287 | 23S rRNA (adenine2503-C2)-methyltransferase | K06941 | 235 | |
| DSAG12_01288 | long-chain acyl-CoA synthetase | K01897 | 143 | |
| DSAG12_01289 | serine protease AprX | K17734 | 196 | |
| DSAG12_01290 | hypothetical protein | -- | 184 | |
| DSAG12_01291 | NADH-quinone oxidoreductase subunit C/D | K13378 | 96 | |
| DSAG12_01292 | hypothetical protein | -- | 207 | |
| DSAG12_01293 | hypothetical protein | -- | 243 | |
| DSAG12_01294 | dUTP pyrophosphatase | K01520 | 155 | |
| DSAG12_01295 | hypothetical protein | -- | 245 | |
| DSAG12_01296 | cysteine desulfurase | K04487 | 212 | |
| DSAG12_01297 | arginyl-tRNA synthetase | K01887 | 231 | |
| DSAG12_01298 | ribose 5-phosphate isomerase A | K01807 | 159 | |
| DSAG12_01299 | hypothetical protein | -- | 190 | |
| DSAG12_01300 | hypothetical protein | -- | 159 | |
| DSAG12_01301 | prolyl-tRNA synthetase | K01881 | 174 | |
| DSAG12_01302 | hypothetical protein | -- | 250 | |
| DSAG12_01303 | hypothetical protein | -- | 174 | |
| DSAG12_01304 | mycothiol synthase | K15520 | 125 | |
| DSAG12_01305 | hypothetical protein | -- | 149 | |
| DSAG12_01306 | adenosylhomocysteinase | K01251 | 175 | |
| DSAG12_01307 | hypothetical protein | -- | 204 | |
| DSAG12_01308 | hypothetical protein | -- | 168 | |
| DSAG12_01309 | | K00257 | 110 | |
| DSAG12_01310 | hypothetical protein | -- | 0 | |
| DSAG12_01311 | hypothetical protein | -- | 224 | |
| DSAG12_01312 | hypothetical protein | -- | 160 | |
| DSAG12_01313 | proteasome beta subunit | K03433 | 233 | |
| DSAG12_01314 | hypothetical protein | K09730 | 54 | |
| DSAG12_01315 | hypothetical protein | -- | 164 | |
| DSAG12_01316 | hypothetical protein | -- | 232 | |

| | | | |
|---|---|---|---|
| DSAG12_01317 | endonuclease III | K10773 | 262 |
| DSAG12_01318 | two-component system, NtrC family, response regulator AtoC | K07714 | 170 |
| DSAG12_01319 | two-component system, NtrC family, sensor kinase | K02482 | 227 |
| DSAG12_01320 | protein-tyrosine phosphatase | K01104 | 187 |
| DSAG12_01321 | histidinol-phosphatase (PHP family) | K04486 | 262 |
| DSAG12_01322 | hypothetical protein | -- | 164 |
| DSAG12_01323 | hydroxyacylglutathione hydrolase | K01069 | 201 |
| DSAG12_01324 | hypothetical protein | -- | 257 |
| DSAG12_01325 | hypothetical protein | -- | 273 |
| DSAG12_01326 | | K06944 | 155 |
| DSAG12_01327 | hypothetical protein | -- | 150 |
| DSAG12_01328 | drug/metabolite transporter, DME family | K03298 | 222 |
| DSAG12_01329 | hypothetical protein | -- | 102 |
| DSAG12_01330 | | K00540 | 126 |
| DSAG12_01331 | GTP-binding protein | K03979 | 201 |
| DSAG12_01332 | glycoside/pentoside/hexuronide:cation symporter, GPH family | K03292 | 180 |
| DSAG12_01333 | hypothetical protein | -- | 138 |
| DSAG12_01334 | hypothetical protein | -- | 154 |
| DSAG12_01335 | glycoside/pentoside/hexuronide:cation symporter, GPH family | K03292 | 146 |
| DSAG12_01336 | hypothetical protein | -- | 159 |
| DSAG12_01337 | hypothetical protein | -- | 140 |
| DSAG12_01338 | NitT/TauT family transport system ATP-binding protein | K02049 | 163 |
| DSAG12_01339 | NitT/TauT family transport system permease protein | K02050 | 139 |
| DSAG12_01340 | hypothetical protein | -- | 159 |
| DSAG12_01341 | NitT/TauT family transport system substrate-binding protein | K02051 | 78 |
| DSAG12_01342 | hypothetical protein | -- | 254 |
| DSAG12_01343 | hypothetical protein | -- | 150 |
| DSAG12_01344 | hypothetical protein | -- | 187 |
| DSAG12_01345 | tRNA-Thr | Intron(14498 | 166 |
| DSAG12_01346 | hypothetical protein | -- | 243 |
| DSAG12_01347 | hypothetical protein | -- | 55 |
| DSAG12_01348 | hypothetical protein | -- | 68 |
| DSAG12_01349 | hypothetical protein | -- | 0 |
| DSAG12_01350 | protein-S-isoprenylcysteine O-methyltransferase | K00587 | 270 |
| DSAG12_01351 | hypothetical protein | -- | 176 |
| DSAG12_01352 | endoglucanase | K01179 | 36 |
| DSAG12_01353 | tRNA-Met | Intron(14569 | 162 |
| DSAG12_01354 | hypothetical protein | -- | 210 |
| DSAG12_01355 | hypothetical protein | -- | 154 |
| DSAG12_01356 | hypothetical protein | -- | 229 |
| DSAG12_01357 | hypothetical protein | -- | 190 |
| DSAG12_01358 | hypothetical protein | -- | 211 |
| DSAG12_01359 | hypothetical protein | -- | 198 |
| DSAG12_01360 | hypothetical protein | -- | 111 |
| DSAG12_01361 | MFS transporter, DHA3 family, macrolide efflux protein | K08217 | 260 |
| DSAG12_01362 | hypothetical protein | -- | 170 |
| DSAG12_01363 | | K00540 | 199 |
| DSAG12_01364 | D-alanine-D-alanine ligase | K01921 | 163 |
| DSAG12_01365 | Ras-related protein Rab-4B | K07880 | 622 |
| DSAG12_01366 | hypothetical protein | -- | 283 |
| DSAG12_01367 | methylthioribose-1-phosphate isomerase | K08963 | 131 |
| DSAG12_01368 | translation initiation factor IF-2 | K02519 | 75 |
| DSAG12_01369 | | K07005 | 189 |
| DSAG12_01370 | hypothetical protein | -- | 344 |
| DSAG12_01371 | 7, 8-dihydropterin-6-yl-methyl-4-(beta-D- ribofuranosyl)aminobenzene 5'-phosphate synthase | K06897 | 393 |
| DSAG12_01372 | hypothetical protein | -- | 198 |
| DSAG12_01373 | hypothetical protein | -- | 125 |
| DSAG12_01374 | hypothetical protein | K09133 | 243 |
| DSAG12_01375 | hypothetical protein | -- | 107 |
| DSAG12_01376 | hypothetical protein | -- | 187 |
| DSAG12_01377 | trans-2,3-dihydro-3-hydroxyanthranilate isomerase | K06998 | 167 |
| DSAG12_01378 | hypothetical protein | -- | 353 |
| DSAG12_01379 | hypothetical protein | -- | 198 |
| DSAG12_01380 | hypothetical protein | -- | 242 |
| DSAG12_01381 | putative acetyltransferase | K03826 | 255 |
| DSAG12_01382 | hypothetical protein | -- | 158 |
| DSAG12_01383 | hypothetical protein | -- | 202 |
| DSAG12_01384 | hypothetical protein | -- | 182 |
| DSAG12_01385 | hypothetical protein | -- | 188 |
| DSAG12_01386 | hypothetical protein | -- | 172 |
| DSAG12_01387 | hypothetical protein | -- | 208 |
| DSAG12_01388 | hypothetical protein | -- | 195 |
| DSAG12_01389 | tRNA-Asp | Intron(14894 | 188 |
| DSAG12_01390 | phosphoglycolate phosphatase | K01091 | 137 |
| DSAG12_01391 | hypothetical protein | -- | 186 |
| DSAG12_01392 | hypothetical protein | -- | 118 |
| DSAG12_01393 | hypothetical protein | -- | 169 |
| DSAG12_01394 | hypothetical protein | -- | 186 |

| | | | | |
|---|---|---|---|---|
| DSAG12_01395 | hypothetical protein | -- | 197 | |
| DSAG12_01396 | aminotransferase | K10907 | 196 | |
| DSAG12_01397 | hypothetical protein | -- | 161 | |
| DSAG12_01398 | homoserine dehydrogenase | K00003 | 78 | |
| DSAG12_01399 | transposase | K07486 | 233 | |
| DSAG12_01400 | hypothetical protein | -- | 0 | |
| DSAG12_01401 | exodeoxyribonuclease III | K01142 | 142 | |
| DSAG12_01402 | aldehyde:ferredoxin oxidoreductase | K03738 | 153 | |
| DSAG12_01403 | two-component system, cell cycle sensor histidine kinase and response regulator CckA | K13587 | 188 | |
| DSAG12_01404 | hypothetical protein | -- | 130 | |
| DSAG12_01405 | retinol dehydrogenase 13 | K11161 | 198 | |
| DSAG12_01406 | hypothetical protein | -- | 191 | |
| DSAG12_01407 | hypothetical protein | -- | 256 | |
| DSAG12_01408 | hydroxyacylglutathione hydrolase | K01069 | 135 | |
| DSAG12_01409 | Ras-related protein Rab-11A | K07904 | 266 | |
| DSAG12_01410 | hypothetical protein | -- | 192 | |
| DSAG12_01411 | hypothetical protein | -- | 171 | |
| DSAG12_01412 | inorganic pyrophosphatase | K01507 | 175 | |
| DSAG12_01413 | hypothetical protein | -- | 277 | |
| DSAG12_01414 | cystathione beta-lyase | K14155 | 193 | |
| DSAG12_01415 | hypothetical protein | -- | 251 | |
| DSAG12_01416 | hypothetical protein | -- | 191 | |
| DSAG12_01417 | hypothetical protein | -- | 0 | |
| DSAG12_01418 | | K00540 | 239 | |
| DSAG12_01419 | geranylgeranyl diphosphate synthase, type I | K13787 | 560 | |
| DSAG12_01420 | hypothetical protein | -- | 215 | |
| DSAG12_01421 | hypothetical protein | -- | 162 | small GTP-binding domain protein |
| DSAG12_01422 | selenide, water dikinase | K01008 | 191 | |
| DSAG12_01423 | hypothetical protein | -- | 169 | |
| DSAG12_01424 | translation initiation factor 2 subunit 3 | K03242 | 187 | |
| DSAG12_01425 | acetylornithine aminotransferase | K00818 | 99 | |
| DSAG12_01426 | hypothetical protein | -- | 216 | |
| DSAG12_01427 | | K00680 | 176 | |
| DSAG12_01428 | arsenate reductase | K03741 | 133 | |
| DSAG12_01429 | hypothetical protein | -- | 154 | |
| DSAG12_01430 | hypothetical protein | -- | 163 | |
| DSAG12_01431 | TatD DNase family protein | K03424 | 188 | |
| DSAG12_01432 | hypothetical protein | -- | 130 | |
| DSAG12_01433 | hypothetical protein | -- | 150 | |
| DSAG12_01434 | hypothetical protein | -- | 261 | |
| DSAG12_01435 | hypothetical protein | -- | 165 | |
| DSAG12_01436 | hypothetical protein | -- | 220 | |
| DSAG12_01437 | hypothetical protein | -- | 226 | |
| DSAG12_01438 | hypothetical protein | -- | 216 | |
| DSAG12_01439 | hypothetical protein | -- | 155 | |
| DSAG12_01440 | hypothetical protein | -- | 182 | |
| DSAG12_01441 | hypothetical protein | -- | 102 | |
| DSAG12_01442 | hypothetical protein | -- | 152 | |
| DSAG12_01443 | hypothetical protein | -- | 212 | |
| DSAG12_01444 | hypothetical protein | -- | 7 | |
| DSAG12_01445 | integrase/recombinase XerD | K04763 | 1 | |
| DSAG12_01446 | hypothetical protein | -- | 5 | |
| DSAG12_01447 | hypothetical protein | -- | 0 | |
| DSAG12_01448 | lysophospholipase | K01048 | 0 | |
| DSAG12_01449 | hypothetical protein | -- | 1 | |
| DSAG12_01450 | hypothetical protein | -- | 0 | |
| DSAG12_01451 | hypothetical protein | -- | 4 | |
| DSAG12_01452 | GMP synthase (glutamine-hydrolysing) | K01951 | 6 | |
| DSAG12_01453 | | K00680 | 1 | |
| DSAG12_01454 | hypothetical protein | -- | 7 | |
| DSAG12_01455 | hypothetical protein | -- | 6 | |
| DSAG12_01456 | hypothetical protein | -- | 1 | |
| DSAG12_01457 | tRNA-Val | | 6 | |
| DSAG12_01458 | DNA-directed RNA polymerase subunit K | K03055 | 7 | |
| DSAG12_01459 | hypothetical protein | -- | 147 | |
| DSAG12_01460 | hypothetical protein | -- | 159 | |
| DSAG12_01461 | fanconi anemia group M protein | K10896 | 225 | |
| DSAG12_01462 | histidinol-phosphatase (PHP family) | K04486 | 146 | |
| DSAG12_01463 | hypothetical protein | -- | 168 | |
| DSAG12_01464 | actin beta/gamma 1 | K05692 | 253 | |
| DSAG12_01465 | hypothetical protein | -- | 257 | |
| DSAG12_01466 | hypothetical protein | -- | 0 | |
| DSAG12_01467 | hypothetical protein | -- | 148 | |
| DSAG12_01468 | hypothetical protein | -- | 163 | |
| DSAG12_01469 | 4-aminobutyrate aminotransferase | K00823 | 138 | |
| DSAG12_01470 | hypothetical protein | -- | 149 | |
| DSAG12_01471 | hypothetical protein | -- | 0 | |
| DSAG12_01472 | alanyl-tRNA synthetase | K01872 | 131 | |

| | | | | |
|---|---|---|---|---|
| DSAG12_01473 | hypothetical protein | -- | 82 | |
| DSAG12_01474 | hypothetical protein | -- | 107 | |
| DSAG12_01475 | hypothetical protein | -- | 208 | |
| DSAG12_01476 | hypothetical protein | -- | 139 | |
| DSAG12_01477 | hypothetical protein | -- | 136 | |
| DSAG12_01478 | hypothetical protein | -- | 155 | |
| DSAG12_01479 | ATP-binding cassette, subfamily B, bacterial | K06147 | 159 | |
| DSAG12_01480 | hypothetical protein | -- | 169 | |
| DSAG12_01481 | hypothetical protein | -- | 221 | |
| DSAG12_01482 | Lrp/AsnC family transcriptional regulator, regulator for asnA, asnC and gidA | K03718 | 45 | |
| DSAG12_01483 | Lrp/AsnC family transcriptional regulator, regulator for asnA, asnC and gidA | K03718 | 244 | |
| DSAG12_01484 | hypothetical protein | -- | 241 | |
| DSAG12_01485 | hypothetical protein | -- | 189 | |
| DSAG12_01486 | hypothetical protein | -- | 257 | |
| DSAG12_01487 | glycogen(starch) synthase | K16150 | 123 | |
| DSAG12_01488 | mannose-1-phosphate guanylyltransferase / phosphomannomutase | K16881 | 163 | |
| DSAG12_01489 | mannose-1-phosphate guanylyltransferase | K00966 | 219 | |
| DSAG12_01490 | Ras-related protein Rab-21 | K07890 | 178 | |
| DSAG12_01491 | hypothetical protein | -- | 200 | |
| DSAG12_01492 | hypothetical protein | -- | 222 | |
| DSAG12_01493 | large subunit ribosomal protein L38e | K02923 | 0 | |
| DSAG12_01494 | peptidyl-tRNA hydrolase, PTH2 family | K04794 | 228 | |
| DSAG12_01495 | CopG family transcriptional regulator, nickel-responsive regulator | K07722 | 219 | |
| DSAG12_01496 | nicotinate-nucleotide pyrophosphorylase (carboxylating) | K00767 | 183 | |
| DSAG12_01497 | | K03423 | 38 | |
| DSAG12_01498 | hypothetical protein | K09141 | 34 | |
| DSAG12_01499 | dihydroorotase | K01465 | 317 | |
| DSAG12_01500 | signal recognition particle subunit SRP54 | K03106 | 64 | |
| DSAG12_01501 | hypothetical protein | -- | 127 | |
| DSAG12_01502 | hypothetical protein | -- | 292 | |
| DSAG12_01503 | putative transposase | K07491 | 199 | |
| DSAG12_01504 | putative transposase | K07491 | 218 | |
| DSAG12_01505 | beta-phosphoglucomutase | K01838 | 205 | small GTP-binding domain protein |
| DSAG12_01506 | hypothetical protein | -- | 115 | |
| DSAG12_01507 | translation initiation factor 5A | K03263 | 285 | |
| DSAG12_01508 | NAD+ kinase | K00858 | 50 | |
| DSAG12_01509 | UPF0271 protein | K07060 | 168 | |
| DSAG12_01510 | flap endonuclease-1 | K04799 | 157 | |
| DSAG12_01511 | transitional endoplasmic reticulum ATPase | K13525 | 221 | |
| DSAG12_01512 | hypothetical protein | -- | 222 | |
| DSAG12_01513 | | K06915 | 249 | |
| DSAG12_01514 | hypothetical protein | -- | 199 | |
| DSAG12_01515 | hypothetical protein | -- | 279 | |
| DSAG12_01516 | hypothetical protein | -- | 191 | |
| DSAG12_01517 | GMP synthase (glutamine-hydrolysing) | K01951 | 0 | |
| DSAG12_01518 | hypothetical protein | -- | 113 | |
| DSAG12_01519 | hypothetical protein | -- | 120 | |
| DSAG12_01520 | hypothetical protein | -- | 91 | |
| DSAG12_01521 | hypothetical protein | -- | 166 | |
| DSAG12_01522 | hypothetical protein | -- | 220 | |
| DSAG12_01523 | Ras-related protein Rab-3D | K07884 | 196 | |
| DSAG12_01524 | hypothetical protein | -- | 216 | |
| DSAG12_01525 | | K07095 | 173 | |
| DSAG12_01526 | hypothetical protein | -- | 224 | |
| DSAG12_01527 | phosphoribosylformylglycinamidine cyclo-ligase | K01933 | 306 | |
| DSAG12_01528 | adenylosuccinate lyase | K01756 | 88 | |
| DSAG12_01529 | hypothetical protein | -- | 195 | |
| DSAG12_01530 | hypothetical protein | -- | 170 | |
| DSAG12_01531 | TldD protein | K03568 | 189 | |
| DSAG12_01532 | PmbA protein | K03592 | 186 | |
| DSAG12_01533 | hypothetical protein | -- | 154 | |
| DSAG12_01534 | hypothetical protein | -- | 267 | |
| DSAG12_01535 | UDP-glucuronate decarboxylase | K08678 | 466 | |
| DSAG12_01536 | hypothetical protein | -- | 194 | |
| DSAG12_01537 | hypothetical protein | -- | 155 | |
| DSAG12_01538 | hypothetical protein | -- | 148 | |
| DSAG12_01539 | hypothetical protein | -- | 147 | |
| DSAG12_01540 | hypothetical protein | -- | 147 | |
| DSAG12_01541 | hypothetical protein | -- | 173 | |
| DSAG12_01542 | hypothetical protein | -- | 151 | |
| DSAG12_01543 | hypothetical protein | -- | 219 | |
| DSAG12_01544 | hypothetical protein | -- | 244 | |
| DSAG12_01545 | hypothetical protein | -- | 114 | |
| DSAG12_01546 | ATP-binding cassette, subfamily B, bacterial | K06147 | 120 | |
| DSAG12_01547 | ATP-binding cassette, subfamily B, bacterial | K06147 | 234 | |
| DSAG12_01548 | hypothetical protein | -- | 182 | |
| DSAG12_01549 | hypothetical protein | -- | 229 | |
| DSAG12_01550 | replication factor C small subunit | K04801 | 143 | |

| | | | |
|---|---|---|---|
| DSAG12_01551 | replication factor C small subunit | K04801 | 230 |
| DSAG12_01552 | replication factor C large subunit | K04800 | 110 |
| DSAG12_01553 | hypothetical protein | -- | 102 |
| DSAG12_01554 | putative adenine-specific DNA-methyltransferase | K07319 | 217 |
| DSAG12_01555 | putative adenine-specific DNA-methyltransferase | K07319 | 139 |
| DSAG12_01556 | MFS transporter, DHA1 family, multidrug resistance protein | K08161 | 202 |
| DSAG12_01557 | hypothetical protein | -- | 143 |
| DSAG12_01558 | hypothetical protein | -- | 170 |
| DSAG12_01559 | hypothetical protein | -- | 192 |
| DSAG12_01560 | transcription initiation factor TFIIB | K03124 | 203 |
| DSAG12_01561 | hypothetical protein | -- | 191 |
| DSAG12_01562 | protein-tyrosine phosphatase | K01104 | 283 |
| DSAG12_01563 | Lrp/AsnC family transcriptional regulator, regulator for asnA, asnC and gidA | K03718 | 308 |
| DSAG12_01564 | hypothetical protein | -- | 113 |
| DSAG12_01565 | peptide/nickel transport system permease protein | K02033 | 115 |
| DSAG12_01566 | peptide/nickel transport system permease protein | K02034 | 178 |
| DSAG12_01567 | hypothetical protein | -- | 161 |
| DSAG12_01568 | ribosomal-protein-alanine N-acetyltransferase | K03790 | 201 |
| DSAG12_01569 | hypothetical protein | -- | 109 |
| DSAG12_01570 | glycine hydroxymethyltransferase | K00600 | 94 |
| DSAG12_01571 | | K06889 | 157 |
| DSAG12_01572 | hypothetical protein | -- | 201 |
| DSAG12_01573 | hypothetical protein | -- | 109 |
| DSAG12_01574 | DNA-directed RNA polymerase subunit D | K03047 | 161 |
| DSAG12_01575 | hypothetical protein | -- | 180 |
| DSAG12_01576 | TetR/AcrR family transcriptional regulator, multidrug resistance operon repressor | K18136 | 0 |
| DSAG12_01577 | hypothetical protein | -- | 243 |
| DSAG12_01578 | small subunit ribosomal protein S9 | K02996 | 49 |
| DSAG12_01579 | large subunit ribosomal protein L13 | K02871 | 150 |
| DSAG12_01580 | large subunit ribosomal protein L18e | K02883 | 202 |
| DSAG12_01581 | hypothetical protein | -- | 230 |
| DSAG12_01582 | hypothetical protein | -- | 193 |
| DSAG12_01583 | hypothetical protein | -- | 223 |
| DSAG12_01584 | hypothetical protein | -- | 320 |
| DSAG12_01585 | hypothetical protein | -- | 178 |
| DSAG12_01586 | hypothetical protein | -- | 178 |
| DSAG12_01587 | 3-phosphoshikimate 1-carboxyvinyltransferase | K00800 | 286 |
| DSAG12_01588 | hypothetical protein | -- | 278 |
| DSAG12_01589 | hypothetical protein | -- | 171 |
| DSAG12_01590 | hypothetical protein | -- | 199 |
| DSAG12_01591 | hypothetical protein | -- | 203 |
| DSAG12_01592 | hypothetical protein | -- | 209 |
| DSAG12_01593 | hypothetical protein | -- | 146 |
| DSAG12_01594 | hypothetical protein | -- | 191 |
| DSAG12_01595 | hypothetical protein | -- | 196 |
| DSAG12_01596 | Lrp/AsnC family transcriptional regulator, leucine-responsive regulatory protein | K03719 | 200 |
| DSAG12_01597 | hypothetical protein | -- | 158 |
| DSAG12_01598 | hypothetical protein | -- | 146 |
| DSAG12_01599 | | K06997 | 158 |
| DSAG12_01600 | hypothetical protein | -- | 90 |
| DSAG12_01601 | phosphoenolpyruvate phosphomutase | K01841 | 291 |
| DSAG12_01602 | 1,3-propanediol dehydrogenase | K00086 | 292 |
| DSAG12_01603 | phosphonopyruvate decarboxylase | K09459 | 485 |
| DSAG12_01604 | phosphoenolpyruvate phosphomutase | K01841 | 232 |
| DSAG12_01605 | serine racemase | K12235 | 185 |
| DSAG12_01606 | hypothetical protein | -- | 247 |
| DSAG12_01607 | hypothetical protein | -- | 122 |
| DSAG12_01608 | hypothetical protein | -- | 169 |
| DSAG12_01609 | hypothetical protein | -- | 72 |
| DSAG12_01610 | | K07090 | 243 |
| DSAG12_01611 | | K07050 | 41 |
| DSAG12_01612 | hypothetical protein | -- | 241 |
| DSAG12_01613 | hypothetical protein | -- | 97 |
| DSAG12_01614 | glycerol kinase | K00864 | 202 |
| DSAG12_01615 | hypothetical protein | -- | 138 |
| DSAG12_01616 | hypothetical protein | -- | 179 |
| DSAG12_01617 | hypothetical protein | -- | 175 |
| DSAG12_01618 | hypothetical protein | -- | 161 |
| DSAG12_01619 | hypothetical protein | -- | 206 |
| DSAG12_01620 | hypothetical protein | -- | 218 |
| DSAG12_01621 | hypothetical protein | -- | 217 |
| DSAG12_01622 | hypothetical protein | -- | 205 |
| DSAG12_01623 | DtxR family transcriptional regulator, Mn-dependent transcriptional regulator | K03709 | 200 |
| DSAG12_01624 | ferrous iron transport protein B | K04759 | 299 |
| DSAG12_01625 | hypothetical protein | -- | 24 |
| DSAG12_01626 | hypothetical protein | K09165 | 161 |
| DSAG12_01627 | carboxylesterase | K03928 | 202 |
| DSAG12_01628 | hypothetical protein | -- | 207 |

| | | | | |
|---|---|---|---|---|
| DSAG12_01629 | beta-glucosidase | K05349 | 185 | |
| DSAG12_01630 | hypothetical protein | -- | 127 | |
| DSAG12_01631 | cobalt/nickel transport system ATP-binding protein / cobalt/nickel transport system permease protein | K02006 K02 | 255 | |
| DSAG12_01632 | cobalt/nickel transport system permease protein | K02008 | 178 | |
| DSAG12_01633 | hypothetical protein | -- | 105 | |
| DSAG12_01634 | cobalt/nickel transport system permease protein | K02007 | 130 | |
| DSAG12_01635 | hypothetical protein | -- | 170 | |
| DSAG12_01636 | hypothetical protein | -- | 131 | |
| DSAG12_01637 | hypothetical protein | -- | 153 | |
| DSAG12_01638 | hypothetical protein | -- | 133 | |
| DSAG12_01639 | molecular chaperone GrpE | K03687 | 199 | |
| DSAG12_01640 | hypothetical protein | -- | 114 | |
| DSAG12_01641 | phosphoenolpyruvate carboxykinase (GTP) | K01596 | 214 | |
| DSAG12_01642 | putative MFS transporter, AGZA family, xanthine/uracil permease | K06901 | 83 | |
| DSAG12_01643 | hypothetical protein | -- | 33 | |
| DSAG12_01644 | hypothetical protein | -- | 184 | |
| DSAG12_01645 | hypothetical protein | -- | 344 | |
| DSAG12_01646 | hypothetical protein | -- | 213 | |
| DSAG12_01647 | hypothetical protein | -- | 142 | |
| DSAG12_01648 | hypothetical protein | -- | 136 | |
| DSAG12_01649 | hypothetical protein | -- | 128 | |
| DSAG12_01650 | hypothetical protein | -- | 109 | |
| DSAG12_01651 | hypothetical protein | -- | 241 | |
| DSAG12_01652 | transcription initiation factor TFIIB | K03124 | 167 | |
| DSAG12_01653 | gelsolin | K05768 | 168 | |
| DSAG12_01654 | hypothetical protein | -- | 188 | |
| DSAG12_01655 | hypothetical protein | -- | 73 | |
| DSAG12_01656 | hypothetical protein | -- | 197 | |
| DSAG12_01657 | hypothetical protein | -- | 154 | small GTP-binding domain protein |
| DSAG12_01658 | hypothetical protein | -- | 0 | |
| DSAG12_01659 | transitional endoplasmic reticulum ATPase | K13525 | 252 | |
| DSAG12_01660 | HSP20 family protein | K13993 | 261 | |
| DSAG12_01661 | HSP20 family protein | K13993 | 267 | |
| DSAG12_01662 | MFS transporter, DHA3 family, macrolide efflux protein | K08217 | 192 | |
| DSAG12_01663 | | K07052 | 376 | |
| DSAG12_01664 | hypothetical protein | -- | 245 | |
| DSAG12_01665 | fructokinase | K00847 | 211 | |
| DSAG12_01666 | | K06990 | 256 | |
| DSAG12_01667 | | K06948 | 178 | |
| DSAG12_01668 | hypothetical protein | -- | 180 | |
| DSAG12_01669 | hypothetical protein | -- | 223 | |
| DSAG12_01670 | hypothetical protein | -- | 257 | |
| DSAG12_01671 | carbon-nitrogen hydrolase family protein | K08590 | 340 | hypothetical proteins with gelsolin-like domain |
| DSAG12_01672 | tRNA-intron endonuclease, archaea type | K01170 | 250 | |
| DSAG12_01673 | hypothetical protein | -- | 27 | |
| DSAG12_01674 | V/A-type H+-transporting ATPase subunit I | K02123 | 173 | |
| DSAG12_01675 | V/A-type H+-transporting ATPase subunit D | K02120 | 106 | |
| DSAG12_01676 | V/A-type H+-transporting ATPase subunit B | K02118 | 262 | |
| DSAG12_01677 | V/A-type H+-transporting ATPase subunit A | K02117 | 419 | |
| DSAG12_01678 | hypothetical protein | -- | 399 | |
| DSAG12_01679 | V/A-type H+-transporting ATPase subunit F | K02122 | 194 | |
| DSAG12_01680 | hypothetical protein | -- | 237 | |
| DSAG12_01681 | V/A-type H+-transporting ATPase subunit C | K02119 | 251 | |
| DSAG12_01682 | V/A-type H+-transporting ATPase subunit K | K02124 | 171 | |
| DSAG12_01683 | hypothetical protein | K07220 | 191 | |
| DSAG12_01684 | hypothetical protein | -- | 183 | |
| DSAG12_01685 | hypothetical protein | -- | 210 | |
| DSAG12_01686 | UDPglucose 6-dehydrogenase | K00012 | 125 | |
| DSAG12_01687 | PUA domain protein | K07575 | 206 | |
| DSAG12_01688 | NADH dehydrogenase / NADH dehydrogenase | K00329 K0C | 195 | |
| DSAG12_01689 | hypothetical protein | -- | 171 | |
| DSAG12_01690 | U6 snRNA-associated Sm-like protein LSm5 | K12624 | 151 | |
| DSAG12_01691 | large subunit ribosomal protein L37e | K02922 | 127 | |
| DSAG12_01692 | hypothetical protein | -- | 183 | |
| DSAG12_01693 | hypothetical protein | -- | 138 | |
| DSAG12_01694 | glycoside/pentoside/hexuronide:cation symporter, GPH family | K03292 | 218 | |
| DSAG12_01695 | hypothetical protein | -- | 223 | |
| DSAG12_01696 | | K07053 | 108 | |
| DSAG12_01697 | NAD-dependent deacetylase sirtuin 7 | K11417 | 47 | |
| DSAG12_01698 | glycoside/pentoside/hexuronide:cation symporter, GPH family | K03292 | 128 | |
| DSAG12_01699 | hypothetical protein | -- | 148 | |
| DSAG12_01700 | hypothetical protein | -- | 253 | |
| DSAG12_01701 | dolichol-phosphate mannosyltransferase | K00721 | 308 | |
| DSAG12_01702 | hypothetical protein | -- | 226 | |
| DSAG12_01703 | hypothetical protein | -- | 145 | |
| DSAG12_01704 | hypothetical protein | -- | 223 | |

| | | | |
|---|---|---|---|
| DSAG12_01705 | all-trans-retinol 13,14-reductase | K09516 | 147 |
| DSAG12_01706 | glycoside/pentoside/hexuronide:cation symporter, GPH family | K03292 | 169 |
| DSAG12_01707 | all-trans-retinol 13,14-reductase | K09516 | 137 |
| DSAG12_01708 | hypothetical protein | -- | 80 |
| DSAG12_01709 | hypothetical protein | -- | 125 |
| DSAG12_01710 | putative ABC transport system permease protein | K02004 | 161 |
| DSAG12_01711 | hypothetical protein | -- | 202 |
| DSAG12_01712 | hypothetical protein | -- | 266 |
| DSAG12_01713 | putative ABC transport system permease protein | K02004 | 216 |
| DSAG12_01714 | putative ABC transport system ATP-binding protein | K02003 | 201 |
| DSAG12_01715 | hypothetical protein | -- | 165 |
| DSAG12_01716 | hypothetical protein | -- | 274 |
| DSAG12_01717 | malate dehydrogenase (oxaloacetate-decarboxylating) | K00027 | 205 |
| DSAG12_01718 | glycerol-3-phosphate dehydrogenase | K00111 | 162 |
| DSAG12_01719 | sarcosine oxidase, subunit alpha | K00302 | 191 |
| DSAG12_01720 | hypothetical protein | -- | 172 |
| DSAG12_01721 | amidohydrolase | K01436 | 203 |
| DSAG12_01722 | polar amino acid transport system substrate-binding protein | K02030 | 242 |
| DSAG12_01723 | hypothetical protein | -- | 241 |
| DSAG12_01724 | hypothetical protein | -- | 253 |
| DSAG12_01725 | hypothetical protein | -- | 236 |
| DSAG12_01726 | 3-oxoacyl-[acyl-carrier protein] reductase | K00059 | 95 |
| DSAG12_01727 | glycerophosphoryl diester phosphodiesterase | K01126 | 145 |
| DSAG12_01728 | hypothetical protein | -- | 231 |
| DSAG12_01729 | transcription initiation factor TFIID TATA-box-binding protein | K03120 | 205 |
| DSAG12_01730 | hypothetical protein | -- | 170 |
| DSAG12_01731 | hypothetical protein | -- | 198 |
| DSAG12_01732 | hypothetical protein | -- | 231 |
| DSAG12_01733 | hypothetical protein | -- | 179 |
| DSAG12_01734 | transcription initiation factor TFIID TATA-box-binding protein | K03120 | 131 |
| DSAG12_01735 | Ras-related protein Rab-6A | K07893 | 249 |
| DSAG12_01736 | hypothetical protein | -- | 248 |
| DSAG12_01737 | acetyltransferase | K09181 | 238 |
| DSAG12_01738 | L-2-hydroxycarboxylate dehydrogenase (NAD+) | K05884 | 154 |
| DSAG12_01739 | hypothetical protein | -- | 205 |
| DSAG12_01740 | hypothetical protein | -- | 198 |
| DSAG12_01741 | hypothetical protein | -- | 129 |
| DSAG12_01742 | hypothetical protein | -- | 217 |
| DSAG12_01743 | beta-galactosidase | K01190 | 176 |
| DSAG12_01744 | ADP-dependent NAD(P)H-hydrate dehydratase / NAD(P)H-hydrate epimerase | K17758 K17 | 312 |
| DSAG12_01745 | hypothetical protein | -- | 202 |
| DSAG12_01746 | hypothetical protein | -- | 0 |
| DSAG12_01747 | hypothetical protein | -- | 84 |
| DSAG12_01748 | hypothetical protein | -- | 216 |
| DSAG12_01749 | hypothetical protein | -- | 165 |
| DSAG12_01750 | hypothetical protein | -- | 161 |
| DSAG12_01751 | hypothetical protein | -- | 125 |
| DSAG12_01752 | hypothetical protein | -- | 168 |
| DSAG12_01753 | hypothetical protein | -- | 134 |
| DSAG12_01754 | hypothetical protein | -- | 122 |
| DSAG12_01755 | hypothetical protein | -- | 176 |
| DSAG12_01756 | serpin B11/12 | K13966 | 250 |
| DSAG12_01757 | serpin B11/12 | K13966 | 270 |
| DSAG12_01758 | serpin B | K13963 | 182 |
| DSAG12_01759 | hypothetical protein | -- | 274 |
| DSAG12_01760 | methionyl aminopeptidase | K01265 | 211 |
| DSAG12_01761 | hypothetical protein | -- | 235 |
| DSAG12_01762 | hypothetical protein | -- | 231 |
| DSAG12_01763 | tRNA-Arg | | 232 |
| DSAG12_01764 | isopentenyl-diphosphate delta-isomerase | K01823 | 143 |
| DSAG12_01765 | hypothetical protein | -- | 288 |
| DSAG12_01766 | large subunit ribosomal protein L11 | K02867 | 150 |
| DSAG12_01767 | large subunit ribosomal protein L1 | K02863 | 105 |
| DSAG12_01768 | large subunit ribosomal protein L10 | K02864 | 181 |
| DSAG12_01769 | large subunit ribosomal protein L12 | K02869 | 237 |
| DSAG12_01770 | hypothetical protein | -- | 228 |
| DSAG12_01771 | hypothetical protein | -- | 157 |
| DSAG12_01772 | hypothetical protein | -- | 160 |
| DSAG12_01773 | hypothetical protein | K09717 | 168 |
| DSAG12_01774 | hypothetical protein | -- | 53 |
| DSAG12_01775 | hypothetical protein | -- | 164 |
| DSAG12_01776 | GTP cyclohydrolase I | K01495 | 190 |
| DSAG12_01777 | oleandomycin transport system permease protein | K18233 | 153 |
| DSAG12_01778 | ABC-2 type transport system ATP-binding protein | K01990 | 233 |
| DSAG12_01779 | hypothetical protein | -- | 149 |
| DSAG12_01780 | / putative redox protein | K06889 K07 | 131 |
| DSAG12_01781 | hypothetical protein | -- | 0 |
| DSAG12_01782 | dihydromethanopterin reductase (acceptor) | K18853 | 0 |

| | | | | |
|---|---|---|---|---|
| DSAG12_01783 | hypothetical protein | -- | 195 | |
| DSAG12_01784 | hypothetical protein | K09154 | 176 | |
| DSAG12_01785 | hypothetical protein | K09739 | 213 | |
| DSAG12_01786 | beta-ribofuranosylaminobenzene 5'-phosphate synthase | K06984 | 175 | |
| DSAG12_01787 | hydrogenase expression/formation protein | K07388 | 253 | |
| DSAG12_01788 | protein-tyrosine phosphatase | K01104 | 163 | |
| DSAG12_01789 | small subunit ribosomal protein S2 | K02967 | 213 | |
| DSAG12_01790 | hypothetical protein | -- | 134 | |
| DSAG12_01791 | hypothetical protein | K09955 | 37 | |
| DSAG12_01792 | hypothetical protein | -- | 183 | |
| DSAG12_01793 | small subunit ribosomal protein S8e | K02995 | 163 | |
| DSAG12_01794 | hypothetical protein | -- | 58 | |
| DSAG12_01795 | hypothetical protein | K09736 | 200 | |
| DSAG12_01796 | ribose-phosphate pyrophosphokinase | K00948 | 188 | |
| DSAG12_01797 | hypothetical protein | -- | 193 | |
| DSAG12_01798 | Ca2+-transporting ATPase | K01537 | 0 | |
| DSAG12_01799 | FAD synthetase | K14656 | 190 | |
| DSAG12_01800 | CDP-paratose 2-epimerase | K12454 | 152 | |
| DSAG12_01801 | translin | K07477 | 163 | |
| DSAG12_01802 | ribulose-phosphate 3-epimerase | K01783 | 200 | |
| DSAG12_01803 | Ras-related protein Rab-21 | K07890 | 111 | |
| DSAG12_01804 | hypothetical protein | -- | 126 | |
| DSAG12_01805 | hypothetical protein | -- | 149 | |
| DSAG12_01806 | hypothetical protein | -- | 206 | |
| DSAG12_01807 | hypothetical protein | -- | 139 | |
| DSAG12_01808 | hypothetical protein | -- | 284 | |
| DSAG12_01809 | archaetidylinositol phosphate synthase | K17884 | 165 | |
| DSAG12_01810 | hypothetical protein | -- | 186 | |
| DSAG12_01811 | phosphomannomutase | K17497 | 90 | |
| DSAG12_01812 | phosphatidylinositol alpha-mannosyltransferase | K08256 | 174 | |
| DSAG12_01813 | mannose-1-phosphate guanylyltransferase | K00971 | 147 | |
| DSAG12_01814 | dTDP-4-dehydrorhamnose reductase | K00067 | 122 | |
| DSAG12_01815 | glucose-1-phosphate thymidylyltransferase | K00973 | 222 | |
| DSAG12_01816 | dTDP-4-dehydrorhamnose reductase | K00067 | 169 | |
| DSAG12_01817 | dTDP-glucose 4,6-dehydratase | K01710 | 208 | |
| DSAG12_01818 | GDPmannose 4,6-dehydratase | K01711 | 157 | |
| DSAG12_01819 | hypothetical protein | -- | 171 | |
| DSAG12_01820 | hypothetical protein | -- | 150 | |
| DSAG12_01821 | hypothetical protein | -- | 188 | |
| DSAG12_01822 | hypothetical protein | -- | 221 | small GTP-binding domain protein |
| DSAG12_01823 | | K00680 | 89 | |
| DSAG12_01824 | hypothetical protein | -- | 107 | |
| DSAG12_01825 | hypothetical protein | -- | 172 | |
| DSAG12_01826 | hypothetical protein | -- | 130 | |
| DSAG12_01827 | | K01043 | 42 | |
| DSAG12_01828 | hypothetical protein | -- | 120 | |
| DSAG12_01829 | dTDP-4-amino-4,6-dideoxygalactose transaminase | K02805 | 208 | |
| DSAG12_01830 | GDP-L-fucose synthase | K02377 | 214 | |
| DSAG12_01831 | hypothetical protein | -- | 210 | |
| DSAG12_01832 | hypothetical protein | -- | 203 | |
| DSAG12_01833 | CDP-6-deoxy-D-xylo-4-hexulose-3-dehydrase | K12452 | 187 | |
| DSAG12_01834 | hypothetical protein | -- | 199 | |
| DSAG12_01835 | | K00680 | 203 | |
| DSAG12_01836 | hypothetical protein | -- | 185 | |
| DSAG12_01837 | pseudaminic acid synthase | K15898 | 179 | |
| DSAG12_01838 | | K00837 | 0 | |
| DSAG12_01839 | | K00837 | 4 | |
| DSAG12_01840 | hypothetical protein | -- | 4 | |
| DSAG12_01841 | ribosomal-protein-alanine N-acetyltransferase | K03790 | 8 | |
| DSAG12_01842 | hypothetical protein | -- | 9 | |
| DSAG12_01843 | UDP-N-acetylglucosamine 4,6-dehydratase | K15894 | 4 | |
| DSAG12_01844 | hypothetical protein | -- | 2 | |
| DSAG12_01845 | glycosyltransferase | K13002 | 9 | |
| DSAG12_01846 | dolichol-phosphate mannosyltransferase | K00721 | 2 | |
| DSAG12_01847 | hypothetical protein | -- | 8 | |
| DSAG12_01848 | hypothetical protein | -- | 12 | |
| DSAG12_01849 | hypothetical protein | -- | 5 | |
| DSAG12_01850 | hypothetical protein | -- | 4 | |
| DSAG12_01851 | hypothetical protein | -- | 8 | |
| DSAG12_01852 | hypothetical protein | -- | 4 | |
| DSAG12_01853 | UDP-N-acetylglucosamine 2-epimerase (non-hydrolysing) | K01791 | 1 | |
| DSAG12_01854 | | K01795 | 0 | |
| DSAG12_01855 | dTDP-glucose 4,6-dehydratase | K01710 | 12 | |
| DSAG12_01856 | hypothetical protein | -- | 9 | |
| DSAG12_01857 | D-inositol-3-phosphate glycosyltransferase | K15521 | 6 | |
| DSAG12_01858 | hypothetical protein | -- | 8 | |
| DSAG12_01859 | hypothetical protein | -- | 2 | |
| DSAG12_01860 | hypothetical protein | -- | 6 | |

| | | | | |
|---|---|---|---|---|
| DSAG12_01861 | UDP-N-acetylglucosamine 4-epimerase | K02473 | 11 | |
| DSAG12_01862 | hypothetical protein | -- | 0 | |
| DSAG12_01863 | putative acetyltransferase | K03828 | 4 | |
| DSAG12_01864 | leucyl-tRNA synthetase | K01869 | 2 | |
| DSAG12_01865 | UDP-N-acetyl-2-amino-2-deoxyglucuronate dehydrogenase | K13020 | 13 | |
| DSAG12_01866 | hypothetical protein | -- | 4 | |
| DSAG12_01867 | hypothetical protein | -- | 6 | |
| DSAG12_01868 | hypothetical protein | -- | 8 | |
| DSAG12_01869 | CDP-glycerol glycerophosphotransferase | K09809 | 7 | |
| DSAG12_01870 | hypothetical protein | -- | 2 | |
| DSAG12_01871 | | K01726 | 7 | |
| DSAG12_01872 | phosphoenolpyruvate phosphomutase | K01841 | 0 | |
| DSAG12_01873 | MFS transporter, OPA family, glycerol-3-phosphate transporter | K02445 | 8 | |
| DSAG12_01874 | ADP-ribose pyrophosphatase | K01515 | 10 | |
| DSAG12_01875 | hypothetical protein | -- | 7 | |
| DSAG12_01876 | | K00754 | 5 | |
| DSAG12_01877 | riboflavin kinase, archaea type | K07732 | 7 | |
| DSAG12_01878 | hypothetical protein | -- | 7 | |
| DSAG12_01879 | hypothetical protein | -- | 8 | |
| DSAG12_01880 | hypothetical protein | -- | 5 | |
| DSAG12_01881 | putative ABC transport system ATP-binding protein | K02003 | 0 | |
| DSAG12_01882 | hypothetical protein | -- | 24 | |
| DSAG12_01883 | hypothetical protein | -- | 17 | |
| DSAG12_01884 | hypothetical protein | -- | 55 | |
| DSAG12_01885 | slit 3 | K06850 | 206 | |
| DSAG12_01886 | | K07047 | 226 | |
| DSAG12_01887 | hypothetical protein | -- | 112 | |
| DSAG12_01888 | hypothetical protein | -- | 6 | |
| DSAG12_01889 | hypothetical protein | -- | 5 | |
| DSAG12_01890 | 3-hydroxybutyryl-CoA dehydratase | K17865 | 6 | |
| DSAG12_01891 | hypothetical protein | -- | 12 | |
| DSAG12_01892 | malonyl-CoA O-methyltransferase | K02169 | 7 | |
| DSAG12_01893 | glyceraldehyde 3-phosphate dehydrogenase | K00134 | 0 | |
| DSAG12_01894 | 7, 8-dihydropterin-6-yl-methyl-4-(beta-D- ribofuranosyl)aminobenzene 5'-phosphate synthase | K06897 | 11 | |
| DSAG12_01895 | hypothetical protein | -- | 12 | |
| DSAG12_01896 | enoyl-CoA hydratase / 3-hydroxyacyl-CoA dehydrogenase | K15016 | 3 | |
| DSAG12_01897 | hypothetical protein | -- | 100 | |
| DSAG12_01898 | phosphoglycerate kinase | K00927 | 141 | |
| DSAG12_01899 | uncharacterized protein | K06959 | 169 | |
| DSAG12_01900 | hypothetical protein | -- | 246 | |
| DSAG12_01901 | hypothetical protein | -- | 196 | |
| DSAG12_01902 | hypothetical protein | -- | 116 | |
| DSAG12_01903 | tRNA-Val | | 305 | |
| DSAG12_01904 | hypothetical protein | -- | 170 | |
| DSAG12_01905 | hypothetical protein | -- | 146 | profilin-like domain protein |
| DSAG12_01906 | zeta-carotene isomerase | K15744 | 132 | |
| DSAG12_01907 | hypothetical protein | -- | 192 | |
| DSAG12_01908 | hypothetical protein | -- | 229 | |
| DSAG12_01909 | transposase | K07486 | 190 | |
| DSAG12_01910 | hypothetical protein | -- | 209 | |
| DSAG12_01911 | | K06898 | 216 | |
| DSAG12_01912 | site-specific DNA-methyltransferase (cytosine-N4-specific) | K00590 | 176 | |
| DSAG12_01913 | putative adenine-specific DNA-methyltransferase | K07319 | 97 | |
| DSAG12_01914 | | K00936 | 130 | |
| DSAG12_01915 | hypothetical protein | -- | 259 | |
| DSAG12_01916 | DNA polymerase IV (archaeal DinB-like DNA polymerase) | K04479 | 188 | |
| DSAG12_01917 | hypothetical protein | -- | 203 | |
| DSAG12_01918 | hypothetical protein | -- | 187 | |
| DSAG12_01919 | hypothetical protein | -- | 210 | |
| DSAG12_01920 | hypothetical protein | -- | 251 | |
| DSAG12_01921 | hypothetical protein | -- | 253 | |
| DSAG12_01922 | | K07062 | 65 | |
| DSAG12_01923 | hypothetical protein | -- | 173 | |
| DSAG12_01924 | hypothetical protein | -- | 186 | |
| DSAG12_01925 | hypothetical protein | -- | 0 | |
| DSAG12_01926 | hypothetical protein | -- | 54 | |
| DSAG12_01927 | hypothetical protein | -- | 193 | |
| DSAG12_01928 | phosphoglycolate phosphatase | K01091 | 109 | |
| DSAG12_01929 | acetoin utilization protein AcuC | K04768 | 229 | |
| DSAG12_01930 | hypothetical protein | -- | 171 | |
| DSAG12_01931 | hypothetical protein | -- | 0 | |
| DSAG12_01932 | hypothetical protein | -- | 127 | |
| DSAG12_01933 | hypothetical protein | -- | 134 | |
| DSAG12_01934 | hypothetical protein | -- | 214 | |
| DSAG12_01935 | hypothetical protein | -- | 173 | |
| DSAG12_01936 | Ras-related protein Rab-7A | K07897 | 199 | |
| DSAG12_01937 | hypothetical protein | -- | 221 | |
| DSAG12_01938 | hypothetical protein | -- | 0 | |

| | | | | |
|---|---|---|---|---|
| DSAG12_01939 | hypothetical protein | -- | 11 | |
| DSAG12_01940 | hypothetical protein | -- | 176 | |
| DSAG12_01941 | cell division protein FtsZ | K03531 | 204 | |
| DSAG12_01942 | hypothetical protein | -- | 146 | |
| DSAG12_01943 | | K06995 | 10 | |
| DSAG12_01944 | hypothetical protein | -- | 148 | |
| DSAG12_01945 | | K07065 | 161 | |
| DSAG12_01946 | hypothetical protein | -- | 147 | |
| DSAG12_01947 | hypothetical protein | -- | 97 | |
| DSAG12_01948 | hypothetical protein | -- | 0 | |
| DSAG12_01949 | NADPH2:quinone reductase | K00344 | 287 | |
| DSAG12_01950 | hypothetical protein | -- | 305 | |
| DSAG12_01951 | hypothetical protein | -- | 184 | |
| DSAG12_01952 | hypothetical protein | -- | 131 | |
| DSAG12_01953 | hypothetical protein | -- | 182 | |
| DSAG12_01954 | helicase | K03726 | 271 | |
| DSAG12_01955 | replicative DNA helicase Mcm | K10726 | 230 | |
| DSAG12_01956 | hypothetical protein | -- | 155 | |
| DSAG12_01957 | AAA family ATPase | K07392 | 152 | |
| DSAG12_01958 | hypothetical protein | -- | 216 | |
| DSAG12_01959 | hypothetical protein | -- | 190 | |
| DSAG12_01960 | 2-dehydropantoate 2-reductase | K00077 | 214 | |
| DSAG12_01961 | | K00936 | 312 | small GTP-binding domain protein |
| DSAG12_01962 | glycyl-tRNA synthetase | K01880 | 0 | |
| DSAG12_01963 | hypothetical protein | -- | 6 | |
| DSAG12_01964 | hypothetical protein | -- | 235 | |
| DSAG12_01965 | hypothetical protein | -- | 241 | |
| DSAG12_01966 | carboxypeptidase Taq | K01299 | 245 | |
| DSAG12_01967 | glycoside/pentoside/hexuronide:cation symporter, GPH family | K03292 | 202 | |
| DSAG12_01968 | hypothetical protein | -- | 97 | |
| DSAG12_01969 | hypothetical protein | -- | 51 | |
| DSAG12_01970 | hypothetical protein | -- | 89 | |
| DSAG12_01971 | hypothetical protein | -- | 0 | |
| DSAG12_01972 | hypothetical protein | -- | 195 | |
| DSAG12_01973 | glutamate dehydrogenase (NAD(P)+) | K00261 | 0 | |
| DSAG12_01974 | glutamate dehydrogenase (NADP+) | K00262 | 289 | |
| DSAG12_01975 | hypothetical protein | -- | 257 | |
| DSAG12_01976 | hypothetical protein | -- | 190 | |
| DSAG12_01977 | hypothetical protein | -- | 7 | |
| DSAG12_01978 | glutamate dehydrogenase (NADP+) | K00262 | 200 | |
| DSAG12_01979 | hypothetical protein | -- | 234 | |
| DSAG12_01980 | hypothetical protein | -- | 223 | |
| DSAG12_01981 | hypothetical protein | -- | 238 | |
| DSAG12_01982 | hypothetical protein | -- | 188 | |
| DSAG12_01983 | Ras-related protein Rab-34 | K07921 | 208 | |
| DSAG12_01984 | hypothetical protein | -- | 58 | |
| DSAG12_01985 | small subunit ribosomal protein S30e | K02983 | 76 | |
| DSAG12_01986 | tRNA-Gln | | 185 | |
| DSAG12_01987 | hypothetical protein | -- | 132 | |
| DSAG12_01988 | hypothetical protein | -- | 251 | |
| DSAG12_01989 | hypothetical protein | -- | 13 | |
| DSAG12_01990 | acetolactate synthase I/II/III large subunit | K01652 | 144 | |
| DSAG12_01991 | alkenylglycerophosphocholine/alkenylglycerophosphoethano lamine hydrolase | K18575 | 108 | |
| DSAG12_01992 | dihydroxy-acid dehydratase | K01687 | 266 | |
| DSAG12_01993 | hypothetical protein | -- | 230 | |
| DSAG12_01994 | glycoside/pentoside/hexuronide:cation symporter, GPH family | K03292 | 99 | |
| DSAG12_01995 | hypothetical protein | -- | 97 | |
| DSAG12_01996 | flavin-binding kelch repeat F-box protein 1 | K12116 | 154 | |
| DSAG12_01997 | hypothetical protein | -- | 186 | |
| DSAG12_01998 | hypothetical protein | -- | 241 | |
| DSAG12_01999 | hypothetical protein | -- | 1313 | |
| DSAG12_02000 | hypothetical protein | -- | 627 | |
| DSAG12_02001 | beta-phosphoglucomutase | K01838 | 232 | |
| DSAG12_02002 | hypothetical protein | -- | 129 | |
| DSAG12_02003 | hypothetical protein | -- | 94 | |
| DSAG12_02004 | serine/threonine-protein phosphatase PP1 catalytic subunit | K06269 | 295 | |
| DSAG12_02005 | hypothetical protein | -- | 233 | |
| DSAG12_02006 | hypothetical protein | -- | 213 | |
| DSAG12_02007 | UDP-glucose 4-epimerase | K01784 | 233 | putative membrane-bound protein with ubiquitin- like domain |
| DSAG12_02008 | hypothetical protein | -- | 116 | |
| DSAG12_02009 | methionyl aminopeptidase | K01265 | 202 | |
| DSAG12_02010 | | K07131 | 213 | |
| DSAG12_02011 | dTMP kinase | K00943 | 73 | |
| DSAG12_02012 | hypothetical protein | -- | 0 | |
| DSAG12_02013 | hypothetical protein | -- | 130 | |

| | | | | |
|---|---|---|---|---|
| DSAG12_02014 | hypothetical protein | -- | 229 | [illegible] (dolichyl-diphosphooligosaccharide--protein glycosyltransferase subunit 1) |
| DSAG12_02015 | hypothetical protein | -- | 58 | |
| DSAG12_02016 | MoxR-like ATPase | K03924 | 266 | |
| DSAG12_02017 | superkiller protein 3 | K12600 | 119 | |
| DSAG12_02018 | two-component system, cell cycle sensor histidine kinase and response regulator CckA | K13587 | 250 | |
| DSAG12_02019 | hypothetical protein | -- | 233 | |
| DSAG12_02020 | hypothetical protein | -- | 205 | |
| DSAG12_02021 | hypothetical protein | K07502 | 162 | |
| DSAG12_02022 | hypothetical protein | K09122 | 237 | |
| DSAG12_02023 | hypothetical protein | -- | 198 | |
| DSAG12_02024 | hypothetical protein | -- | 180 | |
| DSAG12_02025 | formylmethanofuran dehydrogenase subunit E | K11261 | 192 | |
| DSAG12_02026 | MFS transporter, DHA1 family, multidrug resistance protein | K08153 | 98 | |
| DSAG12_02027 | long-chain acyl-CoA synthetase | K01897 | 139 | |
| DSAG12_02028 | long-chain acyl-CoA synthetase | K01897 | 6 | |
| DSAG12_02029 | cysteine synthase A | K01738 | 91 | |
| DSAG12_02030 | serine O-acetyltransferase | K00640 | 147 | |
| DSAG12_02031 | hypothetical protein | -- | 144 | |
| DSAG12_02032 | hypothetical protein | K09711 | 201 | |
| DSAG12_02033 | lipoate-protein ligase A | K03800 | 237 | |
| DSAG12_02034 | biotin synthase | K01012 | 70 | |
| DSAG12_02035 | hypothetical protein | -- | 266 | |
| DSAG12_02036 | hypothetical protein | -- | 199 | |
| DSAG12_02037 | hypothetical protein | -- | 171 | |
| DSAG12_02038 | hypothetical protein | -- | 245 | |
| DSAG12_02039 | TldD protein | K03568 | 195 | |
| DSAG12_02040 | PmbA protein | K03592 | 156 | |
| DSAG12_02041 | Ras-related GTP-binding protein A/B | K16185 | 259 | |
| DSAG12_02042 | Ras-related GTP-binding protein A/B | K16185 | 299 | |
| DSAG12_02043 | hypothetical protein | -- | 216 | |
| DSAG12_02044 | thioredoxin 1 | K03671 | 0 | |
| DSAG12_02045 | hypothetical protein | -- | 261 | |
| DSAG12_02046 | superkiller protein 3 | K12600 | 270 | |
| DSAG12_02047 | hypothetical protein | -- | 171 | |
| DSAG12_02048 | hypothetical protein | -- | 152 | |
| DSAG12_02049 | triosephosphate isomerase (TIM) | K01803 | 171 | |
| DSAG12_02050 | hypothetical protein | -- | 182 | |
| DSAG12_02051 | hypothetical protein | -- | 160 | |
| DSAG12_02052 | energy-coupling factor transport system substrate-specific component | K16923 | 93 | |
| DSAG12_02053 | hypothetical protein | -- | 294 | |
| DSAG12_02054 | threonine synthase | K01733 | 251 | |
| DSAG12_02055 | drug/metabolite transporter, DME family | K03298 | 226 | |
| DSAG12_02056 | hypothetical protein | -- | 218 | |
| DSAG12_02057 | hypothetical protein | -- | 209 | |
| DSAG12_02058 | hypothetical protein | -- | 185 | |
| DSAG12_02059 | hypothetical protein | -- | 224 | |
| DSAG12_02060 | hypothetical protein | -- | 204 | |
| DSAG12_02061 | DNA excision repair protein ERCC-2 | K10844 | 138 | |
| DSAG12_02062 | hemolysin III | K11068 | 289 | |
| DSAG12_02063 | starch synthase | K00703 | 205 | |
| DSAG12_02064 | uncharacterized protein | K06869 | 192 | |
| DSAG12_02065 | large subunit ribosomal protein L35Ae | K02917 | 186 | |
| DSAG12_02066 | hypothetical protein | -- | 178 | |
| DSAG12_02067 | hypothetical protein | -- | 189 | |
| DSAG12_02068 | | K07062 | 218 | |
| DSAG12_02069 | hypothetical protein | -- | 130 | |
| DSAG12_02070 | hypothetical protein | -- | 300 | |
| DSAG12_02071 | hypothetical protein | -- | 0 | |
| DSAG12_02072 | hypothetical protein | -- | 186 | |
| DSAG12_02073 | hypothetical protein | -- | 176 | |
| DSAG12_02074 | charged multivesicular body protein 4 | K12194 | 154 | |
| DSAG12_02075 | hypothetical protein | -- | 213 | |
| DSAG12_02076 | hypothetical protein | -- | 106 | |
| DSAG12_02077 | | K07065 | 142 | |
| DSAG12_02078 | hypothetical protein | -- | 478 | |
| DSAG12_02079 | leucine-rich repeat-containing G protein-coupled receptor 6 | K08399 | 146 | |
| DSAG12_02080 | pyruvate kinase | K00873 | 207 | |
| DSAG12_02081 | hypothetical protein | -- | 179 | |
| DSAG12_02082 | hypothetical protein | -- | 192 | |
| DSAG12_02083 | hypothetical protein | -- | 148 | |
| DSAG12_02084 | putative ABC transport system ATP-binding protein | K02021 | 189 | |
| DSAG12_02085 | ATP-binding cassette, subfamily B, bacterial | K06147 | 22 | |
| DSAG12_02086 | hypothetical protein | -- | 76 | |
| DSAG12_02087 | hypothetical protein | -- | 210 | |
| DSAG12_02088 | hypothetical protein | -- | 239 | |
| DSAG12_02089 | hypothetical protein | -- | 237 | |

| | | | | |
|---|---|---|---|---|
| DSAG12_02090 | hypothetical protein | -- | 232 | |
| DSAG12_02091 | hypothetical protein | -- | 134 | |
| DSAG12_02092 | hypothetical protein | -- | 266 | |
| DSAG12_02093 | transposase | K07486 | 256 | |
| DSAG12_02094 | hypothetical protein | -- | 64 | |
| DSAG12_02095 | hypothetical protein | -- | 64 | |
| DSAG12_02096 | hypothetical protein | -- | 189 | |
| DSAG12_02097 | Ras-related protein Rab-1A | K07874 | 0 | |
| DSAG12_02098 | hypothetical protein | -- | 211 | |
| DSAG12_02099 | hypothetical protein | -- | 133 | |
| DSAG12_02100 | hypothetical protein | -- | 351 | III) |
| DSAG12_02101 | hypothetical protein | -- | 189 | |
| DSAG12_02102 | hypothetical protein | -- | 166 | |
| DSAG12_02103 | fanconi anemia group M protein | K10896 | 93 | |
| DSAG12_02104 | hypothetical protein | -- | 69 | |
| DSAG12_02105 | hypothetical protein | -- | 111 | |
| DSAG12_02106 | nitrous oxidase accessory protein | K07218 | 232 | |
| DSAG12_02107 | Ras-related protein Rab-11A | K07904 | 170 | |
| DSAG12_02108 | hypothetical protein | -- | 198 | |
| DSAG12_02109 | helicase | K03726 | 437 | |
| DSAG12_02110 | hypothetical protein | -- | 216 | |
| DSAG12_02111 | hypothetical protein | -- | 228 | |
| DSAG12_02112 | hypothetical protein | -- | 203 | |
| DSAG12_02113 | DNA repair protein RadA | K04483 | 186 | |
| DSAG12_02114 | hypothetical protein | -- | 124 | |
| DSAG12_02115 | hypothetical protein | -- | 225 | |
| DSAG12_02116 | hypothetical protein | -- | 164 | |
| DSAG12_02117 | hypothetical protein | -- | 58 | |
| DSAG12_02118 | flap endonuclease-1 | K04799 | 388 | |
| DSAG12_02119 | hypothetical protein | -- | 164 | |
| DSAG12_02120 | hypothetical protein | -- | 118 | |
| DSAG12_02121 | hypothetical protein | -- | 168 | |
| DSAG12_02122 | hypothetical protein | -- | 183 | small GTP-binding domain protein |
| DSAG12_02123 | hypothetical protein | -- | 254 | |
| DSAG12_02124 | | K06883 | 71 | |
| DSAG12_02125 | hypothetical protein | -- | 252 | |
| DSAG12_02126 | hypothetical protein | -- | 107 | |
| DSAG12_02127 | hypothetical protein | -- | 173 | |
| DSAG12_02128 | hypothetical protein | -- | 225 | |
| DSAG12_02129 | hypothetical protein | -- | 172 | |
| DSAG12_02130 | hypothetical protein | -- | 182 | |
| DSAG12_02131 | hypothetical protein | -- | 211 | |
| DSAG12_02132 | hypothetical protein | -- | 0 | |
| DSAG12_02133 | fanconi anemia group M protein | K10896 | 129 | |
| DSAG12_02134 | hypothetical protein | -- | 226 | |
| DSAG12_02135 | hypothetical protein | -- | 261 | |
| DSAG12_02136 | hypothetical protein | -- | 211 | |
| DSAG12_02137 | hypothetical protein | -- | 220 | |
| DSAG12_02138 | hypothetical protein | -- | 191 | |
| DSAG12_02139 | hypothetical protein | -- | 256 | |
| DSAG12_02140 | hypothetical protein | -- | 96 | |
| DSAG12_02141 | hypothetical protein | -- | 0 | |
| DSAG12_02142 | hypothetical protein | -- | 239 | |
| DSAG12_02143 | hypothetical protein | -- | 224 | |
| DSAG12_02144 | hypothetical protein | -- | 153 | |
| DSAG12_02145 | fanconi anemia group M protein | K10896 | 267 | |
| DSAG12_02146 | hypothetical protein | -- | 163 | |
| DSAG12_02147 | hypothetical protein | -- | 97 | |
| DSAG12_02148 | hypothetical protein | -- | 223 | |
| DSAG12_02149 | hypothetical protein | -- | 162 | |
| DSAG12_02150 | hypothetical protein | -- | 247 | |
| DSAG12_02151 | hypothetical protein | -- | 212 | |
| DSAG12_02152 | hypothetical protein | -- | 225 | |
| DSAG12_02153 | | K07131 | 235 | |
| DSAG12_02154 | hypothetical protein | -- | 231 | |
| DSAG12_02155 | aspartyl-tRNA synthetase | K01876 | 292 | |
| DSAG12_02156 | tRNA pseudouridine13 synthase | K06176 | 106 | |
| DSAG12_02157 | small subunit ribosomal protein S11 | K02948 | 248 | |
| DSAG12_02158 | small subunit ribosomal protein S4 | K02986 | 155 | |
| DSAG12_02159 | small subunit ribosomal protein S13 | K02952 | 222 | |
| DSAG12_02160 | 4-hydroxybutyryl-CoA synthetase (ADP-forming) | K18593 | 101 | |
| DSAG12_02161 | hypothetical protein | -- | 276 | |
| DSAG12_02162 | NADPH2:quinone reductase | K00344 | 248 | |
| DSAG12_02163 | hypothetical protein | -- | 165 | |
| DSAG12_02164 | hypothetical protein | -- | 202 | |
| DSAG12_02165 | hypothetical protein | -- | 117 | |
| DSAG12_02166 | hypothetical protein | -- | 150 | |
| DSAG12_02167 | hypothetical protein | -- | 223 | |

| | | | | |
|---|---|---|---|---|
| DSAG12_02168 | amidophosphoribosyltransferase | K00764 | 185 | |
| DSAG12_02169 | phosphoribosylformylglycinamidine synthase | K01952 | 240 | |
| DSAG12_02170 | phosphoribosylformylglycinamidine synthase | K01952 | 179 | |
| DSAG12_02171 | hypothetical protein | -- | 276 | |
| DSAG12_02172 | fusion protein PurCD | K13713 | 215 | |
| DSAG12_02173 | dCTP diphosphatase | K16904 | 245 | |
| DSAG12_02174 | 3-oxoadipate enol-lactonase | K01055 | 246 | |
| DSAG12_02175 | | K07131 | 278 | |
| DSAG12_02176 | | K01932 | 220 | |
| DSAG12_02177 | dihydropteroate synthase | K00796 | 268 | |
| DSAG12_02178 | DNA gyrase subunit B | K02470 | 213 | |
| DSAG12_02179 | DNA gyrase subunit A | K02469 | 246 | |
| DSAG12_02180 | hypothetical protein | -- | 87 | |
| DSAG12_02181 | hypothetical protein | -- | 222 | |
| DSAG12_02182 | hypothetical protein | -- | 172 | |
| DSAG12_02183 | hypothetical protein | -- | 274 | |
| DSAG12_02184 | hypothetical protein | -- | 188 | |
| DSAG12_02185 | hypothetical protein | -- | 213 | |
| DSAG12_02186 | hypothetical protein | -- | 228 | |
| DSAG12_02187 | | K06883 | 159 | |
| DSAG12_02188 | hypothetical protein | -- | 215 | |
| DSAG12_02189 | hypothetical protein | -- | 144 | |
| DSAG12_02190 | flotillin | K07192 | 157 | |
| DSAG12_02191 | hypothetical protein | -- | 217 | |
| DSAG12_02192 | hypothetical protein | -- | 111 | |
| DSAG12_02193 | hypothetical protein | -- | 191 | |
| DSAG12_02194 | hypothetical protein | -- | 230 | |
| DSAG12_02195 | hypothetical protein | -- | 193 | |
| DSAG12_02196 | hypothetical protein | -- | 221 | |
| DSAG12_02197 | hypothetical protein | -- | 173 | |
| DSAG12_02198 | 3-deoxy-7-phosphoheptulonate synthase | K03856 | 195 | |
| DSAG12_02199 | 3-dehydroquinate synthase | K01735 | 100 | |
| DSAG12_02200 | hypothetical protein | -- | 164 | |
| DSAG12_02201 | asparaginyl-tRNA synthetase | K01893 | 172 | |
| DSAG12_02202 | hypothetical protein | -- | 206 | |
| DSAG12_02203 | hypothetical protein | K09989 | 200 | |
| DSAG12_02204 | hypothetical protein | -- | 226 | |
| DSAG12_02205 | hypothetical protein | -- | 237 | |
| DSAG12_02206 | hypothetical protein | -- | 88 | |
| DSAG12_02207 | hypothetical protein | -- | 186 | |
| DSAG12_02208 | DNA ligase 1 | K10747 | 207 | |
| DSAG12_02209 | deoxyhypusine synthase | K00809 | 287 | |
| DSAG12_02210 | arginine decarboxylase | K02626 | 38 | |
| DSAG12_02211 | hypothetical protein | -- | 107 | |
| DSAG12_02212 | hypothetical protein | -- | 177 | |
| DSAG12_02213 | D-alanine-D-alanine ligase | K01921 | 234 | small GTP-binding domain protein |
| DSAG12_02214 | D-alanine-D-alanine ligase | K01921 | 102 | |
| DSAG12_02215 | hypothetical protein | -- | 57 | |
| DSAG12_02216 | AMP phosphorylase | K18931 | 308 | |
| DSAG12_02217 | hypothetical protein | -- | 147 | |
| DSAG12_02218 | hypothetical protein | -- | 235 | |
| DSAG12_02219 | hypothetical protein | -- | 94 | |
| DSAG12_02220 | hypothetical protein | -- | 176 | |
| DSAG12_02221 | hypothetical protein | -- | 133 | |
| DSAG12_02222 | hypothetical protein | -- | 148 | |
| DSAG12_02223 | hypothetical protein | -- | 114 | |
| DSAG12_02224 | hypothetical protein | -- | 214 | |
| DSAG12_02225 | F420-non-reducing hydrogenase iron-sulfur subunit | K14127 | 201 | |
| DSAG12_02226 | heterodisulfide reductase subunit A | K03388 | 137 | |
| DSAG12_02227 | heterodisulfide reductase subunit A | K03388 | 225 | |
| DSAG12_02228 | heterodisulfide reductase subunit A | K03388 | 194 | |
| DSAG12_02229 | hypothetical protein | -- | 184 | |
| DSAG12_02230 | hypothetical protein | -- | 185 | |
| DSAG12_02231 | hypothetical protein | -- | 90 | |
| DSAG12_02232 | phosphonate transport system ATP-binding protein | K02041 | 146 | |
| DSAG12_02233 | hemin transport system ATP-binding protein | K09814 | 237 | |
| DSAG12_02234 | hypothetical protein | -- | 197 | |
| DSAG12_02235 | hypothetical protein | -- | 184 | |
| DSAG12_02236 | hypothetical protein | -- | 151 | |
| DSAG12_02237 | hypothetical protein | -- | 181 | |
| DSAG12_02238 | hypothetical protein | -- | 57 | |
| DSAG12_02239 | hypothetical protein | -- | 226 | |
| DSAG12_02240 | hypothetical protein | -- | 177 | |
| DSAG12_02241 | hypothetical protein | -- | 50 | |
| DSAG12_02242 | hypothetical protein | -- | 0 | |
| DSAG12_02243 | hypothetical protein | -- | 231 | |
| DSAG12_02244 | hypothetical protein | -- | 192 | |
| DSAG12_02245 | hypothetical protein | -- | 218 | |

| | | | | |
|---|---|---|---|---|
| DSAG12_02246 | hypothetical protein | -- | 76 | |
| DSAG12_02247 | hypothetical protein | -- | 137 | |
| DSAG12_02248 | hypothetical protein | -- | 247 | |
| DSAG12_02249 | ribokinase | K00852 | 39 | |
| DSAG12_02250 | hypothetical protein | -- | 89 | |
| DSAG12_02251 | glycoside/pentoside/hexuronide:cation symporter, GPH family | K03292 | 115 | |
| DSAG12_02252 | glucosamine--fructose-6-phosphate aminotransferase (isomerizing) | K00820 | 349 | |
| DSAG12_02253 | hypothetical protein | -- | 183 | |
| DSAG12_02254 | hypothetical protein | -- | 5 | |
| DSAG12_02255 | hypothetical protein | -- | 215 | |
| DSAG12_02256 | hypothetical protein | -- | 100 | |
| DSAG12_02257 | isoleucyl-tRNA synthetase | K01870 | 102 | |
| DSAG12_02258 | hypothetical protein | -- | 87 | |
| DSAG12_02259 | hypothetical protein | -- | 105 | |
| DSAG12_02260 | hypothetical protein | -- | 134 | |
| DSAG12_02261 | hypothetical protein | -- | 80 | |
| DSAG12_02262 | hypothetical protein | -- | 101 | |
| DSAG12_02263 | hypothetical protein | -- | 162 | |
| DSAG12_02264 | hypothetical protein | -- | 120 | |
| DSAG12_02265 | phosphoribosylamine--glycine ligase | K01945 | 125 | |
| DSAG12_02266 | N-glycosylase/DNA lyase | K03660 | 94 | |
| DSAG12_02267 | hypothetical protein | -- | 125 | |
| DSAG12_02268 | hypothetical protein | -- | 121 | |
| DSAG12_02269 | hypothetical protein | -- | 0 | |
| DSAG12_02270 | hypothetical protein | -- | 139 | |
| DSAG12_02271 | phosphomannomutase / phosphoglucomutase | K15778 | 176 | |
| DSAG12_02272 | nitric oxide dioxygenase | K05916 | 146 | |
| DSAG12_02273 | hypothetical protein | -- | 150 | |
| DSAG12_02274 | | K06940 | 0 | |
| DSAG12_02275 | alkylated DNA repair protein alkB homolog 8 | K10770 | 8 | |
| DSAG12_02276 | hypothetical protein | -- | 184 | |
| DSAG12_02277 | hypothetical protein | -- | 203 | |
| DSAG12_02278 | hypothetical protein | -- | 224 | |
| DSAG12_02279 | hypothetical protein | -- | 274 | |
| DSAG12_02280 | hypothetical protein | -- | 174 | |
| DSAG12_02281 | hypothetical protein | -- | 110 | |
| DSAG12_02282 | hypothetical protein | -- | 88 | |
| DSAG12_02283 | hypothetical protein | -- | 185 | |
| DSAG12_02284 | hypothetical protein | -- | 215 | |
| DSAG12_02285 | hypothetical protein | -- | 245 | |
| DSAG12_02286 | hypothetical protein | -- | 244 | |
| DSAG12_02287 | glycerophosphoryl diester phosphodiesterase | K01126 | 130 | |
| DSAG12_02288 | | K06966 | 181 | |
| DSAG12_02289 | | K07123 | 152 | |
| DSAG12_02290 | hypothetical protein | -- | 118 | |
| DSAG12_02291 | hypothetical protein | -- | 187 | |
| DSAG12_02292 | hypothetical protein | -- | 165 | |
| DSAG12_02293 | hypothetical protein | -- | 221 | |
| DSAG12_02294 | endoglucanase | K01179 | 138 | |
| DSAG12_02295 | hypothetical protein | -- | 166 | |
| DSAG12_02296 | menaquinone-dependent protoporphyrinogen oxidase | K00230 | 192 | |
| DSAG12_02297 | ornithine--oxo-acid transaminase | K00819 | 90 | |
| DSAG12_02298 | carbamate kinase | K00926 | 102 | |
| DSAG12_02299 | ornithine carbamoyltransferase | K00611 | 245 | |
| DSAG12_02300 | hypothetical protein | -- | 245 | |
| DSAG12_02301 | 16S rRNA (cytosine967-C5)-methyltransferase | K03500 | 105 | |
| DSAG12_02302 | hypothetical protein | -- | 185 | |
| DSAG12_02303 | | K01463 | 172 | |
| DSAG12_02304 | hypothetical protein | -- | 154 | |
| DSAG12_02305 | hypothetical protein | -- | 126 | |
| DSAG12_02306 | hypothetical protein | -- | 143 | |
| DSAG12_02307 | hypothetical protein | -- | 195 | |
| DSAG12_02308 | acetyl-CoA C-acetyltransferase | K00626 | 157 | |
| DSAG12_02309 | hypothetical protein | -- | 138 | |
| DSAG12_02310 | hypothetical protein | -- | 307 | |
| DSAG12_02311 | hypothetical protein | -- | 486 | |
| DSAG12_02312 | 2,4-dienoyl-CoA reductase (NADPH2) | K00219 | 249 | |
| DSAG12_02313 | hypothetical protein | -- | 194 | |
| DSAG12_02314 | hypothetical protein | -- | 156 | |
| DSAG12_02315 | hypothetical protein | -- | 173 | |
| DSAG12_02316 | hypothetical protein | -- | 215 | |
| DSAG12_02317 | hypothetical protein | -- | 175 | |
| DSAG12_02318 | beta-glucuronidase | K01195 | 184 | small GTP-binding domain protein |
| DSAG12_02319 | hypothetical protein | -- | 249 | |
| DSAG12_02320 | hypothetical protein | -- | 159 | |
| DSAG12_02321 | HSP20 family protein | K13993 | 162 | |
| DSAG12_02322 | hypothetical protein | -- | 248 | |
| DSAG12_02323 | hypothetical protein | -- | 90 | |

| | | | | |
|---|---|---|---|---|
| DSAG12_02324 | | K07062 | 169 | |
| DSAG12_02325 | aldehyde:ferredoxin oxidoreductase | K03738 | 347 | |
| DSAG12_02326 | hypothetical protein | -- | 285 | |
| DSAG12_02327 | hypothetical protein | -- | 286 | |
| DSAG12_02328 | hypothetical protein | -- | 479 | |
| DSAG12_02329 | orotate phosphoribosyltransferase | K00762 | 164 | |
| DSAG12_02330 | signal peptidase, endoplasmic reticulum-type | K13280 | 152 | |
| DSAG12_02331 | signal peptidase, endoplasmic reticulum-type | K13280 | 213 | |
| DSAG12_02332 | carboxylesterase | K03928 | 194 | |
| DSAG12_02333 | modification methylase | K13581 | 255 | |
| DSAG12_02334 | hypothetical protein | -- | 110 | |
| DSAG12_02335 | NAD-dependent deacetylase | K12410 | 123 | |
| DSAG12_02336 | hypothetical protein | -- | 329 | |
| DSAG12_02337 | diaminopropionate ammonia-lyase | K01751 | 187 | |
| DSAG12_02338 | indolepyruvate ferredoxin oxidoreductase, beta subunit | K00180 | 208 | |
| DSAG12_02339 | indolepyruvate ferredoxin oxidoreductase, alpha subunit | K00179 | 147 | |
| DSAG12_02340 | putative membrane protein | K00389 | 215 | |
| DSAG12_02341 | D-glycerate 3-kinase | K15918 | 185 | |
| DSAG12_02342 | hypothetical protein | -- | 146 | |
| DSAG12_02343 | hypothetical protein | -- | 114 | |
| DSAG12_02344 | hypothetical protein | -- | 147 | |
| DSAG12_02345 | hypothetical protein | -- | 193 | small GTP-binding domain protein |
| DSAG12_02346 | hypothetical protein | -- | 164 | |
| DSAG12_02347 | hypothetical protein | -- | 235 | |
| DSAG12_02348 | hypothetical protein | -- | 132 | |
| DSAG12_02349 | internalin A | K13730 | 188 | |
| DSAG12_02350 | hypothetical protein | -- | 154 | |
| DSAG12_02351 | hypothetical protein | -- | 29 | |
| DSAG12_02352 | 16S rRNA (cytosine967-C5)-methyltransferase | K03500 | 118 | |
| DSAG12_02353 | hypothetical protein | -- | 224 | |
| DSAG12_02354 | hypothetical protein | -- | 176 | |
| DSAG12_02355 | inorganic phosphate transporter, PiT family | K03306 | 255 | |
| DSAG12_02356 | hypothetical protein | -- | 222 | |
| DSAG12_02357 | CBS domain-containing membrane protein | K07168 | 157 | |
| DSAG12_02358 | | K00540 | 218 | |
| DSAG12_02359 | hypothetical protein | -- | 188 | |
| DSAG12_02360 | fanconi anemia group M protein | K10896 | 181 | |
| DSAG12_02361 | hypothetical protein | -- | 186 | |
| DSAG12_02362 | hypothetical protein | -- | 170 | |
| DSAG12_02363 | hypothetical protein | -- | 192 | |
| DSAG12_02364 | NAD+ diphosphatase | K03426 | 163 | |
| DSAG12_02365 | hypothetical protein | -- | 203 | |
| DSAG12_02366 | hypothetical protein | -- | 141 | |
| DSAG12_02367 | sarcosine oxidase, subunit beta | K00303 | 202 | |
| DSAG12_02368 | hypothetical protein | -- | 121 | |
| DSAG12_02369 | hypothetical protein | -- | 162 | |
| DSAG12_02370 | hypothetical protein | -- | 148 | |
| DSAG12_02371 | formylmethanofuran--tetrahydromethanopterin N-formyltransferase | K00672 | 89 | |
| DSAG12_02372 | hypothetical protein | -- | 0 | |
| DSAG12_02373 | hypothetical protein | -- | 207 | |
| DSAG12_02374 | archaetidylinositol phosphate synthase | K17884 | 81 | |
| DSAG12_02375 | hypothetical protein | -- | 176 | |
| DSAG12_02376 | hypothetical protein | -- | 232 | |
| DSAG12_02377 | Ras-related protein Rab-30 | K07917 | 203 | |
| DSAG12_02378 | hypothetical protein | -- | 205 | |
| DSAG12_02379 | hypothetical protein | -- | 303 | |
| DSAG12_02380 | hypothetical protein | -- | 248 | small GTP-binding domain protein |
| DSAG12_02381 | hypothetical protein | -- | 237 | |
| DSAG12_02382 | glycolate oxidase | K00104 | 268 | |
| DSAG12_02383 | heterodisulfide reductase subunit D | K08264 | 56 | |
| DSAG12_02384 | NAD-dependent deacetylase | K12410 | 247 | |
| DSAG12_02385 | | K06911 | 180 | |
| DSAG12_02386 | hypothetical protein | -- | 131 | |
| DSAG12_02387 | hypothetical protein | -- | 230 | |
| DSAG12_02388 | hypothetical protein | -- | 160 | |
| DSAG12_02389 | hypothetical protein | -- | 202 | |
| DSAG12_02390 | hypothetical protein | -- | 117 | |
| DSAG12_02391 | hypothetical protein | -- | 86 | |
| DSAG12_02392 | hypothetical protein | -- | 187 | |
| DSAG12_02393 | hypothetical protein | -- | 199 | |
| DSAG12_02394 | laminin, alpha 3/5 | K06240 | 83 | |
| DSAG12_02395 | hypothetical protein | -- | 237 | |
| DSAG12_02396 | hypothetical protein | -- | 236 | |
| DSAG12_02397 | hypothetical protein | -- | 279 | |
| DSAG12_02398 | hypothetical protein | -- | 97 | |
| DSAG12_02399 | hypothetical protein | -- | 133 | |
| DSAG12_02400 | glyoxylate reductase | K00015 | 156 | |
| DSAG12_02401 | hypothetical protein | -- | 73 | |

| | | | | |
|---|---|---|---|---|
| DSAG12_02402 | 2'-5' RNA ligase | K01975 | 126 | |
| DSAG12_02403 | Ras-related GTP-binding protein C/D | K16186 | 162 | |
| DSAG12_02404 | orotidine-5'-phosphate decarboxylase | K01591 | 150 | |
| DSAG12_02405 | | K07024 | 115 | |
| DSAG12_02406 | hypothetical protein | -- | 172 | small GTP-binding domain protein |
| DSAG12_02407 | ADP-ribosylation factor-like protein 8 | K07955 | 89 | |
| DSAG12_02408 | hypothetical protein | -- | 148 | |
| DSAG12_02409 | hypothetical protein | -- | 154 | |
| DSAG12_02410 | hypothetical protein | -- | 155 | |
| DSAG12_02411 | elongation factor 2 | K03234 | 318 | |
| DSAG12_02412 | | K00936 | 326 | |
| DSAG12_02413 | two-component system, cell cycle response regulator DivK | K11443 | 201 | |
| DSAG12_02414 | two-component system, cell cycle response regulator DivK | K11443 | 245 | |
| DSAG12_02415 | hypothetical protein | -- | 224 | |
| DSAG12_02416 | | K06944 | 213 | |
| DSAG12_02417 | two-component system, cell cycle response regulator DivK | K11443 | 187 | |
| DSAG12_02418 | hypothetical protein | -- | 203 | |
| DSAG12_02419 | hypothetical protein | -- | 208 | |
| DSAG12_02420 | hypothetical protein | -- | 140 | |
| DSAG12_02421 | hypothetical protein | -- | 131 | |
| DSAG12_02422 | two-component system, cell cycle response regulator DivK | K11443 | 67 | |
| DSAG12_02423 | hypothetical protein | -- | 0 | |
| DSAG12_02424 | hypothetical protein | -- | 258 | |
| DSAG12_02425 | Ca-activated chloride channel homolog | K07114 | 217 | |
| DSAG12_02426 | hypothetical protein | -- | 218 | |
| DSAG12_02427 | energy-coupling factor transport system permease protein / energy-coupling factor transport system ATP-binding protein / energy-coupling factor transport system ATP-binding protein | K16785 K16 | 187 | |
| DSAG12_02428 | energy-coupling factor transport system substrate-specific component | K16925 | 37 | |
| DSAG12_02429 | hypothetical protein | -- | 73 | |
| DSAG12_02430 | hypothetical protein | -- | 195 | |
| DSAG12_02431 | hypothetical protein | -- | 144 | |
| DSAG12_02432 | hypothetical protein | -- | 132 | |
| DSAG12_02433 | isocitrate dehydrogenase | K00031 | 209 | |
| DSAG12_02434 | ATP-citrate lyase alpha-subunit | K15230 | 204 | |
| DSAG12_02435 | ATP-citrate lyase beta-subunit | K15231 | 176 | |
| DSAG12_02436 | aconitate hydratase | K01681 | 169 | |
| DSAG12_02437 | hypothetical protein | -- | 265 | small GTP-binding domain protein |
| DSAG12_02438 | sulfur carrier protein | K03154 | 206 | |
| DSAG12_02439 | peptidyl-prolyl isomerase F (cyclophilin D) | K09565 | 183 | |
| DSAG12_02440 | hypothetical protein | -- | 152 | |
| DSAG12_02441 | acetylornithine deacetylase | K01438 | 234 | |
| DSAG12_02442 | hypothetical protein | -- | 166 | |
| DSAG12_02443 | hypothetical protein | -- | 66 | |
| DSAG12_02444 | aldehyde:ferredoxin oxidoreductase | K03738 | 61 | |
| DSAG12_02445 | CopG family transcriptional regulator, nickel-responsive regulator | K07722 | 125 | |
| DSAG12_02446 | hypothetical protein | -- | 197 | |
| DSAG12_02447 | acetyl-CoA synthetase (ADP-forming) | K01905 | 143 | |
| DSAG12_02448 | hypothetical protein | -- | 168 | |
| DSAG12_02449 | hypothetical protein | -- | 143 | |
| DSAG12_02450 | large subunit ribosomal protein L10e | K02866 | 185 | |
| DSAG12_02451 | hypothetical protein | -- | 109 | |
| DSAG12_02452 | DtxR family transcriptional regulator, Mn-dependent transcriptional regulator | K03709 | 0 | |
| DSAG12_02453 | hypothetical protein | -- | 84 | |
| DSAG12_02454 | hypothetical protein | -- | 175 | |
| DSAG12_02455 | hypothetical protein | -- | 127 | |
| DSAG12_02456 | hypothetical protein | -- | 237 | |
| DSAG12_02457 | DNA topoisomerase I | K03168 | 150 | |
| DSAG12_02458 | DNA replication ATP-dependent helicase Dna2 | K10742 | 157 | |
| DSAG12_02459 | hypothetical protein | -- | 136 | |
| DSAG12_02460 | hypothetical protein | -- | 172 | |
| DSAG12_02461 | dynein light chain roadblock-type | K10419 | 205 | |
| DSAG12_02462 | hypothetical protein | -- | 461 | |
| DSAG12_02463 | hypothetical protein | -- | 159 | |
| DSAG12_02464 | bacterial/archaeal transporter family protein | K08978 | 309 | |
| DSAG12_02465 | UDP-N-acetylglucosamine 3-dehydrogenase | K18855 | 315 | |
| DSAG12_02466 | 3-amino-5-hydroxybenzoate synthase | K16016 | 242 | |
| DSAG12_02467 | hypothetical protein | -- | 283 | |
| DSAG12_02468 | aldehyde:ferredoxin oxidoreductase | K03738 | 157 | |
| DSAG12_02469 | hypothetical protein | -- | 18 | |
| DSAG12_02470 | LAO/AO transport system kinase | K07588 | 177 | |
| DSAG12_02471 | methylmalonyl-CoA mutase, C-terminal domain | K01849 | 138 | |
| DSAG12_02472 | methylmalonyl-CoA mutase, N-terminal domain | K01848 | 236 | |
| DSAG12_02473 | cob(I)alamin adenosyltransferase | K00798 | 177 | |
| DSAG12_02474 | hypothetical protein | -- | 219 | |
| DSAG12_02475 | digeranylgeranylglycerophospholipid reductase | K17830 | 283 | |
| DSAG12_02476 | hypothetical protein | -- | 151 | |
| DSAG12_02477 | hypothetical protein | -- | 179 | |
| DSAG12_02478 | | K07124 | 197 | |

| | | | | |
|---|---|---|---|---|
| DSAG12_02479 | MFS transporter, DHA1 family, multidrug resistance protein | K08153 | 224 | |
| DSAG12_02480 | Ras-related protein Rab-9A | K07899 | 277 | |
| DSAG12_02481 | Ras-related protein Rab-27A | K07885 | 0 | |
| DSAG12_02482 | thioredoxin 1 | K03671 | 373 | |
| DSAG12_02483 | | K07013 | 206 | |
| DSAG12_02484 | 2-isopropylmalate synthase | K01649 | 151 | |
| DSAG12_02485 | hypothetical protein | -- | 165 | |
| DSAG12_02486 | putative endonuclease | K07461 | 157 | |
| DSAG12_02487 | O-phosphoseryl-tRNA(Sec) kinase | K10837 | 179 | |
| DSAG12_02488 | transitional endoplasmic reticulum ATPase | K13525 | 142 | |
| DSAG12_02489 | NAD+ kinase | K00858 | 217 | |
| DSAG12_02490 | hypothetical protein | -- | 191 | |
| DSAG12_02491 | tyrosine phenol-lyase | K01668 | 224 | |
| DSAG12_02492 | indolepyruvate ferredoxin oxidoreductase, beta subunit | K00180 | 146 | |
| DSAG12_02493 | indolepyruvate ferredoxin oxidoreductase, alpha subunit | K00179 | 139 | |
| DSAG12_02494 | penicillin amidase | K01434 | 153 | |
| DSAG12_02495 | hypothetical protein | -- | 132 | |
| DSAG12_02496 | hypothetical protein | -- | 111 | |
| DSAG12_02497 | acetyl-CoA synthetase (ADP-forming) | K01905 | 206 | |
| DSAG12_02498 | hypothetical protein | -- | 208 | |
| DSAG12_02499 | hypothetical protein | -- | 191 | |
| DSAG12_02500 | hypothetical protein | -- | 238 | |
| DSAG12_02501 | methionyl-tRNA synthetase | K01874 | 130 | |
| DSAG12_02502 | hypothetical protein | -- | 219 | |
| DSAG12_02503 | hypothetical protein | K09721 | 194 | |
| DSAG12_02504 | beta-galactosidase | K01190 | 255 | |
| DSAG12_02505 | hypothetical protein | -- | 158 | |
| DSAG12_02506 | alpha-mannosidase | K01191 | 175 | |
| DSAG12_02507 | hypothetical protein | -- | 193 | |
| DSAG12_02508 | hypothetical protein | -- | 163 | |
| DSAG12_02509 | hypothetical protein | -- | 36 | |
| DSAG12_02510 | hypothetical protein | -- | 147 | |
| DSAG12_02511 | hypothetical protein | -- | 156 | |
| DSAG12_02512 | hypothetical protein | -- | 175 | small GTP-binding domain protein |
| DSAG12_02513 | peroxiredoxin Q/BCP | K03564 | 195 | |
| DSAG12_02514 | hypothetical protein | -- | 107 | |
| DSAG12_02515 | hypothetical protein | -- | 178 | |
| DSAG12_02516 | hypothetical protein | -- | 21 | |
| DSAG12_02517 | hypothetical protein | -- | 167 | |
| DSAG12_02518 | hypothetical protein | -- | 119 | |
| DSAG12_02519 | | K07052 | 139 | |
| DSAG12_02520 | hypothetical protein | -- | 260 | |
| DSAG12_02521 | hypothetical protein | -- | 206 | |
| DSAG12_02522 | hypothetical protein | -- | 193 | hypothetical proteins with gelsolin-like domain |
| DSAG12_02523 | hypothetical protein | -- | 279 | |
| DSAG12_02524 | putative hydrolase of the HAD superfamily | K07025 | 203 | |
| DSAG12_02525 | hypothetical protein | -- | 261 | |
| DSAG12_02526 | hypothetical protein | -- | 238 | |
| DSAG12_02527 | hypothetical protein | -- | 149 | |
| DSAG12_02528 | hypothetical protein | -- | 252 | |
| DSAG12_02529 | hypothetical protein | -- | 226 | |
| DSAG12_02530 | hypothetical protein | -- | 54 | |
| DSAG12_02531 | hypothetical protein | -- | 145 | |
| DSAG12_02532 | hypothetical protein | -- | 223 | |
| DSAG12_02533 | hypothetical protein | -- | 195 | |
| DSAG12_02534 | two-component system, OmpR family, sensor kinase | K02484 | 213 | |
| DSAG12_02535 | hypothetical protein | -- | 116 | |
| DSAG12_02536 | hypothetical protein | -- | 205 | |
| DSAG12_02537 | arylformamidase | K07130 | 128 | |
| DSAG12_02538 | protein phosphatase | K01090 | 177 | |
| DSAG12_02539 | hypothetical protein | -- | 175 | |
| DSAG12_02540 | hypothetical protein | -- | 78 | |
| DSAG12_02541 | O-acetylserine/cysteine efflux transporter | K15268 | 158 | |
| DSAG12_02542 | hypothetical protein | -- | 159 | |
| DSAG12_02543 | menaquinone-dependent protoporphyrinogen oxidase | K00230 | 0 | |
| DSAG12_02544 | hypothetical protein | -- | 166 | |
| DSAG12_02545 | alpha-mannosidase | K01191 | 307 | |
| DSAG12_02546 | hypothetical protein | -- | 0 | |
| DSAG12_02547 | hypothetical protein | -- | 102 | |
| DSAG12_02548 | hypothetical protein | -- | 250 | |
| DSAG12_02549 | hypothetical protein | -- | 306 | |
| DSAG12_02550 | two-component system, cell cycle sensor histidine kinase and response regulator CckA | K13587 | 233 | |
| DSAG12_02551 | hypothetical protein | -- | 125 | |
| DSAG12_02552 | hypothetical protein | -- | 125 | |
| DSAG12_02553 | hypothetical protein | -- | 188 | |
| DSAG12_02554 | hypothetical protein | -- | 96 | |
| DSAG12_02555 | DNA excision repair protein ERCC-2 | K10844 | 49 | |

| | | | |
|---|---|---|---|
| DSAG12_02556 | putative acetyltransferase | K03828 | 42 |
| DSAG12_02557 | L-serine dehydratase | K01752 | 157 |
| DSAG12_02558 | putative ABC transport system ATP-binding protein | K02003 | 166 |
| DSAG12_02559 | hypothetical protein | -- | 146 |
| DSAG12_02560 | putative ABC transport system ATP-binding protein | K02003 | 203 |
| DSAG12_02561 | hypothetical protein | -- | 164 |
| DSAG12_02562 | hypothetical protein | -- | 195 |
| DSAG12_02563 | hypothetical protein | -- | 61 |
| DSAG12_02564 | hypothetical protein | -- | 128 |
| DSAG12_02565 | protease I | K05520 | 231 |
| DSAG12_02566 | hypothetical protein | -- | 0 |
| DSAG12_02567 | hypothetical protein | -- | 48 |
| DSAG12_02568 | hypothetical protein | -- | 183 |
| DSAG12_02569 | hypothetical protein | -- | 139 |
| DSAG12_02570 | hypothetical protein | -- | 148 |
| DSAG12_02571 | hypothetical protein | -- | 198 |
| DSAG12_02572 | | K06883 | 115 |
| DSAG12_02573 | hypothetical protein | -- | 188 |
| DSAG12_02574 | hypothetical protein | -- | 116 |
| DSAG12_02575 | hypothetical protein | -- | 206 |
| DSAG12_02576 | hypothetical protein | -- | 123 |
| DSAG12_02577 | xylulokinase | K00854 | 188 |
| DSAG12_02578 | glycerol kinase | K00864 | 194 |
| DSAG12_02579 | hypothetical protein | -- | 117 |
| DSAG12_02580 | hypothetical protein | -- | 188 |
| DSAG12_02581 | hypothetical protein | -- | 136 |
| DSAG12_02582 | hypothetical protein | -- | 158 |
| DSAG12_02583 | hypothetical protein | -- | 68 |
| DSAG12_02584 | hypothetical protein | -- | 188 |
| DSAG12_02585 | hypothetical protein | -- | 184 |
| DSAG12_02586 | hypothetical protein | -- | 139 |
| DSAG12_02587 | hypothetical protein | -- | 182 |
| DSAG12_02588 | hypothetical protein | -- | 158 |
| DSAG12_02589 | hypothetical protein | -- | 172 |
| DSAG12_02590 | hypothetical protein | -- | 186 |
| DSAG12_02591 | hypothetical protein | -- | 118 |
| DSAG12_02592 | hypothetical protein | -- | 220 |
| DSAG12_02593 | hypothetical protein | -- | 236 |
| DSAG12_02594 | hypothetical protein | -- | 265 |
| DSAG12_02595 | hypothetical protein | -- | 243 |
| DSAG12_02596 | ATP-dependent helicase IRC3 | K17677 | 137 |
| DSAG12_02597 | hypothetical protein | -- | 127 |
| DSAG12_02598 | hypothetical protein | -- | 306 |
| DSAG12_02599 | ribonuclease III | K03685 | 179 |
| DSAG12_02600 | hypothetical protein | -- | 135 |
| DSAG12_02601 | hypothetical protein | -- | 159 |
| DSAG12_02602 | hypothetical protein | -- | 241 |
| DSAG12_02603 | hypothetical protein | -- | 151 |
| DSAG12_02604 | RelE protein | K06218 | 203 |
| DSAG12_02605 | hypothetical protein | -- | 163 |
| DSAG12_02606 | hypothetical protein | -- | 96 |
| DSAG12_02607 | hypothetical protein | -- | 171 |
| DSAG12_02608 | hypothetical protein | -- | 134 |
| DSAG12_02609 | hypothetical protein | -- | 292 |
| DSAG12_02610 | hypothetical protein | -- | 217 |
| DSAG12_02611 | hypothetical protein | -- | 214 |
| DSAG12_02612 | hypothetical protein | -- | 274 |
| DSAG12_02613 | protein scribble | K16175 | 0 |
| DSAG12_02614 | hypothetical protein | -- | 55 |
| DSAG12_02615 | hypothetical protein | -- | 134 |
| DSAG12_02616 | hypothetical protein | -- | 88 |
| DSAG12_02617 | hypothetical protein | -- | 152 |
| DSAG12_02618 | hypothetical protein | -- | 0 |
| DSAG12_02619 | hypothetical protein | -- | 0 |
| DSAG12_02620 | hypothetical protein | -- | 100 |
| DSAG12_02621 | hypothetical protein | -- | 201 |
| DSAG12_02622 | hypothetical protein | -- | 108 |
| DSAG12_02623 | hypothetical protein | -- | 203 |
| DSAG12_02624 | hypothetical protein | -- | 130 |
| DSAG12_02625 | hypothetical protein | -- | 138 |
| DSAG12_02626 | hypothetical protein | -- | 135 |
| DSAG12_02627 | acetyl-CoA synthetase | K01895 | 125 |
| DSAG12_02628 | hypothetical protein | -- | 200 |
| DSAG12_02629 | hypothetical protein | -- | 139 |
| DSAG12_02630 | putative RecB family exonuclease | K07464 | 113 |
| DSAG12_02631 | hypothetical protein | -- | 226 |
| DSAG12_02632 | hypothetical protein | -- | 243 |
| DSAG12_02633 | hypothetical protein | -- | 184 |

| | | | |
|---|---|---|---|
| DSAG12_02634 | hypothetical protein | -- | 31 |
| DSAG12_02635 | hypothetical protein | -- | 143 |
| DSAG12_02636 | hypothetical protein | -- | 165 |
| DSAG12_02637 | | K07052 | 168 |
| DSAG12_02638 | hypothetical protein | -- | 82 |
| DSAG12_02639 | hypothetical protein | -- | 0 |
| DSAG12_02640 | transposase | K07486 | 180 |
| DSAG12_02641 | acetyl-CoA synthetase | K01895 | 0 |
| DSAG12_02642 | hypothetical protein | -- | 88 |
| DSAG12_02643 | hypothetical protein | -- | 93 |
| DSAG12_02644 | hypothetical protein | -- | 107 |
| DSAG12_02645 | hypothetical protein | -- | 126 |
| DSAG12_02646 | hypothetical protein | -- | 65 |
| DSAG12_02647 | hypothetical protein | -- | 172 |
| DSAG12_02648 | hypothetical protein | -- | 0 |
| DSAG12_02649 | hypothetical protein | -- | 98 |
| DSAG12_02650 | hypothetical protein | -- | 107 |
| DSAG12_02651 | hypothetical protein | -- | 126 |
| DSAG12_02652 | hypothetical protein | -- | 103 |
| DSAG12_02653 | hypothetical protein | -- | 33 |
| DSAG12_02654 | hypothetical protein | -- | 227 |
| DSAG12_02655 | hypothetical protein | -- | 132 |
| DSAG12_02656 | hypothetical protein | -- | 124 |
| DSAG12_02657 | hypothetical protein | -- | 114 |
| DSAG12_02658 | hypothetical protein | -- | 184 |
| DSAG12_02659 | inner membrane protein | K07038 | 32 |
| DSAG12_02660 | hypothetical protein | -- | 148 |
| DSAG12_02661 | hypothetical protein | -- | 78 |
| DSAG12_02662 | hypothetical protein | -- | 170 |
| DSAG12_02663 | pyruvate formate lyase activating enzyme | K04069 | 165 |
| DSAG12_02664 | hypothetical protein | -- | 72 |
| DSAG12_02665 | hypothetical protein | -- | 170 |
| DSAG12_02666 | hypothetical protein | -- | 0 |
| DSAG12_02667 | hypothetical protein | -- | 84 |
| DSAG12_02668 | archaea-specific RecJ-like exonuclease | K07463 | 65 |
| DSAG12_02669 | hypothetical protein | -- | 189 |
| DSAG12_02670 | kynureninase | K01556 | 164 |
| DSAG12_02671 | hypothetical protein | -- | 212 |
| DSAG12_02672 | hypothetical protein | -- | 138 |
| DSAG12_02673 | putative protease | K08303 | 83 |
| DSAG12_02674 | ATP-dependent RNA helicase DeaD | K05592 | 92 |
| DSAG12_02675 | hypothetical protein | -- | 108 |
| DSAG12_02676 | hypothetical protein | -- | 202 |
| DSAG12_02677 | hypothetical protein | -- | 265 |
| DSAG12_02678 | hypothetical protein | -- | 155 |
| DSAG12_02679 | bifunctional UDP-N-acetylglucosamine pyrophosphorylase / Glucosamine-1-phosphate N-acetyltransferase | K04042 | 27 |
| DSAG12_02680 | | K00100 | 391 |
| DSAG12_02681 | small subunit ribosomal protein S3Ae | K02984 | 142 |
| DSAG12_02682 | small subunit ribosomal protein S15 | K02956 | 139 |
| DSAG12_02683 | hypothetical protein | -- | 150 |
| DSAG12_02684 | hypothetical protein | -- | 149 |
| DSAG12_02685 | hypothetical protein | -- | 145 |
| DSAG12_02686 | demethylmenaquinone methyltransferase / 2-methoxy-6-polyprenyl-1,4-benzoquinol methylase | K03183 | 181 |
| DSAG12_02687 | XTP/dITP diphosphohydrolase | K02428 | 143 |
| DSAG12_02688 | hypothetical protein | -- | 217 |
| DSAG12_02689 | hypothetical protein | -- | 45 |
| DSAG12_02690 | Xaa-Pro dipeptidase | K01271 | 123 |
| DSAG12_02691 | hypothetical protein | -- | 74 |
| DSAG12_02692 | hypothetical protein | -- | 134 |
| DSAG12_02693 | hypothetical protein | -- | 147 |
| DSAG12_02694 | guanidinopropionase | K18459 | 335 |
| DSAG12_02695 | deoxyhypusine synthase | K00809 | 41 |
| DSAG12_02696 | | K07079 | 118 |
| DSAG12_02697 | hypothetical protein | -- | 120 |
| DSAG12_02698 | hypothetical protein | -- | 187 |
| DSAG12_02699 | hypothetical protein | -- | 144 |
| DSAG12_02700 | demethylmenaquinone methyltransferase / 2-methoxy-6-polyprenyl-1,4-benzoquinol methylase | K03183 | 185 |
| DSAG12_02701 | hypothetical protein | -- | 193 |
| DSAG12_02702 | | K07052 | 189 |
| DSAG12_02703 | MFS transporter, DHA1 family, multidrug resistance protein | K08153 | 229 |
| DSAG12_02704 | hypothetical protein | -- | 116 |
| DSAG12_02705 | hypothetical protein | -- | 252 |
| DSAG12_02706 | acyl-CoA synthetase | K00142 | 35 |
| DSAG12_02707 | hypothetical protein | -- | 231 |
| DSAG12_02708 | hypothetical protein | -- | 122 |
| DSAG12_02709 | small subunit ribosomal protein S10 | K02946 | 198 |
| DSAG12_02710 | hypothetical protein | -- | 79 |

| | | | |
|---|---|---|---|
| DSAG12_02711 | hypothetical protein | -- | 167 |
| DSAG12_02712 | hypothetical protein | -- | 217 |
| DSAG12_02713 | | K07057 | 194 |
| DSAG12_02714 | hypothetical protein | -- | 161 |
| DSAG12_02715 | hypothetical protein | -- | 187 |
| DSAG12_02716 | tRNA-Ser | | 134 |
| DSAG12_02717 | inorganic phosphate transporter, PiT family | K03306 | 169 |
| DSAG12_02718 | hypothetical protein | -- | 180 |
| DSAG12_02719 | hypothetical protein | -- | 239 |
| DSAG12_02720 | hypothetical protein | -- | 205 |
| DSAG12_02721 | ribosomal-protein-alanine N-acetyltransferase | K03790 | 228 |
| DSAG12_02722 | hypothetical protein | -- | 58 |
| DSAG12_02723 | hypothetical protein | -- | 0 |
| DSAG12_02724 | glycine C-acetyltransferase | K00639 | 214 |
| DSAG12_02725 | UDP-glucose 4-epimerase | K01784 | 190 |
| DSAG12_02726 | hypothetical protein | -- | 166 |
| DSAG12_02727 | hypothetical protein | -- | 125 |
| DSAG12_02728 | | K07018 | 65 |
| DSAG12_02729 | hypothetical protein | -- | 142 |
| DSAG12_02730 | hypothetical protein | -- | 212 |
| DSAG12_02731 | Xaa-Pro aminopeptidase | K01262 | 138 |
| DSAG12_02732 | hypothetical protein | -- | 170 |
| DSAG12_02733 | hypothetical protein | -- | 170 |
| DSAG12_02734 | hypothetical protein | -- | 191 |
| DSAG12_02735 | hypothetical protein | -- | 187 |
| DSAG12_02736 | COMPASS component SWD3 | K14963 | 201 |
| DSAG12_02737 | spermidine synthase | K00797 | 217 |
| DSAG12_02738 | hypothetical protein | -- | 128 |
| DSAG12_02739 | hypothetical protein | -- | 5 |
| DSAG12_02740 | tRNA 2-thiouridine synthesizing protein A | K04085 | 95 |
| DSAG12_02741 | adenylylsulfate reductase, subunit B | K00395 | 148 |
| DSAG12_02742 | hypothetical protein | -- | 172 |
| DSAG12_02743 | formate dehydrogenase, beta subunit | K00125 | 146 |
| DSAG12_02744 | hypothetical protein | -- | 202 |
| DSAG12_02745 | hypothetical protein | -- | 170 |
| DSAG12_02746 | hypothetical protein | -- | 274 |
| DSAG12_02747 | hypothetical protein | -- | 299 |
| DSAG12_02748 | hypothetical protein | -- | 30 |
| DSAG12_02749 | hypothetical protein | -- | 128 |
| DSAG12_02750 | hypothetical protein | -- | 156 |
| DSAG12_02751 | hypothetical protein | -- | 174 |
| DSAG12_02752 | hypothetical protein | -- | 87 |
| DSAG12_02753 | hypothetical protein | -- | 147 |
| DSAG12_02754 | hypothetical protein | -- | 142 |
| DSAG12_02755 | hypothetical protein | -- | 161 |
| DSAG12_02756 | hypothetical protein | -- | 0 |
| DSAG12_02757 | hypothetical protein | -- | 241 |
| DSAG12_02758 | hypothetical protein | -- | 202 |
| DSAG12_02759 | hypothetical protein | -- | 118 |
| DSAG12_02760 | phenylglyoxylate dehydrogenase beta subunit | K18356 | 164 |
| DSAG12_02761 | aldehyde:ferredoxin oxidoreductase | K03738 | 100 |
| DSAG12_02762 | hypothetical protein | -- | 155 |
| DSAG12_02763 | F420-non-reducing hydrogenase iron-sulfur subunit | K14127 | 230 |
| DSAG12_02764 | heterodisulfide reductase subunit A | K03388 | 254 |
| DSAG12_02765 | heterodisulfide reductase subunit C | K03390 | 221 |
| DSAG12_02766 | hypothetical protein | -- | 129 |
| DSAG12_02767 | 4Fe-4S ferredoxin | K00205 | 112 |
| DSAG12_02768 | hypothetical protein | -- | 164 |
| DSAG12_02769 | hypothetical protein | -- | 196 |
| DSAG12_02770 | | K07052 | 195 |
| DSAG12_02771 | hypothetical protein | -- | 233 |
| DSAG12_02772 | antitoxin MazE | K07172 | 135 |
| DSAG12_02773 | mRNA interferase MazF | K07171 | 120 |
| DSAG12_02774 | 16S rRNA (cytosine967-C5)-methyltransferase | K03500 | 164 |
| DSAG12_02775 | hypothetical protein | -- | 211 |
| DSAG12_02776 | hypothetical protein | -- | 141 |
| DSAG12_02777 | hypothetical protein | -- | 130 |
| DSAG12_02778 | hypothetical protein | -- | 195 |
| DSAG12_02779 | cysteine synthase A | K01738 | 147 |
| DSAG12_02780 | hypothetical protein | -- | 243 |
| DSAG12_02781 | hypothetical protein | -- | 54 |
| DSAG12_02782 | AraC family transcriptional regulator, regulatory protein of adaptative response / methylated-DNA-[protein]-cysteine methyltransferase | K10778 | 136 |
| DSAG12_02783 | methylated-DNA-[protein]-cysteine S-methyltransferase | K00567 | 233 |
| DSAG12_02784 | hypothetical protein | -- | 123 |
| DSAG12_02785 | hypothetical protein | -- | 204 |
| DSAG12_02786 | small conductance mechanosensitive channel | K03442 | 109 |
| DSAG12_02787 | hypothetical protein | -- | 231 |

| | | | | |
|---|---|---|---|---|
| DSAG12_02788 | 3-hydroxyisobutyrate dehydrogenase | K00020 | 318 | |
| DSAG12_02789 | hypothetical protein | -- | 26 | |
| DSAG12_02790 | hypothetical protein | -- | 102 | |
| DSAG12_02791 | NADH oxidase | K00359 | 125 | |
| DSAG12_02792 | hypothetical protein | -- | 137 | |
| DSAG12_02793 | integrase/recombinase XerC | K03733 | 31 | |
| DSAG12_02794 | hypothetical protein | -- | 154 | |
| DSAG12_02795 | hypothetical protein | -- | 171 | |
| DSAG12_02796 | hypothetical protein | -- | 136 | |
| DSAG12_02797 | hypothetical protein | -- | 124 | |
| DSAG12_02798 | dGTPase | K01129 | 138 | |
| DSAG12_02799 | hypothetical protein | -- | 148 | |
| DSAG12_02800 | site-specific DNA-methyltransferase (cytosine-N4-specific) | K00590 | 190 | |
| DSAG12_02801 | hypothetical protein | -- | 295 | |
| DSAG12_02802 | hypothetical protein | -- | 171 | |
| DSAG12_02803 | hypothetical protein | -- | 146 | |
| DSAG12_02804 | hypothetical protein | -- | 257 | |
| DSAG12_02805 | hypothetical protein | -- | 171 | |
| DSAG12_02806 | ATP-binding cassette, subfamily B, bacterial MsbA | K11085 | 208 | |
| DSAG12_02807 | ATP-binding cassette, subfamily B, bacterial | K06147 | 194 | |
| DSAG12_02808 | hypothetical protein | -- | 0 | |
| DSAG12_02809 | hypothetical protein | -- | 186 | |
| DSAG12_02810 | hypothetical protein | -- | 163 | |
| DSAG12_02811 | proline iminopeptidase | K01259 | 156 | |
| DSAG12_02812 | hypothetical protein | -- | 0 | |
| DSAG12_02813 | actin beta/gamma 1 | K05692 | 64 | |
| DSAG12_02814 | Ras-related protein Rab-14 | K07881 | 107 | |
| DSAG12_02815 | hypothetical protein | -- | 0 | |
| DSAG12_02816 | hypothetical protein | -- | 181 | |
| DSAG12_02817 | hypothetical protein | -- | 192 | |
| DSAG12_02818 | hypothetical protein | -- | 124 | |
| DSAG12_02819 | MFS transporter, OFA family, oxalate/formate antiporter | K08177 | 0 | |
| DSAG12_02820 | hypothetical protein | -- | 233 | |
| DSAG12_02821 | hypothetical protein | -- | 131 | |
| DSAG12_02822 | hypothetical protein | -- | 258 | |
| DSAG12_02823 | hypothetical protein | -- | 106 | |
| DSAG12_02824 | hypothetical protein | -- | 173 | |
| DSAG12_02825 | hypothetical protein | -- | 153 | |
| DSAG12_02826 | hypothetical protein | -- | 171 | |
| DSAG12_02827 | thioredoxin reductase (NADPH) | K00384 | 143 | |
| DSAG12_02828 | thioredoxin 1 | K03671 | 144 | |
| DSAG12_02829 | hypothetical protein | -- | 157 | |
| DSAG12_02830 | hypothetical protein | -- | 173 | |
| DSAG12_02831 | hypothetical protein | -- | 203 | |
| DSAG12_02832 | hypothetical protein | -- | 174 | |
| DSAG12_02833 | putative transcriptional regulator | K07729 | 211 | |
| DSAG12_02834 | hypothetical protein | -- | 238 | |
| DSAG12_02835 | ABC-2 type transport system ATP-binding protein | K01990 | 135 | |
| DSAG12_02836 | hypothetical protein | -- | 133 | |
| DSAG12_02837 | hypothetical protein | -- | 82 | |
| DSAG12_02838 | hypothetical protein | -- | 239 | |
| DSAG12_02839 | hypothetical protein | -- | 61 | |
| DSAG12_02840 | hypothetical protein | -- | 122 | |
| DSAG12_02841 | hypothetical protein | -- | 197 | |
| DSAG12_02842 | hypothetical protein | -- | 175 | |
| DSAG12_02843 | thiosulfate/3-mercaptopyruvate sulfurtransferase | K01011 | 189 | |
| DSAG12_02844 | 2-oxoglutarate ferredoxin oxidoreductase subunit delta | K00176 | 193 | |
| DSAG12_02845 | 2-oxoglutarate ferredoxin oxidoreductase subunit alpha | K00174 | 7 | |
| DSAG12_02846 | 2-oxoglutarate ferredoxin oxidoreductase subunit beta | K00175 | 6 | |
| DSAG12_02847 | 2-oxoglutarate ferredoxin oxidoreductase subunit gamma | K00177 | 253 | |
| DSAG12_02848 | MFS transporter, DHA1 family, tetracycline resistance protein | K08151 | 157 | |
| DSAG12_02849 | hypothetical protein | -- | 217 | |
| DSAG12_02850 | Ras-related protein Rab-30 | K07917 | 185 | |
| DSAG12_02851 | hypothetical protein | -- | 140 | |
| DSAG12_02852 | hypothetical protein | -- | 151 | |
| DSAG12_02853 | hypothetical protein | -- | 0 | |
| DSAG12_02854 | actin-related protein 6 | K11662 | 104 | |
| DSAG12_02855 | hypothetical protein | -- | 176 | |
| DSAG12_02856 | hypothetical protein | -- | 47 | |
| DSAG12_02857 | archaeal cell division control protein 6 | K10725 | 212 | related) |
| DSAG12_02858 | serine/threonine protein kinase, bacterial | K08884 | 202 | small GTP-binding domain protein |
| DSAG12_02859 | hypothetical protein | -- | 244 | |
| DSAG12_02860 | | K00936 | 121 | |
| DSAG12_02861 | NAD+ synthase | K01916 | 191 | |
| DSAG12_02862 | aspartate aminotransferase | K00812 | 199 | |
| DSAG12_02863 | hypothetical protein | -- | 11 | |
| DSAG12_02864 | hypothetical protein | -- | 179 | |
| DSAG12_02865 | hypothetical protein | -- | 71 | |

| | | | |
|---|---|---|---|
| DSAG12_02866 | hypothetical protein | -- | 215 |
| DSAG12_02867 | acetyltransferase | K09181 | 205 |
| DSAG12_02868 | hypothetical protein | -- | 225 |
| DSAG12_02869 | hypothetical protein | -- | 206 |
| DSAG12_02870 | hypothetical protein | -- | 10 |
| DSAG12_02871 | hypothetical protein | -- | 283 |
| DSAG12_02872 | hypothetical protein | -- | 173 |
| DSAG12_02873 | adenylosuccinate synthase | K01939 | 223 |
| DSAG12_02874 | energy-coupling factor transport system substrate-specific component | K16927 | 157 |
| DSAG12_02875 | energy-coupling factor transport system ATP-binding protein / energy-coupling factor transport system ATP-binding protein | K16786 K16 | 133 |
| DSAG12_02876 | energy-coupling factor transport system permease protein | K16785 | 208 |
| DSAG12_02877 | 2,3-bisphosphoglycerate-dependent phosphoglycerate mutase | K01834 | 21 |
| DSAG12_02878 | 2,3-bisphosphoglycerate-dependent phosphoglycerate mutase | K01834 | 196 |
| DSAG12_02879 | 2,3-bisphosphoglycerate-dependent phosphoglycerate mutase | K01834 | 186 |
| DSAG12_02880 | hypothetical protein | -- | 158 |
| DSAG12_02881 | hypothetical protein | -- | 93 |
| DSAG12_02882 | histidine triad (HIT) family protein | K02503 | 115 |
| DSAG12_02883 | | K00680 | 164 |
| DSAG12_02884 | hypothetical protein | -- | 218 |
| DSAG12_02885 | hypothetical protein | -- | 141 |
| DSAG12_02886 | | K06930 | 115 |
| DSAG12_02887 | hypothetical protein | -- | 174 |
| DSAG12_02888 | hypothetical protein | -- | 72 |
| DSAG12_02889 | hypothetical protein | -- | 255 |
| DSAG12_02890 | hypothetical protein | -- | 281 |
| DSAG12_02891 | hypothetical protein | -- | 164 |
| DSAG12_02892 | hypothetical protein | -- | 194 |
| DSAG12_02893 | MFS transporter, DHA1 family, tetracycline resistance protein | K08151 | 138 |
| DSAG12_02894 | hypothetical protein | -- | 231 |
| DSAG12_02895 | fructose-bisphosphate aldolase, class II | K01624 | 172 |
| DSAG12_02896 | arabinose-5-phosphate isomerase | K06041 | 138 |
| DSAG12_02897 | dipeptidase D | K01270 | 132 |
| DSAG12_02898 | | K07059 | 145 |
| DSAG12_02899 | hypothetical protein | -- | 0 |
| DSAG12_02900 | 2-hydroxy-3-keto-5-methylthiopentenyl-1-phosphate phosphatase | K08966 | 195 |
| DSAG12_02901 | hypothetical protein | -- | 55 |
| DSAG12_02902 | hypothetical protein | -- | 219 |
| DSAG12_02903 | hypothetical protein | -- | 205 |
| DSAG12_02904 | hypothetical protein | -- | 184 |
| DSAG12_02905 | alkyldihydroxyacetonephosphate synthase | K00803 | 171 |
| DSAG12_02906 | heterodisulfide reductase subunit D | K08264 | 188 |
| DSAG12_02907 | glycerol kinase | K00864 | 228 |
| DSAG12_02908 | hypothetical protein | -- | 125 |
| DSAG12_02909 | hypothetical protein | -- | 226 |
| DSAG12_02910 | hypothetical protein | -- | 227 |
| DSAG12_02911 | tRNA-intron endonuclease, archaea type | K01170 | 181 |
| DSAG12_02912 | ATPase | K06865 | 178 |
| DSAG12_02913 | CDP-diacylglycerol---serine O-phosphatidyltransferase | K17103 | 180 |
| DSAG12_02914 | glycoside/pentoside/hexuronide:cation symporter, GPH family | K03292 | 130 |
| DSAG12_02915 | hypothetical protein | -- | 141 |
| DSAG12_02916 | hypothetical protein | -- | 275 |
| DSAG12_02917 | hypothetical protein | -- | 174 |
| DSAG12_02918 | protein scribble | K16175 | 219 |
| DSAG12_02919 | hypothetical protein | -- | 161 |
| DSAG12_02920 | hypothetical protein | -- | 168 |
| DSAG12_02921 | | K06883 | 100 |
| DSAG12_02922 | hypothetical protein | -- | 217 |
| DSAG12_02923 | 2,3-diketo-5-methylthiopentyl-1-phosphate enolase | K08965 | 193 |
| DSAG12_02924 | hypothetical protein | -- | 176 |
| DSAG12_02925 | xylulokinase | K00854 | 199 |
| DSAG12_02926 | hypothetical protein | -- | 171 |
| DSAG12_02927 | glutamate-1-semialdehyde 2,1-aminomutase | K01845 | 165 |
| DSAG12_02928 | hypothetical protein | -- | 224 |
| DSAG12_02929 | hypothetical protein | -- | 224 |
| DSAG12_02930 | hypothetical protein | -- | 161 |
| DSAG12_02931 | hypothetical protein | -- | 195 |
| DSAG12_02932 | hypothetical protein | -- | 262 |
| DSAG12_02933 | | K06930 | 405 |
| DSAG12_02934 | hypothetical protein | -- | 93 |
| DSAG12_02935 | protoheme IX farnesyltransferase | K02301 | 300 |
| DSAG12_02936 | phytol kinase | K18678 | 571 |
| DSAG12_02937 | isopentenyl phosphate kinase | K06981 | 269 |
| DSAG12_02938 | hypothetical protein | -- | 208 |
| DSAG12_02939 | hypothetical protein | -- | 97 |
| DSAG12_02940 | hypothetical protein | -- | 0 |
| DSAG12_02941 | hypothetical protein | -- | 247 |
| DSAG12_02942 | hypothetical protein | -- | 194 |

| | | | | |
|---|---|---|---|---|
| DSAG12_02943 | hypothetical protein | -- | 196 | |
| DSAG12_02944 | hypothetical protein | -- | 146 | |
| DSAG12_02945 | hypothetical protein | -- | 173 | |
| DSAG12_02946 | hypothetical protein | -- | 155 | |
| DSAG12_02947 | UDP-glucose 4-epimerase | K01784 | 97 | |
| DSAG12_02948 | hypothetical protein | -- | 51 | |
| DSAG12_02949 | | K00936 | 150 | |
| DSAG12_02950 | hypothetical protein | K07502 | 64 | |
| DSAG12_02951 | hypothetical protein | -- | 247 | |
| DSAG12_02952 | putative RecB family exonuclease | K07464 | 241 | |
| DSAG12_02953 | hypothetical protein | -- | 205 | |
| DSAG12_02954 | hypothetical protein | -- | 127 | |
| DSAG12_02955 | hypothetical protein | -- | 81 | |
| DSAG12_02956 | | K07076 | 181 | |
| DSAG12_02957 | | K07076 | 170 | |
| DSAG12_02958 | hypothetical protein | -- | 209 | |
| DSAG12_02959 | hypothetical protein | -- | 170 | |
| DSAG12_02960 | 2-(1,2-epoxy-1,2-dihydrophenyl)acetyl-CoA isomerase | K15866 | 211 | |
| DSAG12_02961 | hypothetical protein | -- | 117 | |
| DSAG12_02962 | hypothetical protein | -- | 172 | |
| DSAG12_02963 | hypothetical protein | -- | 153 | |
| DSAG12_02964 | hypothetical protein | -- | 241 | |
| DSAG12_02965 | DNA (cytosine-5)-methyltransferase 1 | K00558 | 0 | |
| DSAG12_02966 | A/G-specific adenine glycosylase | K03575 | 235 | |
| DSAG12_02967 | hypothetical protein | -- | 181 | |
| DSAG12_02968 | | K06915 | 197 | |
| DSAG12_02969 | exonuclease SbcC | K03546 | 65 | |
| DSAG12_02970 | hypothetical protein | -- | 276 | |
| DSAG12_02971 | | K00936 | 64 | |
| DSAG12_02972 | hypothetical protein | -- | 240 | |
| DSAG12_02973 | hypothetical protein | -- | 183 | |
| DSAG12_02974 | hypothetical protein | -- | 264 | |
| DSAG12_02975 | hypothetical protein | -- | 175 | |
| DSAG12_02976 | hypothetical protein | -- | 170 | |
| DSAG12_02977 | hypothetical protein | -- | 178 | |
| DSAG12_02978 | hypothetical protein | -- | 356 | hypothetical proteins with gelsolin-like domain |
| DSAG12_02979 | COMPASS component SWD3 | K14963 | 221 | |
| DSAG12_02980 | E3 ubiquitin-protein ligase TRAF7 | K10646 | 117 | |
| DSAG12_02981 | hypothetical protein | -- | 209 | |
| DSAG12_02982 | transcription initiation factor TFIID TATA-box-binding protein | K03120 | 153 | |
| DSAG12_02983 | hypothetical protein | -- | 170 | |
| DSAG12_02984 | hypothetical protein | -- | 201 | |
| DSAG12_02985 | | K07065 | 205 | |
| DSAG12_02986 | hypothetical protein | -- | 172 | |
| DSAG12_02987 | hypothetical protein | -- | 182 | |
| DSAG12_02988 | hypothetical protein | -- | 200 | |
| DSAG12_02989 | hypothetical protein | -- | 244 | |
| DSAG12_02990 | hypothetical protein | -- | 132 | |
| DSAG12_02991 | hypothetical protein | -- | 217 | |
| DSAG12_02992 | hypothetical protein | -- | 178 | |
| DSAG12_02993 | hypothetical protein | -- | 204 | |
| DSAG12_02994 | tryptophan synthase beta chain | K01696 | 246 | |
| DSAG12_02995 | hypothetical protein | -- | 16 | |
| DSAG12_02996 | hypothetical protein | -- | 0 | |
| DSAG12_02997 | hypothetical protein | -- | 0 | |
| DSAG12_02998 | hypothetical protein | -- | 238 | |
| DSAG12_02999 | A/G-specific adenine glycosylase | K03575 | 176 | |
| DSAG12_03000 | hypothetical protein | -- | 188 | |
| DSAG12_03001 | hypothetical protein | -- | 218 | |
| DSAG12_03002 | ribosomal-protein-alanine N-acetyltransferase | K03790 | 59 | |
| DSAG12_03003 | tripeptide aminopeptidase | K01258 | 164 | |
| DSAG12_03004 | hypothetical protein | -- | 216 | |
| DSAG12_03005 | hypothetical protein | -- | 209 | |
| DSAG12_03006 | ABC-2 type transport system ATP-binding protein | K01990 | 131 | |
| DSAG12_03007 | hypothetical protein | -- | 11 | |
| DSAG12_03008 | hypothetical protein | -- | 15 | |
| DSAG12_03009 | hypothetical protein | -- | 214 | |
| DSAG12_03010 | hypothetical protein | -- | 8 | |
| DSAG12_03011 | hypothetical protein | -- | 5 | |
| DSAG12_03012 | MFS transporter, AAHS family, benzoate transport protein | K05548 | 9 | |
| DSAG12_03013 | heterodisulfide reductase subunit D | K08264 | 6 | |
| DSAG12_03014 | hypothetical protein | -- | 12 | |
| DSAG12_03015 | hypothetical protein | -- | 4 | |
| DSAG12_03016 | Ras-related protein Rab-7L1 | K07916 | 7 | |
| DSAG12_03017 | hypothetical protein | -- | 5 | |
| DSAG12_03018 | hypothetical protein | -- | 0 | |
| DSAG12_03019 | hypothetical protein | -- | 2 | |

| | | | | |
|---|---|---|---|---|
| DSAG12_03020 | hypothetical protein | -- | 4 | |
| DSAG12_03021 | hypothetical protein | -- | 8 | |
| DSAG12_03022 | methylated-DNA-protein-cysteine methyltransferase related protein | K07443 | 7 | |
| DSAG12_03023 | butyryl-CoA dehydrogenase | K00248 | 299 | |
| DSAG12_03024 | tryptophanase | K01667 | 9 | |
| DSAG12_03025 | hypothetical protein | -- | 68 | |
| DSAG12_03026 | hypothetical protein | -- | 108 | |
| DSAG12_03027 | myo-inositol-1-phosphate synthase | K01858 | 192 | |
| DSAG12_03028 | hypothetical protein | -- | 262 | |
| DSAG12_03029 | hypothetical protein | -- | 245 | |
| DSAG12_03030 | hypothetical protein | -- | 205 | |
| DSAG12_03031 | phosphomannomutase | K01840 | 160 | |
| DSAG12_03032 | hypothetical protein | -- | 269 | |
| DSAG12_03033 | hypothetical protein | -- | 52 | |
| DSAG12_03034 | peptide/nickel transport system permease protein | K02034 | 83 | |
| DSAG12_03035 | peptide/nickel transport system permease protein | K02033 | 119 | |
| DSAG12_03036 | hypothetical protein | -- | 251 | |
| DSAG12_03037 | oligopeptide transport system substrate-binding protein | K15580 | 197 | |
| DSAG12_03038 | hypothetical protein | -- | 225 | |
| DSAG12_03039 | hypothetical protein | -- | 209 | |
| DSAG12_03040 | hypothetical protein | -- | 253 | |
| DSAG12_03041 | hypothetical protein | -- | 81 | |
| DSAG12_03042 | oligopeptide transport system ATP-binding protein | K15583 | 123 | |
| DSAG12_03043 | peptide/nickel transport system ATP-binding protein / peptide/nickel transport system ATP-binding protein | K02031 K02 | 107 | |
| DSAG12_03044 | hypothetical protein | -- | 0 | |
| DSAG12_03045 | hypothetical protein | -- | 256 | |
| DSAG12_03046 | quinolinate synthase | K03517 | 115 | |
| DSAG12_03047 | thioredoxin reductase (NADPH) | K00384 | 65 | |
| DSAG12_03048 | formate C-acetyltransferase | K00656 | 177 | |
| DSAG12_03049 | pyruvate formate lyase activating enzyme | K04069 | 232 | |
| DSAG12_03050 | hypothetical protein | -- | 193 | |
| DSAG12_03051 | hypothetical protein | K01163 | 108 | |
| DSAG12_03052 | hypothetical protein | -- | 155 | |
| DSAG12_03053 | K(+)-stimulated pyrophosphate-energized sodium pump | K15987 | 184 | |
| DSAG12_03054 | hypothetical protein | -- | 223 | |
| DSAG12_03055 | hypothetical protein | -- | 73 | |
| DSAG12_03056 | ABC-2 type transport system ATP-binding protein | K01990 | 160 | |
| DSAG12_03057 | hypothetical protein | -- | 175 | |
| DSAG12_03058 | hypothetical protein | -- | 209 | |
| DSAG12_03059 | hypothetical protein | -- | 22 | |
| DSAG12_03060 | hypothetical protein | -- | 118 | |
| DSAG12_03061 | hypothetical protein | -- | 134 | |
| DSAG12_03062 | 8-amino-7-oxononanoate synthase | K00652 | 436 | |
| DSAG12_03063 | hypothetical protein | -- | 231 | |
| DSAG12_03064 | hypothetical protein | -- | 250 | |
| DSAG12_03065 | hypothetical protein | -- | 203 | |
| DSAG12_03066 | | K06944 | 42 | |
| DSAG12_03067 | hypothetical protein | -- | 117 | small GTP-binding domain protein |
| DSAG12_03068 | hypothetical protein | -- | 0 | |
| DSAG12_03069 | hypothetical protein | -- | 0 | |
| DSAG12_03070 | hypothetical protein | -- | 228 | |
| DSAG12_03071 | hypothetical protein | -- | 177 | |
| DSAG12_03072 | hypothetical protein | -- | 41 | |
| DSAG12_03073 | hypothetical protein | -- | 66 | |
| DSAG12_03074 | hypothetical protein | -- | 100 | |
| DSAG12_03075 | hypothetical protein | -- | 275 | |
| DSAG12_03076 | hypothetical protein | -- | 223 | |
| DSAG12_03077 | probable phosphoglycerate mutase | K15634 | 178 | |
| DSAG12_03078 | hypothetical protein | -- | 114 | |
| DSAG12_03079 | tRNA-Ala | | 211 | |
| DSAG12_03080 | NADPH2:quinone reductase | K00344 | 176 | |
| DSAG12_03081 | hypothetical protein | -- | 132 | |
| DSAG12_03082 | hypothetical protein | -- | 184 | small GTP-binding domain protein |
| DSAG12_03083 | hypothetical protein | -- | 198 | |
| DSAG12_03084 | hypothetical protein | -- | 142 | |
| DSAG12_03085 | hypothetical protein | -- | 4 | |
| DSAG12_03086 | hypothetical protein | -- | 319 | |
| DSAG12_03087 | hypothetical protein | -- | 266 | |
| DSAG12_03088 | tRNA-dihydrouridine synthase B | K05540 | 148 | |
| DSAG12_03089 | hypothetical protein | -- | 4647 | |
| DSAG12_03090 | hypothetical protein | -- | 160 | |
| DSAG12_03091 | hypothetical protein | -- | 270 | |
| DSAG12_03092 | hypothetical protein | -- | 135 | |
| DSAG12_03093 | hypothetical protein | -- | 215 | |
| DSAG12_03094 | hypothetical protein | -- | 252 | |
| DSAG12_03095 | hypothetical protein | -- | 362 | |
| DSAG12_03096 | putative adenine-specific DNA-methyltransferase | K07319 | 120 | |

| | | | |
|---|---|---|---|
| DSAG12_03097 | hypothetical protein | -- | 205 |
| DSAG12_03098 | putative adenine-specific DNA-methyltransferase | K07319 | 0 |
| DSAG12_03099 | hypothetical protein | -- | 224 |
| DSAG12_03100 | 1-acyl-sn-glycerol-3-phosphate acyltransferase | K00655 | 224 |
| DSAG12_03101 | hypothetical protein | -- | 228 |
| DSAG12_03102 | hypothetical protein | -- | 158 |
| DSAG12_03103 | | K07052 | 1002 |
| DSAG12_03104 | hypothetical protein | -- | 224 |
| DSAG12_03105 | hypothetical protein | -- | 0 |
| DSAG12_03106 | actin-related protein 2 | K17260 | 195 |
| DSAG12_03107 | hypothetical protein | -- | 408 |
| DSAG12_03108 | hypothetical protein | -- | 143 |
| DSAG12_03109 | hypothetical protein | -- | 138 |
| DSAG12_03110 | hypothetical protein | -- | 197 |
| DSAG12_03111 | hypothetical protein | -- | 165 |
| DSAG12_03112 | hypothetical protein | -- | 121 |
| DSAG12_03113 | hypothetical protein | -- | 109 |
| DSAG12_03114 | diacylglycerol kinase (ATP) | K07029 | 167 |
| DSAG12_03115 | hypothetical protein | -- | 206 |
| DSAG12_03116 | hypothetical protein | -- | 265 |
| DSAG12_03117 | hypothetical protein | -- | 176 |
| DSAG12_03118 | DNA-3-methyladenine glycosylase I | K01246 | 193 |
| DSAG12_03119 | undecaprenyl-diphosphatase | K06153 | 197 |
| DSAG12_03120 | alcohol dehydrogenase, propanol-preferring | K13953 | 0 |
| DSAG12_03121 | hypothetical protein | -- | 203 |
| DSAG12_03122 | cold shock protein (beta-ribbon, CspA family) | K03704 | 160 |
| DSAG12_03123 | hypothetical protein | -- | 170 |
| DSAG12_03124 | hypothetical protein | -- | 0 |
| DSAG12_03125 | hypothetical protein | -- | 202 |
| DSAG12_03126 | hypothetical protein | -- | 188 |
| DSAG12_03127 | peroxiredoxin Q/BCP | K03564 | 191 |
| DSAG12_03128 | transcription initiation factor TFIIB | K03124 | 284 |
| DSAG12_03129 | hypothetical protein | -- | 163 |
| DSAG12_03130 | hypothetical protein | -- | 183 |
| DSAG12_03131 | hypothetical protein | -- | 217 |
| DSAG12_03132 | hypothetical protein | -- | 149 |
| DSAG12_03133 | exonuclease SbcC | K03546 | 197 |
| DSAG12_03134 | hypothetical protein | -- | 0 |
| DSAG12_03135 | hypothetical protein | -- | 180 |
| DSAG12_03136 | hypothetical protein | -- | 120 |
| DSAG12_03137 | hypothetical protein | -- | 0 |
| DSAG12_03138 | hypothetical protein | -- | 209 |
| DSAG12_03139 | hypothetical protein | -- | 139 |
| DSAG12_03140 | hypothetical protein | -- | 229 |
| DSAG12_03141 | hypothetical protein | -- | 135 |
| DSAG12_03142 | hypothetical protein | -- | 184 |
| DSAG12_03143 | ribonuclease P protein subunit RPR2 | K03540 | 209 |
| DSAG12_03144 | rRNA small subunit pseudouridine methyltransferase Nep1 | K14568 | 188 |
| DSAG12_03145 | hypothetical protein | -- | 156 |
| DSAG12_03146 | hypothetical protein | -- | 197 |
| DSAG12_03147 | hypothetical protein | -- | 209 |
| DSAG12_03148 | hypothetical protein | -- | 238 |
| DSAG12_03149 | hypothetical protein | -- | 186 |
| DSAG12_03150 | nitric oxide reductase NorQ protein | K04748 | 225 |
| DSAG12_03151 | ABC-2 type transport system ATP-binding protein | K01990 | 194 |
| DSAG12_03152 | ABC-2 type transport system permease protein | K01992 | 54 |
| DSAG12_03153 | hypothetical protein | -- | 9 |
| DSAG12_03154 | hypothetical protein | -- | 0 |
| DSAG12_03155 | hypothetical protein | -- | 182 |
| DSAG12_03156 | hypothetical protein | -- | 159 |
| DSAG12_03157 | hypothetical protein | -- | 257 |
| DSAG12_03158 | tRNA-Gly | | 0 |
| DSAG12_03159 | glycoside/pentoside/hexuronide:cation symporter, GPH family | K03292 | 297 |
| DSAG12_03160 | hypothetical protein | -- | 40 |
| DSAG12_03161 | heterodisulfide reductase subunit C | K03390 | 52 |
| DSAG12_03162 | | K05912 | 171 |
| DSAG12_03163 | heterodisulfide reductase subunit A | K03388 | 201 |
| DSAG12_03164 | anaerobic sulfite reductase subunit A | K16950 | 162 |
| DSAG12_03165 | sulfhydrogenase subunit gamma (sulfur reductase) | K17995 | 119 |
| DSAG12_03166 | hypothetical protein | -- | 186 |
| DSAG12_03167 | hypothetical protein | -- | 190 |
| DSAG12_03168 | hypothetical protein | -- | 124 |
| DSAG12_03169 | hypothetical protein | -- | 182 |
| DSAG12_03170 | hypothetical protein | -- | 225 |
| DSAG12_03171 | tRNA-Phe | | 169 |
| DSAG12_03172 | isovaleryl-CoA dehydrogenase | K00253 | 205 |
| DSAG12_03173 | hypothetical protein | -- | 227 |
| DSAG12_03174 | hypothetical protein | -- | 101 |

| | | | |
|---|---|---|---|
| DSAG12_03175 | MFS transporter, DHA3 family, macrolide efflux protein | K08217 | 213 |
| DSAG12_03176 | hypothetical protein | -- | 151 |
| DSAG12_03177 | hypothetical protein | -- | 157 |
| DSAG12_03178 | bifunctional enzyme Fae/Hps | K13812 | 214 |
| DSAG12_03179 | hypothetical protein | -- | 195 |
| DSAG12_03180 | hypothetical protein | -- | 62 |
| DSAG12_03181 | fanconi anemia group M protein | K10896 | 171 |
| DSAG12_03182 | fanconi anemia group M protein | K10896 | 265 |
| DSAG12_03183 | 8-oxo-dGTP diphosphatase | K03574 | 1044 |
| DSAG12_03184 | hypothetical protein | -- | 180 |
| DSAG12_03185 | hypothetical protein | -- | 150 |
| DSAG12_03186 | 5'-nucleotidase | K01081 | 186 |
| DSAG12_03187 | 5'-nucleotidase | K01081 | 197 |
| DSAG12_03188 | methylenetetrahydrofolate dehydrogenase (NADP+) / methenyltetrahydrofolate cyclohydrolase | K01491 | 178 |
| DSAG12_03189 | amidophosphoribosyltransferase | K00764 | 191 |
| DSAG12_03190 | hypothetical protein | K09126 | 157 |
| DSAG12_03191 | chloride channel protein, ClC family | K03281 | 137 |
| DSAG12_03192 | hypothetical protein | -- | 109 |
| DSAG12_03193 | hypothetical protein | -- | 187 |
| DSAG12_03194 | hypothetical protein | -- | 151 |
| DSAG12_03195 | hypothetical protein | -- | 148 |
| DSAG12_03196 | butyryl-CoA dehydrogenase | K00248 | 229 |
| DSAG12_03197 | glycolate oxidase | K00104 | 206 |
| DSAG12_03198 | hypothetical protein | -- | 189 |
| DSAG12_03199 | excinuclease ABC subunit B | K03702 | 133 |
| DSAG12_03200 | excinuclease ABC subunit A | K03701 | 149 |
| DSAG12_03201 | hypothetical protein | -- | 147 |
| DSAG12_03202 | hypothetical protein | -- | 141 |
| DSAG12_03203 | 3'-5' exoribonuclease | K03698 | 271 |
| DSAG12_03204 | hypothetical protein | -- | 188 |
| DSAG12_03205 | hypothetical protein | -- | 5 |
| DSAG12_03206 | hypothetical protein | -- | 9 |
| DSAG12_03207 | 3-oxo-5-alpha-steroid 4-dehydrogenase 1 | K12343 | 5 |
| DSAG12_03208 | hypothetical protein | -- | 222 |
| DSAG12_03209 | long-chain acyl-CoA synthetase | K01897 | 196 |
| DSAG12_03210 | hypothetical protein | -- | 172 |
| DSAG12_03211 | | K01362 | 92 |
| DSAG12_03212 | hypothetical protein | -- | 198 |
| DSAG12_03213 | hypothetical protein | -- | 171 |
| DSAG12_03214 | | K06940 | 0 |
| DSAG12_03215 | cysteinyl-tRNA synthetase | K01883 | 219 |
| DSAG12_03216 | hypothetical protein | -- | 233 |
| DSAG12_03217 | tRNA nucleotidyltransferase (CCA-adding enzyme) | K07558 | 227 |
| DSAG12_03218 | hypothetical protein | -- | 214 |
| DSAG12_03219 | elongation factor 2 | K03234 | 273 |
| DSAG12_03220 | hypothetical protein | -- | 174 |
| DSAG12_03221 | hypothetical protein | -- | 228 |
| DSAG12_03222 | | K06936 | 213 |
| DSAG12_03223 | archaeosine synthase | K07557 | 334 |
| DSAG12_03224 | hypothetical protein | -- | 230 |
| DSAG12_03225 | arsenite-transporting ATPase | K01551 | 181 |
| DSAG12_03226 | ATP-dependent Clp protease ATP-binding subunit ClpB | K03695 | 195 |
| DSAG12_03227 | hypothetical protein | -- | 0 |
| DSAG12_03228 | putative copper resistance protein D | K07245 | 0 |
| DSAG12_03229 | DNA primase | K02316 | 277 |
| DSAG12_03230 | hypothetical protein | -- | 0 |
| DSAG12_03231 | fused signal recognition particle receptor | K03110 | 290 |
| DSAG12_03232 | prefoldin alpha subunit | K04797 | 196 |
| DSAG12_03233 | large subunit ribosomal protein LX | K02944 | 213 |
| DSAG12_03234 | translation initiation factor 6 | K03264 | 150 |
| DSAG12_03235 | hypothetical protein | -- | 434 |
| DSAG12_03236 | programmed cell death protein 5 | K06875 | 536 |
| DSAG12_03237 | small subunit ribosomal protein S19e | K02966 | 193 |
| DSAG12_03238 | methionyl-tRNA synthetase | K01874 | 130 |
| DSAG12_03239 | diacylglycerol kinase (ATP) | K07029 | 0 |
| DSAG12_03240 | hypothetical protein | -- | 194 |
| DSAG12_03241 | hypothetical protein | -- | 141 |
| DSAG12_03242 | hypothetical protein | -- | 124 |
| DSAG12_03243 | macrolide transport system ATP-binding/permease protein | K05685 | 201 |
| DSAG12_03244 | hypothetical protein | -- | 300 |
| DSAG12_03245 | archaea-specific helicase | K03725 | 179 |
| DSAG12_03246 | glycoside/pentoside/hexuronide:cation symporter, GPH family | K03292 | 116 |
| DSAG12_03247 | hypothetical protein | -- | 165 |
| DSAG12_03248 | hypothetical protein | -- | 212 |
| DSAG12_03249 | hypothetical protein | -- | 159 |
| DSAG12_03250 | hypothetical protein | -- | 220 |
| DSAG12_03251 | hypothetical protein | -- | 194 |
| DSAG12_03252 | hypothetical protein | -- | 157 |

| | | | |
|---|---|---|---|
| DSAG12_03253 | MarR family transcriptional regulator, transcriptional regulator for hemolysin | K06075 | 147 |
| DSAG12_03254 | MFS transporter, DHA3 family, macrolide efflux protein | K08217 | 225 |
| DSAG12_03255 | | K00358 | 355 |
| DSAG12_03256 | dihydrolipoamide dehydrogenase | K00382 | 191 |
| DSAG12_03257 | hypothetical protein | -- | 104 |
| DSAG12_03258 | maltose/moltooligosaccharide transporter | K16211 | 244 |
| DSAG12_03259 | hypothetical protein | -- | 238 |
| DSAG12_03260 | hypothetical protein | -- | 24 |
| DSAG12_03261 | 3-oxoacyl-[acyl-carrier protein] reductase | K00059 | 242 |
| DSAG12_03262 | hypothetical protein | -- | 201 |
| DSAG12_03263 | hypothetical protein | -- | 159 |
| DSAG12_03264 | hypothetical protein | -- | 210 |
| DSAG12_03265 | 2-dehydro-3-deoxyphosphogluconate aldolase / (4S)-4-hydroxy-2-oxoglutarate aldolase | K01625 | 183 |
| DSAG12_03266 | 2-dehydro-3-deoxygluconokinase | K00874 | 218 |
| DSAG12_03267 | hypothetical protein | -- | 186 |
| DSAG12_03268 | hypothetical protein | -- | 232 |
| DSAG12_03269 | hypothetical protein | -- | 192 |
| DSAG12_03270 | NADH-quinone oxidoreductase subunit E | K00334 | 180 |
| DSAG12_03271 | NADH-quinone oxidoreductase subunit F | K00335 | 171 |
| DSAG12_03272 | formate dehydrogenase major subunit | K00123 | 277 |
| DSAG12_03273 | hypothetical protein | -- | 120 |
| DSAG12_03274 | energy-coupling factor transport system ATP-binding protein | K16786 | 256 |
| DSAG12_03275 | energy-coupling factor transport system permease protein | K16785 | 166 |
| DSAG12_03276 | hypothetical protein | -- | 179 |
| DSAG12_03277 | hypothetical protein | -- | 226 |
| DSAG12_03278 | ribonuclease Z | K00784 | 268 |
| DSAG12_03279 | hypothetical protein | -- | 203 |
| DSAG12_03280 | hypothetical protein | -- | 165 |
| DSAG12_03281 | serine/threonine-protein phosphatase 5 | K04460 | 232 |
| DSAG12_03282 | | K07022 | 232 |
| DSAG12_03283 | hypothetical protein | -- | 85 |
| DSAG12_03284 | zinc transporter, ZIP family | K07238 | 200 |
| DSAG12_03285 | alpha-mannosidase | K01191 | 389 |
| DSAG12_03286 | hypothetical protein | -- | 144 |
| DSAG12_03287 | hypothetical protein | -- | 134 |
| DSAG12_03288 | serine/threonine-protein phosphatase 5 | K04460 | 251 |
| DSAG12_03289 | indolepyruvate ferredoxin oxidoreductase, beta subunit | K00180 | 75 |
| DSAG12_03290 | indolepyruvate ferredoxin oxidoreductase, alpha subunit | K00179 | 178 |
| DSAG12_03291 | cytochrome c-type biogenesis protein | K06196 | 142 |
| DSAG12_03292 | hypothetical protein | -- | 66 |
| DSAG12_03293 | hypothetical protein | -- | 220 |
| DSAG12_03294 | hypothetical protein | -- | 158 |
| DSAG12_03295 | pimeloyl-[acyl-carrier protein] methyl ester esterase | K02170 | 159 |
| DSAG12_03296 | hypothetical protein | -- | 86 |
| DSAG12_03297 | hypothetical protein | -- | 128 |
| DSAG12_03298 | pimeloyl-[acyl-carrier protein] methyl ester esterase | K02170 | 0 |
| DSAG12_03299 | alpha-N-acetylglucosaminidase | K01205 | 240 |
| DSAG12_03300 | molybdopterin synthase sulfur carrier subunit | K03636 | 259 |
| DSAG12_03301 | hypothetical protein | -- | 21 |
| DSAG12_03302 | all-trans-retinol 13,14-reductase | K09516 | 87 |
| DSAG12_03303 | hypothetical protein | -- | 215 |
| DSAG12_03304 | hypothetical protein | -- | 215 |
| DSAG12_03305 | hypothetical protein | -- | 192 |
| DSAG12_03306 | hypothetical protein | -- | 185 |
| DSAG12_03307 | hypothetical protein | -- | 168 |
| DSAG12_03308 | hypothetical protein | -- | 122 |
| DSAG12_03309 | hypothetical protein | -- | 0 |
| DSAG12_03310 | hypothetical protein | -- | 111 |
| DSAG12_03311 | hypothetical protein | -- | 116 |
| DSAG12_03312 | hypothetical protein | -- | 193 |
| DSAG12_03313 | tRNA-Arg | Intron(37429 | 158 |
| DSAG12_03314 | demethylmenaquinone methyltransferase / 2-methoxy-6-polyprenyl-1,4-benzoquinol methylase | K03183 | 151 |
| DSAG12_03315 | hypothetical protein | -- | 248 |
| DSAG12_03316 | hypothetical protein | -- | 145 |
| DSAG12_03317 | translation initiation factor 5B | K03243 | 244 |
| DSAG12_03318 | hypothetical protein | -- | 153 |
| DSAG12_03319 | hypothetical protein | -- | 216 |
| DSAG12_03320 | small subunit ribosomal protein S6e | K02991 | 235 |
| DSAG12_03321 | hypothetical protein | -- | 209 |
| DSAG12_03322 | hypothetical protein | -- | 174 |
| DSAG12_03323 | hypothetical protein | -- | 198 |
| DSAG12_03324 | small subunit ribosomal protein S28e | K02979 | 144 |
| DSAG12_03325 | large subunit ribosomal protein L7Ae | K02936 | 231 |
| DSAG12_03326 | hypothetical protein | -- | 197 |
| DSAG12_03327 | hypothetical protein | -- | 215 |
| DSAG12_03328 | tRNA-His | Intron(37590 | 266 |
| DSAG12_03329 | hypothetical protein | -- | 209 |
| DSAG12_03330 | hypothetical protein | -- | 109 |

| | | | |
|---|---|---|---|
| DSAG12_03331 | hypothetical protein | -- | 166 |
| DSAG12_03332 | hypothetical protein | -- | 231 |
| DSAG12_03333 | hypothetical protein | -- | 212 |
| DSAG12_03334 | histidyl-tRNA synthetase | K01892 | 175 |
| DSAG12_03335 | hypothetical protein | -- | 176 |
| DSAG12_03336 | hypothetical protein | -- | 149 |
| DSAG12_03337 | cold shock protein (beta-ribbon, CspA family) | K03704 | 146 |
| DSAG12_03338 | | K01112 | 54 |
| DSAG12_03339 | bilin biosynthesis protein | K05384 | 230 |
| DSAG12_03340 | bifunctional tRNA threonylcarbamoyladenosine biosynthesis protein | K15904 | 53 |
| DSAG12_03341 | small subunit ribosomal protein S27Ae | K02977 | 207 |
| DSAG12_03342 | small subunit ribosomal protein S24e | K02974 | 156 |
| DSAG12_03343 | hypothetical protein | K09735 | 187 |
| DSAG12_03344 | hypothetical protein | -- | 185 |
| DSAG12_03345 | hypothetical protein | -- | 201 |
| DSAG12_03346 | hypothetical protein | -- | 225 |
| DSAG12_03347 | hypothetical protein | -- | 172 |
| DSAG12_03348 | enterochelin esterase and related enzymes | K07214 | 50 |
| DSAG12_03349 | DNA-directed RNA polymerase subunit E' | K03049 | 203 |
| DSAG12_03350 | hypothetical protein | -- | 213 |
| DSAG12_03351 | medium-chain acyl-[acyl-carrier-protein] hydrolase | K01071 | 245 |
| DSAG12_03352 | hypothetical protein | -- | 178 |
| DSAG12_03353 | hypothetical protein | -- | 221 |
| DSAG12_03354 | hypothetical protein | -- | 102 |
| DSAG12_03355 | hypothetical protein | -- | 204 |
| DSAG12_03356 | hypothetical protein | -- | 160 |
| DSAG12_03357 | hypothetical protein | -- | 36 |
| DSAG12_03358 | 3-oxoacyl-[acyl-carrier protein] reductase | K00059 | 182 |
| DSAG12_03359 | 3-oxoacyl-[acyl-carrier protein] reductase | K00059 | 188 |
| DSAG12_03360 | hypothetical protein | -- | 55 |
| DSAG12_03361 | glycerophosphoinositol inositolphosphodiesterase | K01124 | 246 |
| DSAG12_03362 | 3-oxoacyl-[acyl-carrier protein] reductase | K00059 | 195 |
| DSAG12_03363 | 3-oxoacyl-[acyl-carrier protein] reductase | K00059 | 139 |
| DSAG12_03364 | hypothetical protein | -- | 168 |
| DSAG12_03365 | | K00540 | 47 |
| DSAG12_03366 | | K07095 | 173 |
| DSAG12_03367 | hypothetical protein | -- | 178 |
| DSAG12_03368 | hypothetical protein | -- | 122 |
| DSAG12_03369 | hypothetical protein | -- | 161 |
| DSAG12_03370 | LL-diaminopimelate aminotransferase | K10206 | 151 |
| DSAG12_03371 | electron transfer flavoprotein alpha subunit | K03522 | 140 |
| DSAG12_03372 | electron transfer flavoprotein beta subunit | K03521 | 182 |
| DSAG12_03373 | hypothetical protein | -- | 159 |
| DSAG12_03374 | acyl-CoA dehydrogenase | K00249 | 192 |
| DSAG12_03375 | ATP-binding protein involved in chromosome partitioning | K03593 | 261 |
| DSAG12_03376 | | K01529 | 239 |
| DSAG12_03377 | | K07076 | 144 |
| DSAG12_03378 | hypothetical protein | -- | 86 |
| DSAG12_03379 | indolepyruvate ferredoxin oxidoreductase, alpha subunit | K00179 | 233 |
| DSAG12_03380 | indolepyruvate ferredoxin oxidoreductase, beta subunit | K00180 | 0 |
| DSAG12_03381 | Lrp/AsnC family transcriptional regulator, regulator for asnA, asnC and gidA | K03718 | 192 |
| DSAG12_03382 | | K07076 | 125 |
| DSAG12_03383 | hypothetical protein | -- | 227 |
| DSAG12_03384 | hypothetical protein | -- | 107 |
| DSAG12_03385 | hypothetical protein | -- | 184 |
| DSAG12_03386 | hypothetical protein | -- | 165 |
| DSAG12_03387 | anaerobic nitric oxide reductase flavorubredoxin | K12264 | 187 |
| DSAG12_03388 | phosphoglycolate phosphatase | K01091 | 107 |
| DSAG12_03389 | hypothetical protein | -- | 151 |
| DSAG12_03390 | heterodisulfide reductase subunit D | K08264 | 186 |
| DSAG12_03391 | hypothetical protein | -- | 160 |
| DSAG12_03392 | hypothetical protein | -- | 205 |
| DSAG12_03393 | phosphoglycolate phosphatase | K01091 | 231 |
| DSAG12_03394 | hypothetical protein | -- | 181 |
| DSAG12_03395 | hypothetical protein | -- | 182 |
| DSAG12_03396 | hypothetical protein | -- | 103 |
| DSAG12_03397 | adenine-specific DNA-methyltransferase | K07317 | 211 |
| DSAG12_03398 | hypothetical protein | -- | 251 |
| DSAG12_03399 | hypothetical protein | -- | 221 |
| DSAG12_03400 | hypothetical protein | -- | 164 |
| DSAG12_03401 | digeranylgeranylglycerophospholipid reductase | K17830 | 169 |
| DSAG12_03402 | hypothetical protein | -- | 181 |
| DSAG12_03403 | superkiller protein 3 | K12600 | 88 |
| DSAG12_03404 | hypothetical protein | -- | 257 |
| DSAG12_03405 | thioredoxin 1 | K03671 | 151 |
| DSAG12_03406 | oligopeptide transport system ATP-binding protein | K10823 | 189 |
| DSAG12_03407 | peptide/nickel transport system ATP-binding protein | K02031 | 181 |
| DSAG12_03408 | oligopeptide transport system permease protein | K15582 | 194 |

| | | | |
|---|---|---|---|
| DSAG12_03409 | peptide/nickel transport system permease protein | K02033 | 0 |
| DSAG12_03410 | hypothetical protein | -- | 143 |
| DSAG12_03411 | peptide/nickel transport system substrate-binding protein | K02035 | 106 |
| DSAG12_03412 | hypothetical protein | -- | 249 |
| DSAG12_03413 | hypothetical protein | -- | 208 |
| DSAG12_03414 | hypothetical protein | -- | 0 |
| DSAG12_03415 | 26S proteasome regulatory subunit N11 | K03030 | 113 |
| DSAG12_03416 | hypothetical protein | -- | 323 |
| DSAG12_03417 | transcription initiation factor TFIIB | K03124 | 216 |
| DSAG12_03418 | hypothetical protein | -- | 214 |
| DSAG12_03419 | hypothetical protein | -- | 243 |
| DSAG12_03420 | internalin A | K13730 | 437 |
| DSAG12_03421 | | K07131 | 178 |
| DSAG12_03422 | hypothetical protein | -- | 201 |
| DSAG12_03423 | hypothetical protein | -- | 283 |
| DSAG12_03424 | lysophospholipase | K01048 | 572 |
| DSAG12_03425 | hypothetical protein | -- | 2408 |
| DSAG12_03426 | ribonuclease Z | K00784 | 218 |
| DSAG12_03427 | hypothetical protein | -- | 165 |
| DSAG12_03428 | hypothetical protein | -- | 179 |
| DSAG12_03429 | hypothetical protein | -- | 183 |
| DSAG12_03430 | ribosomal-protein-alanine N-acetyltransferase | K03790 | 145 |
| DSAG12_03431 | hypothetical protein | -- | 209 |
| DSAG12_03432 | glutamine synthetase | K01915 | 292 |
| DSAG12_03433 | hypothetical protein | -- | 226 |
| DSAG12_03434 | CTP synthase | K01937 | 172 |
| DSAG12_03435 | magnesium chelatase subunit D | K03404 | 366 |
| DSAG12_03436 | hypothetical protein | -- | 98 |
| DSAG12_03437 | hypothetical protein | -- | 209 |
| DSAG12_03438 | hypothetical protein | -- | 150 |
| DSAG12_03439 | hypothetical protein | -- | 131 |
| DSAG12_03440 | L-ribulose-5-phosphate 4-epimerase | K01786 | 258 |
| DSAG12_03441 | segregation and condensation protein B | K06024 | 200 |
| DSAG12_03442 | acylglycerol lipase | K01054 | 156 |
| DSAG12_03443 | | K01453 | 81 |
| DSAG12_03444 | hypothetical protein | -- | 11 |
| DSAG12_03445 | chromosome segregation protein | K03529 | 188 |
| DSAG12_03446 | hypothetical protein | -- | 126 |
| DSAG12_03447 | | K07024 | 0 |
| DSAG12_03448 | Trp repressor binding protein | K03809 | 313 |
| DSAG12_03449 | chorismate synthase | K01736 | 215 |
| DSAG12_03450 | ribose 1,5-bisphosphokinase | K05774 | 159 |
| DSAG12_03451 | hypothetical protein | -- | 218 |
| DSAG12_03452 | hypothetical protein | -- | 207 |
| DSAG12_03453 | hypothetical protein | -- | 119 |
| DSAG12_03454 | hypothetical protein | -- | 134 |
| DSAG12_03455 | hypothetical protein | -- | 180 |
| DSAG12_03456 | peptide/nickel transport system substrate-binding protein | K02035 | 55 |
| DSAG12_03457 | molecular chaperone DnaK | K04043 | 159 |
| DSAG12_03458 | molecular chaperone DnaJ | K03686 | 139 |
| DSAG12_03459 | hypothetical protein | -- | 163 |
| DSAG12_03460 | hypothetical protein | -- | 236 |
| DSAG12_03461 | tRNA-Gly | | 220 |
| DSAG12_03462 | glycerophosphoryl diester phosphodiesterase | K01126 | 180 |
| DSAG12_03463 | uncharacterized protein | K06950 | 232 |
| DSAG12_03464 | hypothetical protein | -- | 185 |
| DSAG12_03465 | Rab family, other | K07976 | 164 |
| DSAG12_03466 | ribonuclease J | K12574 | 175 |
| DSAG12_03467 | transcription initiation factor TFIID TATA-box-binding protein | K03120 | 127 |
| DSAG12_03468 | hypothetical protein | -- | 214 |
| DSAG12_03469 | ribosomal-protein-alanine N-acetyltransferase | K03790 | 190 |
| DSAG12_03470 | phosphatidylserine decarboxylase | K01613 | 226 |
| DSAG12_03471 | hypothetical protein | -- | 210 |
| DSAG12_03472 | hypothetical protein | -- | 104 |
| DSAG12_03473 | hypothetical protein | -- | 570 |
| DSAG12_03474 | hypothetical protein | -- | 133 |
| DSAG12_03475 | hypothetical protein | -- | 90 |
| DSAG12_03476 | hydrogenase expression/formation protein HypE | K04655 | 248 |
| DSAG12_03477 | esterase / lipase | K01066 | 213 |
| DSAG12_03478 | fumarate hydratase, class I | K01676 | 199 |
| DSAG12_03479 | fumarate hydratase subunit alpha | K01677 | 183 |
| DSAG12_03480 | hypothetical protein | -- | 231 |
| DSAG12_03481 | molybdopterin-guanine dinucleotide biosynthesis protein B | K03753 | 25 |
| DSAG12_03482 | hypothetical protein | -- | 157 |
| DSAG12_03483 | hypothetical protein | -- | 167 |
| DSAG12_03484 | hypothetical protein | -- | 49 |
| DSAG12_03485 | hypothetical protein | -- | 161 |
| DSAG12_03486 | hypothetical protein | -- | 178 |

| | | | |
|---|---|---|---|
| DSAG12_03487 | hypothetical protein | -- | 171 |
| DSAG12_03488 | putative transposase | K07491 | 173 |
| DSAG12_03489 | hypothetical protein | -- | 16 |
| DSAG12_03490 | hypothetical protein | -- | 177 |
| DSAG12_03491 | hypothetical protein | -- | 121 |
| DSAG12_03492 | hypothetical protein | -- | 157 |
| DSAG12_03493 | hypothetical protein | -- | 191 |
| DSAG12_03494 | hypothetical protein | -- | 205 |
| DSAG12_03495 | hypothetical protein | -- | 173 |
| DSAG12_03496 | hypothetical protein | -- | 217 |
| DSAG12_03497 | | K07102 | 232 |
| DSAG12_03498 | hypothetical protein | -- | 189 |
| DSAG12_03499 | hypothetical protein | -- | 150 |
| DSAG12_03500 | hypothetical protein | -- | 218 |
| DSAG12_03501 | fanconi anemia group M protein | K10896 | 198 |
| DSAG12_03502 | hypothetical protein | -- | 187 |
| DSAG12_03503 | hypothetical protein | -- | 156 |
| DSAG12_03504 | hypothetical protein | -- | 148 |
| DSAG12_03505 | hypothetical protein | -- | 218 |
| DSAG12_03506 | hypothetical protein | -- | 179 |
| DSAG12_03507 | fanconi anemia group M protein | K10896 | 213 |
| DSAG12_03508 | hypothetical protein | -- | 184 |
| DSAG12_03509 | hypothetical protein | -- | 178 |
| DSAG12_03510 | hypothetical protein | -- | 135 |
| DSAG12_03511 | hypothetical protein | -- | 214 |
| DSAG12_03512 | hypothetical protein | -- | 180 |
| DSAG12_03513 | hypothetical protein | -- | 180 |
| DSAG12_03514 | hypothetical protein | -- | 183 |
| DSAG12_03515 | hypothetical protein | -- | 75 |
| DSAG12_03516 | | K06883 | 146 |
| DSAG12_03517 | hypothetical protein | -- | 169 |
| DSAG12_03518 | hypothetical protein | -- | 244 |
| DSAG12_03519 | hypothetical protein | -- | 641 |
| DSAG12_03520 | hypothetical protein | -- | 341 |
| DSAG12_03521 | hypothetical protein | -- | 111 |
| DSAG12_03522 | hypothetical protein | -- | 196 |
| DSAG12_03523 | hypothetical protein | -- | 0 |
| DSAG12_03524 | hypothetical protein | -- | 183 |
| DSAG12_03525 | leucine-rich repeat kinase 2 | K08844 | 204 |
| DSAG12_03526 | flap endonuclease-1 | K04799 | 212 |
| DSAG12_03527 | hypothetical protein | -- | 162 |
| DSAG12_03528 | hypothetical protein | -- | 203 |
| DSAG12_03529 | hypothetical protein | -- | 275 |
| DSAG12_03530 | DNA repair protein RadA | K04483 | 159 |
| DSAG12_03531 | hypothetical protein | -- | 150 |
| DSAG12_03532 | hypothetical protein | -- | 207 |
| DSAG12_03533 | hypothetical protein | -- | 166 |
| DSAG12_03534 | hypothetical protein | -- | 131 |
| DSAG12_03535 | hypothetical protein | -- | 148 |
| DSAG12_03536 | hypothetical protein | -- | 213 |
| DSAG12_03537 | hypothetical protein | -- | 196 |
| DSAG12_03538 | helicase | K03726 | 193 |
| DSAG12_03539 | hypothetical protein | -- | 203 |
| DSAG12_03540 | hypothetical protein | -- | 210 |
| DSAG12_03541 | hypothetical protein | -- | 217 |
| DSAG12_03542 | hypothetical protein | -- | 148 |
| DSAG12_03543 | helicase | K03726 | 117 |
| DSAG12_03544 | hypothetical protein | -- | 213 |
| DSAG12_03545 | hypothetical protein | -- | 0 |
| DSAG12_03546 | fanconi anemia group M protein | K10896 | 0 |
| DSAG12_03547 | hypothetical protein | -- | 0 |
| DSAG12_03548 | hypothetical protein | -- | 0 |
| DSAG12_03549 | hypothetical protein | -- | 0 |
| DSAG12_03550 | mTERF domain-containing protein, mitochondrial | K15032 | 0 |
| DSAG12_03551 | hypothetical protein | -- | 0 |
| DSAG12_03552 | hypothetical protein | -- | 0 |
| DSAG12_03553 | hypothetical protein | -- | 0 |
| DSAG12_03554 | hypothetical protein | -- | 0 |
| DSAG12_03555 | hypothetical protein | -- | 0 |
| DSAG12_03556 | hypothetical protein | -- | 0 |
| DSAG12_03557 | hypothetical protein | -- | 0 |
| DSAG12_03558 | hypothetical protein | -- | 0 |
| DSAG12_03559 | hypothetical protein | -- | 0 |
| DSAG12_03560 | small subunit ribosomal protein S26e | K02976 | 0 |
| DSAG12_03561 | archaetidylinositol phosphate synthase | K17884 | 0 |
| DSAG12_03562 | hypothetical protein | -- | 0 |
| DSAG12_03563 | hypothetical protein | -- | 0 |
| DSAG12_03564 | hypothetical protein | -- | 0 |

| | | | | |
|---|---|---|---|---|
| DSAG12_03565 | hypothetical protein | -- | 0 | |
| DSAG12_03566 | hypothetical protein | -- | 0 | |
| DSAG12_03567 | transcription initiation factor TFIIB | K03124 | 0 | |
| DSAG12_03568 | RNA-binding protein | K07569 | 0 | |
| DSAG12_03569 | GTP-binding protein | K03978 | 0 | |
| DSAG12_03570 | hypothetical protein | -- | 0 | |
| DSAG12_03571 | phosphoglycolate phosphatase | K01091 | 0 | |
| DSAG12_03572 | threonyl-tRNA synthetase | K01868 | 0 | |
| DSAG12_03573 | tRNA wybutosine-synthesizing protein 1 | K15449 | 0 | |
| DSAG12_03574 | hypothetical protein | -- | 0 | |
| DSAG12_03575 | hypothetical protein | -- | 0 | |
| DSAG12_03576 | hypothetical protein | -- | 0 | |
| DSAG12_03577 | hypothetical protein | -- | 6 | |
| DSAG12_03578 | hypothetical protein | -- | 0 | |
| DSAG12_03579 | hypothetical protein | -- | 0 | |
| DSAG12_03580 | hypothetical protein | -- | 0 | |
| DSAG12_03581 | hypothetical protein | -- | 0 | |
| DSAG12_03582 | putative ABC transport system ATP-binding protein | K02003 | 0 | |
| DSAG12_03583 | hypothetical protein | -- | 0 | |
| DSAG12_03584 | hypothetical protein | -- | 0 | |
| DSAG12_03585 | hypothetical protein | -- | 0 | |
| DSAG12_03586 | hypothetical protein | -- | 0 | |
| DSAG12_03587 | putative ABC transport system ATP-binding protein | K02003 | 0 | |
| DSAG12_03588 | deoxyribonuclease IV | K01151 | 0 | |
| DSAG12_03589 | hypothetical protein | -- | 0 | |
| DSAG12_03590 | hypothetical protein | -- | 0 | |
| DSAG12_03591 | 5-formyltetrahydrofolate cyclo-ligase | K01934 | 0 | |
| DSAG12_03592 | phosphoribosylaminoimidazolecarboxamide formyltransferase / IMP cyclohydrolase | K00602 | 0 | |
| DSAG12_03593 | hypothetical protein | -- | 0 | |
| DSAG12_03594 | IMP dehydrogenase | K00088 | 0 | |
| DSAG12_03595 | Ras-related protein Rab-1A | K07874 | 0 | |
| DSAG12_03596 | uncharacterized protein | K06950 | 0 | |
| DSAG12_03597 | hypothetical protein | -- | 0 | |
| DSAG12_03598 | adenylyltransferase and sulfurtransferase | K11996 | 0 | |
| DSAG12_03599 | sulfite reductase (ferredoxin) | K00392 | 0 | |
| DSAG12_03600 | methanogen homoaconitase small subunit | K16793 | 0 | |
| DSAG12_03601 | methanogen homoaconitase large subunit | K16792 | 0 | |
| DSAG12_03602 | adenylate kinase | K00939 | 0 | |
| DSAG12_03603 | hypothetical protein | -- | 0 | |
| DSAG12_03604 | branched-chain amino acid aminotransferase | K00826 | 0 | |
| DSAG12_03605 | branched-chain amino acid aminotransferase | K00826 | 0 | |
| DSAG12_03606 | transposase | K07486 | 0 | |
| DSAG12_03607 | hypothetical protein | -- | 0 | |
| DSAG12_03608 | hypothetical protein | -- | 0 | |
| DSAG12_03609 | | K00936 | 0 | |
| DSAG12_03610 | hypothetical protein | -- | 0 | |
| DSAG12_03611 | hypothetical protein | -- | 0 | |
| DSAG12_03612 | hypothetical protein | -- | 0 | |
| DSAG12_03613 | acylphosphatase | K01512 | 0 | |
| DSAG12_03614 | hypothetical protein | -- | 0 | |
| DSAG12_03615 | pantetheine-phosphate adenylyltransferase | K02201 | 0 | |
| DSAG12_03616 | phenylalanyl-tRNA synthetase alpha chain | K01889 | 0 | |
| DSAG12_03617 | phenylalanyl-tRNA synthetase beta chain | K01890 | 0 | |
| DSAG12_03618 | hypothetical protein | -- | 0 | |
| DSAG12_03619 | ribonuclease PH | K00989 | 0 | |
| DSAG12_03620 | chorismate mutase / prephenate dehydratase | K14170 | 0 | |
| DSAG12_03621 | hypothetical protein | -- | 0 | |
| DSAG12_03622 | Ca2+-transporting ATPase | K01537 | 0 | |
| DSAG12_03623 | hypothetical protein | -- | 0 | |
| DSAG12_03624 | hypothetical protein | -- | 0 | |
| DSAG12_03625 | ribonuclease III | K03685 | 0 | |
| DSAG12_03626 | demethylmenaquinone methyltransferase / 2-methoxy-6-polyprenyl-1,4-benzoquinol methylase | K03183 | 125 | |
| DSAG12_03627 | hypothetical protein | -- | 209 | |
| DSAG12_03628 | aminoglycoside 2''-phosphotransferase | K17910 | 188 | |
| DSAG12_03629 | ESCRT-II complex subunit VPS22 | K12188 | 151 | |
| DSAG12_03630 | ESCRT-II complex subunit VPS25 | K12189 | 82 | |
| DSAG12_03631 | vacuolar protein-sorting-associated protein 4 | K12196 | 150 | |
| DSAG12_03632 | hypothetical protein | -- | 127 | |
| DSAG12_03633 | hypothetical protein | -- | 226 | |
| DSAG12_03634 | hypothetical protein | -- | 150 | |
| DSAG12_03635 | hypothetical protein | -- | 142 | small GTP-binding domain protein |
| DSAG12_03636 | hypothetical protein | -- | 191 | |
| DSAG12_03637 | hypothetical protein | -- | 216 | |
| DSAG12_03638 | hypothetical protein | -- | 249 | |
| DSAG12_03639 | hypothetical protein | -- | 174 | |
| DSAG12_03640 | 3-dehydrosphinganine reductase | K04708 | 72 | |
| DSAG12_03641 | hypothetical protein | -- | 159 | |
| DSAG12_03642 | charged multivesicular body protein 1 | K12197 | 188 | |

| | | | | |
|---|---|---|---|---|
| DSAG12_03643 | oligoendopeptidase F | K08602 | 231 | |
| DSAG12_03644 | charged multivesicular body protein 1 | K12197 | 251 | |
| DSAG12_03645 | two-component system, NtrC family, sensor kinase | K02482 | 181 | |
| DSAG12_03646 | hypothetical protein | -- | 111 | |
| DSAG12_03647 | hypothetical protein | -- | 223 | |
| DSAG12_03648 | hypothetical protein | -- | 257 | |
| DSAG12_03649 | hypothetical protein | -- | 232 | |
| DSAG12_03650 | hypothetical protein | -- | 221 | |
| DSAG12_03651 | F420-non-reducing hydrogenase iron-sulfur subunit | K14127 | 133 | |
| DSAG12_03652 | heterodisulfide reductase subunit A | K03388 | 159 | |
| DSAG12_03653 | hypothetical protein | -- | 179 | |
| DSAG12_03654 | heterodisulfide reductase subunit A | K03388 | 237 | |
| DSAG12_03655 | heterodisulfide reductase subunit A | K03388 | 135 | |
| DSAG12_03656 | heterodisulfide reductase subunit B | K03389 | 119 | |
| DSAG12_03657 | heterodisulfide reductase subunit C | K03390 | 106 | |
| DSAG12_03658 | 2-iminobutanoate/2-iminopropanoate deaminase | K09022 | 200 | |
| DSAG12_03659 | hypothetical protein | -- | 96 | |
| DSAG12_03660 | thiamine biosynthesis protein ThiI | K03151 | 243 | |
| DSAG12_03661 | ribokinase | K00852 | 197 | |
| DSAG12_03662 | hypothetical protein | -- | 178 | |
| DSAG12_03663 | ATP-binding cassette, sub-family E, member 1 | K06174 | 129 | |
| DSAG12_03664 | hypothetical protein | -- | 191 | |
| DSAG12_03665 | hypothetical protein | -- | 33 | |
| DSAG12_03666 | hypothetical protein | -- | 152 | |
| DSAG12_03667 | 2,3-bisphosphoglycerate-dependent phosphoglycerate mutase | K01834 | 241 | |
| DSAG12_03668 | | K06900 | 175 | |
| DSAG12_03669 | hypothetical protein | -- | 86 | |
| DSAG12_03670 | hypothetical protein | -- | 243 | |
| DSAG12_03671 | glutamine synthetase | K01915 | 232 | |
| DSAG12_03672 | glucosamine--fructose-6-phosphate aminotransferase (isomerizing) | K00820 | 9 | |
| DSAG12_03673 | hypothetical protein | -- | 501 | |
| DSAG12_03674 | COP9 signalosome complex subunit 5 | K09613 | 52 | |
| DSAG12_03675 | tRNA-dihydrouridine synthase B | K05540 | 95 | |
| DSAG12_03676 | starch phosphorylase | K00688 | 161 | |
| DSAG12_03677 | hypothetical protein | -- | 216 | |
| DSAG12_03678 | DNA repair protein RadA | K04483 | 136 | |
| DSAG12_03679 | | K00540 | 65 | |
| DSAG12_03680 | dimethylargininase | K01482 | 241 | |
| DSAG12_03681 | hypothetical protein | -- | 116 | |
| DSAG12_03682 | hypothetical protein | -- | 196 | |
| DSAG12_03683 | | K07131 | 204 | |
| DSAG12_03684 | glucose-6-phosphate isomerase | K01810 | 0 | |
| DSAG12_03685 | hypothetical protein | -- | 187 | |
| DSAG12_03686 | acetyltransferase | K09181 | 0 | |
| DSAG12_03687 | | K06889 | 122 | |
| DSAG12_03688 | hypothetical protein | -- | 239 | |
| DSAG12_03689 | hypothetical protein | -- | 139 | |
| DSAG12_03690 | lipid A 4'-phosphatase | K12978 | 153 | |
| DSAG12_03691 | hypothetical protein | -- | 155 | |
| DSAG12_03692 | hypothetical protein | -- | 0 | |
| DSAG12_03693 | putative hydrolase of the HAD superfamily | K07025 | 168 | |
| DSAG12_03694 | hypothetical protein | -- | 181 | |
| DSAG12_03695 | hypothetical protein | -- | 136 | |
| DSAG12_03696 | hypothetical protein | -- | 197 | EAP30 domain protein (ESCRT-II) (Vps22/36-like) |
| DSAG12_03697 | hypothetical protein | -- | 185 | |
| DSAG12_03698 | hypothetical protein | -- | 256 | Vps4-like ATPase |
| DSAG12_03699 | hypothetical protein | -- | 182 | |
| DSAG12_03700 | hypothetical protein | -- | 113 | |
| DSAG12_03701 | hypothetical protein | -- | 141 | |
| DSAG12_03702 | NTE family protein | K07001 | 93 | |
| DSAG12_03703 | hypothetical protein | -- | 178 | |
| DSAG12_03704 | hypothetical protein | -- | 45 | |
| DSAG12_03705 | hypothetical protein | -- | 174 | |
| DSAG12_03706 | hypothetical protein | -- | 0 | |
| DSAG12_03707 | hypothetical protein | -- | 157 | |
| DSAG12_03708 | hypothetical protein | -- | 202 | |
| DSAG12_03709 | hypothetical protein | -- | 173 | |
| DSAG12_03710 | hypothetical protein | -- | 235 | III) |
| DSAG12_03711 | hypothetical protein | -- | 232 | |
| DSAG12_03712 | hypothetical protein | -- | 244 | |
| DSAG12_03713 | hypothetical protein | -- | 191 | |
| DSAG12_03714 | translation initiation factor 4G | K03260 | 549 | |
| DSAG12_03715 | hypothetical protein | -- | 229 | |
| DSAG12_03716 | hypothetical protein | -- | 213 | |
| DSAG12_03717 | hypothetical protein | -- | 215 | |
| DSAG12_03718 | hypothetical protein | -- | 193 | |
| DSAG12_03719 | hypothetical protein | -- | 223 | |

| | | | |
|---|---|---|---|
| DSAG12_03720 | hypothetical protein | -- | 277 |
| DSAG12_03721 | hypothetical protein | -- | 198 |
| DSAG12_03722 | hypothetical protein | -- | 231 |
| DSAG12_03723 | hypothetical protein | -- | 244 |
| DSAG12_03724 | alpha-ribazole phosphatase | K02226 | 210 |
| DSAG12_03725 | hypothetical protein | -- | 205 |
| DSAG12_03726 | hypothetical protein | -- | 151 |
| DSAG12_03727 | hypothetical protein | -- | 77 |
| DSAG12_03728 | hypothetical protein | -- | 179 |
| DSAG12_03729 | hypothetical protein | -- | 228 |
| DSAG12_03730 | tyrosyl-tRNA synthetase | K01866 | 160 |
| DSAG12_03731 | hypothetical protein | -- | 214 |
| DSAG12_03732 | ABC-2 type transport system ATP-binding protein | K01990 | 208 |
| DSAG12_03733 | pyruvate, orthophosphate dikinase | K01006 | 205 |
| DSAG12_03734 | Ras-related protein Rab-11A | K07904 | 284 |
| DSAG12_03735 | hypothetical protein | -- | 175 |
| DSAG12_03736 | ribonucleoside-triphosphate reductase | K00527 | 227 |
| DSAG12_03737 | aspartate carbamoyltransferase catalytic subunit | K00609 | 146 |
| DSAG12_03738 | hypothetical protein | -- | 128 |
| DSAG12_03739 | large subunit ribosomal protein L44e | K02929 | 239 |
| DSAG12_03740 | small subunit ribosomal protein S27e | K02978 | 191 |
| DSAG12_03741 | transcription elongation factor | K03057 | 164 |
| DSAG12_03742 | hypothetical protein | -- | 172 |
| DSAG12_03743 | proliferating cell nuclear antigen | K04802 | 206 |
| DSAG12_03744 | pyruvate, water dikinase | K01007 | 304 |
| DSAG12_03745 | hypothetical protein | -- | 208 |
| DSAG12_03746 | high-affinity iron transporter | K07243 | 290 |
| DSAG12_03747 | hypothetical protein | -- | 192 |
| DSAG12_03748 | hypothetical protein | -- | 196 |
| DSAG12_03749 | hypothetical protein | -- | 195 |
| DSAG12_03750 | DNA primase large subunit | K18882 | 160 |
| DSAG12_03751 | DNA primase small subunit | K02683 | 460 |
| DSAG12_03752 | hypothetical protein | -- | 231 |
| DSAG12_03753 | H/ACA ribonucleoprotein complex subunit 3 | K11130 | 168 |
| DSAG12_03754 | translation initiation factor 2 subunit 1 | K03237 | 195 |
| DSAG12_03755 | hypothetical protein | -- | 201 |
| DSAG12_03756 | hypothetical protein | -- | 244 |
| DSAG12_03757 | hypothetical protein | -- | 213 |
| DSAG12_03758 | Lrp/AsnC family transcriptional regulator, regulator for asnA, asnC and gidA | K03718 | 187 |
| DSAG12_03759 | cation:H+ antiporter | K07301 | 182 |
| DSAG12_03760 | hypothetical protein | -- | 234 |
| DSAG12_03761 | adenylylsulfate reductase, subunit B | K00395 | 154 |
| DSAG12_03762 | hypothetical protein | -- | 155 |
| DSAG12_03763 | arsenite transporter, ACR3 family | K03325 | 152 |
| DSAG12_03764 | hypothetical protein | -- | 209 |
| DSAG12_03765 | protein-S-isoprenylcysteine O-methyltransferase | K00587 | 171 |
| DSAG12_03766 | putative protease | K08303 | 141 |
| DSAG12_03767 | hypothetical protein | -- | 195 |
| DSAG12_03768 | phosphoglycolate phosphatase | K01091 | 253 |
| DSAG12_03769 | MFS transporter, DHA3 family, tetracycline resistance protein | K18214 | 193 |
| DSAG12_03770 | hypothetical protein | -- | 208 |
| DSAG12_03771 | hypothetical protein | -- | 138 |
| DSAG12_03772 | hypothetical protein | -- | 258 |
| DSAG12_03773 | hypothetical protein | -- | 212 |
| DSAG12_03774 | HSP20 family protein | K13993 | 173 |
| DSAG12_03775 | hypothetical protein | -- | 186 |
| DSAG12_03776 | hypothetical protein | -- | 211 |
| DSAG12_03777 | hypothetical protein | -- | 232 |
| DSAG12_03778 | hypothetical protein | -- | 149 |
| DSAG12_03779 | hypothetical protein | -- | 163 |
| DSAG12_03780 | hypothetical protein | -- | 142 |
| DSAG12_03781 | hypothetical protein | -- | 229 |
| DSAG12_03782 | Arf/Sar family, other | K07977 | 222 |
| DSAG12_03783 | hypothetical protein | -- | 102 |
| DSAG12_03784 | ADP-ribose pyrophosphatase | K01515 | 125 |
| DSAG12_03785 | | K07076 | 212 |
| DSAG12_03786 | | K07076 | 188 |
| DSAG12_03787 | hypothetical protein | -- | 194 |
| DSAG12_03788 | glycerol-3-phosphate dehydrogenase subunit B | K00112 | 180 |
| DSAG12_03789 | glycerol-3-phosphate dehydrogenase | K00111 | 236 |
| DSAG12_03790 | hypothetical protein | -- | 102 |
| DSAG12_03791 | sulfopyruvate decarboxylase subunit beta | K13039 | 176 |
| DSAG12_03792 | phosphonopyruvate decarboxylase | K09459 | 146 |
| DSAG12_03793 | phosphinothricin acetyltransferase | K03823 | 216 |
| DSAG12_03794 | hypothetical protein | -- | 218 |
| DSAG12_03795 | carboxy-terminal domain RNA polymerase II polypeptide A small phosphatase | K15731 | 233 |
| DSAG12_03796 | hypothetical protein | -- | 204 |
| DSAG12_03797 | hypothetical protein | -- | 143 |

| | | | |
|---|---|---|---|
| DSAG12_03798 | MFS transporter, DHA1 family, multidrug resistance protein | K08153 | 226 |
| DSAG12_03799 | thiamine pyrophosphokinase | K00949 | 169 |
| DSAG12_03800 | hypothetical protein | -- | 210 |
| DSAG12_03801 | thiamine transport system substrate-binding protein | K02064 | 285 |
| DSAG12_03802 | thiamine transport system permease protein | K02063 | 170 |
| DSAG12_03803 | iron(III) transport system ATP-binding protein | K02010 | 122 |
| DSAG12_03804 | hypothetical protein | -- | 145 |
| DSAG12_03805 | tRNA-Arg | Intron(43144 | 246 |
| DSAG12_03806 | hypothetical protein | -- | 199 |
| DSAG12_03807 | hypothetical protein | -- | 73 |
| DSAG12_03808 | small subunit ribosomal protein S7 | K02992 | 69 |
| DSAG12_03809 | small subunit ribosomal protein S12 | K02950 | 179 |
| DSAG12_03810 | large subunit ribosomal protein L30e | K02908 | 115 |
| DSAG12_03811 | hypothetical protein | -- | 183 |
| DSAG12_03812 | hypothetical protein | -- | 224 |
| DSAG12_03813 | hypothetical protein | -- | 129 |
| DSAG12_03814 | hypothetical protein | -- | 229 |
| DSAG12_03815 | hypothetical protein | -- | 154 |
| DSAG12_03816 | tryptophanyl-tRNA synthetase | K01867 | 157 |
| DSAG12_03817 | hypothetical protein | -- | 194 |
| DSAG12_03818 | hypothetical protein | -- | 180 |
| DSAG12_03819 | DNA-directed RNA polymerase subunit A" | K03042 | 207 |
| DSAG12_03820 | DNA-directed RNA polymerase subunit A' | K03041 | 113 |
| DSAG12_03821 | hypothetical protein | -- | 32 |
| DSAG12_03822 | hypothetical protein | -- | 242 |
| DSAG12_03823 | D-3-phosphoglycerate dehydrogenase | K00058 | 13 |
| DSAG12_03824 | hypothetical protein | -- | 3 |
| DSAG12_03825 | hypothetical protein | -- | 3 |
| DSAG12_03826 | acetyl-CoA synthetase (ADP-forming) | K01905 | 111 |
| DSAG12_03827 | hypothetical protein | -- | 154 |
| DSAG12_03828 | Ca2+-transporting ATPase | K01537 | 148 |
| DSAG12_03829 | nicotinamide-nucleotide amidase | K03742 | 108 |
| DSAG12_03830 | hypothetical protein | -- | 173 |
| DSAG12_03831 | hypothetical protein | -- | 203 |
| DSAG12_03832 | two-component system, NtrC family, C4-dicarboxylate transport sensor histidine kinase DctB | K10125 | 105 |
| DSAG12_03833 | acetyl-CoA C-acetyltransferase | K00626 | 91 |
| DSAG12_03834 | DNA polymerase I | K02319 | 221 |
| DSAG12_03835 | hypothetical protein | -- | 240 |
| DSAG12_03836 | tRNA-Thr | Intron(43643 | 177 |
| DSAG12_03837 | hypothetical protein | -- | 203 |
| DSAG12_03838 | hypothetical protein | -- | 203 |
| DSAG12_03839 | hypothetical protein | -- | 243 |
| DSAG12_03840 | hypothetical protein | -- | 0 |
| DSAG12_03841 | ABC-2 type transport system permease protein | K01992 | 151 |
| DSAG12_03842 | ABC-2 type transport system ATP-binding protein | K01990 | 56 |
| DSAG12_03843 | hypothetical protein | -- | 268 |
| DSAG12_03844 | hypothetical protein | -- | 277 |
| DSAG12_03845 | hypothetical protein | -- | 450 |
| DSAG12_03846 | hypothetical protein | -- | 297 |
| DSAG12_03847 | hypothetical protein | -- | 0 |
| DSAG12_03848 | glucans biosynthesis protein C | K11941 | 245 |
| DSAG12_03849 | phenylacetate-CoA ligase | K01912 | 157 |
| DSAG12_03850 | hypothetical protein | -- | 258 |
| DSAG12_03851 | ribosomal-protein-alanine N-acetyltransferase | K03790 | 0 |
| DSAG12_03852 | hypothetical protein | -- | 165 |
| DSAG12_03853 | hypothetical protein | -- | 0 |
| DSAG12_03854 | hypothetical protein | -- | 186 |
| DSAG12_03855 | hypothetical protein | -- | 838 |
| DSAG12_03856 | hypothetical protein | -- | 165 |
| DSAG12_03857 | MFS transporter, DHA3 family, macrolide efflux protein | K08217 | 116 |
| DSAG12_03858 | aminopeptidase | K01269 | 121 |
| DSAG12_03859 | Fur family transcriptional regulator, peroxide stress response regulator | K09825 | 105 |
| DSAG12_03860 | catalase-peroxidase | K03782 | 221 |
| DSAG12_03861 | hypothetical protein | -- | 258 |
| DSAG12_03862 | hypothetical protein | -- | 0 |
| DSAG12_03863 | hypothetical protein | -- | 99 |
| DSAG12_03864 | hypothetical protein | -- | 164 |
| DSAG12_03865 | hypothetical protein | -- | 178 |
| DSAG12_03866 | hydrogenase expression/formation protein HypD | K04654 | 165 |
| DSAG12_03867 | hydrogenase 3 maturation protease | K08315 | 172 |
| DSAG12_03868 | hydrogenase expression/formation protein HypE | K04655 | 0 |
| DSAG12_03869 | hydrogenase maturation protein HypF | K04656 | 133 |
| DSAG12_03870 | lysophospholipase | K01048 | 157 |
| DSAG12_03871 | putative hydrolase of the HAD superfamily | K07025 | 187 |
| DSAG12_03872 | hypothetical protein | -- | 196 |
| DSAG12_03873 | hydrogenase expression/formation protein HypC | K04653 | 195 |
| DSAG12_03874 | ArsR family transcriptional regulator | K03892 | 94 |
| DSAG12_03875 | Cd2+/Zn2+-exporting ATPase | K01534 | 212 |

| | | | | |
|---|---|---|---|---|
| DSAG12_03876 | hypothetical protein | -- | 224 | |
| DSAG12_03877 | F420-non-reducing hydrogenase large subunit | K14126 | 187 | |
| DSAG12_03878 | heterodisulfide reductase subunit B | K03389 | 163 | hypothetical protein with RING-/ and TPR domain |
| DSAG12_03879 | heterodisulfide reductase subunit C | K03390 | 286 | |
| DSAG12_03880 | F420-non-reducing hydrogenase small subunit | K14128 | 229 | |
| DSAG12_03881 | hypothetical protein | -- | 230 | |
| DSAG12_03882 | tRNA 2-thiouridine synthesizing protein C | K07236 | 190 | |
| DSAG12_03883 | tRNA 2-thiouridine synthesizing protein D | K07235 | 181 | |
| DSAG12_03884 | | K07112 | 5 | |
| DSAG12_03885 | tRNA 2-thiouridine synthesizing protein A | K04085 | 8 | |
| DSAG12_03886 | hypothetical protein | -- | 6 | |
| DSAG12_03887 | mono-ADP-ribosyltransferase sirtuin 6 | K11416 | 8 | |
| DSAG12_03888 | hypothetical protein | -- | 4 | |
| DSAG12_03889 | anaerobic magnesium-protoporphyrin IX monomethyl ester cyclase | K04034 | 244 | |
| DSAG12_03890 | hypothetical protein | -- | 301 | |
| DSAG12_03891 | UDP-glucose 4-epimerase | K01784 | 138 | |
| DSAG12_03892 | aminoglycoside N3'-acetyltransferase | K00662 | 232 | |
| DSAG12_03893 | hypothetical protein | -- | 269 | |
| DSAG12_03894 | hypothetical protein | -- | 0 | |
| DSAG12_03895 | hypothetical protein | -- | 178 | |
| DSAG12_03896 | hypothetical protein | -- | 144 | |
| DSAG12_03897 | hypothetical protein | -- | 179 | |
| DSAG12_03898 | titin | K12567 | 113 | |
| DSAG12_03899 | | | 166 | |
| DSAG12_03900 | | | 197 | |
| DSAG12_03901 | | | 217 | small GTP-binding domain protein |
| DSAG12_03902 | | | 268 | |
| DSAG12_03903 | | | 163 | |
| DSAG12_03904 | | | 216 | |
| DSAG12_03905 | | | 104 | |
| DSAG12_03906 | | | 156 | |
| DSAG12_03907 | | | 340 | |
| DSAG12_03908 | | | 187 | |
| DSAG12_03909 | | | 233 | |
| DSAG12_03910 | | | 83 | |
| DSAG12_03911 | | | 117 | |
| DSAG12_03912 | | | 226 | |
| DSAG12_03913 | | | 205 | |
| DSAG12_03914 | | | 177 | |
| DSAG12_03915 | | | 171 | |
| DSAG12_03916 | | | 114 | |
| DSAG12_03917 | | | 137 | |
| DSAG12_03918 | | | 175 | |
| DSAG12_03919 | | | 218 | |
| DSAG12_03920 | | | 163 | |
| DSAG12_03921 | | | 178 | |
| DSAG12_03922 | | | 253 | |
| DSAG12_03923 | | | 200 | |
| DSAG12_03924 | | | 173 | |
| DSAG12_03925 | | | 79 | |
| DSAG12_03926 | | | 210 | |
| DSAG12_03927 | | | 225 | |
| DSAG12_03928 | | | 217 | |
| DSAG12_03929 | | | 200 | |
| DSAG12_03930 | | | 204 | |
| DSAG12_03931 | | | 232 | |
| DSAG12_03932 | | | 126 | |
| DSAG12_03933 | | | 323 | |
| DSAG12_03934 | | | 180 | |
| DSAG12_03935 | | | 133 | |
| DSAG12_03936 | | | 178 | |
| DSAG12_03937 | | | 221 | |
| DSAG12_03938 | | | 160 | |
| DSAG12_03939 | | | 243 | |
| DSAG12_03940 | | | 172 | |
| DSAG12_03941 | | | 233 | |
| DSAG12_03942 | | | 248 | |
| DSAG12_03943 | | | 229 | |
| DSAG12_03944 | | | 171 | |
| DSAG12_03945 | | | 158 | |
| DSAG12_03946 | | | 64 | |
| DSAG12_03947 | | | 96 | |
| DSAG12_03948 | | | 224 | |
| DSAG12_03949 | | | 290 | |
| DSAG12_03950 | | | 285 | |
| DSAG12_03951 | | | 248 | |
| DSAG12_03952 | | | 273 | |
| DSAG12_03953 | | | 266 | |
| DSAG12_03954 | | | 63 | |
| DSAG12_03955 | | | 155 | |

| | |
|---|---|
| DSAG12_03956 | 129 |
| DSAG12_03957 | 217 |
| DSAG12_03958 | 57 |
| DSAG12_03959 | 174 |
| DSAG12_03960 | 73 |
| DSAG12_03961 | 60 |
| DSAG12_03962 | 327 |
| DSAG12_03963 | 179 |
| DSAG12_03964 | 157 |
| DSAG12_03965 | 250 |
| DSAG12_03966 | 175 |
| DSAG12_03967 | 159 |
| DSAG12_03968 | 203 |
| DSAG12_03969 | 335 |
| DSAG12_03970 | 126 |
| DSAG12_03971 | 224 |
